# Peptidic Tryptophan Halogenation by a Promiscuous Flavin-dependent Enzyme

**DOI:** 10.1101/2025.01.12.632611

**Authors:** Andrew J. Rice, Mayuresh G. Gadgil, Paola Bisignano, Richard A Stein, Hassane S. Mchaourab, Douglas A. Mitchell

**Affiliations:** Department of Biochemistry, Vanderbilt University School of Medicine, Department of Chemistry, Vanderbilt University, Nashville, TN 37232 USA; Department of Chemistry, University of Illinois Urbana-Champaign, Urbana, IL 61801 USA; Department of Molecular Physiology and Biophysics, Vanderbilt University, Nashville, TN 37232 USA

**Keywords:** biocatalysis, halogenation, peptides, proteins, chlorine

## Abstract

Amino acids undergo numerous enzymatic modifications. However, the broad applicability of amino acid-modifying enzymes for synthetic purposes is limited by narrow substrate scope and often unknown regulatory or accessory factor requirements. Here, we characterize ChlH, a flavin-dependent halogenase (FDH) from the chlorolassin biosynthetic gene cluster. Unlike characterized peptide-modifying FDHs, which are limited to either specifically modified peptides or the termini of linear peptides, ChlH halogenates internal Trp residues of linear peptides, as well as N- and C-terminal Trp. Scanning mutagenesis of the substrate peptide ChlA revealed Trp was tolerated by ChlH at nearly every position. Molecular dynamics simulations corroborated the importance of a C-terminal motif in ChlA and provided insight into the lack of Trp14 chlorination in native chlorolassin. Furthermore, halogenation of disparate ribosomally synthesized and post-translationally modified peptide (RiPP) precursor peptides, pharmacologically relevant peptides, and an internal Trp of a protein was achieved using wild-type ChlH. A rapid cell-free biosynthetic assay provided insight into ChlH’s preferences. In contrast to characterized FDHs, ChlH halogenates diverse peptide sequences, and we predict this promiscuity may find utility in the modification of additional peptide and protein substrates of biotechnological value.

**Entry for the Table of Contents:** ChlH, a flavin-dependent tryptophan halogenase, is reconstituted *in vitro* and found to be capable of modifying a wide array of diverse peptidic substrates despite showing selectivity on its native substrate peptide ChlA. This highlights its potential use in the biocatalytic production of chlorinated peptides.

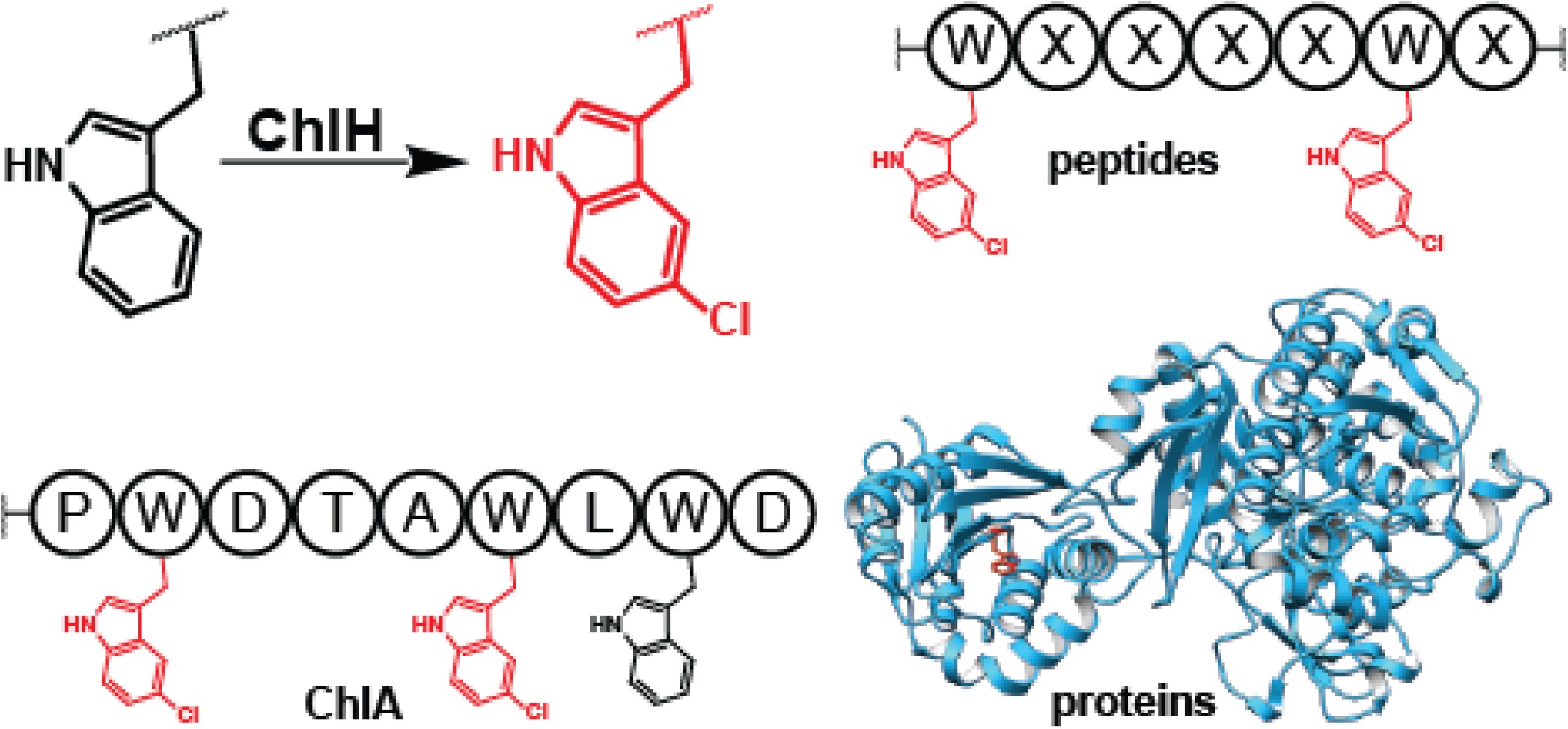

## Introduction

Natural products have historically been a source of inspiration for drug development.^1^ However, most natural products must be further derivatized to become suitable for clinical use. One common modification is halogenation. Many FDA-approved drugs contain at least a single halogen atom,^2^ and numerous research groups have devised creative methods for robust enzymatic halogen incorporation into small-molecule scaffolds.^3–9^

Natural product biosynthetic pathways similarly employ various strategies for bioactivity optimization. Many flavin-dependent halogenases (FDHs) act on the free amino acid Trp, and play a critical role in modifying bioactivity.^10–13^ Unfortunately, FDHs frequently exhibit poor activity *in vitro* and require additional recognition motifs for robust enzymatic activity.^14–16^ Recently, approaches have been taken to obtain FDHs that halogenate peptidic Trp.^17–19^ Several recently characterized FDHs are derived from ribosomally synthesized and post-translationally modified peptide (RiPP) biosynthetic gene clusters.

While relatively rare in RiPP biosynthesis, examples of Trp halogenation have been reported. NAI-107 (synonym: microbisporicin) is a 24-residue lanthipeptide with Trp4 being chlorinated at the 5-position by the FDH MibH (NCBI accession code: WP_036325874.1).^20^ However, MibH only accepts the lanthionine-containing, multicyclic peptide *deschloro*-NAI-107 as a substrate. This feature disqualifies wild-type MibH as a broadly applicable biocatalyst, as it does not tolerate linear substrates or lanthipeptides structurally similar to NAI-107. Another RiPP-associated FDH, SrpI, forms 6-bromoTrp at the C-terminus of the cognate substrate peptide.^21,22^ Substrate tolerance studies showed that SrpI-catalyzed bromination is limited to the C-terminus and required an ∼18-residue segment of the leader peptide for efficient product formation, narrowing its potential biocatalytic utility.^23^ Separately, Thal (ABK79936.1), which modifies free *L*-Trp during the biosynthesis of thienodolin,^24^ can also site-selectively chlorinate and brominate polypeptides, but is restricted to C-terminal Trp. An evolved Thal could also selectively brominate Trp when placed at the C-terminus of a protein.^25^

Chlorolassin is a lasso peptide isolated from *Lentzia jiangxiensis*.^26^ The mature product features ChlH-dependent 5-chlorination of Trp8 and Trp12 (Figure 1). Here, we present the *in vitro* reconstitution of the chlorolassin halogenase ChlH (WP_090100303.1) and the cognate flavin reductase ChlR (WP_090100388.1). ChlH modifies the linear precursor peptide ChlA (WP_143022795.1) and does not modify threaded *des*-chlorolassin (chlorolassin lacking halogenation). In contrast to the stricter substrate requirements of MibH and SrpI, ChlH exhibited broad substrate tolerance and modified a wide range of ChlA variants. Furthermore, ChlH chlorinates Trp in unrelated peptides of various lengths, a macrocyclic peptide, and an internal residue of an unrelated 1,054-amino acid protein. Our molecular dynamics simulations provide valuable insights into the dynamics of an enzyme-substrate interaction. We glean insight into the substrate preferences of ChlH by extensively evaluating numerous substrates using either purified proteins or cell-free biosynthetic methods. These insights are then utilized to significantly improve processing of a poor substrate. This work demonstrates that wild-type ChlH is a versatile Trp chlorinating enzyme.

**Figure 1.**
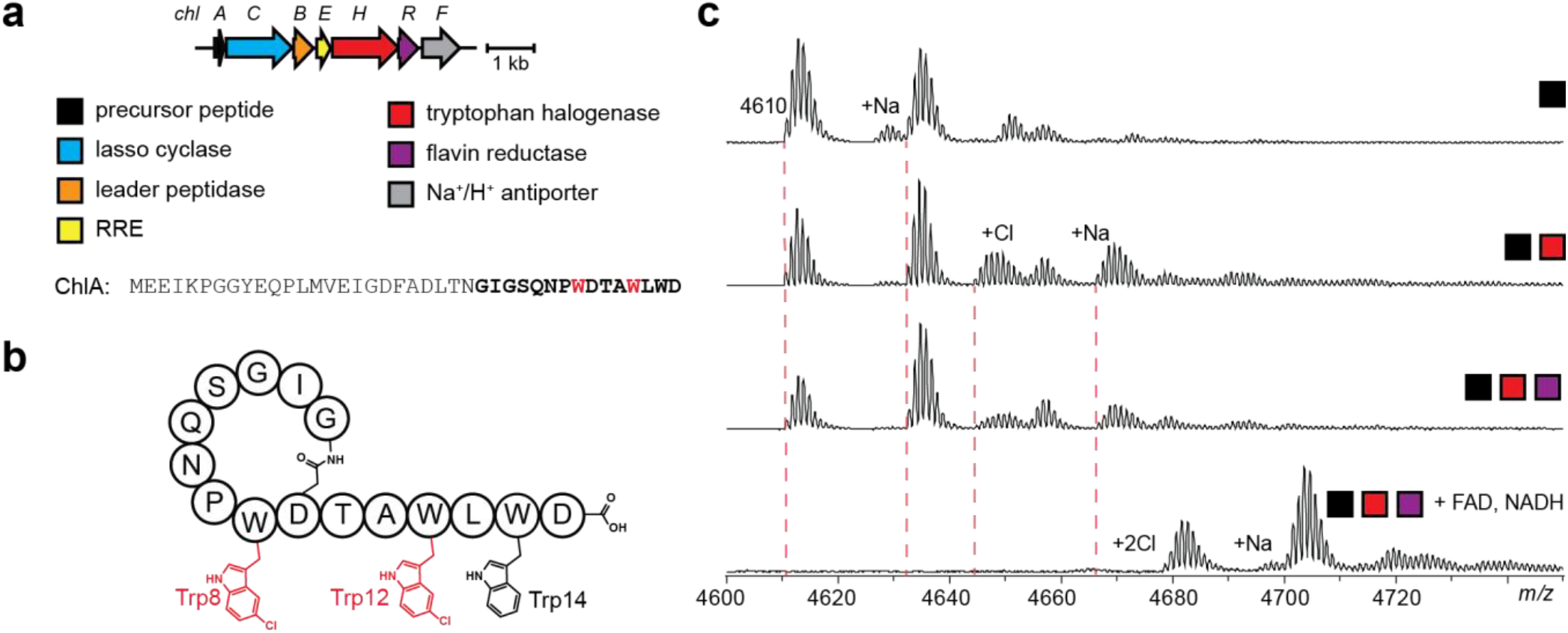
Chlorolassin biosynthesis, structure, and reconstitution. (a) Biosynthetic gene cluster and precursor peptide sequence for ChlA. (b) The structure of chlorolassin shown in an unthreaded conformation for clarity. Trp8 and Trp12 are chlorinated at indole position 5 (Trp14 is unmodified). (c) *In vitro* reconstitution of ChlH/R. Expected m/z values for unmodified, mono-, and dichlorinated ChlA are 4610, 4644, and 4678 respectively. Colored squares represent reaction components as defined in panel a.

## Results and Discussion

### *In vitro* reconstitution of ChlHR

We first sought to determine when Trp8 and Trp12 were halogenated during chlorolassin biosynthesis. With Trp14 being less accessible in the folded lasso peptide (PDB code: 8UKC),^26^ and strict substrate requirements reported for NAI-107 chlorination, we hypothesized that ChlH modified *des*-chlorolassin (folded lasso peptide). However, given the observation of *des*-chlorolassin during the isolation of chlorolassin, lasso peptide cyclization was independent of the chlorination status of ChlA.^26^ To address the timing of ChlH modification, we reconstituted the chlorination activity *in vitro. E. coli*-optimized genes encoding ChlH and ChlR (see Supporting Methods) were individually cloned with N-terminal His^6^ tags. Following co-expression with the chaperones GroES/EL (pGro7),^14^ ChlH and ChlR were purified in high yield using immobilized metal-affinity chromatography (Figure S1). Reactions were initiated using purified ChlH/R and an excess quantity of NADH and FAD before analysis by matrix-assisted laser desorption/ionization time-of-flight mass spectrometry (MALDI-TOF-MS). Under no circumstances was des-chlorolassin or free L-Trp chlorinated, even after prolonged reaction times and relatively high enzyme concentrations. No chlorination of *des*-chlorolassin and the free amino acid *L*-Trp was observed after reaction with ChlH/R (Figures S2-3).

Next, we prepared a construct encoding the ChlA precursor peptide with an N-terminal maltose-binding protein (MBP) tag. Removal of MBP via treatment with tobacco etch virus (TEV) protease resulted in linear ChlA peptide. Upon reaction with ChlH/R, NADH, FAD, and Cl^-^, two chlorinations were observed on ChlA within 2 h (Figure 1). Although ChlH alone partially chlorinated ChlA, the addition of excess FAD, NADH, and ChlR yielded full dichlorination. All subsequent reactions therefore included ChlR, FAD, and NADH. MALDI-LIFT-TOF/TOF-MS (hereafter, MALDI-LIFT-MS) analysis established Trp8 and Trp12 as the chlorination sites, in agreement with the site-selectivity observed in native chlorolassin (Figures 1, S4). Analysis of monochlorinated ChlA from earlier timepoints revealed that Trp12 was modified before Trp8 (Figure S5).

After extended reaction times (24 h), a low intensity ion indicative of three ChlA chlorinations was detected (Figure S6). Localization by MALDI-LIFT-MS revealed Trp14 as the third chlorination site (Figure S7).^27^ Previous work on the FDH RebH (CAC93722.1) identified Lys139 of ChlH as a potential catalytic residue (Figure S8). Characterized FDHs use chloride, O_2_, and FADH_2_ to generate hypochlorous acid. A conserved catalytic Lys then interacts with hypochlorous acid either through chloramine formation or by serving as a proton donor to chlorinate Trp.^10,15^ A previous report showed that hypohalous acid can also spontaneously react with Trp.^28^ To determine if the conserved Lys139 of ChlH was necessary for Trp chlorination, we prepared the K139A variant (Figure S1). Identical reactions were conducted and MS-based analysis showed that the K139A variant was unable to chlorinate ChlA, even after extended reaction times (Figure S9). This confirmed that Trp14 chlorination, while only observed *in vitro*, required catalytically active ChlH.

### ChlH can brominate Trp residues of ChlA

To evaluate if ChlH could utilize other halide sources, a series of reactions were supplemented with 100 mM NaF, NaBr, or NaI (Figure S10). The chloride concentration remained at 50 mM to allow for halide competition. NaF supplementation did not result in ChlA fluorination but rather exclusive chlorination. NaBr supplementation gave a mixture of chlorination and bromination. The stock solutions of ChlA, TEV protease, MBP-ChlA, ChlH, and ChlR were then extensively buffer exchanged to reduce chloride to trace levels. Upon NaBr supplementation, exclusive bromination of ChlA was observed (Figure S11). Unfortunately, ChlH precipitated in the NaI-supplemented buffer and thus iodination was not investigated further.

### ChlA halogenation does not require the leader peptide

To assess the role of the leader peptide in ChlH activity, we cleaved MBP-ChlA at the native leader peptide-core peptide junction using the leader peptidase FusB (WP_011291590.1) and associated RiPP-recognition element FusE (WP_011291591.1) from *Thermobifida fusca*.^29,30^ Despite this non-cognate usage, FusB/E readily yielded the core region of ChlA (residues 1-15, ChlA_core_). The relative chlorination efficiency of full-length ChlA and ChlA_core_ was then compared (Figure S12). ChlA_core_ was fully dichlorinated at 2 h, at the same approximate rate as full-length (40-residue) ChlA. This demonstrated that ChlH activity was independent of the leader peptide, unlike other characterized RiPP-associated Trp-halogenases.^31^ However, we note that ChlA_core_ is poorly soluble in water, thus we opted to use full-length ChlA for most of the *in vitro* experiments hereafter.

### Ala variants of ChlA highlight substrate promiscuity

To explore the site-selective dichlorination of ChlA, we performed an Ala scan of ChlA_core_ (Figure 2). Halogenation activity was assessed *in vitro* at 2 and 24 h by MALDI-TOF-MS (Figures S13-S14).

**Figure 2.**
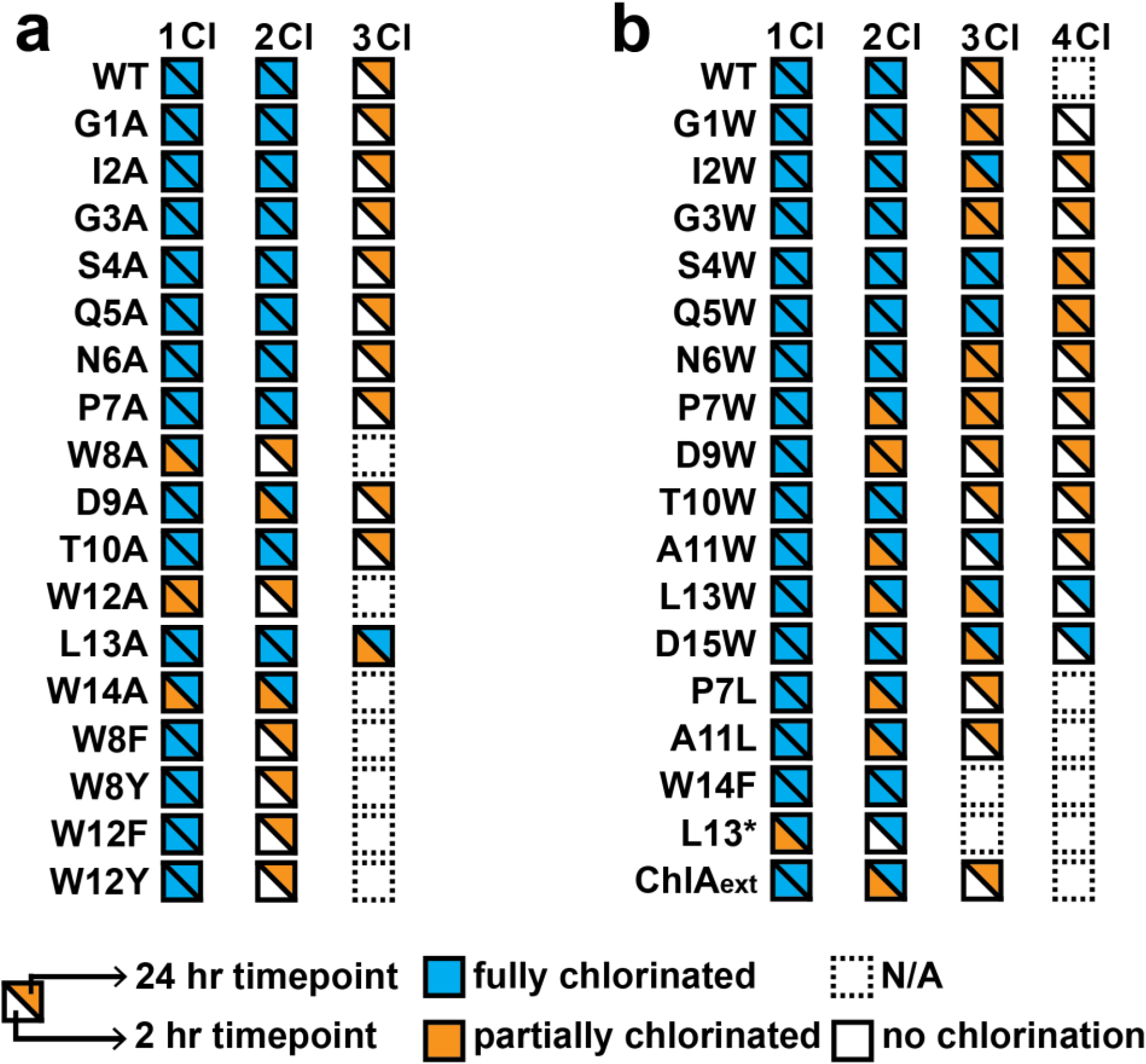
*In vitro* chlorination of ChlA variants. (a) Compiled results from the Ala scan of ChlA and other variants. The lower left half of each square represents chlorination status after 2 h of reaction with ChlH/R. The upper right half represents the same after 24 h. WT = wild-type ChlA. (b) Compiled results from the Trp scan of ChlA with other variants. L13* represents a stop codon at position 13 of ChlA. ChlA_ext_ represents a C-terminally extended variant of ChlA, the sequence of which can be found in Figure S17. N/A, not applicable.

Most Ala substitutions were well-tolerated, with minimal differences observed in processing compared to wild-type ChlA. Single chlorination of the W8A and W12A variants at 2 h demonstrated that chlorination of Trp8 was independent of Trp12 chlorination and *vice versa*. However, we note that neither W8A nor W12A was fully monochlorinated at 2 h. To determine if aromatic amino acids at position 8 or 12 enhanced chlorination efficiency at the non-varied site, we generated and analyzed the W8F, W8Y, W12F, and W12Y variants (Figure 2). After 2 h of reaction with ChlH, each variant was fully monochlorinated (Figure S15).

A modest extent of L13A trichlorination was observed at 2 h and full trichlorination was achieved within 24 h. This result implied Leu13 hindered Trp14 chlorination and could aid in understanding the observed chlorination pattern on chlorolassin. To further probe this idea, we generated ChlA variants P7L and A11L, which place Leu immediately before Trp8 and Trp12, respectively (Figure 2). These variants were poorly modified compared to wild-type ChlA and P7A at 2 h (Figure S14), supporting an inhibitory role of Leu at the (-1) position for Trp chlorination. Similarly, ChlH processing of variant W14A gave a mixture of 0-2 chlorinations at 2 h but was fully dichlorinated at 24 h. Predicting that aromatic amino acids near the C-terminus contribute to efficient substrate recognition, we evaluated W14F, which was fully dichlorinated at 2 h (Figure 2, S15-16). This result underscores the importance of aromatic amino acids near the C-terminus of ChlA for efficient processing, but demonstrates they are dispensable at longer reaction times.

The characterized FDH SrpI recognizes the C-terminal region of SrpE, the cognate precursor peptide.^31^ To assess the importance of the C-terminus of ChlA for ChlH activity, we evaluated a truncated ChlA peptide lacking Leu13-Asp15 (ChlA-L13*, Figure 2). A low-intensity monochlorination product was observed at 2 h while fully dichlorinated product was observed at 24 h (Figures S15-16). Accordingly, a 19-residue addition to the C-terminus of ChlA that included a His^6^ tag (ChlA_ext,_ Figures 2, S17) resulted in complete monochlorination at 2 h and complete dichlorination at 24 h after ChlH processing (Figures S15-S16). These data demonstrate that the identity of the most C-terminal residues of the substrate peptide influences the processing efficiency of ChlH but are otherwise dispensable.

### ChlH chlorinates non-native Trp residues in ChlA

We then performed a Trp scan of the core region of ChlA to assess chlorination at non-native positions (Figures 2, S18-19). ChlA variants S4W and G5W were fully trichlorinated at 2 h while the remaining variants exhibited an array of modification states. The P7W, D9W, A11W, and L13W variants of ChlA were less extensively modified compared to wild-type and all other Trp variants at 2 h. Notably, either partial or full modification was observed for all Trp variants at 24 h.

ChlA variants L13W and D15W were fully tetrachlorinated at 24 h, supporting a role of these positions in delaying Trp14 chlorination in wild-type ChlA. To better gauge the pattern of Trp chlorination, several ChlA Trp variants were reacted with ChlH/R, then FusB/E, purified by high-performance liquid chromatography (HPLC), and analyzed by MALDI-LIFT-MS (Figures S20-S24). This revealed a general trend in which Trp8 and Trp12 are modified more effectively than Trp14 or the non-native Trp residues introduced into ChlA. Overall, these data collectively show that certain positions of the core peptide, when changed to Trp, are preferentially chlorinated compared to other positions. Still, these preferences can be overridden with longer reaction times.

### ChlH recognizes a Trp-Leu motif in ChlA

Given that the leader region of ChlA was dispensable for ChlH processing, we next assessed the manner by which ChlA and ChlH may interact. We began with generating an AlphaFold 2 Multimer (AF2M) model of the ChlH–ChlA complex.^32^ Trp8 was positioned with the C5 at 5.2 Å from the catalytic Lys139 (Figures 3, S25), though not within the distance necessary for catalysis to occur, as it would be too close to include a Chlorine atom. In addition, the model’s confidence in the placement and orientation of the ChlA peptide is very low (Figure S25). Notably, AlphaFold 3 (AF3) does not offer a substantial increase in model quality (Figure S26).^33^ To further explore substrate-enzyme interaction, we first optimized the rotamer of Trp8 in our AF2M model to enhance its proximity and orientation relative to Lys139. Using this as a template, we manually shifted the ChlA peptide to generate additional models in which Trp12 or Trp14 replaced Trp8 in the same spatial location, since AlphaFold did not generate any high-confidence models of Trp12 or Trp14 in the active site. Each of these four configurations (original AF2M model, Trp8-optimized, Trp12-shifted, Trp14-shifted) was simulated in triplicate using unbiased atomistic molecular dynamics, resulting in 12 trajectories (Table S1) and a total of 16.5 μs of aggregate sampling. All MD simulations were performed with full-length ChlA.

**Figure 3.**
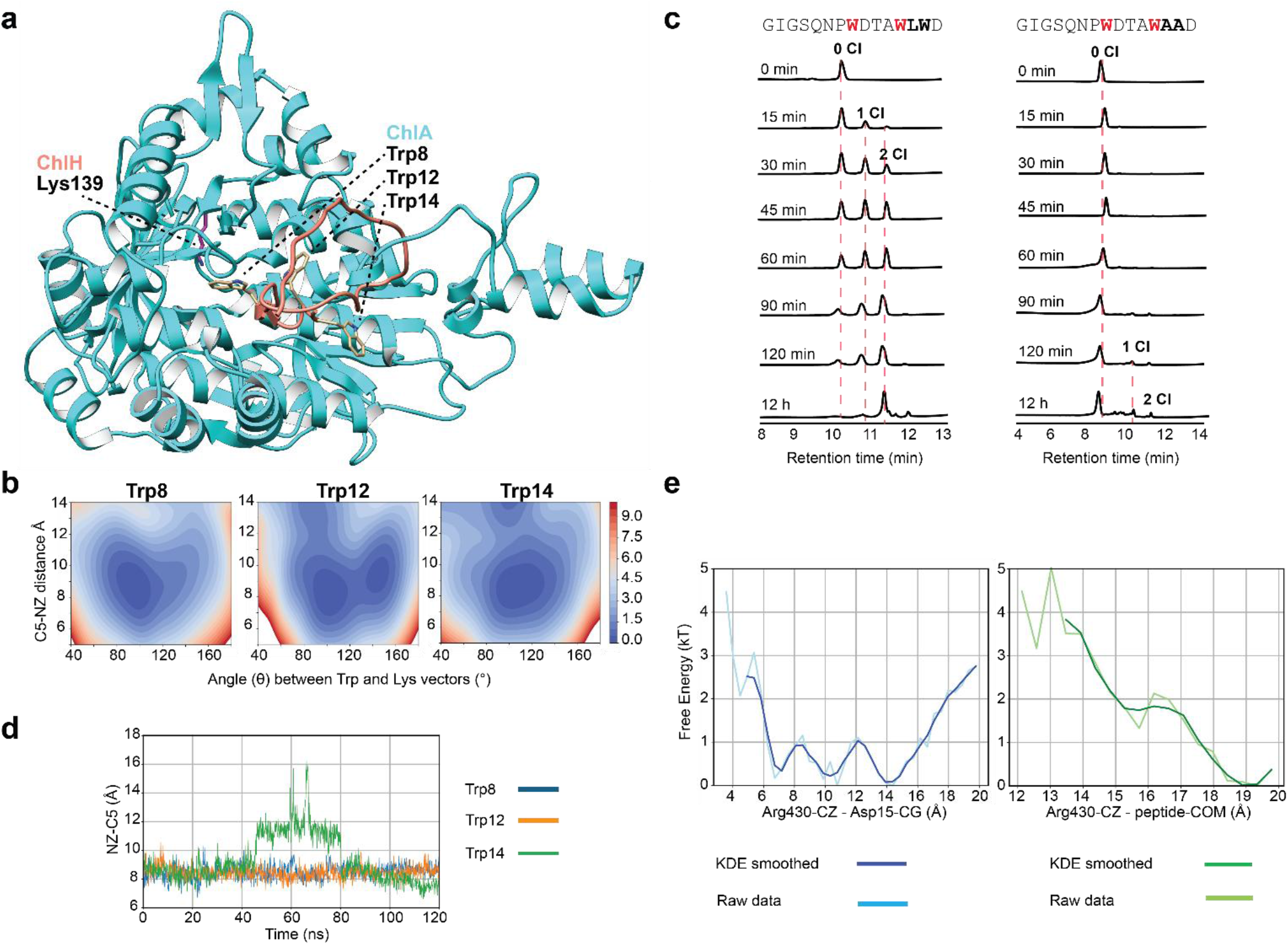
(a) AlphaFold 2 Multimer structure of ChlH and ChlA utilized for MD. Trp8 is placed close to Lys139, albeit with very low confidence. (b) Free energies projected onto the distance between Trp C5 and Lys139 NZ atoms, and the angle (θ) between their respective alignment vectors, for Trp8, Trp12, and Trp14. Lower free energy regions (blue) indicate favorable geometric configurations. The angle θ is defined as the angle between the Trp indole plane vector (CZ3→C5) and the Lys sidechain vector (CE→NZ). (c) Stacked HPLC chromatograms with observation at 220 nm showing the relative reaction rates of ChlA_core_ and ChlA_core(AA)_ with ChlH. (d) Distance over time between the catalytic Lys139 NZ atom and the C5 atom of the Trp residue positioned near the active site in each system: Trp8 (blue), Trp12 (orange), and Trp14 (green). Trajectories were sampled from iteration 12 of the weighted ensemble simulations. (e) One-dimensional free energy profiles projected along the distance between the Arg430 guanidinium carbon (CZ) and either the Asp15 carboxylate (CG) or the peptide center of mass (COM). Raw distributions (light blue/green) and KDE-smoothed profiles (blue/green) are shown.

Across most trajectories, the Trp initially placed near the catalytic lysine disengaged within a few hundred ns. Notably, two Trp8-centered simulations maintained close contact throughout, and in one of these, structural rearrangements in ChlH stabilized the ChlA peptide in a catalytically poised conformation. From this trajectory, we selected three frames (at 730 ns, 1500 ns, and 2500 ns) to serve as seeds for weighted ensemble (WE)^34^ simulations using the highly parallelizable WESTPA 2.0^35^ framework that allow sampling of rare events. For each of the seeds, we generated new configurations by repositioning each Trp into the same catalytic pose, yielding a systematic permutation set for testing Trp retention.

This design allowed us to enrich the initial states for the WE simulations, ensuring each Trp residue was tested under identical structural conditions. The progress coordinates used in the WE framework were: (1) a reaction likelihood score combining Trp–Lys distance and angular alignment (see SI methods), and (2) a discrete Trp identity binning scheme (1, 2, or 3) to track for each trajectory walker which of the three Trp contributed to the score (yielding the best angle/distance parameters). Although the WE simulations were driven by these progress coordinates, we projected the resulting data onto the raw geometric observables, distance and angle, to analyze the underlying free energy landscapes.

WE simulations revealed that both Trp8 and Trp12 repeatedly presented their C5 atom to Lys139, whereas Trp14 failed to maintain proximity. Free energy surfaces projected onto distance and angle before (Figure S27) and after applying Kernel Density Estimation (KDE) smoothing (Figure 3) demonstrated that Trp14 samples higher-energy regions with suboptimal geometry, while Trp8 and Trp12 occupy well-defined low-energy basins consistent with catalytically competent binding. These findings suggest that Trp14 is not effectively retained near the catalytic site and that subtle but critical structural remodeling of the apo state, which is absent in the unrefined AF2M model, may be necessary to permit productive ChlA binding.

To assess our simulations, we performed an additional endpoint assay utilizing both ChlA_core_ and ChlA_core(AA)_, in which both Leu13 and Trp14 were replaced with Ala. HPLC analysis showed that after treatment with ChlH, the conversion was nearly complete for ChlA_core_ while very little conversion was observed for ChlA_core(AA)_, under identical conditions (Figure 3). This further supports the hypothesis that the Leu-Trp motif aids in efficient enzymatic processing. Analysis of representative trajectories from iteration 12 of the WE simulations revealed distinct differences in the behavior of individual Trp residues. Trp8 and Trp12 exhibited persistent interactions with the catalytic Lys139 over time, with the indole C5 remaining within ∼7-10 Å of the Lys NZ, whereas Trp14 dissociated from the catalytic site after ∼40 ns (Supplementary Movies SM1–3, Figure 3). We attribute the rapid dissociation of Trp14 from the active site in part to an interaction with the neighboring Leu13, which caused Trp14 to rotate away from the active site. Additionally, our trajectories revealed a potential compensatory interaction between ChlA Asp15 and ChlH Arg430 in substrate recognition. Free energy profiles were projected onto one-dimensional distances between Arg430–Asp15 and Arg430–peptide center of mass (COM, Figure 3). Both profiles show favorable free energy minima at short distances, suggesting that Arg430 may stabilize the [E-S] complex even in the absence of ideal Trp–Lys geometry. A well-defined free energy minimum is observed at ∼6–7 Å for Arg430– Asp15, indicating a likely salt bridge. A broader minimum in the Arg430–COM profile suggests more diffuse stabilizing interactions between the enzyme and the peptide backbone. Notably, the strongest stabilization is observed when Arg430 interacts directly with ChlA Asp15, indicating a potential anchoring role.

### ChlH modifies peptides unrelated to ChlA

Given that ChlH exhibited broad, leader peptide-independent substrate tolerance, we next asked whether ChlH could halogenate FusA, an unrelated lasso precursor peptide. After enzymatic processing, FusA is converted into fusilassin (also known as fuscanodin),^29,36,37,30^ which contains two Trp. Linear FusA_core_ was reacted with ChlH for 24 h and a combination of unmodified, mono- and dichlorinated FusA_core_ was observed (Figure 4). Analogous to the lack of activity on *des*-chlorolassin, ChlH did not accept the threaded lasso peptide fusilassin as a substrate (Figure S28). Although ChlA and FusA are both lasso precursor peptides, they show no core region similarity (Figure S29). We next assessed if a linear variant of the darobactin precursor peptide was a substrate for ChlH (i.e., DarA-Q8*).^38,39^ We observed monochlorination of DarA-Q8* after reaction with ChlH (Figure 4). As with cyclic lasso peptides, we did not detect chlorination of the fused bicyclic darobactin intermediate (Figure S30). In contrast, pyritide A1, a macrocyclic RiPP featuring a central pyridine and two Trp, was fully monochlorinated and partially dichlorinated by ChlH (Figure 4).^40,41^ HR-MS confirmed the molecular formula of the [M+H]^+^ for the dichlorinated pyritide A1 (C_49_H_59_N_10_O_9_Cl_2_; theoretical, 1001.3838 Da; observed, 1001.3835 Da, 0.3 ppm error) (Figure S31). Attempts to localize the chlorination site in the monochlorinated peptide were inconclusive given the limited fragmentation observed (Figure S31). For the dichlorinated product, we hypothesize each Trp was singly chlorinated, as there has been no evidence that ChlH performs dichlorination of a single Trp.

**Figure 4.**
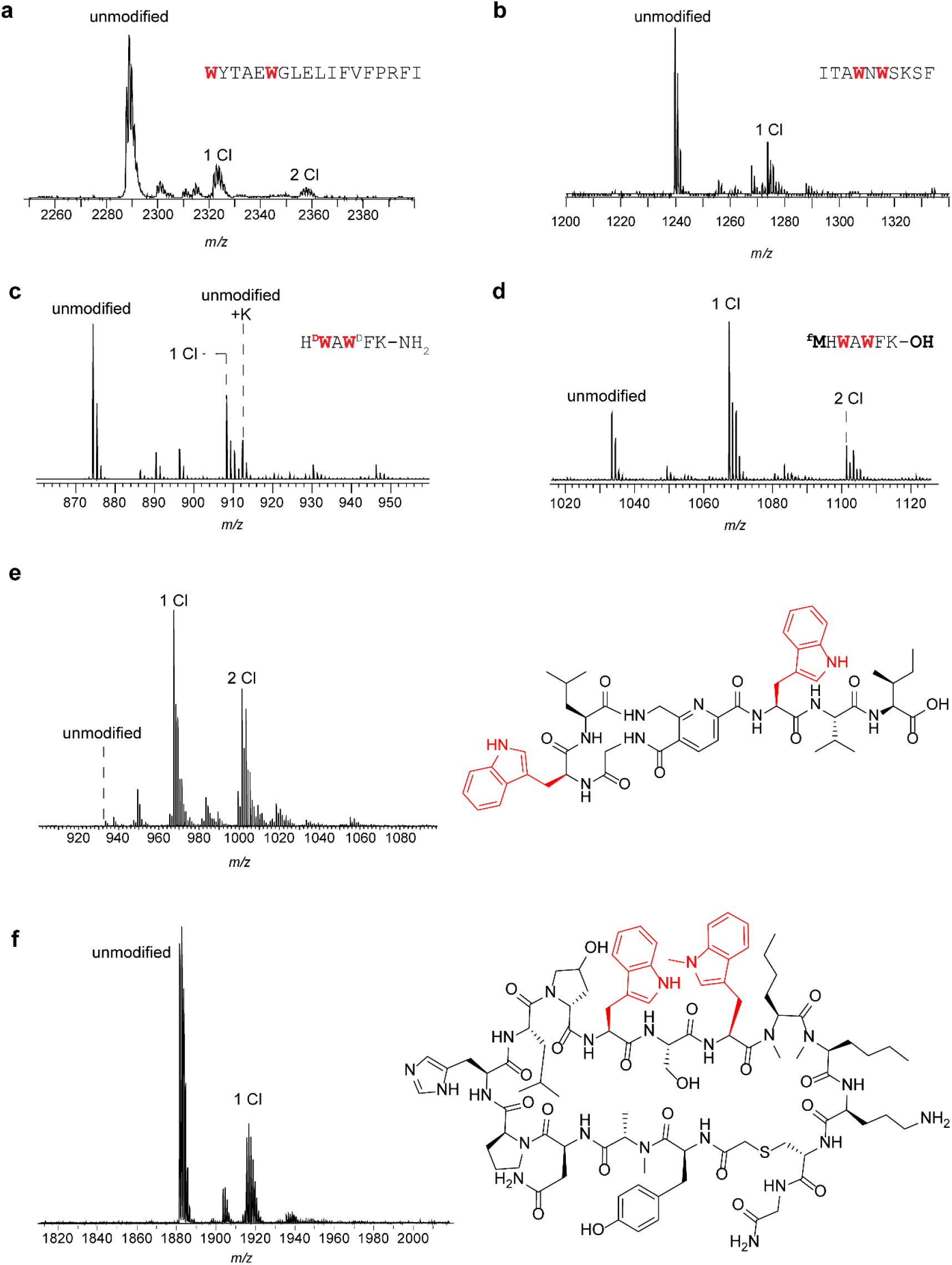
ChlH reactivity towards alternative substrates. (a) MALDI-TOF-MS of FusA_core_ (expected *m/z*: unmodified, 2287; 1 Cl, 2321; 2 Cl, 2355) and (b) DarA-Q8* (shown: GluC fragment, expected *m/z*: unmodified, 1240; 1 Cl, 1274) after treatment with ChlH/R. (c) MALDI-TOF-MS of GHRP-6 (expected *m/*z: unmodified, 873; 1 Cl, 907) and (d) an in vitro-prepared analog (expected *m/z*: unmodified, 1034; 1 Cl, 1067; 2 Cl, 1101) after treatment with ChlH/R. (e) MALDI-TOF-MS of pyritide A1 (expected *m/z*: unmodified, 934; 1 Cl, 967; 2 Cl, 1001). (f) MALDI-TOF-MS of WL12 peptide (expected *m/z*: unmodified, 1881; 1 Cl, 1915).

We also evaluated the capability of ChlH to perform Trp chlorination of two pharmacologically relevant peptides. GHRP-6 (growth hormone-releasing peptide 6) is a hexapeptide containing both *L*-Trp and *D*-Trp that stimulates growth hormone secretion and displays cardioprotective properties.^42–44^ Upon reaction with ChlH, we observed monochlorinated GHRP-6 and a new mass deviating by +1 Da that we hypothesized resulted from amide hydrolysis of the C-terminus (Figure 4).^45^ Tandem MS localized the +1 Da shift to the C-terminal residue and the chlorination event to *D*-Trp2 of GHRP-6 (Figure S32). This result was reminiscent of a recent report on Thal that showed properly placed *L*- and *D*-Trp could be halogenated.^17^ To evaluate if an all-*L*-configured isomer of GHRP-6 was a ChlH substrate, a synthetic DNA template coding for the relevant amino acid sequence was prepared and subjected to *in vitro* transcription/translation. The resulting peptide was treated with ChlH, then analyzed by MALDI-TOF-MS. The highest intensity peak corresponded to monochlorination while a lower intensity ion was consistent with dichlorination (Figure 4). MALDI-LIFT-MS localized the major chlorination site to *L*-Trp2 (Figure S33).

Additionally, we assessed the potential of ChlH to halogenate a head-to-tail cyclized macrocyclic peptide. Our test candidate was WL12 peptide, a known binder of programmed death ligand 1 (PD-L1) which is an immune checkpoint protein involved in T-cell evasion by some types of cancer cells.^46^ This 15-amino acid macrocyclic peptide contains a Trp-Ser-^(N-Me)^Trp motif where the *N*-methylation is on the side chain indole nitrogen. After treatment with ChlH, MS analysis revealed an ion consistent with single chlorination. High-resolution MS (Figure S34) confirmed chlorination, although precise MS/MS localization was not possible because of the close proximity of the two Trp residues and insufficient fragmentation. In summary, this result demonstrates a surprising level of structural diversity in the peptides accepted as substrates, from standard linear to fully macrocyclic scaffolds.

### Cell-free insights into ChlH substrate tolerance

Encouraged by the diversity of peptides modified by ChlH, we designed additional ChlA variants to investigate the extent of substrate tolerance through *in vitro* transcription/translation. Given the leader peptide independence of ChlH, this region was replaced with a single N-terminal Met (installed as formyl-Met, ^f^Met) to allow translation initiation.^47^ These experiments employed a I2K variant of ChlA to enhance MS-based detection. We refer to the *in vitro* transcription/translation prepared ChlA core peptide bearing N(-1)^f^M and I2K substitutions as ChlA_PE_ (Figure 5).

**Figure 5.**
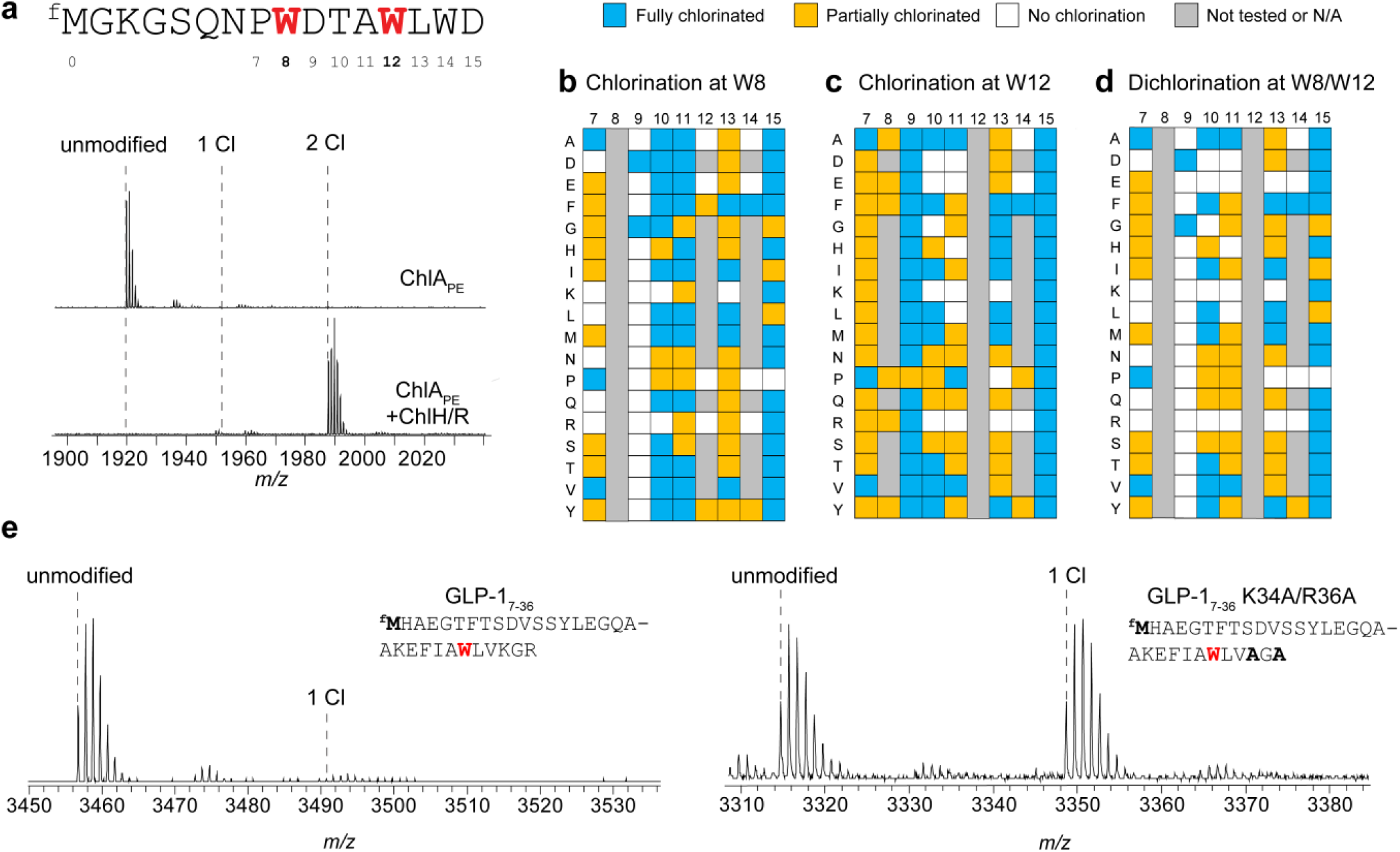
Cell-free mutagenic scanning informs substrate engineering. (a) Sequence and MALDI-TOF-MS of ChlA_PE_ after reaction with ChlH/R. Expected *m/z*: unmodified, 1920; 2 Cl, 1988. (b) Assessment of Trp8 chlorination of the 120-membered mutagenic panel after reaction with ChlH/R. (c) Same as b, except the status of Trp12 chlorination is indicated. (d) Assessment of Trp8 and Trp12 dichlorination. (e) Left spectrum, MALDI-TOF-MS of GLP-1_7-36_ after reaction with ChlH/R. Expected *m/z*: unmodified, 3457; 1 Cl, 3491. Right spectrum, same except with the K34A/R36A variant of GLP-1_7-36_. Expected *m/z*: unmodified, 3315; 1 Cl, 3349.

Analogous to ChlA and ChlA_core_, ChlA_PE_ was dichlorinated at Trp8 and Trp12 after a 1 h reaction with ChlH (Figure 5). A ChlA_PE_ variant with residues 8-15 duplicated and appended to the C-terminus was next tested (Figure S35). This variant contains six Trp, two being preceded by Leu and thus likely to undergo delayed chlorination. After reaction with ChlH for 1 h, a mixture of masses indicative of 3-4 chlorinations was observed. MALDI-LIFT-MS tentatively localized chlorination to Trp6, 10, 14, and 18 (Figure S36), highlighting that motif duplication doubled the number of chlorinations observed. An inverted ChlA_PE_ sequence was also tested, which yielded low levels of chlorination after treatment with ChlH (Figure S37). The position corresponding to Trp14 of ChlA, was preferentially chlorinated, while the position corresponding to Trp12 of ChlA was likely unmodified (Figure S38). This highlights the role of both flanking residues in modification by ChlH.

To reassess the inhibitory role of a preceding Leu, we generated ChlA_PE_ variant P7L/A11L, which placed Leu directly before all three Trp residues. Upon reaction with ChlH for 1 h, monochlorination was observed (Figure S39). MALDI-LIFT-MS localized chlorination to Trp8, indicating that Leu11 effectively delayed Trp12 chlorination yet Leu7 did not delay Trp8 chlorination (Figure S40).

The contribution of core residues was next evaluated by creating an extensive mutagenic panel that probed positions 7-15 of ChlA_PE_. Additionally, all three Trp were individually replaced with Ala, Glu, Phe, Pro, Arg, and Tyr for a total of 120 variants tested. After *in vitro* transcription/translation and reaction with ChlH, the samples were subjected to MS analysis to determine chlorination stoichiometry and location (Figures 5, S41-49, and Supplementary Data Set 1).

We first analyzed monochlorinated products. Most variants tested displayed at least one chlorination, highlighting the general substrate tolerance of ChlH. A total of 43 variants were fully modified at Trp8 while 50 variants were fully modified at Trp12. While ChlH displayed versatility in accepting many substrate sequences, the residues flanking the Trp modification site influenced the reaction outcome. For example, Trp8 was only modified when Asp9 was changed to Gly; however, Trp12 was chlorinated in every tested Asp9 variant. Additionally, many Ala11 variants displayed reduced or no chlorination at Trp12, but over 50% of the tested variants were fully chlorinated at Trp8. Generally, small residues were preferred in the (-1) position (i.e., core positions 7 and 11), while tolerance at the (+1) positions (i.e., core positions 9 and 13) differed substantially between Trp8 and Trp12.

We then analyzed dichlorinated products from the 120-membered panel of ChlA_PE_ variants (Figure 5). Several variants (*n* = 29; 24%) resulted in full dichlorination. Position 15 was the most permissive of those tested, as every substitution except Pro yielded a dichlorinated product. Positions 10 and 13 exhibited similar preferences for hydrophobic residues. Positively charged residues at position 10 prevented modification and broadly inhibited chlorination at all positions except 15.

Additionally, we constructed three variants of ChlH in which an Arg residue within the putative active site was replaced with Ala: R430A, R486A, and R571A (Figure S1). ChlH-R430A was impaired in modifying ChlA_PE_, supporting the findings of our MD that this residue forms important contact points with the substrate (Figure S50). In contrast, ChlH-R486A and -R571A dichlorinated ChlA_PE_ identically to wild-type ChlH under our conditions.

### ChlH modifies a GLP-1 variant

Building on the mutagenic panel, we next tested whether ChlH could halogenate glucagon-like peptide 1 (GLP-1^7-36^). This peptide serves as the basis for a series of type 2 diabetes- and obesity-related drugs.^48–50^ GLP-1^7-36^ contains a single Trp that received only trace chlorination after reaction with ChlH (Figure 5). While Trp25 is flanked by small hydrophobic residues, the nearby Lys28 and Arg30 residues may be inhibitory, as evidenced from the variant library where Lys/Arg residues prevented chlorination of nearby Trp. ChlH processing of the K28A/R30A variant of GLP-1^7-36^ was substantially improved (Figure 5).

### ChlH modifies an internal Trp of an active enzyme

Lastly, we asked if ChlH could modify Trp in a full-length protein. Thus, we tested a set of readily available, purified proteins that are responsible for thiomuracin biosynthesis.^41,51^ MBP-TbtA, -TbtD, -TbtE, -TbtF, and -TbtG and His^6^-TbtB and -TbtC were individually treated with ChlH, trypsinized, and analyzed by MS. Out of 97 unique Trp across seven proteins, we observed modification of only a single Trp, MBP-TbtG Trp471 (WP_013130815.1, Figure 6). As no experimentally determined structure is available for MBP-TbtG, we generated an AF3 model of TbtG. Trp74 of TbtG (corresponding to Trp471 in MBP-TbtG), is not solvent exposed in the predicted structure (ptm score of 0.95). MBP-TbtG was catalytically active, confirming proper folding of the protein (Figure S51).^33^ To determine if ChlH modified the trypsinized MBP-TbtG fragment or intact MBP-TbtG, an in-gel trypsin digest was performed following the reaction. Trp chlorination was still observed, confirming that the non-trypsinyzed protein MBP-TbtG was modified by ChlH. We speculate that TbtG may be flexible, such that ChlH primarily modified an alternate conformation in which Trp471 was more accessible. Potential complexation with the precursor peptide TbtA, TbtE, or TbtF may alter the conformation. Notably, the local sequence of TbtG that was chlorinated is partially analogous to ChlA (Figure S52). Overall, while this is a striking result, we do not expect native ChlH to find broad use in the modification of proteins.

**Figure 6.**
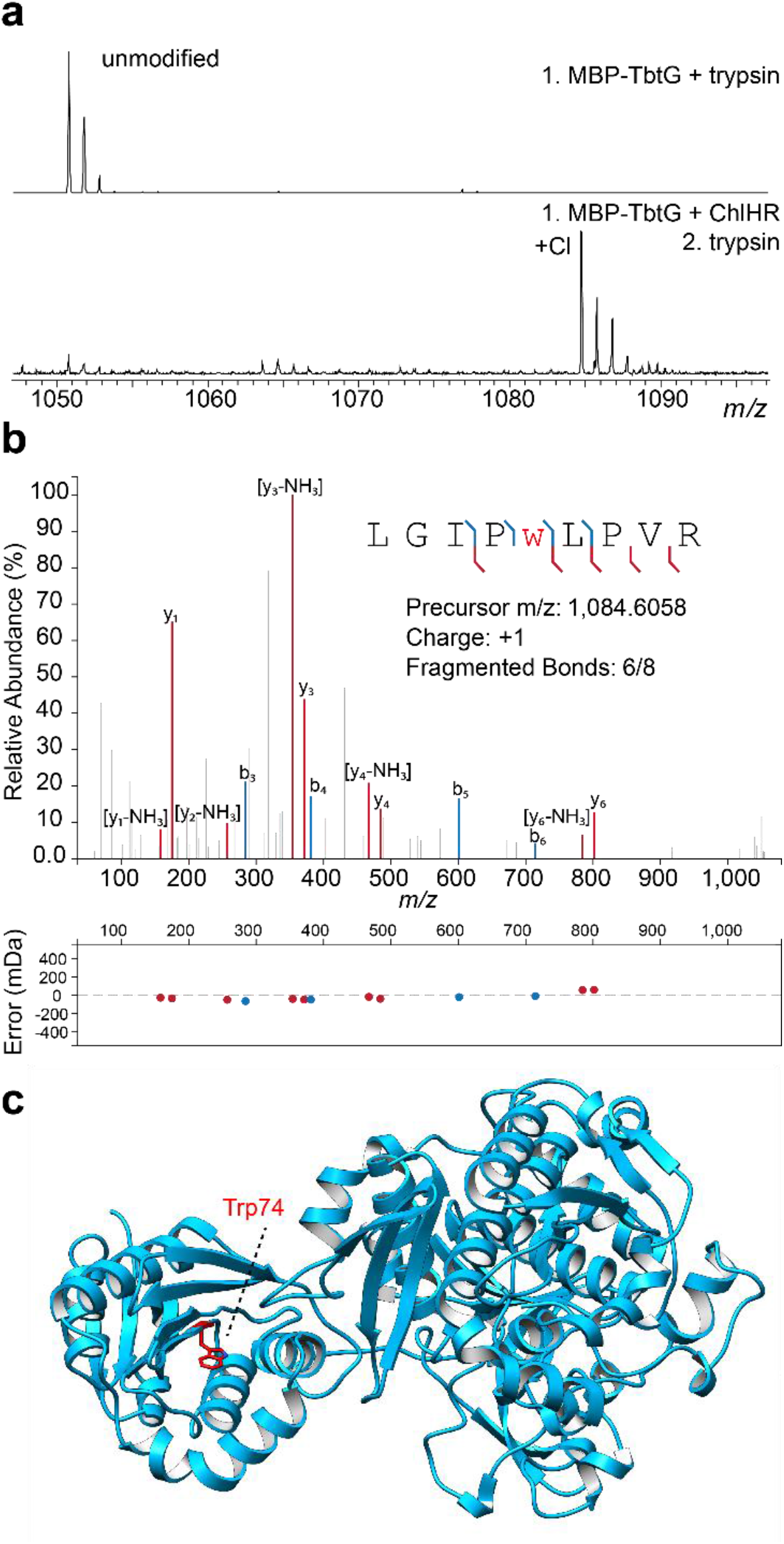
Evaluation of ChlH protein substrate halogenation. (a) MALDI-TOF-MS of MBP-TbtG reacted with ChlH/R with subsequent trypsin digestion. The expected *m/z* values are 1050.6 for the unmodified fragment and 1084.7 for the chlorinated fragment. (b) MALDI-LIFT-MS analysis of the tryptic peptide containing Cl-Trp (lowercase, red w). (c) AF3 structural prediction of TbtG (modified Trp is red).

## Discussion

The data reported here show that ChlH acts on the linear precursor peptide ChlA and not the threaded lasso peptide *des*-chlorolassin. ChlH shares 58% sequence identity with the lanthipeptide-modifying MibH. An AF3-enabled structural alignment of ChlH with the crystal structure of MibH (PDB: 5UAO) shows high predicted structural similarity with a global RMSD of 0.5 Å (Figure S53).^20^ Despite this similarity, these proteins modify their respective substrates at distinct biosynthetic stages. This may be rationalized by the size differences in the active site cavities of each enzyme. CASTpFold calculated a 442 Å^3^ volume for ChlH while the active site cavity for MibH is nearly 50% larger (652 Å^3^, Figure S54).^52^ Caution should be applied in interpreting these values, however, as they are derived from static structures, and our molecular dynamics simulations suggest that ChlH undergoes structural rearrangement upon binding of ChlA. Additionally, while FDHs acting on free *L*-Trp are often sequence-similar, known FDHs acting on peptide substrates can be highly sequence-diverse (Figure S55).

Using ChlA as the substrate led us to wonder why Trp14 was unmodified *in vivo* but minimally modified *in vitro*. Our results indicated that rather than utilizing the leader peptide of ChlA for substrate recognition, ChlH seems to recognize the C-terminal region of the ChlA core peptide (i.e., Leu13-Trp14). While the C-terminal motif expedited modification *in vitro*, it was dispensable, as numerous substrates lacking the motif were successfully modified. Despite contributing to processing elsewhere, Trp14 is slowly modified *in vitro*. Our results show that ChlH is broadly tolerant and capable of modifying disparate Trp-containing peptides. Because of this, ChlH activity may need to be tightly regulated in *Lentzea jiangxiensis*. Indeed, both *des*-chlorolassin and monochlorolassin were observed during the purification of chlorolassin,^26^ demonstrating that during native production, ChlA is incompletely modified prior to lasso cyclization.

Nature often utilizes halogenation to tailor the physical properties of natural products. For example, the FDH DarH naturally brominates the 5-position of Trp1 in darobactin.^53^ While capable of iodination, DarH could not chlorinate darobactin. ChlH, in contrast, can chlorinate and brominate substrates but only acts on the linear precursor of darobactin. The complementary activities of DarH and ChlH could be envisioned to provide access to new-to-nature halogenated darobactin variants.

While several residues of ChlA (Ala11-Trp14) were important for robust processing by ChlH, an atomic-resolution co-crystal structure was not obtained and thus the precise substrate-binding mode remains to be investigated. Furthermore, mutagenic scanning revealed that modification of Trp8 and Trp12 are not entirely independent. For example, most Pro7 variants experienced hindered processing not only at the adjacent Trp8 but also at the more distant Trp12. Further work is needed to shed light on the precise details of the ChlA-ChlH interaction.

Notably, while this manuscript was in preparation, another research group reconstituted ChlH.^54^ These authors reported that ChlH was incapable of chlorinating variants of ChlA which contained additional Trp residues. Although a precise cause of this discrepancy cannot be determined, we believe it may be due to some key differences in protein affinity tags that were employed. While a maltose-binding protein (MBP) tag (∼40 kDa) was used in the other work to purify ChlH, we utilized a hexahistidine tag (0.84 kDa) for ChlH purification. Further, the authors reported that the leader peptide was critical for substrate modification and involved in a predicted binding interaction with ChlH,^54^ while we found the leader peptide to be fully dispensable. Further, we did not observe any compelling evidence for this proposed leader-enzyme interaction *in vitro* or *in silico*. Similar to our initial, low-confidence AF2M predicted model, the simulations reported in the recent publication had ChlA Trp8 to ChlH Lys139 distances from 13–19 Å, which are far too long for catalytic competence.^54^ In contrast, we utilized conformations where all Trp residues were proximal to Lys 139 as starting points for further analyses. Our simulations also show that the inability of Trp14 to maintain functional proximity to Lys139 might be related to an intramolecular interaction of Leu13 that causes Trp14 to move away from the catalytic lysine. This hypothesis is supported by the observed inhibitory role of Leu in the (-1) position. Our use of weighted ensemble sampling and a geometrically informed scoring function provided a dynamic view of Trp–Lys interactions and catalytic competence.

While several residues of ChlA (Ala11-Trp14) were important for robust processing by ChlH, an atomic-resolution co-crystal structure was not obtained, and the precise substrate-binding mode is not completely understood. Furthermore, mutagenic scanning revealed that modification of Trp8 and Trp12 are not entirely independent. For example, most Pro7 variants experienced hindered processing not only at the adjacent Trp8 but also at the more distant Trp12. Further work is needed to shed light on the precise details of the ChlA-ChlH interaction.

When compared with traditional *in vitro* methods, cell-free biosynthesis rapidly expedites the assessment of substrate promiscuity for enzymes acting on peptides.^55^ Several general observations about the substrate preferences of ChlH can be made from our extensive screening efforts. When modifying wild-type ChlA, an N-terminal (-1) Leu is inhibitory. Small hydrophobic residues are often preferred. Positively charged residues are broadly inhibitory. We suspect that this could be due to electrostatic repulsion against positively charged residues within the ChlH active site, although we have shown that single-site mutagenesis is insufficient to modify these substrate preferences.

## Conclusion

In conclusion, we have reconstituted a versatile Trp halogenase enzyme ChlH and its cognate flavin reductase ChlR. Unexpectedly, ChlH modifies the linear precursor peptide ChlA rather than the threaded lasso peptide *des*-chlorolassin. We further assessed the substrate scope of ChlH and achieved chlorination in a broad array of Trp-containing peptidic scaffolds. Linear and macrocyclic peptides, as well as a larger protein, were effectively chlorinated without the need for additional recognition motifs. While several residues of ChlA do contribute to more efficient processing by ChlH, they are not strictly required for modification. The broad applicability of ChlH makes it an attractive candidate for the biocatalytic halogenation of peptidic Trp residues.

## Supporting information

ChlH_SI_BioRxiv

Supplementary Data Set 1

## Supporting Information

Experimental methods and Supporting Figures S1-S55 (PDF)

Supplementary Data Set 1: Assigned ions associated with MS/MS data (.xlsl)

Supplementary Movie 1: Trp8-focused weighted ensemble simulations of ChlH and ChlA

Supplementary Movie 2: Trp12-focused weighted ensemble simulations of ChlH and ChlA

Supplementary Movie 3: Trp14-focused weighted ensemble simulations of ChlH and ChlA

The authors have cited additional references within the Supporting Information.^[56-64]^

## Acknowledgements

The authors thank Hamada Saad and Austin Woodard supplying various substrates.

## References

(1) Atanasov, A. G.; Zotchev, S. B.; Dirsch, V. M.; Supuran, C. T. Natural Products in Drug Discovery: Advances and Opportunities. Nat Rev Drug Discov 2021, 20 (3), 200–216. 10.1038/s41573-020-00114-z.

(2) Benedetto Tiz, D.; Bagnoli, L.; Rosati, O.; Marini, F.; Sancineto, L.; Santi, C. New Halogen-Containing Drugs Approved by FDA in 2021: An Overview on Their Syntheses and Pharmaceutical Use. Molecules 2022, 27 (5), 1643. 10.3390/molecules27051643.

(3) Andorfer, M. C.; Lewis, J. C. Understanding and Improving the Activity of Flavin Dependent Halogenases via Random and Targeted Mutagenesis. Annu Rev Biochem 2018, 87, 159–185. 10.1146/annurev-biochem-062917-012042.

(4) Mori, S.; Pang, A. H.; Thamban Chandrika, N.; Garneau-Tsodikova, S.; Tsodikov, O. V. Unusual Substrate and Halide Versatility of Phenolic Halogenase PltM. Nat Commun 2019, 10 (1), 1255. 10.1038/s41467-019-09215-9.

(5) Gkotsi, D. S.; Ludewig, H.; Sharma, S. V.; Connolly, J. A.; Dhaliwal, J.; Wang, Y.; Unsworth, W. P.; Taylor, R. J. K.; McLachlan, M. M. W.; Shanahan, S.; Naismith, J. H.; Goss, R. J. M. A Marine Viral Halogenase That Iodinates Diverse Substrates. Nature Chemistry 2019, 11 (12), 1091–1097. 10.1038/s41557-019-0349-z.

(6) Gruß, H.; Sewald, N. Late-Stage Diversification of Tryptophan-Derived Biomolecules. Chemistry – A European Journal 2020, 26 (24), 5328–5340. 10.1002/chem.201903756.

(7) Bradley, S. A.; Lehka, B. J.; Hansson, F. G.; Adhikari, K. B.; Rago, D.; Rubaszka, P.; Haidar, A. K.; Chen, L.; Hansen, L. G.; Gudich, O.; Giannakou, K.; Lengger, B.; Gill, R. T.; Nakamura, Y.; de Bernonville, T. D.; Koudounas, K.; Romero-Suarez, D.; Ding, L.; Qiao, Y.; Frimurer, T. M.; Petersen, A. A.; Besseau, S.; Kumar, S.; Gautron, N.; Melin, C.; Marc, J.; Jeanneau, R.; O’Connor, S. E.; Courdavault, V.; Keasling, J. D.; Zhang, J.; Jensen, M. K. Biosynthesis of Natural and Halogenated Plant Monoterpene Indole Alkaloids in Yeast. Nat Chem Biol 2023, 19 (12), 1551–1560. 10.1038/s41589-023-01430-2.

(8) Mondal, H.; Patra, S.; Saha, S.; Nayak, T.; Sengupta, U.; Sudan Maji, M. Late-Stage Halogenation of Peptides, Drugs and (Hetero)Aromatic Compounds with a Nucleophilic Hydrazide Catalyst. Angewandte Chemie International Edition 2023, 62 (51), e202312597. 10.1002/anie.202312597.

(9) Lewis, J. C. Identifying and Engineering Flavin Dependent Halogenases for Selective Biocatalysis. Acc. Chem. Res. 2024, 57 (15), 2067–2079. 10.1021/acs.accounts.4c00172.

(10) Yeh, E.; Blasiak, L. C.; Koglin, A.; Drennan, C. L.; Walsh, C. T. Chlorination by a Long-Lived Intermediate in the Mechanism of Flavin-Dependent Halogenases,. Biochemistry 2007, 46 (5), 1284–1292. 10.1021/bi0621213.

(11) Walsh, C. T.; Wencewicz, T. A. Flavoenzymes: Versatile Catalysts in Biosynthetic Pathways. Nat. Prod. Rep. 2012, 30 (1), 175–200. 10.1039/C2NP20069D.

(12) Schnepel, C.; Sewald, N. Enzymatic Halogenation: A Timely Strategy for Regioselective C-H Activation. Chemistry – A European Journal 2017, 23 (50), 12064–12086. 10.1002/chem.201701209.

(13) Büchler, J.; Papadopoulou, A.; Buller, R. Recent Advances in Flavin-Dependent Halogenase Biocatalysis: Sourcing, Engineering, and Application. Catalysts 2019, 9 (12), 1030. 10.3390/catal9121030.

(14) Payne, J. T.; Andorfer, M. C.; Lewis, J. C. Regioselective Arene Halogenation Using the FAD-Dependent Halogenase RebH. Angewandte Chemie International Edition 2013, 52 (20), 5271–5274. 10.1002/anie.201300762.

(15) Phintha, A.; Prakinee, K.; Jaruwat, A.; Lawan, N.; Visitsatthawong, S.; Kantiwiriyawanitch, C.; Songsungthong, W.; Trisrivirat, D.; Chenprakhon, P.; Mulholland, A.; van Pée, K.-H.; Chitnumsub, P.; Chaiyen, P. Dissecting the Low Catalytic Capability of Flavin-Dependent Halogenases. Journal of Biological Chemistry 2021, 296, 100068. 10.1074/jbc.RA120.016004.

(16) Prakinee, K.; Phintha, A.; Visitsatthawong, S.; Lawan, N.; Sucharitakul, J.; Kantiwiriyawanitch, C.; Damborsky, J.; Chitnumsub, P.; van Pée, K.-H.; Chaiyen, P. Mechanism-Guided Tunnel Engineering to Increase the Efficiency of a Flavin-Dependent Halogenase. Nat Catal 2022, 5 (6), 534–544. 10.1038/s41929-022-00800-8.

(17) Schnepel, C.; Moritzer, A.-C.; Gäfe, S.; Montua, N.; Minges, H.; Nieß, A.; Niemann, H. H.; Sewald, N. Enzymatic Late-Stage Halogenation of Peptides. ChemBioChem 2023, 24 (1), e202200569. 10.1002/cbic.202200569.

(18) Sana, B.; Ke, D.; Li, E. H. Y.; Ho, T.; Seayad, J.; Duong, H. A.; Ghadessy, F. J. Halogenation of Peptides and Proteins Using Engineered Tryptophan Halogenase Enzymes. Biomolecules 2022, 12 (12), 1841. 10.3390/biom12121841.

(19) Montua, N.; Sewald, N. Perfect Partners: Biocatalytic Halogenation and Metal Catalysis for Protein Bioconjugation. ChemBioChem 2024, n/a (n/a), e202400496. 10.1002/cbic.202400496.

(20) Ortega, M. A.; Cogan, D. P.; Mukherjee, S.; Garg, N.; Li, B.; Thibodeaux, G. N.; Maffioli, S. I.; Donadio, S.; Sosio, M.; Escano, J.; Smith, L.; Nair, S. K.; van der Donk, W. A. Two Flavoenzymes Catalyze the Post-Translational Generation of 5-Chlorotryptophan and 2-Aminovinyl-Cysteine during NAI-107 Biosynthesis. ACS Chem. Biol. 2017, 12 (2), 548–557. 10.1021/acschembio.6b01031.

(21) Nguyen, N. A.; Lin, Z.; Mohanty, I.; Garg, N.; Schmidt, E. W.; Agarwal, V. An Obligate Peptidyl Brominase Underlies the Discovery of Highly Distributed Biosynthetic Gene Clusters in Marine Sponge Microbiomes. J. Am. Chem. Soc. 2021. 10.1021/jacs.1c03474.

(22) Nguyen, N. A.; Agarwal, V. A Leader-Guided Substrate Tolerant RiPP Brominase Allows Suzuki–Miyaura Cross-Coupling Reactions for Peptides and Proteins. Biochemistry 2023. 10.1021/acs.biochem.3c00222.

(23) Nguyen, N. A.; Vidya, F. N. U.; Yennawar, N. H.; Wu, H.; McShan, A. C.; Agarwal, V. Disordered Regions in Proteusin Peptides Guide Post-Translational Modification by a Flavin-Dependent RiPP Brominase. Nat Commun 2024, 15 (1), 1265. 10.1038/s41467-024-45593-5.

(24) Milbredt, D.; Patallo, E. P.; van Pée, K.-H. A Tryptophan 6-Halogenase and an Amidotransferase Are Involved in Thienodolin Biosynthesis. ChemBioChem 2014, 15 (7), 1011–1020. 10.1002/cbic.201400016.

(25) Montua, N.; Thye, P.; Hartwig, P.; Kühle, M.; Sewald, N. Enzymatic Peptide and Protein Bromination: The BromoTrp Tag. Angewandte Chemie International Edition 2024, 63 (5), e202314961. 10.1002/anie.202314961.

(26) Harris, L. A.; Saad, H.; Shelton, K. E.; Zhu, L.; Guo, X.; Mitchell, D. A. Tryptophan-Centric Bioinformatics Identifies New Lasso Peptide Modifications. Biochemistry 2024. 10.1021/acs.biochem.4c00035.

(27) Suckau, D.; Resemann, A.; Schuerenberg, M.; Hufnagel, P.; Franzen, J.; Holle, A. A Novel MALDI LIFT-TOF/TOF Mass Spectrometer for Proteomics. Analytical and Bioanalytical Chemistry 2003, 376 (7), 952–965. 10.1007/s00216-003-2057-0.

(28) Jiang, Y.; Snodgrass, H. M.; Zubi, Y. S.; Roof, C. V.; Guan, Y.; Mondal, D.; Honeycutt, N. H.; Lee, J. W.; Lewis, R. D.; Martinez, C. A.; Lewis, J. C. The Single-Component Flavin Reductase/Flavin-Dependent Halogenase AetF Is a Versatile Catalyst for Selective Bromination and Iodination of Arenes and Olefins**. Angewandte Chemie 2022, 134 (51), e202214610. 10.1002/ange.202214610.

(29) DiCaprio, A. J.; Firouzbakht, A.; Hudson, G. A.; Mitchell, D. A. Enzymatic Reconstitution and Biosynthetic Investigation of the Lasso Peptide Fusilassin. J. Am. Chem. Soc. 2019, 141 (1), 290–297. 10.1021/jacs.8b09928.

(30) Kretsch, A. M.; Gadgil, M. G.; DiCaprio, A. J.; Barrett, S. E.; Kille, B. L.; Si, Y.; Zhu, L.; Mitchell, D. A. Peptidase Activation by a Leader Peptide-Bound RiPP Recognition Element. Biochemistry 2023. 10.1021/acs.biochem.2c00700.

(31) Nguyen, N. A.; Agarwal, V. A Leader-Guided Substrate Tolerant RiPP Brominase Allows Suzuki–Miyaura Cross-Coupling Reactions for Peptides and Proteins. Biochemistry 2023, 62 (12), 1838–1843. 10.1021/acs.biochem.3c00222.

(32) Jumper, J.; Evans, R.; Pritzel, A.; Green, T.; Figurnov, M.; Ronneberger, O.; Tunyasuvunakool, K.; Bates, R.; Žídek, A.; Potapenko, A.; Bridgland, A.; Meyer, C.; Kohl, S. A. A.; Ballard, A. J.; Cowie, A.; Romera-Paredes, B.; Nikolov, S.; Jain, R.; Adler, J.; Back, T.; Petersen, S.; Reiman, D.; Clancy, E.; Zielinski, M.; Steinegger, M.; Pacholska, M.; Berghammer, T.; Bodenstein, S.; Silver, D.; Vinyals, O.; Senior, A. W.; Kavukcuoglu, K.; Kohli, P.; Hassabis, D. Highly Accurate Protein Structure Prediction with AlphaFold. Nature 2021, 596 (7873), 583–589. 10.1038/s41586-021-03819-2.

(33) Abramson, J.; Adler, J.; Dunger, J.; Evans, R.; Green, T.; Pritzel, A.; Ronneberger, O.; Willmore, L.; Ballard, A. J.; Bambrick, J.; Bodenstein, S. W.; Evans, D. A.; Hung, C.-C.; O’Neill, M.; Reiman, D.; Tunyasuvunakool, K.; Wu, Z.; Žemgulytė, A.; Arvaniti, E.; Beattie, C.; Bertolli, O.; Bridgland, A.; Cherepanov, A.; Congreve, M.; Cowen-Rivers, A. I.; Cowie, A.; Figurnov, M.; Fuchs, F. B.; Gladman, H.; Jain, R.; Khan, Y. A.; Low, C. M. R.; Perlin, K.; Potapenko, A.; Savy, P.; Singh, S.; Stecula, A.; Thillaisundaram, A.; Tong, C.; Yakneen, S.; Zhong, E. D.; Zielinski, M.; Žídek, A.; Bapst, V.; Kohli, P.; Jaderberg, M.; Hassabis, D.; Jumper, J. M. Accurate Structure Prediction of Biomolecular Interactions with AlphaFold 3. Nature 2024, 630 (8016), 493–500. 10.1038/s41586-024-07487-w.

(34) Zhang, B. W.; Jasnow, D.; Zuckerman, D. M. The “Weighted Ensemble” Path Sampling Method Is Statistically Exact for a Broad Class of Stochastic Processes and Binning Procedures. J Chem Phys 2010, 132 (5), 054107. 10.1063/1.3306345.

(35) Russo, J. D.; Zhang, S.; Leung, J. M. G.; Bogetti, A. T.; Thompson, J. P.; DeGrave, A. J.; Torrillo, P. A.; Pratt, A. J.; Wong, K. F.; Xia, J.; Copperman, J.; Adelman, J. L.; Zwier, M. C.; LeBard, D. N.; Zuckerman, D. M.; Chong, L. T. WESTPA 2.0: High-Performance Upgrades for Weighted Ensemble Simulations and Analysis of Longer-Timescale Applications. J Chem Theory Comput 2022, 18 (2), 638–649. 10.1021/acs.jctc.1c01154.

(36) Koos, J. D.; Link, A. J. Heterologous and in Vitro Reconstitution of Fuscanodin, a Lasso Peptide from Thermobifida Fusca. J. Am. Chem. Soc. 2019, 141 (2), 928–935. 10.1021/jacs.8b10724.

(37) Si, Y.; Kretsch, A. M.; Daigh, L. M.; Burk, M. J.; Mitchell, D. A. Cell-Free Biosynthesis to Evaluate Lasso Peptide Formation and Enzyme–Substrate Tolerance. J. Am. Chem. Soc. 2021. 10.1021/jacs.1c01452.

(38) Kaur, H.; Jakob, R. P.; Marzinek, J. K.; Green, R.; Imai, Y.; Bolla, J. R.; Agustoni, E.; Robinson, C. V.; Bond, P. J.; Lewis, K.; Maier, T.; Hiller, S. The Antibiotic Darobactin Mimics a β-Strand to Inhibit Outer Membrane Insertase. Nature 2021, 1–5. 10.1038/s41586-021-03455-w.

(39) Woodard, A. M.; Peccati, F.; Navo, C. D.; Jiménez-Osés, G.; Mitchell, D. A. Darobactin Substrate Engineering and Computation Show Radical Stability Governs Ether versus C–C Bond Formation. J. Am. Chem. Soc. 2024, 146 (20), 14328–14340. 10.1021/jacs.4c03994.

(40) Nguyen, D. T.; Le, T. T.; Rice, A. J.; Hudson, G. A.; van der Donk, W. A.; Mitchell, D. A. Accessing Diverse Pyridine-Based Macrocyclic Peptides by a Two-Site Recognition Pathway. J. Am. Chem. Soc. 2022, 144 (25), 11263– 11269. 10.1021/jacs.2c02824.

(41) Rice, A. J.; Pelton, J. M.; Kramer, N. J.; Catlin, D. S.; Nair, S. K.; Pogorelov, T. V.; Mitchell, D. A.; Bowers, A. A. Enzymatic Pyridine Aromatization during Thiopeptide Biosynthesis. J. Am. Chem. Soc. 2022, 144 (46), 21116–21124. 10.1021/jacs.2c07377.

(42) Lei, T.; Buchfelder, M.; Fahlbusch, R.; Adams, E. F. Growth Hormone Releasing Peptide (GHRP-6) Stimulates Phosphatidylinositol (PI) Turnover in Human Pituitary Somatotroph Cells. 1995. 10.1677/jme.0.0140135.

(43) Berlanga-Acosta, J.; Abreu-Cruz, A.; Barco Herrera, D. G.; Mendoza-Marí, Y.; Rodríguez-Ulloa, A.; García-Ojalvo, A.; Falcón-Cama, V.; Hernández-Bernal, F.; Beichen, Q.; Guillén-Nieto, G. Synthetic Growth Hormone-Releasing Peptides (GHRPs): A Historical Appraisal of the Evidences Supporting Their Cytoprotective Effects. Clin Med Insights Cardiol 2017, 11, 1179546817694558. 10.1177/1179546817694558.

(44) Berlanga-Acosta, J.; Cibrian, D.; Valiente-Mustelier, J.; Suárez-Alba, J.; García-Ojalvo, A.; Falcón-Cama, V.; Jiang, B.; Wang, L.; Guillén-Nieto, G. Growth Hormone Releasing Peptide-6 (GHRP-6) Prevents Doxorubicin-Induced Myocardial and Extra-Myocardial Damages by Activating Prosurvival Mechanisms. Front. Pharmacol. 2024, 15. 10.3389/fphar.2024.1402138.

(45) Santana, H.; Espinosa, L. A.; Sánchez, A.; Bolaño Alvarez, A.; Besada, V.; González, L. J. Mass Spectrometric and Kinetics Characterization of Modified Species of Growth Hormone Releasing Hexapeptide Generated under Thermal Stress in Different pH and Buffers. Journal of Pharmaceutical and Biomedical Analysis 2021, 194, 113776. 10.1016/j.jpba.2020.113776.

(46) Rapid PD-L1 Detection in Tumors with PET Using a Highly Specific Peptide. Biochemical and Biophysical Research Communications 2017, 483 (1), 258–263. 10.1016/j.bbrc.2016.12.156.

(47) Liu, M.; Thijssen, V.; Jongkees, S. A. K. Suppression of Formylation Provides an Alternative Approach to Vacant Codon Creation in Bacterial In Vitro Translation. Angewandte Chemie International Edition 2020, 59 (49), 21870– 21874. 10.1002/anie.202003779.

(48) Lau, J.; Bloch, P.; Schäffer, L.; Pettersson, I.; Spetzler, J.; Kofoed, J.; Madsen, K.; Knudsen, L. B.; McGuire, J.; Steensgaard, D. B.; Strauss, H. M.; Gram, D. X.; Knudsen, S. M.; Nielsen, F. S.; Thygesen, P.; Reedtz-Runge, S.; Kruse, T. Discovery of the Once-Weekly Glucagon-Like Peptide-1 (GLP-1) Analogue Semaglutide. J. Med. Chem. 2015, 58 (18), 7370–7380. 10.1021/acs.jmedchem.5b00726.

(49) Popoviciu, M.-S.; Păduraru, L.; Yahya, G.; Metwally, K.; Cavalu, S. Emerging Role of GLP-1 Agonists in Obesity: A Comprehensive Review of Randomised Controlled Trials. International Journal of Molecular Sciences 2023, 24 (13), 10449. 10.3390/ijms241310449.

(50) Jastreboff, A. M.; Kushner, R. F. New Frontiers in Obesity Treatment: GLP-1 and Nascent Nutrient-Stimulated Hormone-Based Therapeutics. Annual Review of Medicine 2023, 74 (Volume 74, 2023), 125–139. 10.1146/annurev-med-043021-014919.

(51) Hudson, G. A.; Zhang, Z.; Tietz, J. I.; Mitchell, D. A.; van der Donk, W. A. In Vitro Biosynthesis of the Core Scaffold of the Thiopeptide Thiomuracin. J. Am. Chem. Soc. 2015, 137 (51), 16012–16015. 10.1021/jacs.5b10194.

(52) Ye, B.; Tian, W.; Wang, B.; Liang, J. CASTpFold: Computed Atlas of Surface Topography of the Universe of Protein Folds. Nucleic Acids Research 2024, 52 (W1), W194–W199. 10.1093/nar/gkae415.

(53) Böhringer, N.; Kramer, J.-C.; de la Mora, E.; Padva, L.; Wuisan, Z. G.; Liu, Y.; Kurz, M.; Marner, M.; Nguyen, H.; Amara, P.; Yokoyama, K.; Nicolet, Y.; Mettal, U.; Schäberle, T. F. Genome-and Metabolome-Guided Discovery of Marine BamA Inhibitors Revealed a Dedicated Darobactin Halogenase. Cell Chemical Biology 2023, 30 (8), 943-952.e7. 10.1016/j.chembiol.2023.06.011.

(54) Lu, J.-L.; Cui, J.-J.; Hu, Z.-Y.; Di, J.-M.; Li, Y.-Y.; Xiong, J.; Jiao, Y.-M.; Gao, K.; Min, J.; Luo, S.; Dong, S.-H. Characterization of an Iterative Halogenase Acting on Ribosomal Peptides Underlies the Combinatorial Biosynthesis Logic of Lasso Peptides. J. Nat. Prod. 2025. 10.1021/acs.jnatprod.4c01199.

(55) Rice, A. J.; Sword, T. T.; Chengan, K.; Mitchell, D. A.; Mouncey, N. J.; Moore, S. J.; Bailey, C. B. Cell-Free Synthetic Biology for Natural Product Biosynthesis and Discovery. Chem. Soc. Rev. 2025. 10.1039/D4CS01198H.

(56) Brademan, D. R.; Riley, N. M.; Kwiecien, N. W.; Coon, J. J. Interactive Peptide Spectral Annotator: A Versatile Web-Based Tool for Proteomic Applications*. Mol. Cell. Proteomics 2019, 18 (8, Supplement 1), S193–S201. 10.1074/mcp.TIR118.001209.

(57) Nishihara, K.; Kanemori, M.; Kitagawa, M.; Yanagi, H.; Yura, T. Chaperone Coexpression Plasmids: Differential and Synergistic Roles of DnaK-DnaJ-GrpE and GroEL-GroES in Assisting Folding of an Allergen of Japanese Cedar Pollen, Cryj2, inEscherichia Coli. Appl. Environ. Microbiol. 1998, 64 (5), 1694–1699. 10.1128/AEM.64.5.1694-1699.1998.

(58) Nishihara, K.; Kanemori, M.; Yanagi, H.; Yura, T. Overexpression of Trigger Factor Prevents Aggregation of Recombinant Proteins in Escherichia Coli. Appl. Environ. Microbiol. 2000, 66 (3), 884–889. 10.1128/AEM.66.3.884-889.2000.

(59) Gaussian 16, Revision A.03, Frisch, M. J.; Trucks, G. W.; Schlegel, H. B.; Scuseria, G. E.; Robb, M. A.; Cheeseman, J. R.; Scalmani, G.; Barone, V.; Petersson, G. A.; Nakatsuji, H.; Li, X.; Caricato, M.; Marenich, A. V.; Bloino, J.; Janesko, B. G.; Gomperts, R.; Mennucci, B.; Hratchian, H. P.; Ortiz, J. V.; Izmaylov, A. F.; Sonnenberg, J. L.; Williams-Young, D.; Ding, F.; Lipparini, F.; Egidi, F.; Goings, J.; Peng, B.; Petrone, A.; Henderson, T.; Ranasinghe, D.; Zakrzewski, V. G.; Gao, J.; Rega, N.; Zheng, G.; Liang, W.; Hada, M.; Ehara, M.; Toyota, K.; Fukuda, R.; Hasegawa, J.; Ishida, M.; Nakajima, T.; Honda, Y.; Kitao, O.; Nakai, H.; Vreven, T.; Throssell, K.; Montgomery, J. A., Jr.; Peralta, J. E.; Ogliaro, F.; Bearpark, M. J.; Heyd, J. J.; Brothers, E. N.; Kudin, K. N.; Staroverov, V. N.; Keith, T. A.; Kobayashi, R.; Normand, J.; Raghavachari, K.; Rendell, A. P.; Burant, J. C.; Iyengar, S. S.; Tomasi, J.; Cossi, M.; Millam, J. M.; Klene, M.; Adamo, C.; Cammi, R.; Ochterski, J. W.; Martin, R. L.; Morokuma, K.; Farkas, O.; Foresman, J. B.; Fox, D. J. Gaussian, Inc., Wallingford CT, 2016.

(60) Wang, J.; Wang, W.; Kollman, P. A.; Case, D. A. Automatic Atom Type and Bond Type Perception in Molecular Mechanical Calculations. J. Mol. Graph. Model. 2006, 25 (2), 247–260. 10.1016/j.jmgm.2005.12.005.

(61) Wang, J.; Wolf, R. M.; Caldwell, J. W.; Kollman, P. A.; Case, D. A. Development and Testing of a General Amber Force Field. J. Comput. Chem. 2004, 25 (9), 1157–1174. 10.1002/jcc.20035.

(62) Madeira, F.; Madhusoodanan, N.; Lee, J.; Eusebi, A.; Niewielska, A.; Tivey, A. R. N.; Lopez, R.; Butcher, S. The EMBL-EBI Job Dispatcher Sequence Analysis Tools Framework in 2024. Nucleic Acids Res. 2024, 52 (W1), W521– W525. 10.1093/nar/gkae241.

(63) Zhang, Z.; Hudson, G. A.; Mahanta, N.; Tietz, J. I.; van der Donk, W. A.; Mitchell, D. A. Biosynthetic Timing and Substrate Specificity for the Thiopeptide Thiomuracin. J. Am. Chem. Soc. 2016, 138 (48), 15511–15514. 10.1021/jacs.6b08987.

(64) Pettersen, E. F.; Goddard, T. D.; Huang, C. C.; Couch, G. S.; Greenblatt, D. M.; Meng, E. C.; Ferrin, T. E. UCSF Chimera—A Visualization System for Exploratory Research and Analysis. J. Comput. Chem. 2004, 25 (13), 1605– 1612. 10.1002/jcc.20084.

