## Supplementary material for "Peptidic Tryptophan Halogenation by a Promiscuous Flavin-dependent Enzyme": ChlH_SI_BioRxiv

[a] Departments of Biochemistry and Chemistry, Vanderbilt University, Medical Research Building IV, Nashville, TN 37232. USA.

**Table of Contents:**


Materials and Methods S3
Figure S1: Coomassie-stained SDS-PAGE gels of proteins used in this study S8
Figure S2: *Des*-chlorolassin and monochlorolassin are not substrates for ChlH *in vitro* S9
Figure S3: Free tryptophan was not chlorinated by ChlH *in vitro* S10
Figure S4: MALDI-LIFT-MS of dichlorinated ChlA S11
Figure S5: MALDI-LIFT-MS to assess the order of chlorination on ChlA S12
Figure S6: MALDI-TOF-MS of ChlA/H/R after a 24 h reaction S13
Figure S7: MALDI-LIFT-MS of trichlorinated ChlA S14
Figure S8: Sequence alignment of RebH and ChlH S15
Figure S9: MALDI-TOF-MS for ChlH-K139A variant activity S16
Figure S10: MALDI-TOF-MS for ChlH halogen scope reactions S17
Figure S11: MALDI-TOF-MS analysis of selective ChlA bromination S18
Figure S12: MALDI-TOF-MS for ChlA vs $ChlA_{core}$ halogenation reactions S19
Figure S13: MALDI-TOF-MS for ChlA Ala variants at 2 h S20
Figure S14: MALDI-TOF-MS for ChlA Ala variants at 24 h S21
Figure S15: MALDI-TOF-MS for additional ChlA variants at 2 h S22
Figure S16: MALDI-TOF-MS for additional ChlA variants at 24 h S23
Figure S17: Sequence of $ChlA_{ext}$ S24
Figure S18: MALDI-TOF-MS for ChlA Trp variants at 2 h S25
Figure S19: MALDI-TOF-MS for ChlA Trp variants at 24 h S26
Figures S20-S24: MALDI-LIFT-MS for select Trp scan variants S27-31
Figure S25: AlphaFold 2 Multimer model of ChlA and ChlH S32
Figure S26: AlphaFold 3 model of ChlA and ChlH S33
Table S1: Molecular dynamics setups S34
Figure S27: Raw free energy surfaces S35
Figure S28: MALDI-TOF-MS for attempted chlorination of fusilassin S36
Figure S29: Sequence comparison of ChlA and FusA core regions S37
Figure S30: MALDI-TOF-MS for attempted chlorination of a darobactin biosynthetic intermediate S38
Figure S31: LC-MS/MS of chlorinated pyritide A1 S39
Figure S32: MALDI-LIFT-MS for chlorinated GHRP-6 S40
Figure S33: MALDI-LIFT-MS for chlorinated all-*L*-amino acid GHRP-6 S41
Figure S34: LC-MS/MS for chlorinated WL12 peptide S42
Figure S35: MALDI-TOF-MS for $ChlA_{PE}$ 8-15 duplication substrate chlorination S43
Figure S36: MALDI-LIFT-MS for chlorinated $ChlA_{PE}$ 8-15 duplication substrate S44
Figure S37: MALDI-TOF-MS for $ChlA_{PE}$ inverted substrate chlorination S45
Figure S38: MALDI-LIFT-MS for chlorinated $ChlA_{PE}$ inverted substrate S46
Figure S39: MALDI-TOF-MS for $ChlA_{PE}$ P7L/A11L chlorination S47
Figure S40: MALDI-LIFT-MS for chlorinated $ChlA_{PE}$ P7L/A11L S48
Figures S41-49: MALDI-TOF-MS and -LIFT-MS for $ChlA_{PE}$ single site variants S49-193
Figure S50: ChlH Arg variant activity on $ChlA_{PE}$ S194
Figure S51: Assessment of catalytic activity from MBP-TbtG S195
Figure S52: Alignment highlighting similarities between TbtG-Trp74 and ChlA S196
Figure S53: Alignment of MibH crystal structure and ChlH AlphaFold 3 structure S197
Figure S54: Active site cavity measurement for ChlH AlphaFold 3 structure S198
Figure S55: Sequence identity/similarity matrix for flavin-dependent Trp halogenases S199
Supporting References S200-201

### Materials and Methods.

**General materials and methods.** Reagents used were purchased from New England BioLabs (Ipswich, MA), Thermo Fisher Scientific (Waltham, MA), Gold Biotechnology Inc. (St. Louis, MO), or Sigma-Aldrich (St. Louis, MO). GLP-1 (residues 7-36) was purchased from ProSpec (Rehovot, Israel), GHRP-6 was purchased from Echelon Biosciences (Salt Lake City, UT), WL12 peptide was purchased from CPC Scientific (Milpitas, CA), and synthetic ChlA WT and L13A/W14A core peptides were purchased from Genscript (Piscataway, NJ). Escherichia coli DH5α and BL21(DE3) strains were used for plasmid production and protein expression, respectively. Sanger sequencing was performed by the Core DNA Sequencing Facility at the University of Illinois at Urbana-Champaign. Matrix-assisted laser desorption/ionization time-of-flight mass spectrometry (MALDI-TOF-MS) analysis and MALDI-LIFT-TOF/TOF-MS (herein referred to as MALDI-LIFT-MS) was performed using a Bruker UltrafleXtreme MALDI TOF-TOF mass spectrometer (Bruker Daltonics) in reflector positive mode at the University of Illinois School of Chemical Sciences Mass Spectrometry Laboratory and in the Mass Spectrometry Research Core at Vanderbilt University. All samples were analyzed in positive reflector mode. After collection of MS/MS data, fragment ions were annotated using the Interactive Peptide Spectral Annotator webtool.[1] All HR-MS/MS data was collected in the Mass Spectrometry Research Core at Vanderbilt University on a ThermoFisher Scientific LTQ Orbitrap XL coupled to a Waters Acquity UPLC. A ThermoFisher Scientific Vanquish C18+ column (50 × 2.1mm) was used, employing a flow rate of 0.3 mL/min with water and acetonitrile containing 0.1% formic acid as the mobile phase.

**Cloning.** *E. coli*-optimized genes encoding ChlH and ChlR were individually cloned into multiple cloning site 1 (MCS1) of a pRSFDuet vector containing an N-terminal hexahistidine tag via Gibson ligation. Briefly, DNA sequences corresponding to the gene of interest and pRSFDuet vector were PCR amplified with Q5 polymerase. PCR reactions were analyzed by agarose gel electrophoresis, and bands corresponding to the fragments of interest were excised with a razor blade. Following this, gel pieces were digested in QG buffer (20 mM Tris, 5.5 M guanidinium thiocyanate, pH 6.5) at 60 °C for 10 min with occasional vortexing. Once gel pieces were dissolved, samples were transferred to a spin column and concentrated via centrifugation at 17,000 × g for 2 min. Samples were washed twice with wash buffer (80% ethanol in $H_2O$) and then eluted using elution buffer (50 mM Tris, pH 7). Gibson cloning was achieved using 100-200 ng of DNA in a ratio of 1:5 vector to insert with an equal volume of NEBuilder HiFi DNA Assembly 2x Master Mix (New England Biolabs). Gibson ligations were performed at 55 °C for 15 min, after which 1 μL of each reaction was transformed into *E. coli* DH5α competent cells. Plasmids were amplified, purified, and sequence-confirmed via Sanger sequencing.

For MBP-ChlA, Gibson ligation was performed as above with the following modifications. The insert sequence for ChlA was amplified from genomic DNA of *Lentzia jiangxiensis*. The vector DNA was amplified from an empty pET28a plasmid containing a maltose-binding protein (MBP) tag with a tobacco etch virus (TEV) protease cleavage site and two hexahistidine tags.

*E. coli*-optimized ChlH nucleotide sequence (5’ to 3’):

```
ATGAAAATGAATGCCGGACGTCCCTCAGCTTTGGACGAATTGGTGCAGACATTTAGCGATGACGAGTCTGCCGCGGTCCGTGAATG
GTTAGGCGGTGCAGCAGACGCACCTCAACTGCCAGGGATTTCGTTACGCTCAAATCATGAGCCGCTGGTCGAAATGCTTCCGCGTC
CAGCAGTAGACGATCAGAAGGCAATTCGCCGCGTGGCCGTAGTTGGGGGTGGTACTGCCGGTTATCTTACTGCAATCGCCCTGCGC
ACCAAGCGTCCATGGCTTGAGGTTTCTTTGATTGAATCCCCCTCGATCCCGATTATTGGGGTCGGGGAAGCGACTGTTCCTGGGAT
TGTTATTTTCTTGCATCATTATTTAGGTATCGACGTTGAGGACTTCTACCGTCAGGTTCGTCCCACTTGGAAACAGGGAATTCGCT
TCGAGTGGGGACCGCGCCCAGATGGCTTCATGGCACCGTTTGACTGGGATTCGAACTCGATCGGAACTGCGGGCGCCGTTGACGCC
GGGGGTGGTATCGACGGGTACACCTTACAGTCACTGCTGATGTCAGCTGAGCGCGCTCCGGTTTTCGATGTCGGGGATCGCCATTT
ATCACTTATGAAGTATCTGCCTTTTGCATATCATCTTGACAATGCTCGTTTTGTAAGCTATTTAGCCGACATCGCAAAACAGCGCG
GGGTGCAGCACGTCGAGGCTAAGATTGAAGATGTTGCTTTGTCCGGTGATGATTGGGTCGATCACTTGCACACAACTGATGGCCGT
AAACTGTCCTATGATTTTTACGTCGACTGCTCCGGCTTTCGCTCCATGCTTTTAGGAAAAGCATTTCGTACCCCATTCCATTCGTA
CGCTTCCTCACTTTTCACCGATAGCGCGGTCACGGGTAATGTAGGGCACGGAGGGAACTTGAAGCCGTATACGTCTGCCATTACTA
TGAATTCAGGGTGGTGTTGGGATATTCCGACACCGCAAGACGACCACTTAGGCTACGTGTACTCGTCTAATGCTATCAGCGACGAC
CAGGCGGCGGCGGAATTAGCAGCGCGCTTCCCCGGCGTTTCAGAGCCGCGCTTCGTGCGCTTCCGCGTCGGACGTCACGAAAAGGC
ATGGCGTGGAAACGTAATGGCTATCGGCAATTCGTACGCCTTCGTGGAACCGTTAGAGTCGACAGGAATCCTTATGATTACAAGTA
GTGTGCAGGCGCTTGTCGCCTCGCTGCCGGGAAGCTGGGCCGATCCGCAATGTCGCGATATGGTTAATGTCGGATTAGCTAACCGC
TGGGACGCTTTGCGCTGGTTCCTGTCGATCCACTACCGTTTCAACACTCGTCGTGATACGCCATTTTGGCGCGAGGTGCGCGCCAA
TACGGACATCTCCGGCGCCCAGCCCATGCTGGATATGTATGCAACTGGCGCTCCATTGCGTTACCGTAATCCTCTGTTACGTACCT
TCGTACAGGGCAATGCCCCCACGTTCTACGGGTTAGCTGGGGTTGAGACGATTTTATTGGGCCAACAAGTCCCTACTCGTCTTCTT
```

CACCGTGCAGAGCCGCCGGCCCGTTGGCAGGCTCGTCGCAACGCGGCTGATGCTTTTGTTCGTCGCGCACTGCCGCAGCTTGAAGC
ATTGAGCCGCTTTCATGATGACCCGGCACTTAATCAACAACTGCTTTTTGACGCTGATAGTTGGGCCTCTCCATTACGCGCAGAAC
CAATCGGGATGTTATGA

*E. coli*-optimized ChlR nucleotide sequence (5' to 3'):

ATGGCCGGTTTCCCGACCGGGGTCGCGATCATCACCACCGTCGACACCGACGGCACGCCGCGCGGGATGACGTGCTCGTCGCTGTG
CAGCGTCACGCTCGCCCCACCGACGCTGCTGGCCTGCCTGCGCACCGGCAGCCCGACCCTGGCCGCGGTCCTGCGCCGAGGCGTCT
TCGCCGTCAACCTGGTGCATGGCGAGGCACGCGCTACGGCGGAATTATTTGCAAGCGGAGCCCAGGACCGTTTTGCTCGTGTGCGT
TGGCAAGCAGAAACAGGAGGTCCACATTTGCCCGACGATGCCCACGCTGTCGCCGATTGCGCAGTCCGTCGCGCAGAGCCTGTAGG
GGATCACCACGTGGTTTTCGGGGAGGTCTTACGCATTGCAGGGAGCGAAGCAACCGACCCCCTTTTATACGGTATGCGCCGCTATG
CACGTCTGCCGACGGAGGTGGGAAGCTGA

**Protein expression and purification.** For ChlH and ChlR, pRSFDuet vectors encoding the protein of interest were co-transformed into chemically competent BL21(DE3) *E. coli* cells with pGro7, a plasmid from the Takara chaperone set encoding GroES, GroEL, and chloramphenicol resistance.[2,3] Cells were grown on solid lysogeny broth (LB) media containing 37.5 μg/mL kanamycin and 10 μg/mL chloramphenicol overnight. Single colonies were used to inoculate 10 mL of LB medium containing 37.5 μg/mL kanamycin and 10 μg/mL chloramphenicol and grown at 37 °C with shaking at 250 rpm. After 18 h, 1 L of LB was inoculated with the starter culture after addition of 37.5 μg/mL kanamycin and 10 μg/mL chloramphenicol. Cells were grown to an optical density ($OD_{600}$) of 0.6-0.8. Cells were then placed on ice for 15 min, 0.4 mM (final) isopropyl β-D-1-thiogalactopyranoside (IPTG) and 0.4% w/v L-(+)-arabinose were added, and the cultures grown for 18 h at 25 °C. Cells were harvested by centrifugation at 4,500 × g for 25 min, washed once with phosphate-buffered saline and re-centrifuged at 4,500 × g for 15 min. Cell pellets were flash-frozen in liquid nitrogen and stored at -80 °C for a maximum of one week before use.

For purification, cell pellets were thawed on ice and resuspended in 30 mL of lysis buffer [50 mM $NaH_2PO_4$, 900 mM NaCl, 20 mM imidazole, 15 mM β-mercaptoethanol, 2.5% glycerol (*v/v*), and 0.1% triton X-100 (*v/v*), pH 8] containing 4 mg/mL lysozyme, 2 μM leupeptin, 2 μM benzamidine, 2 μM E64, and 30 mg phenylmethylsulfonyl fluoride (PMSF). Cells were then gently rocked for 30 min at 4 °C before 4 × 45 s of sonication, with 10 min of rocking between sonication rounds. Cellular debris was removed by centrifugation (20,000 × g, 90 min) and the supernatant was applied to pre-equilibrated amylose resin (5 mL of resin per L of cells). The column was washed with 10 column volumes (CV) of lysis buffer, then 10 CV of wash buffer [50 mM $NaH_2PO_4$, 900 mM NaCl, 20 mM imidazole, 15 mM β-mercaptoethanol, and 2.5% glycerol (*v/v*), pH 8]. $His_6$-tagged proteins were eluted using 5 CV of elution buffer [50 mM $NaH_2PO_4$, 900 mM NaCl, 250 mM imidazole, 15 mM β-mercaptoethanol, and 2.5% glycerol (*v/v*), pH 8]. Eluent was concentrated using an Amicon ultracentrifugal filter (EMD Millipore) with an appropriate molecular weight cutoff. Protein was buffer exchanged with 10× volume of protein storage buffer [50 mM 4-(2-hydroxyethyl)-1-piperazineethanesulfonic acid (HEPES), 300 mM NaCl, 2.5% glycerol (*v/v*), and 0.5 mM TCEP, pH 8]. Protein concentrations were estimated using 280 nm absorbance, with theoretical extinction coefficients calculated using the ExPasy ProtParam Tool (http://web.expasy.org/protparam/protpar-ref.html).

MBP-TbtG, MBP-FusB, and MBP-FusE were expressed and purified as described previously.[4,5]

**Halogenation reactions of MBP-ChlA variants**. Halogenation assays were performed in 20 mM sodium phosphate ($NaH_2PO_4$) buffer at pH 7.4, 10 mM NaCl, 5 μM flavin adenine dinucleotide (FAD), 5 μM $His_6$-ChlR, 5 μM $His_6$-ChlH, 5 μM TEV protease, and 50 μM substrate. Reactions were initiated by the addition of 100 μM reduced nicotinamide adenine dinucleotide (NADH). Reactions were allowed to proceed at 30 °C for 2 h or 24 h. After the reaction period, samples were concentrated by centrifugation (17,000 × g, 5 min, 25 °C) to remove any insoluble material, desalted using a C18 Ziptip, and analyzed by MALDI-TOF-MS using Super 2,5-dihydroxybenzoic acid (SDHB) matrix.

**Halogenation reactions of other peptide substrates**. For reactions employing non-MBP tagged peptide substrates, TEV protease was excluded. All other conditions were identical to those stated in the section "Halogenation reactions of MBP-ChlA variants". Substrates which suffered from low water solubility were first dissolved in DMSO to generate a stock solution and then diluted into the reaction mixture to a 50 μM final concentration.

**Site-directed mutagenesis.** Site-directed mutagenesis was performed as described previously.[6]

**Halogen assessment**. Halogenation assays were performed and analyzed as described above with the following changes. Reactions were performed using a buffer containing 20 mM $NaH_2PO_4$ pH 7.4, and 100 mM of NaF, NaCl, NaBr, and NaI. For selective halogenation reactions, MBP-ChlA, TEV protease, $His_6$-ChlH and $His_6$-ChlR were extensively buffer exchanged into bromide-only buffer (50 mM HEPES, 300 mM NaBr, 2.5% glycerol (*v/v*) pH 8) prior to use in bromination reactions. Additionally, 10 mM DTT was added to the final reaction mixture.

**LC-MS analysis of free *L*-Trp halogenation**. *L*-Trp (0.5 mM) was reacted with 20 μM ChlH and 20 μM ChlR. All other buffer and temperature conditions were identical to those listed in the Methods section titled "General halogenation reactions". At the 24 h timepoint, the mixture was analyzed by a Shimadzu LC-MS 2020 single quadrupole system under positive mode electrospray ionization. For the liquid chromatography component, a ThermoFisher Scientific Hypersil GOLD column (150 × 2.1 mm) was employed. Solvents A ($H_2O$) and B (acetonitrile) both contained 0.1% formic acid. The LC method began with equilibration at 5% B for 5 min following injection, followed by a gradient from 5-90% B over 30 min, hold at 90% B for a further 5 min, ramp down to 5% B over 1 min, and finally a hold at 5% B for 10 min.

**HPLC purification of ChlA precursor peptide**. To a 300 μM solution of MBP-ChlA in the following reaction buffer (50 mM Tris-HCl pH 7.5, 125 mM NaCl, 10 mM DTT), 30 μM of TEV protease was added. The reaction was allowed to proceed at 30 °C for 3 h after which an equal volume of acetonitrile was added to quench the reaction. This mixture was subjected to centrifugation at 17,000 × g for 10 min. The supernatant was then purified using a ThermoFisher Scientific Vanquish HPLC instrument equipped with a Hypersil GOLD C18 semipreparative column (250 × 10 mm). The compositions of solvents A and B were 20 mM $NH_4OAc$ in $H_2O$, and acetonitrile, respectively. The following method was used: equilibration for 15 min at 10% B, gradient of 10-50% B for 45 min, 2 min ramp up to 95% B, held at 95% B for 15 min, 2 min ramp down to 10% B, and then held at 10% B for 20 min. Fractions were collected based on absorbance at 220 and 280 nm.

**HPLC purification of ChlA core peptide**. MBP-FusB and MBP-FusE were expressed and purified as described previously.[5] These two proteins were added at concentrations of 15 μM and 30 μM, respectively, to 300 μM MBP-ChlA in the following reaction buffer (50 mM Tris-HCl pH 7.5, 125 mM NaCl, 10 mM DTT). Reactions were allowed to proceed at 37 °C for 3 h after which a 1:1 (*v/v*) amount of acetonitrile was added to quench the reaction. This mixture was subjected to centrifugation at 17,000 × g for 10 min. The isolated supernatant was then subjected to HPLC purification on a ThermoFisher Scientific Vanquish HPLC instrument equipped with an Accucore C18 column (150 × 4.6 mm). Solvents A ($H_2O$) and B (acetonitrile) both contained 0.1% formic acid. The following general method was used: equilibration for 5 min at 10% B, gradient of 10-50% B for 15 min, ramp up to 90% B over 1 min, held at 90% B for 5 min, ramp down to 10% B over 1 min, and then held at 10% B for 10 min. Fractions were collected based on absorbance at 220 and 280 nm.

**HPLC purification of ChlA core peptide variants post-halogenation**. Following halogenation, MBP-ChlA variants were treated with MBP-FusB (15 μM) and MBP-FusE (30 μM). All following steps were identical to those described in the previous section titled "HPLC purification of ChlA core peptide". After concentration of the isolated fractions, MALDI-LIFT-MS was performed to localize the sites of chlorination.

**HPLC purification of fusilassin**. MBP-FusA, MBP-FusB, MBP-FusC, and MBP-FusE were expressed, purified, and reacted *in vitro* to yield crude fusilassin as described previously[7]. The reaction mixture was precipitated with acetonitrile (60% *v/v* final). After centrifugation, the isolated supernatant was then subjected to HPLC purification on a ThermoFisher Scientific Vanquish HPLC instrument equipped with an Accucore C8 column (150 × 4.6 mm). Solvents A ($H_2O$) and B (acetonitrile) both contained 0.1% formic acid. The following general method was used: equilibration for 5 min at 20% B, gradient of 20-55% B for 15 min, ramp to 90% B over 1 min, held at 90% B for 5 min, ramp down to 20% B over 1 min, and then held at 20% B for 10 min. Fractions were collected based on absorbance at 220 and 280 nm.

**HPLC timecourse assay of synthetic ChlA WT and L13A/W14A core peptides**. All halogenation reaction conditions were identical to those stated in the section "Halogenation reactions of other peptide substrates", except for the concentrations of ChlH (10 μM), ChlR (10 μM), and synthetic ChlA core peptide (200 μM). At each time point, a reaction mixture aliquot was obtained, and acetonitrile was added in a 1:1 (v/v) amount. This mixture was subjected to centrifugation at 17,000 × g for 10 min. HPLC analysis of the supernatant was performed on a ThermoFisher Scientific Vanquish HPLC instrument equipped with an Accucore PFP C18 column (150 × 4.6 mm). Solvents A (20 mM ammonium bicarbonate) and B (acetonitrile) were used. The following method was used: equilibration for 5 min at 20% B, gradient of 20-30% B for 15 min, ramp up to 90% B over 1 min, held at 90% B for 5 min, ramp down to 20% B over 1 min, and then held at 20% B for 10 min. Absorbance at 220 and 280 nm was monitored.

**Template DNA for cell-free reactions**. Linear double-stranded DNA templates for cell-free reactions were generated by overhang-extension PCR. An example of DNA template architecture is shown below:

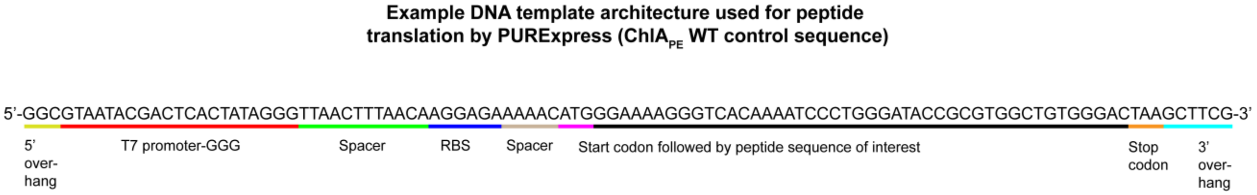


**Cell-free halogenation reactions**. The commercially available PURExpress kit (New England Biolabs) was used for translation of peptides of interest in 2.5 µL scale reactions. 1 µL solution A, 0.75 µL solution B, and 0.75 µL of a DNA template (~300 ng/µL in $H_2O$) were mixed in that order. Translation reactions were allowed to proceed at 37 °C for 3 h. Next, 5 µL of a halogenation reaction mixture was added to the translation reaction, mixed thoroughly, and allowed to react at 30 °C for an additional 1 h. The halogenation reaction mixture consisted of 20 mM $NaH_2PO_4$ buffer at pH 8, 15 µM ChlH, 30 µM ChlR, 5 µM FAD, 100 µM NADH, 10 mM NaCl, and 10 mM DTT. After 1 h, the reactions were diluted by addition of 10 µL $H_2O$, desalted via C18 ZipTip, and analyzed by MALDI-TOF-MS using SDHB matrix.

**Halogenation of MBP-TbtG.** Halogenation assays using protein substrates were performed in 20 mM sodium phosphate buffer at pH 7.4, 10 mM NaCl, 5 μM FAD, 5 μM $His_6$-ChlR, 5 μM $His_6$-ChlH, 5 μM TEV protease, and 50 μM substrate. Reactions were initiated by the addition of 100 μM NADH. Reactions were allowed to proceed at 30 °C for 24 h. After the reaction period, samples were digested with 1 μM trypsin overnight at 37 °C. Reactions were then concentrated via centrifugation (17,000 × g, 5 min, 25 °C) to remove any insoluble material, desalted using a C18 Ziptip, and analyzed by MALDI-TOF-MS using SDHB matrix.

**MBP-TbtG activity.** The MBP-TbtG enzymatic activity was assessed in conjunction with MBP-TbtE and MBP-TbtF, as described previously.[4] These proteins form a trimeric synthetase complex that results in the ATP-dependent conversion of six Cys residues of TbtA into thiazole heterocycles. The TbtF components binds TbtA and delivers the substrate to TbtG, which converts Cys to thiazoline. TbtE transforms thiazolines to thiazoles, which were detected using MS as described previously.[4]

**Ligand Parameterization.** Flavin adenine dinucleotide (FAD) was parameterized using a combination of Gaussian16[8] and Antechamber[9], following standard AMBER force field procedures[10]. Geometry optimization was performed using density functional theory (DFT) at the B3LYP/6-31+G* level with tight self-consistent field (SCF) convergence criteria and the xqc algorithm to ensure robust convergence. The optimized structure, with a –2 total charge and singlet multiplicity, was used to compute the electrostatic potential (ESP) at the HF/6-31+G* level, applying the Merz–Kollman (MK)[10] scheme. RESP partial charges were then derived using Antechamber and assigned alongside AMBER-compatible atom types. The final FAD parameters were incorporated using LEaP.

**System Preparation and Simulation Protocol.** An AlphaFold-predicted model served as a starting structure. The model was solvated and neutralized with NaCl using TLEaP from the AmberTools suite, applying the ff19SB force field for proteins, GAFF2 for ligands, the custom quantum-derived FAD parameters, and the OPC water model. To investigate ChlA binding specificity, three configurations were prepared. In the first, termed the "Trp8-modeled" system, the rotamer of Trp8 was optimized to improve its proximity and orientation relative to Lys139. This optimized pose was then used as a template to manually reposition the ChlA peptide, generating two additional models in which Trp12 or Trp14 replaced Trp8 at the same spatial location. Each of these three systems was simulated in triplicate, resulting in 12 molecular dynamics trajectories with a total of 16.5 μs of aggregate sampling. All systems underwent initial energy minimization using 500 steps of steepest descent with harmonic restraints on the protein backbone. This was followed by NVT equilibration from 10 K to 300 K over 200 ps using a Langevin thermostat ($\gamma\_ln$ = 2 $ps^{-1}$), while backbone restraints were maintained. NPT equilibration followed, employing a stepwise reduction in restraint force constants (from 5.0 to 0.1 kcal/mol·$Å^2$) using a Berendsen barostat and Monte Carlo pressure regulation. Production simulations were carried out under NPT conditions for 0.8-2.5 μs (See Table S1 with a 2 fs timestep and a Langevin thermostat ($\gamma\_ln$ = 1 $ps^{-1}$). In most trajectories, the Trp initially positioned near Lys139 disengaged from the catalytic site within several 100 ns. However, in two simulations of the Trp8-modeled system, Trp8 maintained stable proximity to Lys139. In one of these, the protein underwent rearrangements that promoted persistent retention of the peptide in a bound-like pose. Three frames from this trajectory (at 730 ns, 1500 ns, and 2500 ns) were selected as seeds for weighted ensemble (WE) simulations. From each seed, ChlA was reconfigured to place Trp8, Trp12, or Trp14 near the catalytic Lys139, yielding a complete set of configurations for comparative analysis.

**Weighted Ensemble Simulations**. WE simulations were performed using WESTPA 2.0[11] to enhance sampling of Trp–Lys interactions. Three frames from *original Alfafold_pose_rep 02* trajectory (at 730 ns, 1500 ns, and 2500 ns) were selected as initial configurations. From each seed, ChlA was reconfigured to also place Trp12, or Trp14 near the catalytic lysine, yielding a complete set of configurations for comparative analysis. Each of these configurations was simulated with three replicas, resulting in 27 starting states. WE simulations were run for 28 iterations, generating 2,442 walkers and 280 ns of molecular time, corresponding to ~24.4 μs of aggregated sampling. Progress coordinates were a reaction likelihood score (eq 1) and a discrete index representing Trp identity (Trp8, Trp12, or Trp14). The reaction likelihood score is a combination of a distance and weighted angular function:

$$S = d + \theta \qquad \text{(eq1)}$$

The distance function (*d*) is designed to have a sharp decay above 14 Å (unbound state) and below 5 (clash) and with linear behavior between 5 and 14 Å with optimal distance at 6 Å.

$$d = \begin{cases} \dfrac{1}{1 + e^{\{3\,(|d - 9.5| - 4.5)\}}}, & d < 5 \; or \; d > 14 \\ 1.0 - 0.1(d - 6), & 5 \le d \le 14 \end{cases} \qquad \text{(eq2)}$$

The angular function (*θ*) has a gaussian-like behavior with optimal value for planarity.

$$\theta = e^{\left(-\frac{(180-\theta)^2}{2\sigma_\theta^2}\right)} \qquad \text{(eq3)}$$

The binning was performed using a rectilinear scheme: distance-based bins (score < 2.0) were combined with discrete Trp identity bins. Three walkers per bin were maintained.

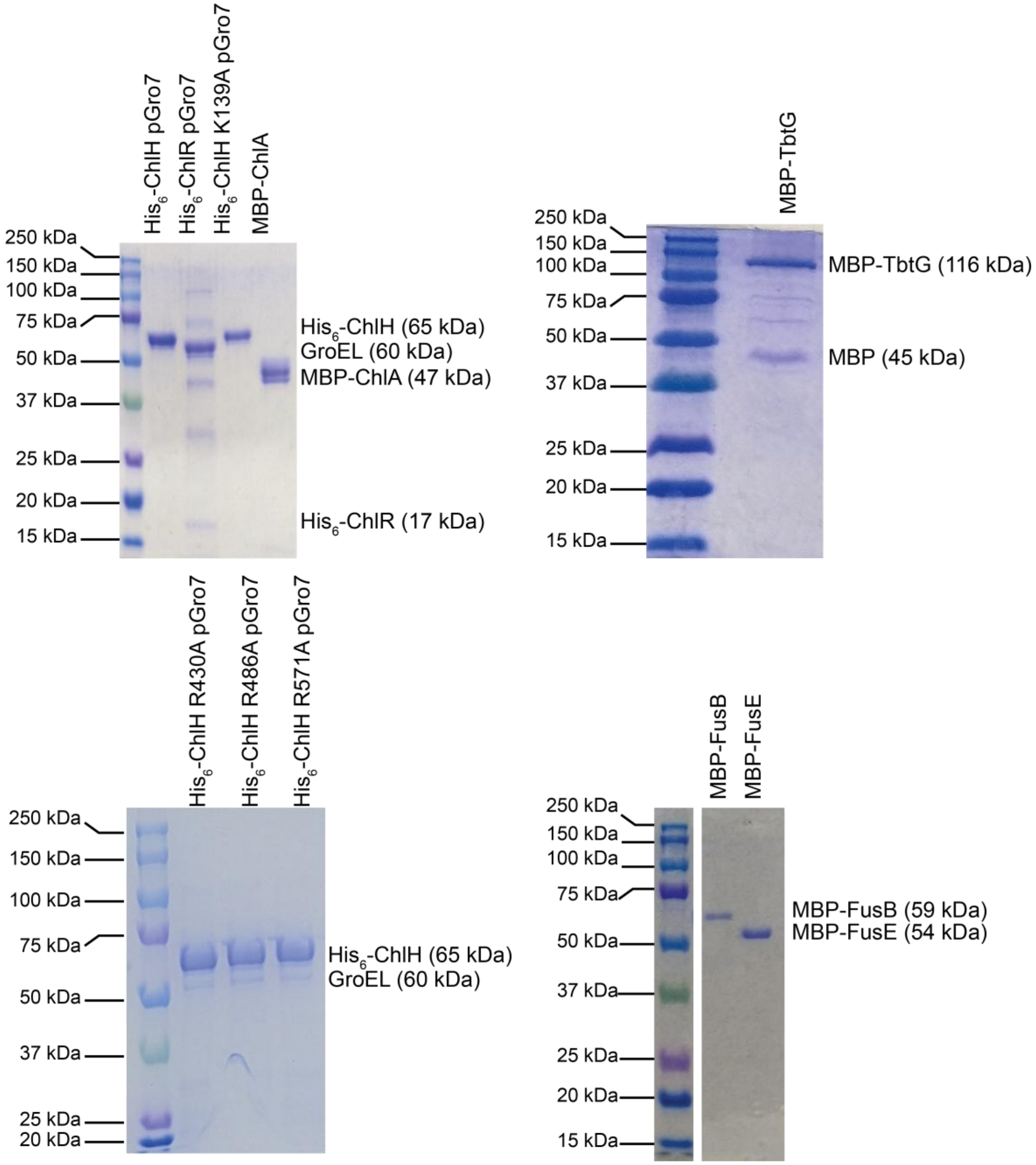


**Figure S1: Coomassie-stained SDS-PAGE gels of proteins used in this study.** pGro7 is a plasmid containing GroES and GroEL.[2,3] Observation of cleaved MBP (maltose-binding protein) arises from interdomain cleavage of MBP-TbtG from endogenous proteases in *E. coli*.

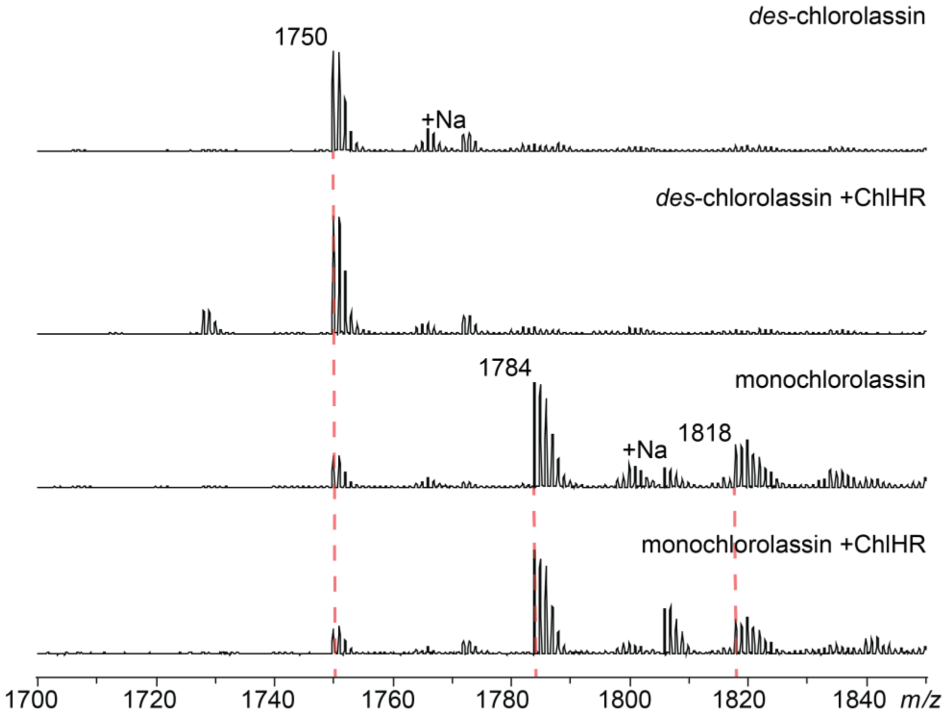


**Figure S2: *Des*-chlorolassin and monochlorolassin are not substrates for ChlH *in vitro*.** The lasso peptides were purified as described previously[12] and tested as described in the Methods section. After 24 h, no increase in chlorination was visible in either sample. The expected *m/z* for *des*-chlorolassin is 1750, and 1784 for monochlorolassin.

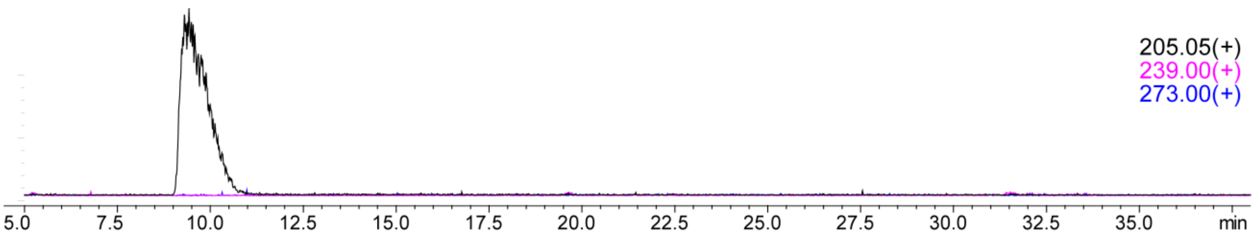


**Figure S3: Free tryptophan was not chlorinated by ChlH *in vitro*.** Extracted ion chromatograms for unmodified, monochlorinated, and dichlorinated Trp are shown in black, pink, and blue, respectively.

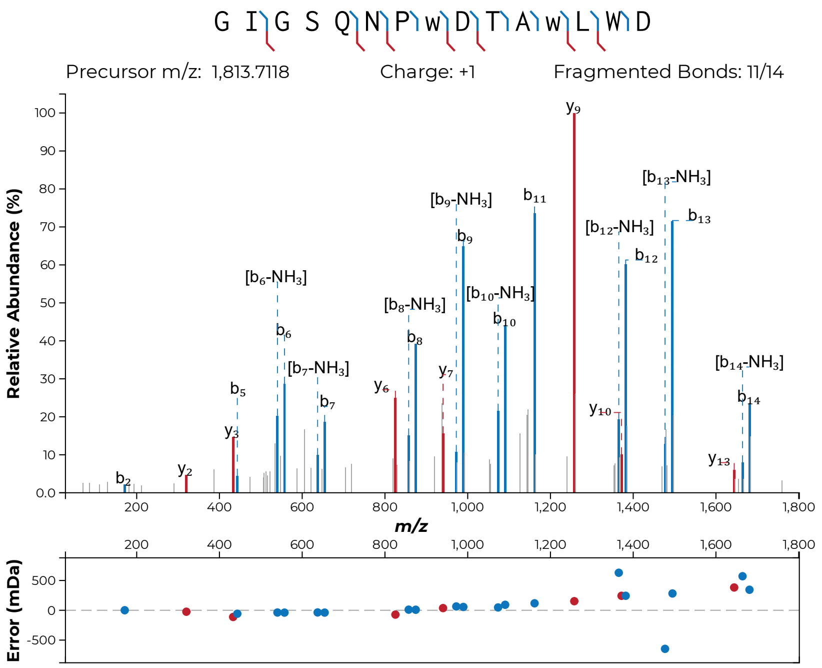


**Figure S4: MALDI-LIFT-MS of dichlorinated ChlA.** Following reaction of ChlA with ChlH/R for 2 h, MS/MS analysis indicated that ChlA Trp8 and Trp12 were both modified to yield a dichlorinated peptide. Chlorinated Trp is indicated in lowercase "w".

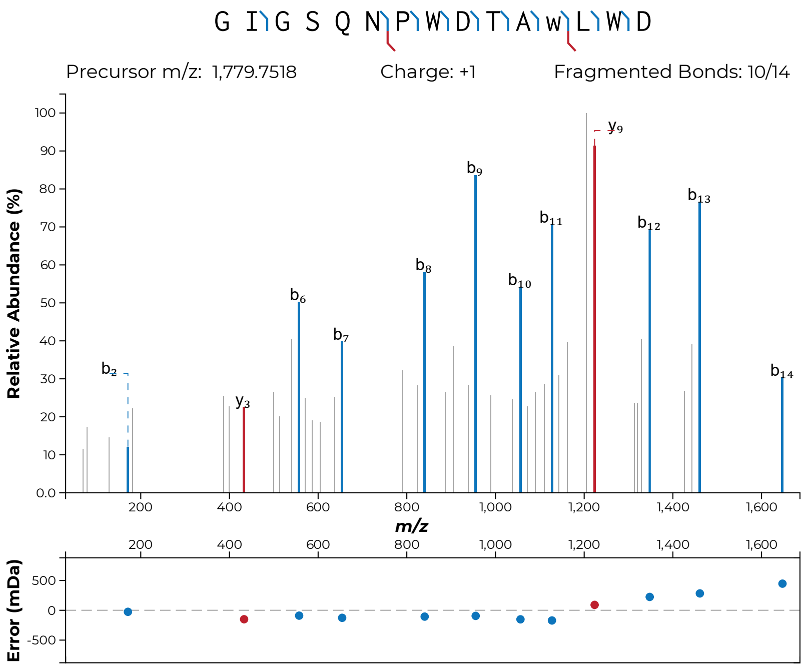


**Figure S5: MALDI-LIFT-MS to assess the order of chlorination on ChlA.** Following reaction of ChlA with ChlH/R for 20 min, MS/MS analysis indicated that the monochlorinated species originated from Trp12 chlorination. Chlorinated Trp is indicated in lowercase “w”.

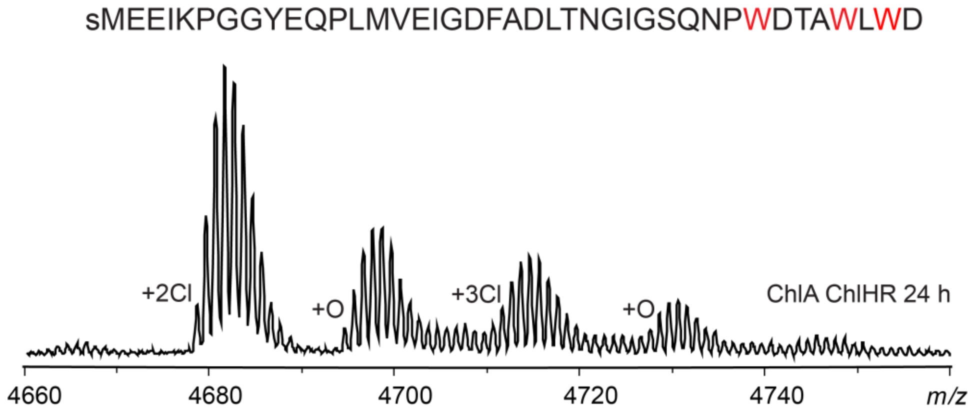


**Figure S6: MALDI-TOF-MS of ChlA/H/R after a 24 h reaction.** MBP-ChlA was partially trichlorinated after a 24 h reaction with ChlH/R *in vitro*. +O indicates presumed substrate oxidation at Met or Trp. Expected *m/z* values for the unmodified, mono-, di-, and trichlorinated peptides are 4610.1, 4644.1, 4678.0, and 4712.0, respectively. The lowercase "s" in the amino acid sequence represents a non-native Ser resulting from TEV protease cleavage of MBP-ChlA.

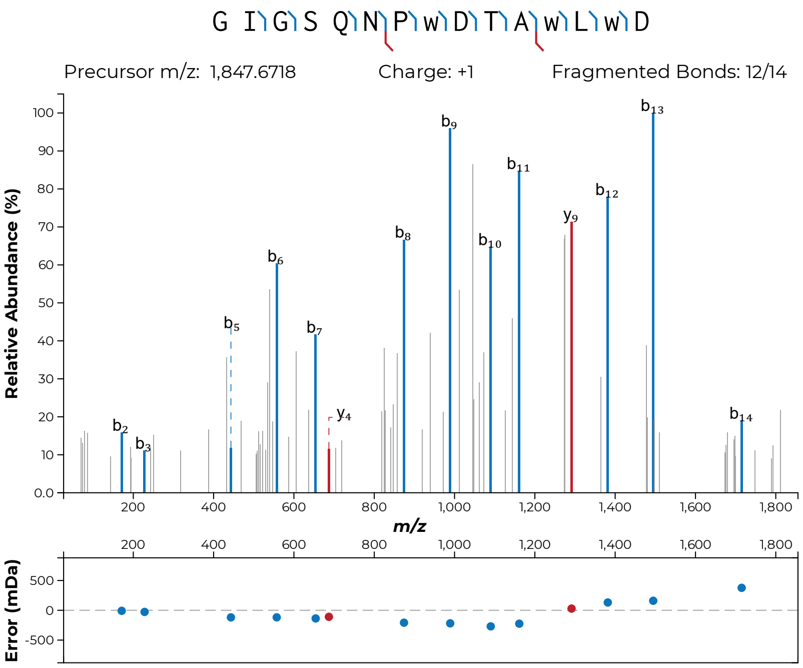


**Figure S7: MALDI-LIFT-MS of trichlorinated ChlA.** MBP-ChlA was partially trichlorinated after a 24 h reaction with ChlH/R *in vitro*. To increase production of the trichlorinated species, this reaction was repeated to a 48 h timepoint. Following FusB/E digestion, MS/MS performed on the trichlorinated species indicated single chlorination at each of the three Trp residues. Chlorinated Trp is indicated in lowercase "w".

```
CLUSTAL multiple sequence alignment by MUSCLE (3.8)


ChlH            MKMNAGRPSALDELVQTFSDDESAAVREWLGGAADAPQLPGISLRSNHEPLVEMLPRPAV
RebH            ---MSGK-----------------------------------------------------
                    :*.

ChlH            DDQKAIRRVAVVGGGTAGYLTAIAL-RTKRPWLEVSLIESPSIPIIGVGEATVPGIVIFL
RebH            -----IDKILIVGGGTAGWMAASYLGKALQGTADITLLQAPDIPTLGVGEATIPNLQTAF
                     * .: :*******:::*  * .: .   :::*:::*.** :******:*.:   :

ChlH            HHYLGIDVEDFYRQVRPTWKQGIRFEWGPRPDGFMAPFDWDSNSIGTAGAVDAGGGID--
RebH            FDFLGIPEDEWMRECNASYKVAIKF------------INWRTAGEGTSEARELDGGPDHF
                . :***  ::: *: ..::* .*.*            ::* : . **: * : .** *

ChlH            -------------------------GYTLQSLLMSAERAPVF---DVGDRHLSLMKYLPF
RebH            YHSFGLLKYHEQIPLSHYWFDRSYRGKTVEPFDYACYKEPVILDANRSPRRLDGSKVTNY
                                         * *::.:  :. . **:   : . *.*.  *   :

ChlH            AYHLDNARFVSYLADIAKQR-GVQHVEAKIEDVALSGDDWVDHLHTTDGRKLSYDFYVDC
RebH            AWHFDAHLVADFLRRFATEKLGVRHVEDRVEHVQRDANGNIESVRTATGRVFDADLFVDC
                *:*:*   ...:*  :*.:. **.*** .:* *  ..:..:: :.*: ** :. *::***

ChlH            SGFRSMLLGKAFRTPFHSYASSLFTDSAVTGNVGHGGN---LKPYTSAITMNSGWCWDIP
RebH            SGFRGLLINKAMEEPFLDMSDHLLNDSAVATQVPHDDDANGVEPFTSAIAMKSGWTWKIP
                ****.:*:.**:  ** . :. *:.****: :* *..:   ::*:****:*:*** *.**

ChlH            TPQDDHLGYVYSSNAISDDQAAAELAARF---PGVSEPRFVRFRVGRHEKAWRGNVMAIG
RebH            MLGRFGTGYVYSSRFATEDEAVREFCEMWHLDPETQPLNRIRFRVGRNRRAWVGNCVSIG
                       ******.  ::*:*. *:.  :   * ..  . :******: .** ** ::**

ChlH            NSYAFVEPLESTGILMITSSVQALVASLPGSWADPQCRDMVNVGLANRWDALRWFLSIHY
RebH            TSSCFVEPLESTGIYFVYAALYQLVKHFPDKSLNPVLTARFNREIETMFDDTRDFIQAHF
                .* .********** :: :::  **  :*..  :*     .*  : . :*  * *:. *:

ChlH            RFNTRRDTPFWREVRANTDISGAQPMLDMYATG----------APLRYRN--PLLRTFVQ
RebH            YFSPRTDTPFWRANKELRLADGMQEKIDMYRAGMAINAPASDDAQLYYGNFEEEFRNFWN
                 *..* ******  .     .* *  :*** :*          * * * *    :*.* :

ChlH            GNAPTFYGLAGVETILLGQQVPTRLLHRAEPPARWQARRNAADAFVRRALPQLEALSRFH
RebH            -NSNYYCVLAGLG--LVPDAPSPRLAHMPQATESVDEVFGAVKDRQRNLLETLPSLHEF-
                 *:  :  ***:   *: :  ..** * .:..   :   .*..   *. *  * :*  *

ChlH            DDPALNQQLLFDADSWASPLRAEPIGML
RebH            ----LRQQH-----------------GR-
                    *.**                 *
```

**Figure S8: Sequence alignment of RebH and ChlH.** Amino acid sequences for ChlH (NCBI accession code: WP_090100303.1) and RebH (CAC93722.1) were aligned using MUSCLE.[13] The catalytically critical Lys79 residue of RebH and the analogous ChlH Lys139 are shown in red.[14]

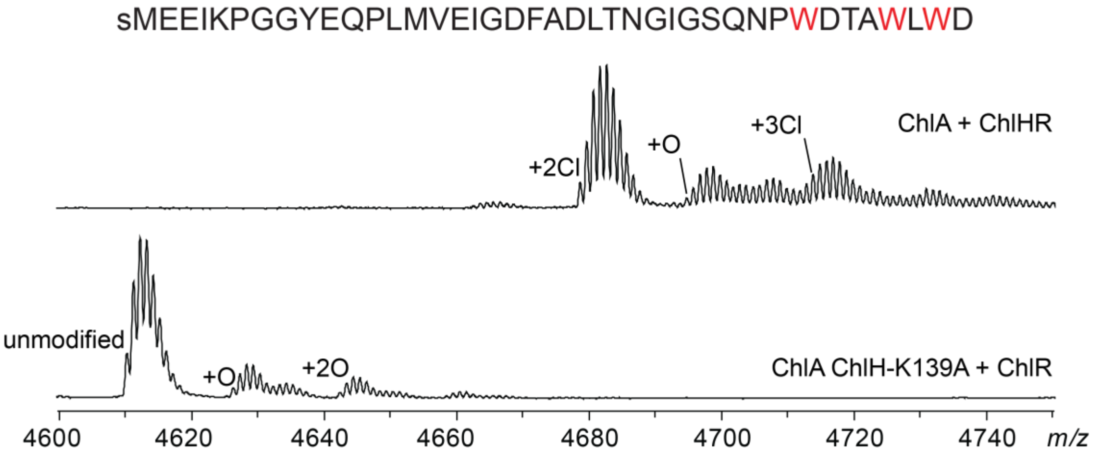


**Figure S9: MALDI-TOF-MS for ChlH-K139A variant activity.** ChlH-K139A was unable to chlorinate ChlA after 24 h of reaction with ChlR. +O indicates presumed substrate oxidation at Met or Trp. Expected *m/z* values for the unmodified, mono-, di-, and trichlorinated peptides are 4610.1, 4644.1, 4678.0, and 4712.0, respectively. The lowercase “s” in the amino acid sequence represents a non-native Ser resulting from TEV protease cleavage of MBP-ChlA.

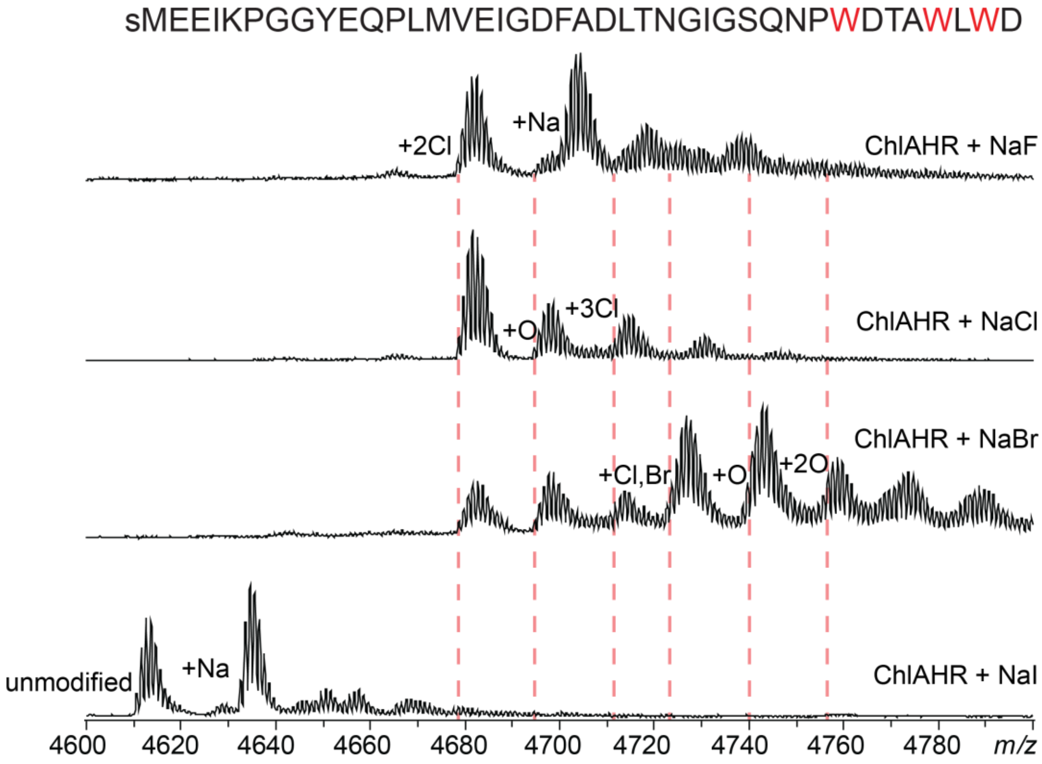


**Figure S10: MALDI-TOF-MS for ChlH halogen scope reactions.** Reactions were run as described in the "Halogen assessment" section of the Methods. Residual chloride (~50 mM) was sufficient for chlorination in the NaF reaction. $His_6$-ChlH precipitated in NaI, preventing further analysis. ChlA was successfully chlorinated and brominated in a NaBr-containing reaction buffer. +O indicates presumed substrate oxidation at Met or Trp. Expected *m/z* values for the unmodified, mono-, di-, and trichlorinated peptides are 4610.1, 4644.1, 4678.0, and 4712.0, respectively. Expected *m/*z values for the mono- and dibrominated peptides are 4688.0 and 4766.9, respectively. The expected *m/*z value for the monobrominated and monochlorinated peptide is 4723.0. The lowercase "s" in the amino acid sequence represents a non-native Ser resulting from TEV protease cleavage of MBP-ChlA.

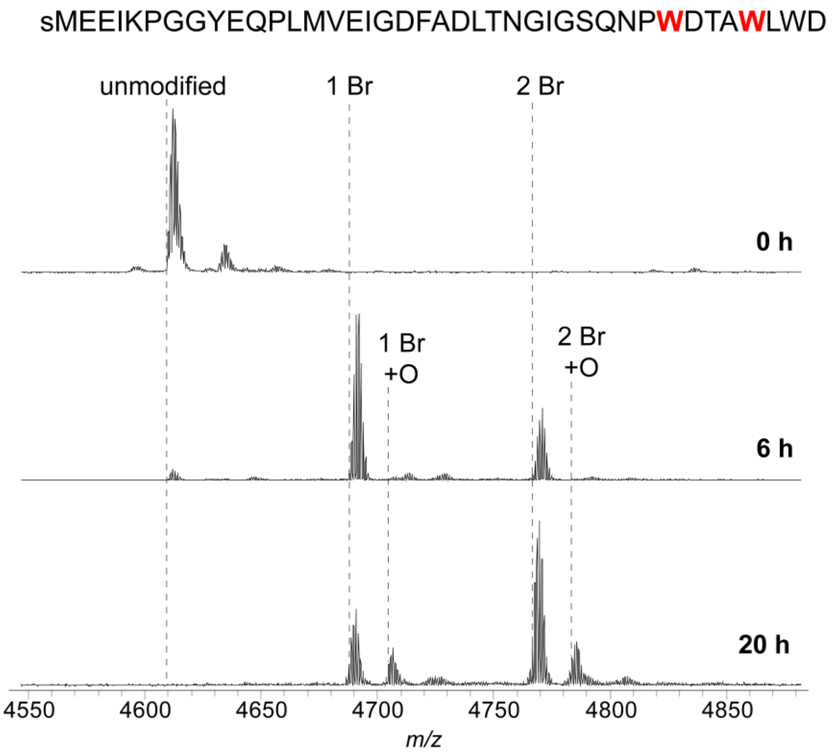


**Figure S11: MALDI-TOF-MS analysis of selective ChlA bromination.** More rigorous removal of chloride by buffer exchange allowed for selective dibromination of ChlA. +O indicates presumed substrate oxidation at Met or Trp. Expected *m/z* values for unmodified, mono-, and dibrominated peptides are 4609.1, 4688.0, and 4766.9, respectively. The lowercase "s" in the amino acid sequence represents a non-native Ser resulting from TEV protease cleavage of MBP-ChlA.

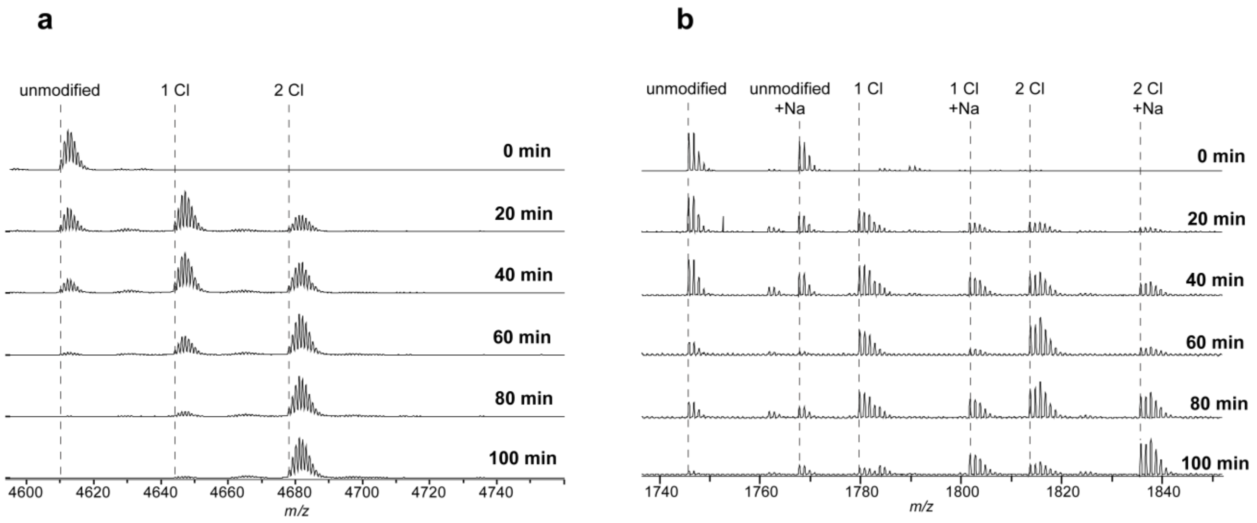


**Figure S12: MALDI-TOF-MS for ChlA vs ChlA$_{core}$ halogenation reactions.** Relative rates of ChlH processing. (a) MALDI-TOF-MS monitoring of ChlA precursor peptide and (b) ChlA$_{core}$. The reactions were identical except for the tested substrate. Expected *m/z* values for unmodified, mono-, and dibrominated full-length ChlA peptides are 4609.1, 4688.0, and 4766.9, respectively, while unmodified, mono-, and dibrominated ChlA$_{core}$ are 1745.8, 1779.8, and 1813.7, respectively. $[M+Na]^+$ peaks are labeled where observed.

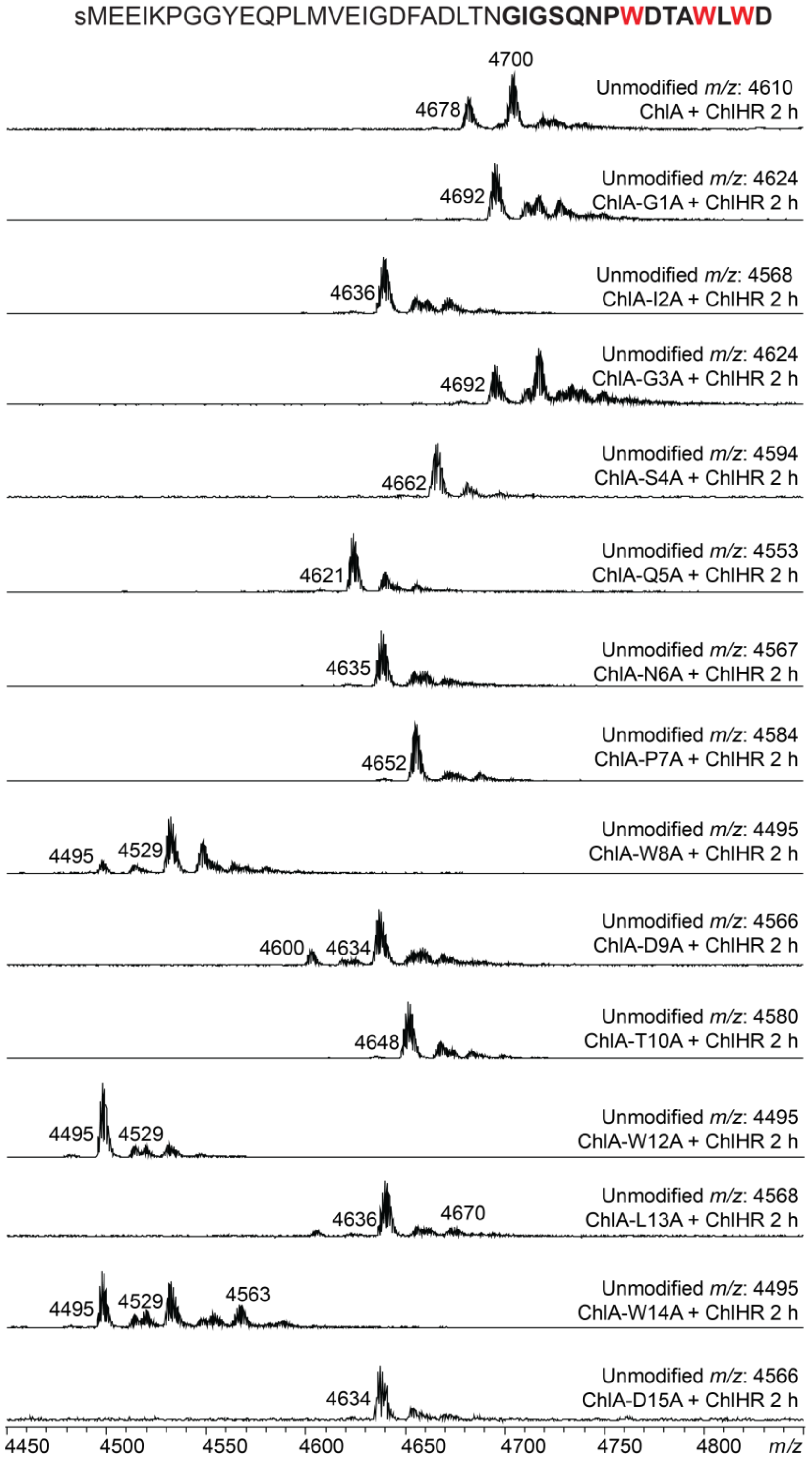


**Figure S13: MALDI-TOF-MS for ChlA Ala variants at 2 h.** MALDI-TOF-MS analysis for Ala-substituted variants of ChlA after a 2 h reaction. The *m/z* for unmodified peptide is supplied, and chlorinated products are labeled. The lowercase “s” in the amino acid sequence represents a non-native Ser resulting from TEV protease cleavage of MBP-ChlA.

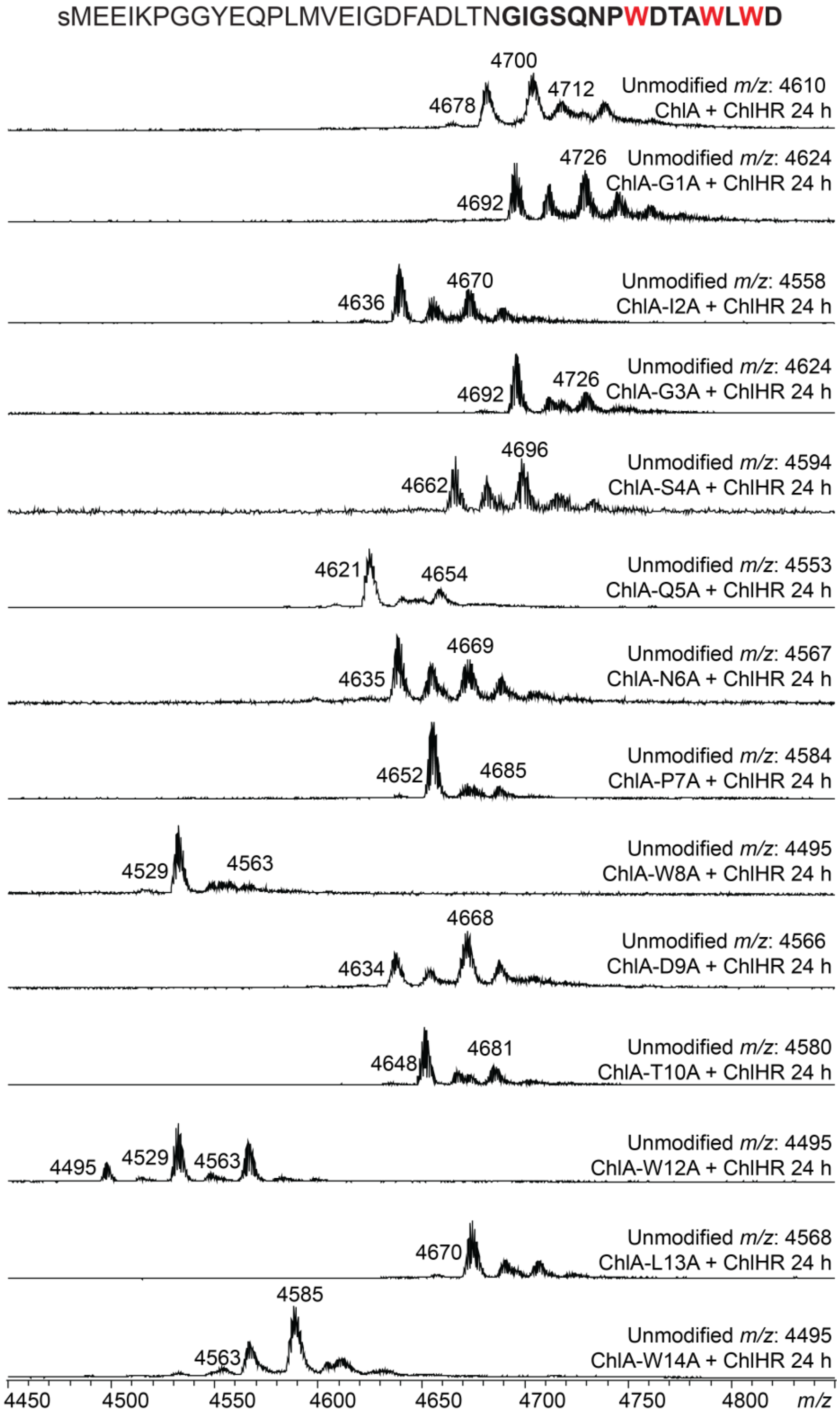


**Figure S14: MALDI-TOF-MS for ChlA Ala variants at 24 h.** MALDI-TOF-MS analysis for Ala-substituted variants of ChlA after a 24 h reaction. The *m/z* for unmodified peptide is supplied, and chlorinated products are labeled. The lowercase “s” in the amino acid sequence represents a non-native Ser resulting from TEV protease cleavage of MBP-ChlA.

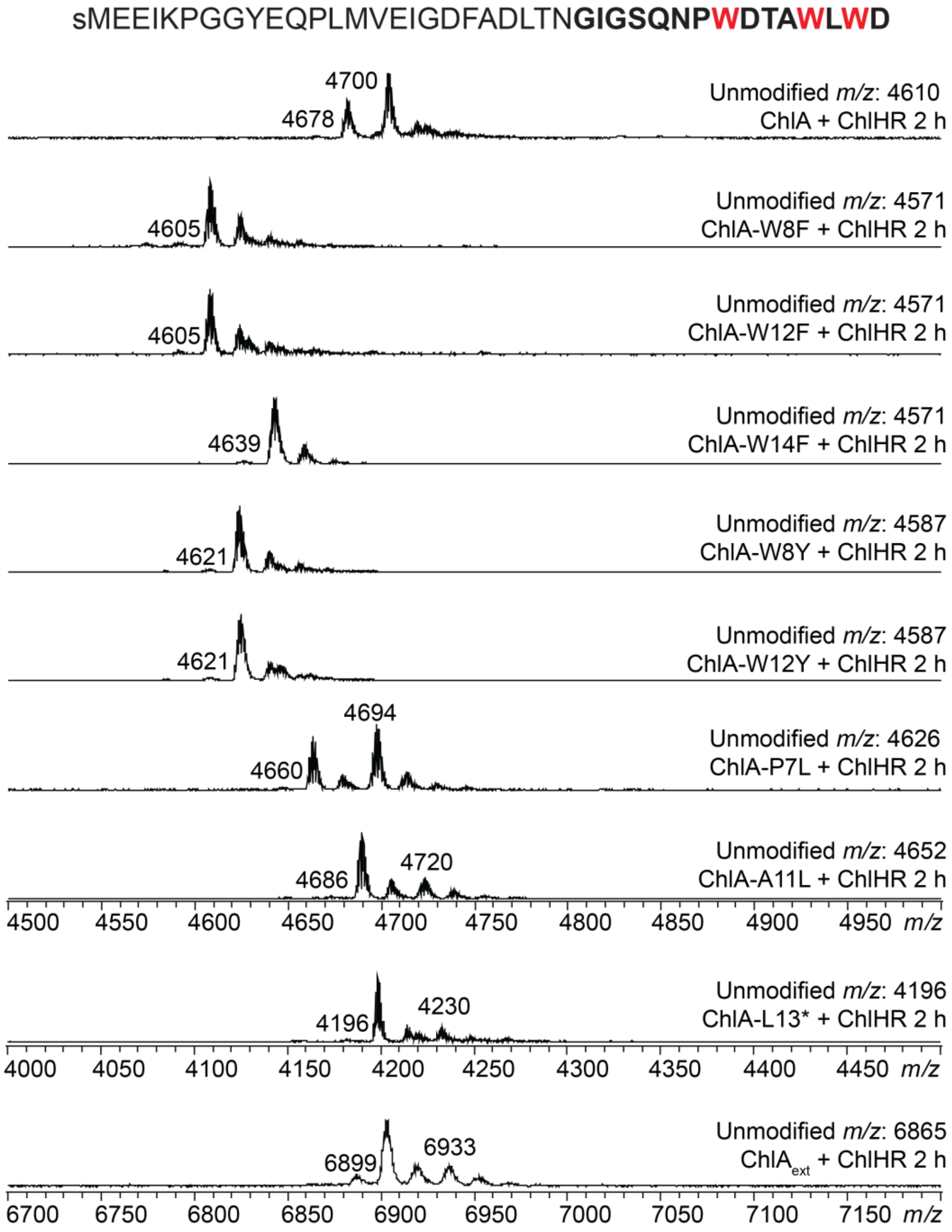


**Figure S15: MALDI-TOF-MS for additional ChlA variants at 2 h.** MALDI-TOF-MS analysis for additional variants of ChlA after a 2 h reaction. The *m/z* for unmodified peptide is supplied, and chlorinated products are labeled. The lowercase "s" in the amino acid sequence represents a non-native Ser resulting from TEV protease cleavage of MBP-ChlA. The sequence of $ChlA_{ext}$, in which several residues have been added to the C-terminus of ChlA, is provided in Figure S16.

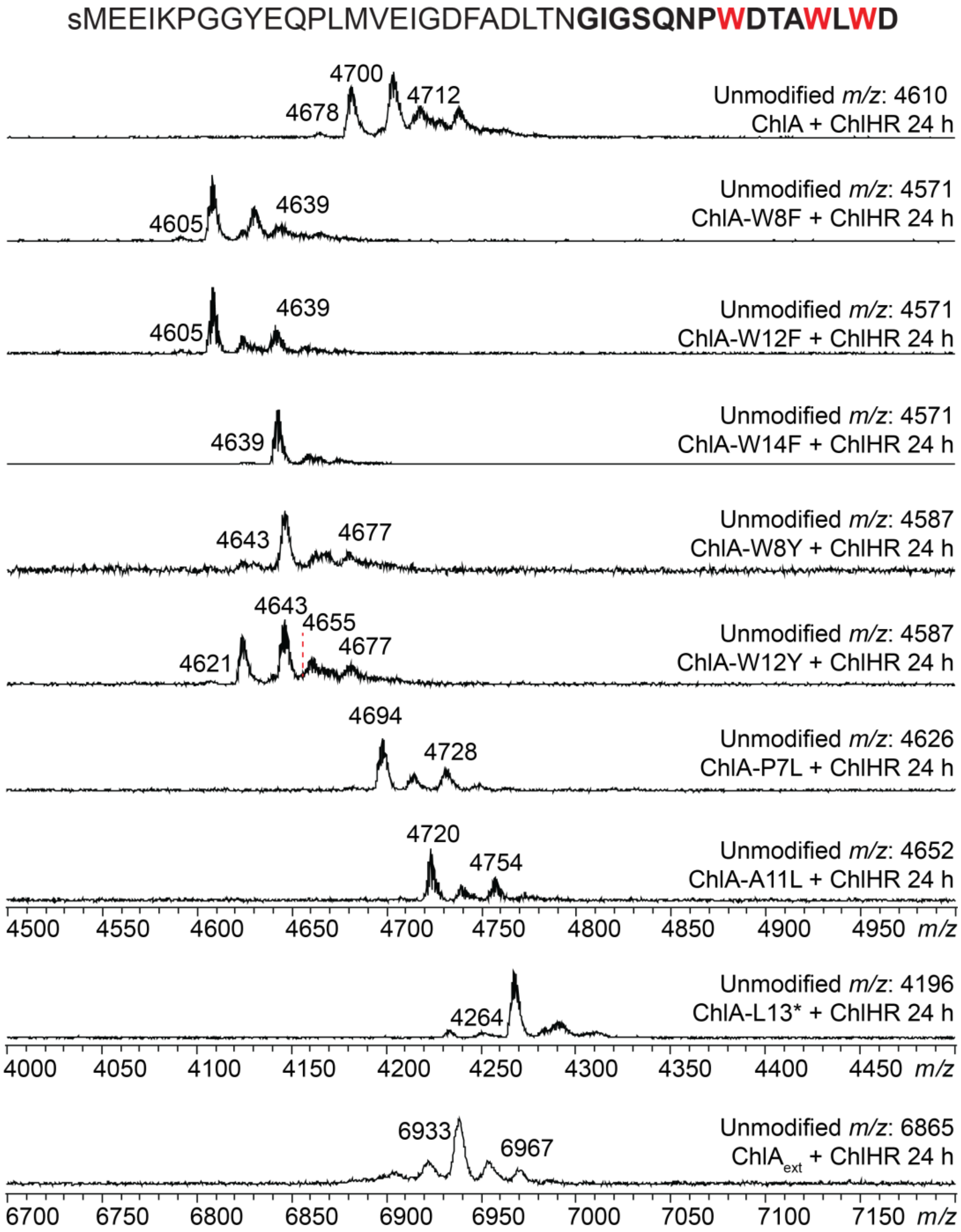


**Figure S16: MALDI-TOF-MS for additional ChlA variants at 24 h.** MALDI-TOF-MS analysis for additional variants of ChlA after a 24 h reaction. The *m/z* for unmodified peptide is supplied, and chlorinated products are labeled. The lowercase "s" in the amino acid sequence represents a non-native Ser resulting from TEV protease cleavage of MBP-ChlA. The sequence of ChlA$_{ext}$, in which several residues have been added to the C-terminus of ChlA, is provided in Figure S16.

ChlA$_{ext}$: sMEEIKPGGYEQPLMVEIGDFADLTNGIGSQNPWDTAWLWDRIPRVDKLAAALEHHHHHH

**Figure S17: Sequence of ChlA$_{ext}$.** Shown is the substrate sequence for ChlA$_{ext}$, which includes a non-native C-terminal "extension" shown in blue. The lowercase "s" in the amino acid sequence represents a non-native Ser resulting from TEV protease cleavage of MBP-ChlA. The MS data deriving from reactions on ChlA$_{ext}$ are in Figures S15-S16.

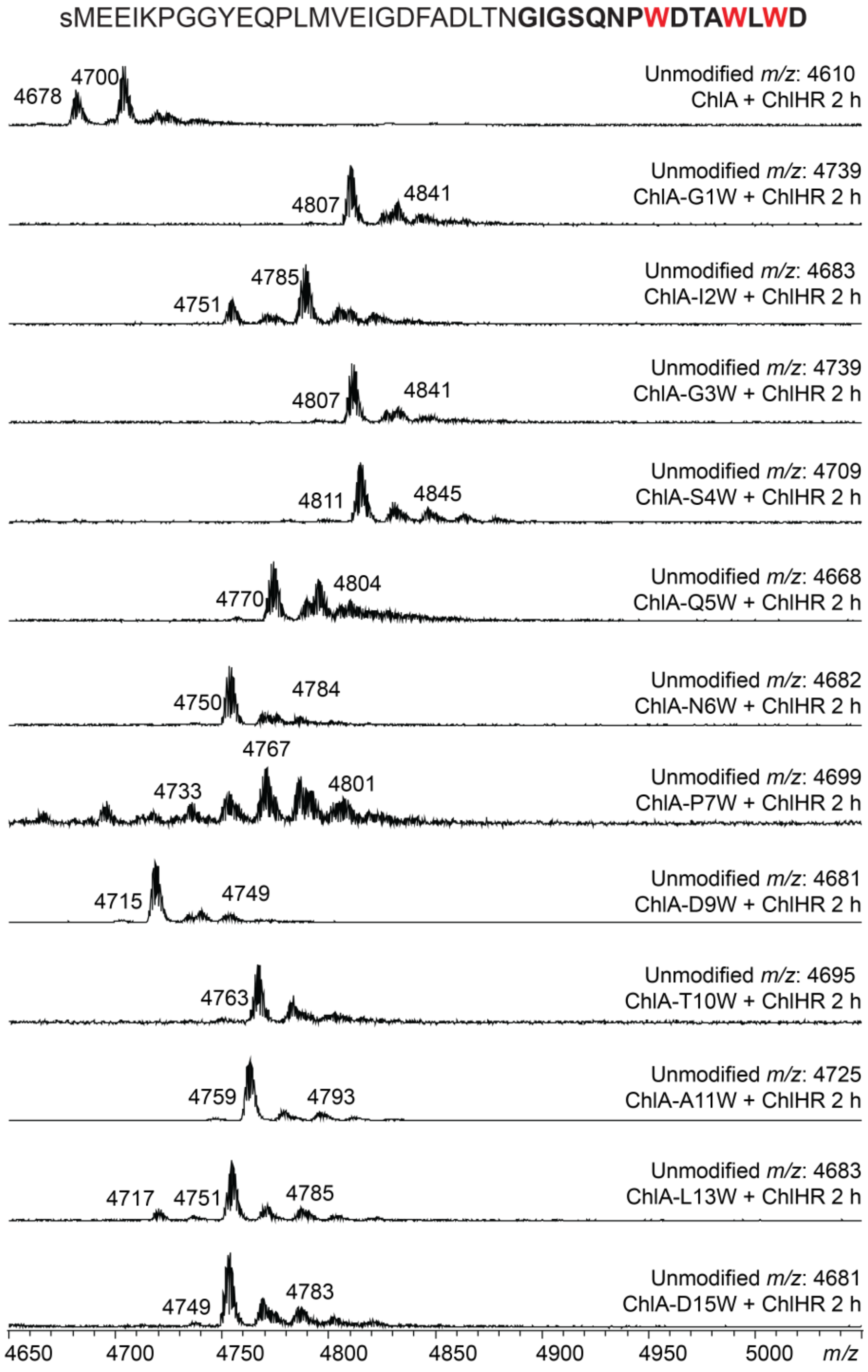


**Figure S18: MALDI-TOF-MS for ChlA Trp variants at 2 h.** MALDI-TOF-MS analysis for Trp-substituted variants of ChlA after a 2 h reaction. The *m/z* for unmodified peptide is supplied, and chlorinated products are labeled. The lowercase “s” in the amino acid sequence represents a non-native Ser resulting from TEV protease cleavage of MBP-ChlA.

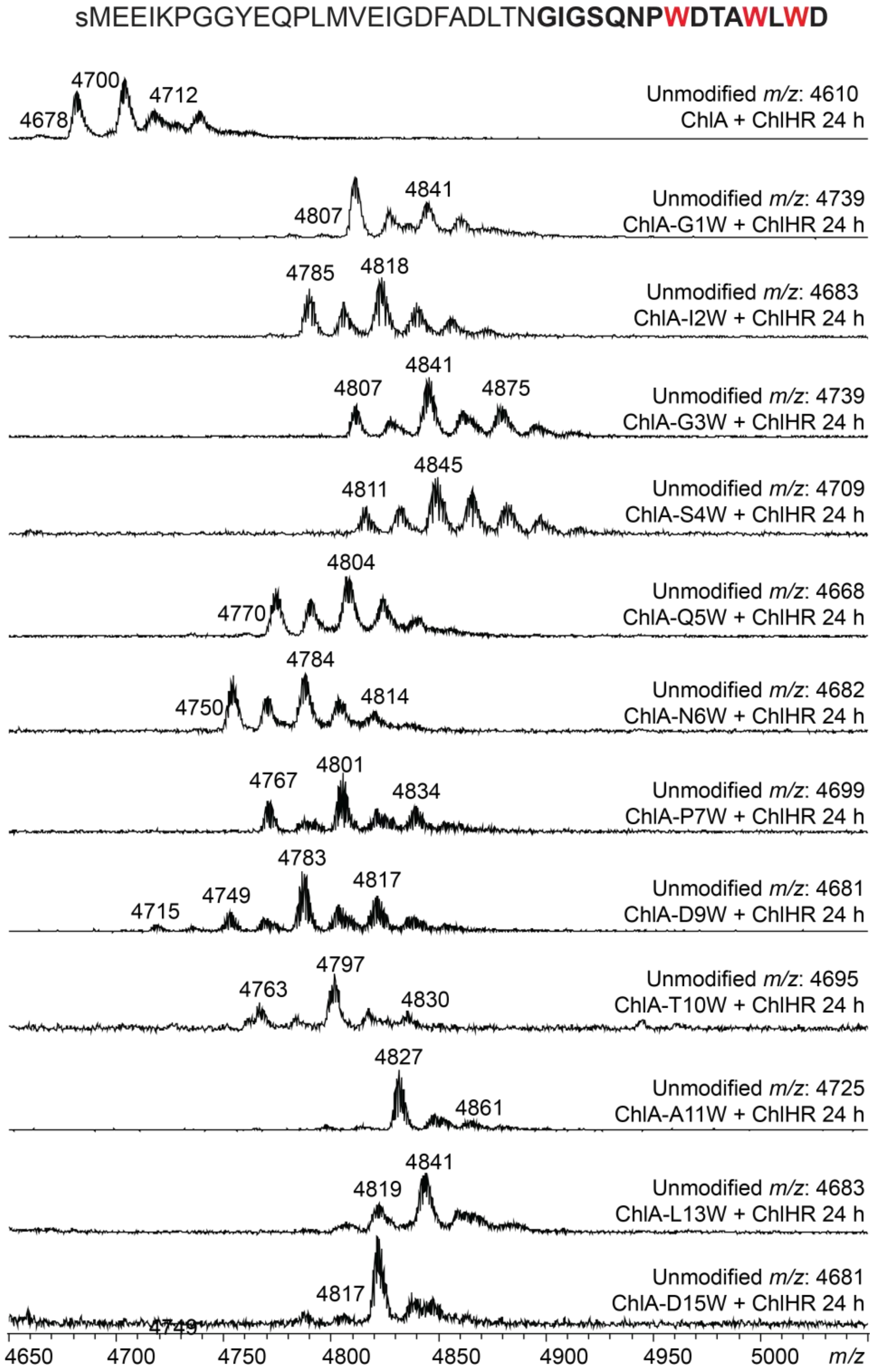


**Figure S19: MALDI-TOF-MS for ChlA Trp variants at 24 h.** MALDI-TOF-MS analysis for Trp-substituted variants of ChlA after a 24 h reaction. The *m/z* for unmodified peptide is supplied, and chlorinated products are labeled. The lowercase "s" in the amino acid sequence represents a non-native Ser resulting from TEV protease cleavage of MBP-ChlA.

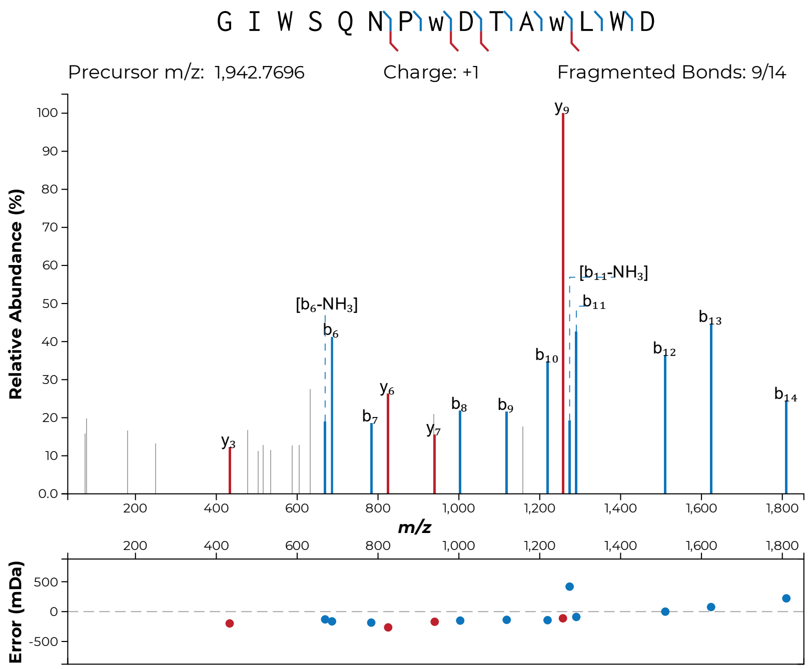


**Figure S20: MALDI-LIFT-MS for ChlA-G3W.** After ChlA-G3W was subjected to a 2 h reaction with ChlH/R, MS/MS indicated that the predominant dichlorinated species originated from Trp8 and Trp12 chlorination. Chlorinated Trp is indicated in lowercase "w".

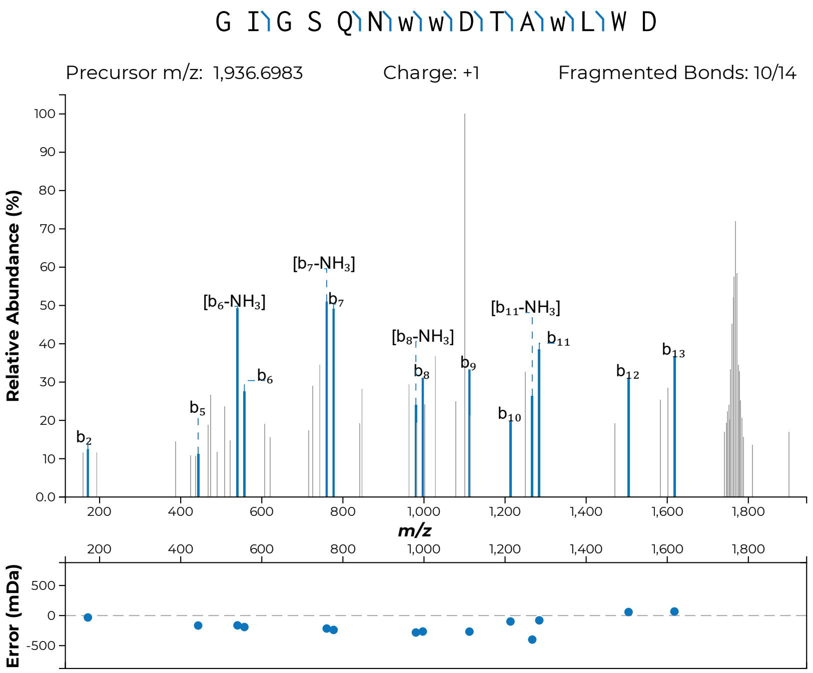


**Figure S21: MALDI-LIFT-MS for ChlA-P7W.** After ChlA-P7W was subjected to a 24 h reaction with ChlH/R, MS/MS indicated that the predominant trichlorinated species originated from Trp7, Trp8, and Trp12 chlorination. Chlorinated Trp is indicated in lowercase "w".

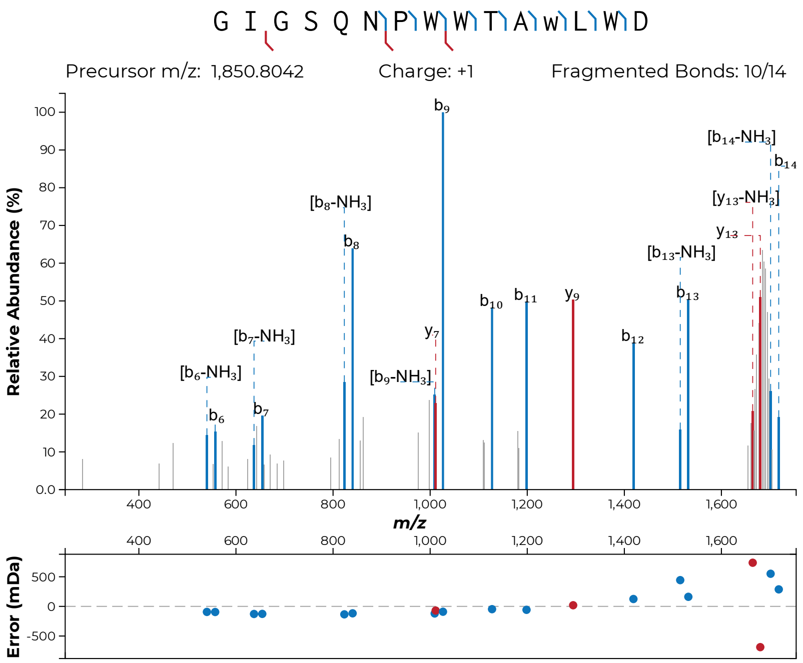


**Figure S22: MALDI-LIFT-MS for ChlA-D9W.** After ChlA-D9W was subjected to a 2 h reaction with ChlH/R, MS/MS indicated that the predominant monochlorinated species originated from Trp12 chlorination. Chlorinated Trp is indicated in lowercase "w".

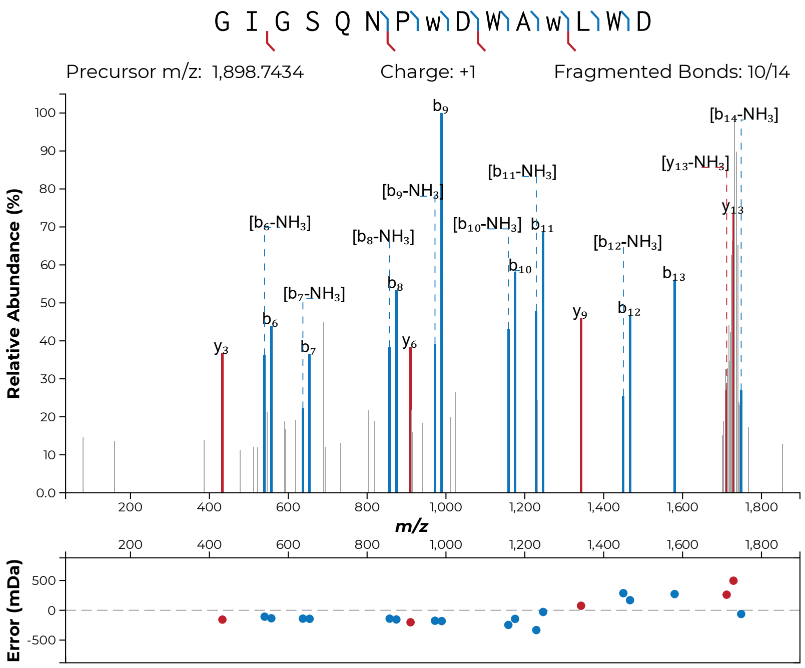


**Figure S23: MALDI-LIFT-MS for ChlA-T10W.** After ChlA-T10W was subjected to a 2 h reaction with ChlH/R, MS/MS indicated that the predominant dichlorinated species originated from Trp8 and Trp12 chlorination. Chlorinated Trp is indicated in lowercase "w".

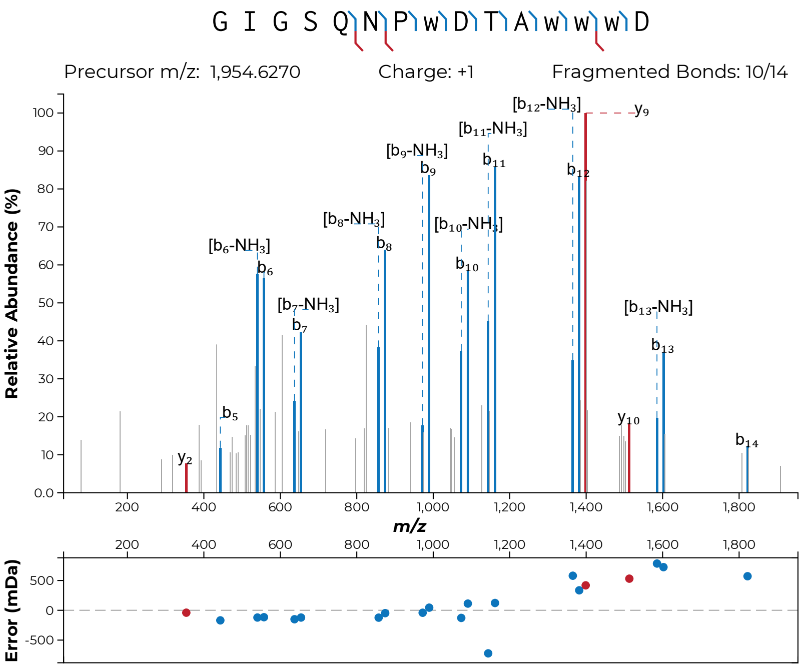


**Figure S24: MALDI-LIFT-MS for ChlA-L13W.** After ChlA-L13W was subjected to a 24 h reaction with ChlH/R, MS/MS indicated that the predominant tetrachlorinated species originated from Trp8, Trp12, Trp13, and Trp14 chlorination. Chlorinated Trp is indicated in lowercase "w".

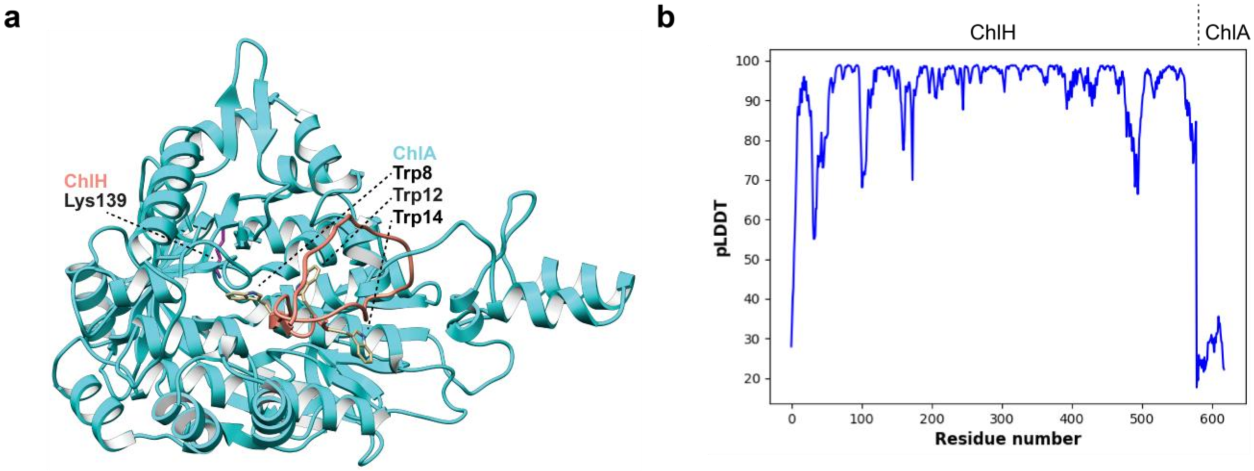


**Figure S25: AlphaFold 2 Multimer model of ChlA and ChlH.** (a) AlphaFold 2 Multimer was used to generate a model of ChlA (salmon) and ChlH (cyan). Relevant residues in ChlA and ChlH are highlighted. (b) pLDDT plot for the model, illustrating a highly uncertain placement of ChlA.

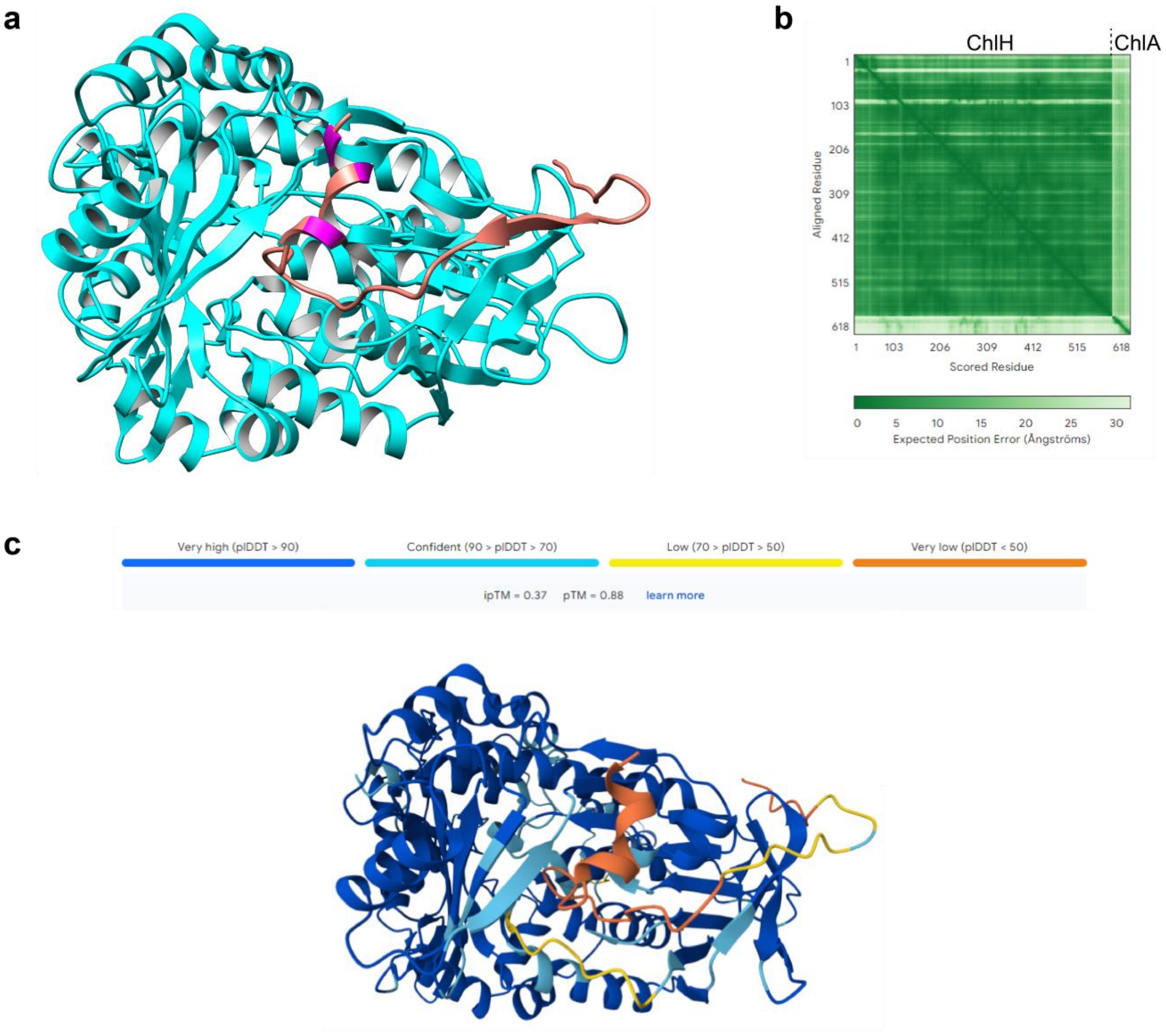


**Figure S26: AlphaFold 3 model of ChlA and ChlH.** (a) AlphaFold 3 was used to generate a model of ChlA (salmon, Trp residues in magenta) and ChlH (cyan).[15] (b) The positional error plot for our ChlA-ChlH model, illustrating that ChlA is not confidently modeled with respect to ChlH. (c) ChlA-ChlH model colored by pLDDT score, highlighting that ChlA is not confidently positioned with respect to ChlA. The core peptide of ChlA is entirely within the orange helix, indicative of very low confidence.

**Table S1. Molecular dynamics setups.**

| **Starting setup** | **# atoms** | **Time (μs)** |
|---|---|---|
| original AlphaFold pose rep 01 | 90,966 | 2.225 |
| original AlphaFold pose rep 02 | 90,966 | 2.500 |
| original AlphaFold pose rep 03 | 90,966 | 2.500 |
| Trp8 – optimized rep 01 | 90,966 | 1.675 |
| Trp8 – optimized rep 02 | 90,966 | 1.325 |
| Trp8 – optimized rep 03 | 90,966 | 1.325 |
| Trp12 –shifted rep 01 | 110,022 | 0.900 |
| Trp12 –shifted rep 02 | 110,022 | 0.750 |
| Trp12 –shifted rep 03 | 110,022 | 0.875 |
| Trp14–shifted rep 01 | 127,294 | 0.850 |
| Trp14–shifted rep 02 | 127,294 | 0.800 |
| Trp14–shifted rep 03 | 127,294 | 0.800 |
| Aggregated Time | | ~16.5 μs |

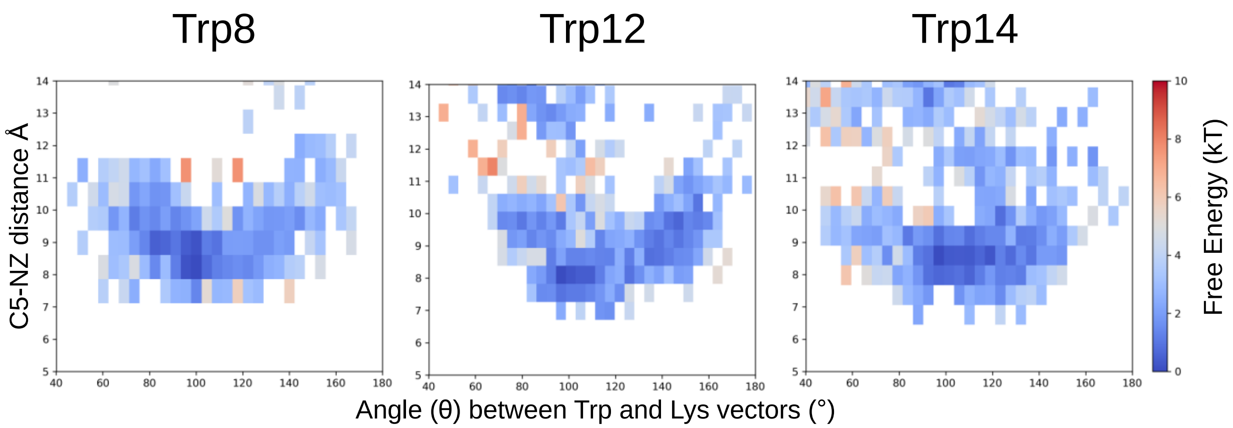


**Figure S27. Raw free energy surfaces.** Results shown for each Trp variant as a function of the distance between the Trp C5 atom and Lys139 NZ atom (y-axis) and the angle between the Trp (CZ3→CE2) and Lys (CE→NZ) vectors (x-axis). These surfaces were computed directly from raw WESTPA trajectory data using iteration-normalized weights, without kernel density estimation (KDE) smoothing. Low-energy basins indicate conformational states with favorable distance and angular alignment for halogen transfer. Trp8 and Trp12 show defined minima, consistent with catalytically competent configurations, while Trp14 samples more diffuse, higher-energy regions.

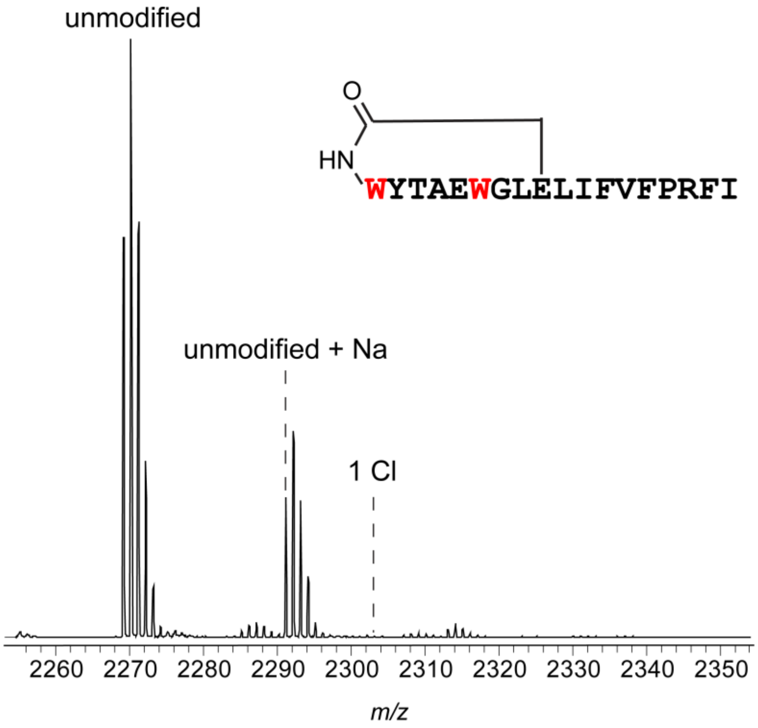


**Figure S28: MALDI-TOF-MS for attempted chlorination of fusilassin.** ChlH rejected fusilassin as a chlorination substrate. Fusilassin contains an isopeptide bond formed between the N-terminal amine and the carboxylate side chain of Glu9, with the remainder of the C-terminal tail threaded through the macrocycle.[7] The structure above is drawn in a non-threaded conformation for clarity. Expected *m/z*: unmodified (2269.2, observed), unmodified + Na (2291.2, observed), 1 Cl (2303.1, not observed), and 2 Cl (2337.1, not observed).

FusA$_{core}$: WYTAEWGLELIFVFPRFI

ChlA$_{core}$: GIGSQNPWDTAWLWD

**Figure S29: Sequence comparison of ChlA and FusA core regions.** The core peptides of ChlA and FusA are listed with Trp highlighted in red. The macrolactam-forming residues are underlined.

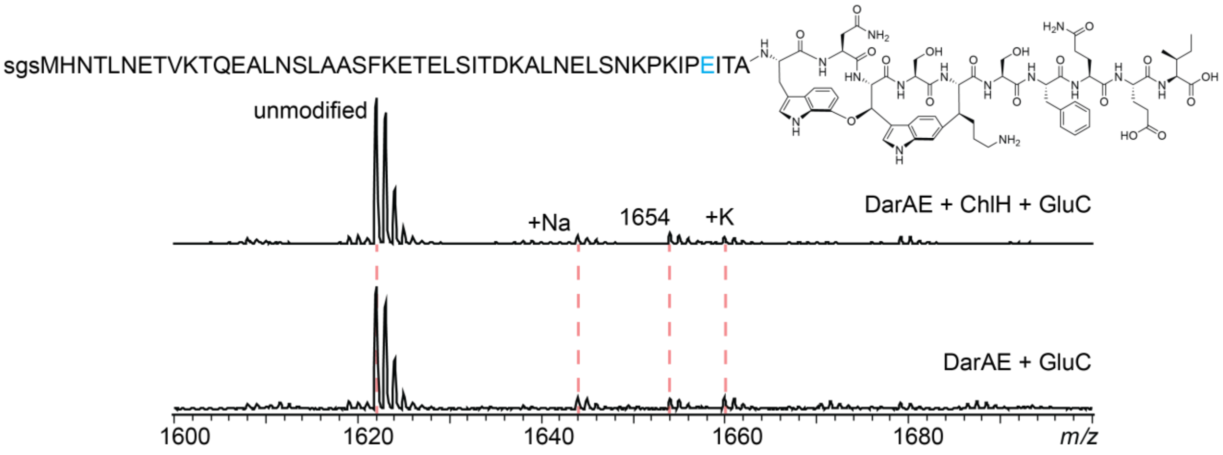


**Figure S30: MALDI-TOF-MS for attempted chlorination of a darobactin biosynthetic intermediate.** ChlH/R were allowed to react with the darobactin biosynthetic intermediate depicted above. The sample was then digested with 1:20 endoproteinase GluC:substrate for 16 h at 37 °C to yield a peptide fragment more amenable for MS-based detection (the analyzed fragment resulted from proteolysis at the blue glutamate residue shown on the sequence). No chlorination was observed. Expected *m/z* values for the unmodified and chlorinated products are 1621.8 and 1655.7, respectively.

a

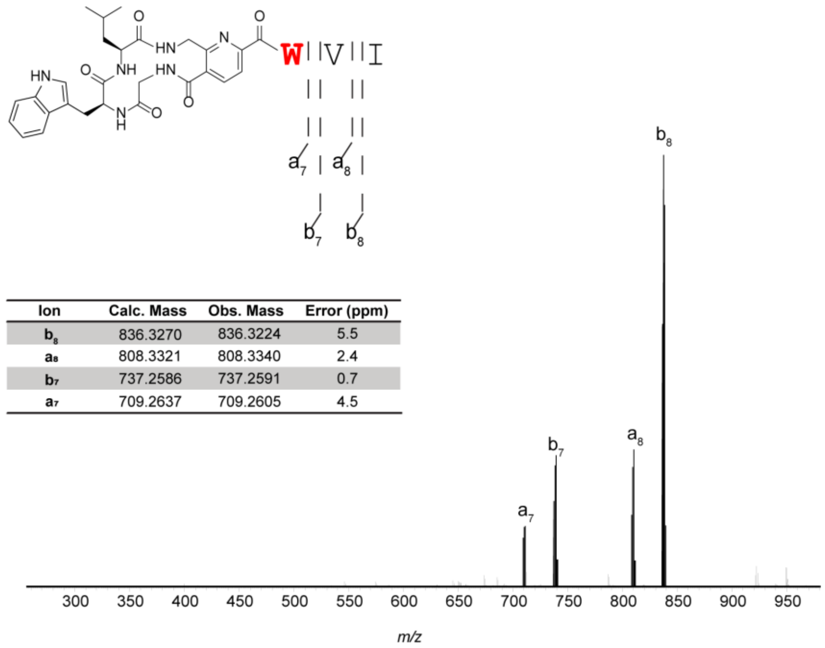


| Ion | Calc. Mass | Obs. Mass | Error (ppm) |
|---|---|---|---|
| $b_8$ | 836.3270 | 836.3224 | 5.5 |
| $a_8$ | 808.3321 | 808.3340 | 2.4 |
| $b_7$ | 737.2586 | 737.2591 | 0.7 |
| $a_7$ | 709.2637 | 709.2605 | 4.5 |

b

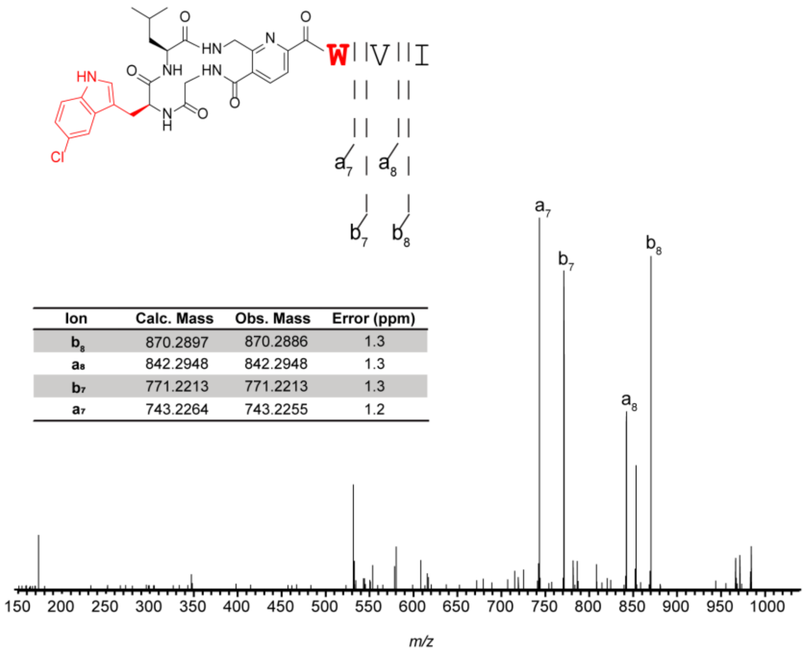


| Ion | Calc. Mass | Obs. Mass | Error (ppm) |
|---|---|---|---|
| $b_8$ | 870.2897 | 870.2886 | 1.3 |
| $a_8$ | 842.2948 | 842.2948 | 1.3 |
| $b_7$ | 771.2213 | 771.2213 | 1.3 |
| $a_7$ | 743.2264 | 743.2255 | 1.2 |

**Figure S31: LC-MS/MS of chlorinated pyritide A1.** HR-MS/MS of (a) monochlorinated and (b) dichlorinated pyritide A1 confirmed that two chlorinations occurred.

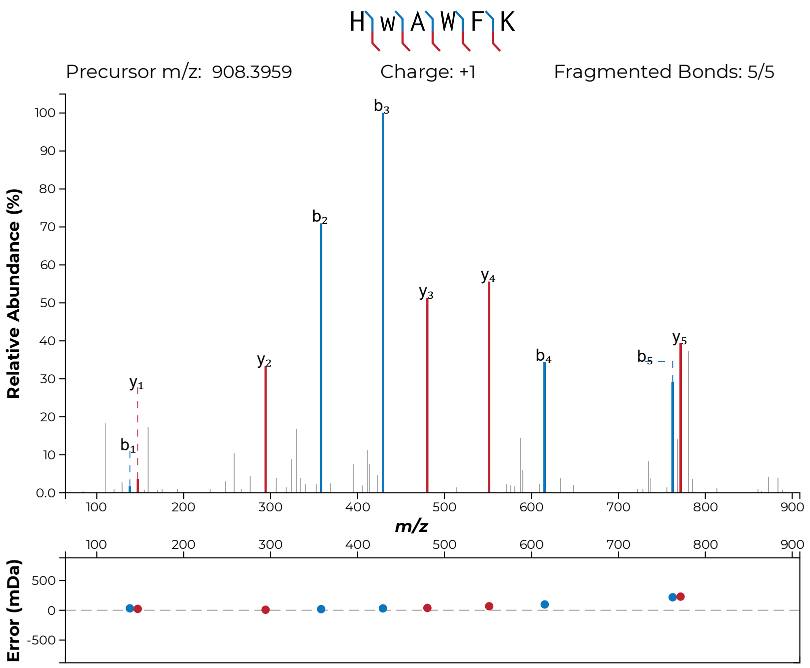


**Figure S32: MALDI-LIFT-MS for chlorinated GHRP-6.** After GHRP-6 was subjected to a 24 h reaction with ChlH/R, MS/MS indicated that the predominant monochlorinated species originated from *D*-Trp2 chlorination. Chlorinated Trp is indicated in lowercase “w”.

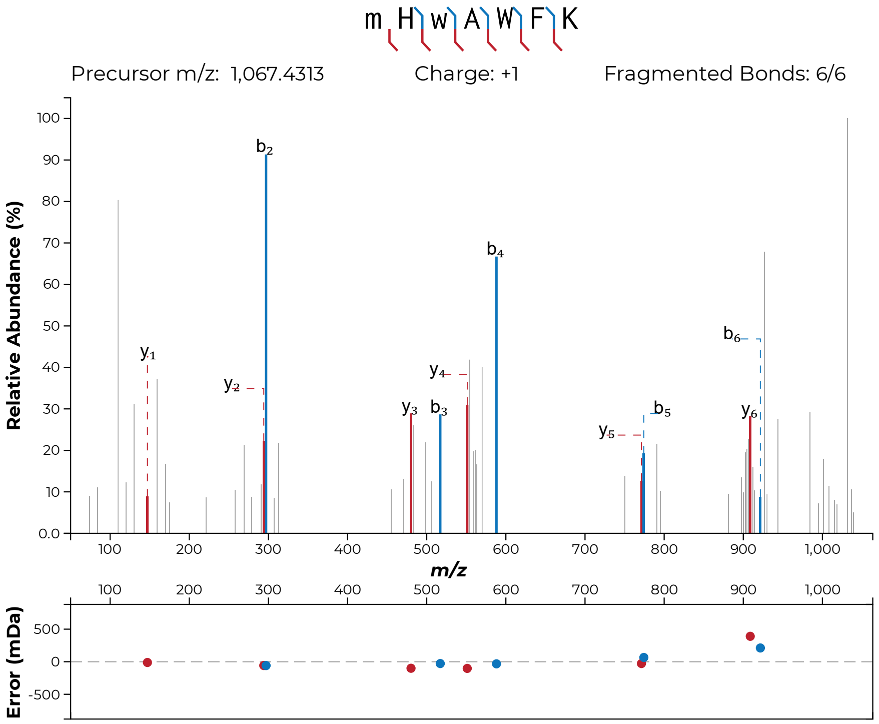


**Figure S33: MALDI-LIFT-MS for chlorinated all-*L*-amino acid GHRP-6.** Following post-translational halogenation of the PURExpress-translated GHRP-6 all-*L*-configured sequence, MS/MS indicated that the predominant monochlorinated species originated from *L*-Trp2 chlorination. Chlorinated Trp is indicated in lowercase "w".

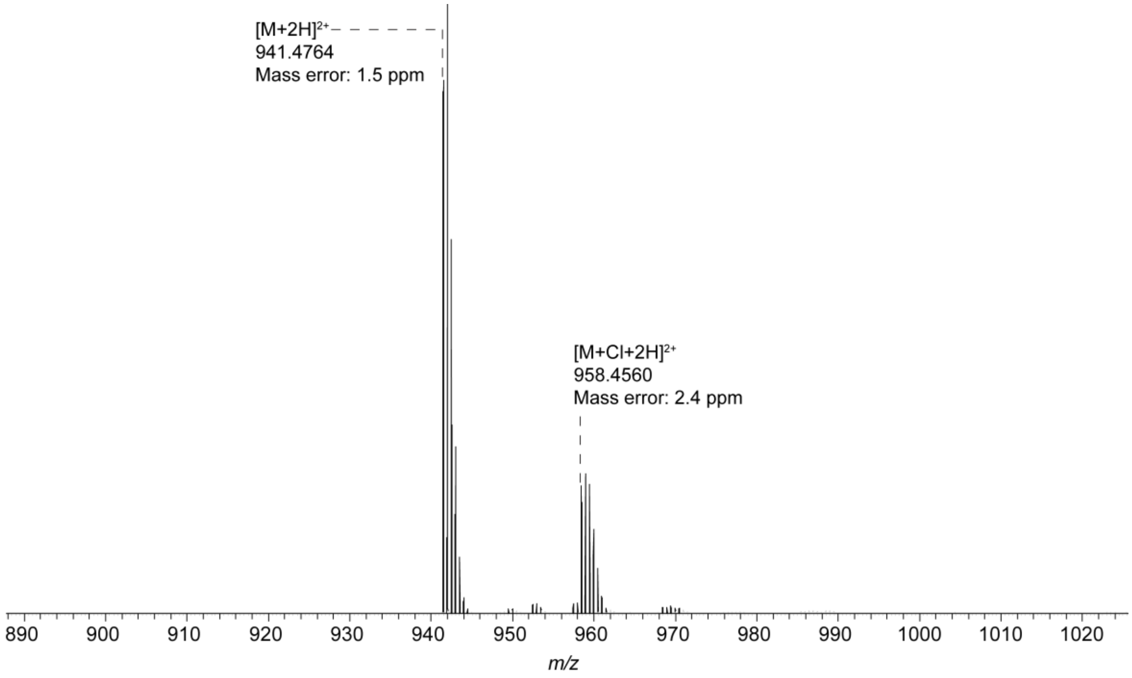


**Figure S34: HR-MS of WL12 peptide chlorination.** High-resolution mass spectrometry indicated that the new ion corresponded to a mass increase matching that of single chlorination, confirming the initial MALDI-TOF-MS result.

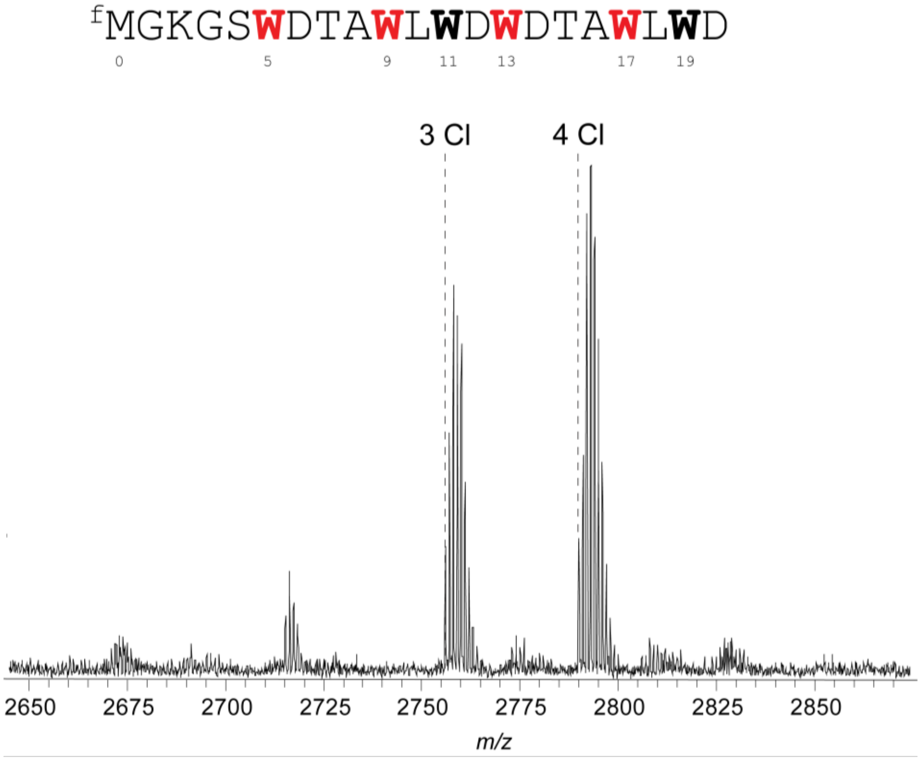


**Figure S35: MALDI-TOF-MS for ChlA$_{PE}$ 8-15 duplication substrate chlorination.** Upon addition of ChlH/R to the translated peptide, formation of tri- and tetrachlorinated species was observed within 1 h. Expected *m/z* values for 3 Cl and 4 Cl are 2756 and 2790, respectively. $^{f}$M indicates formylation of the translation initiating Met residue, a standard occurrence for PURExpress translated polypeptides. All Trp residues are bolded, with presumed sites of chlorination additionally colored red.

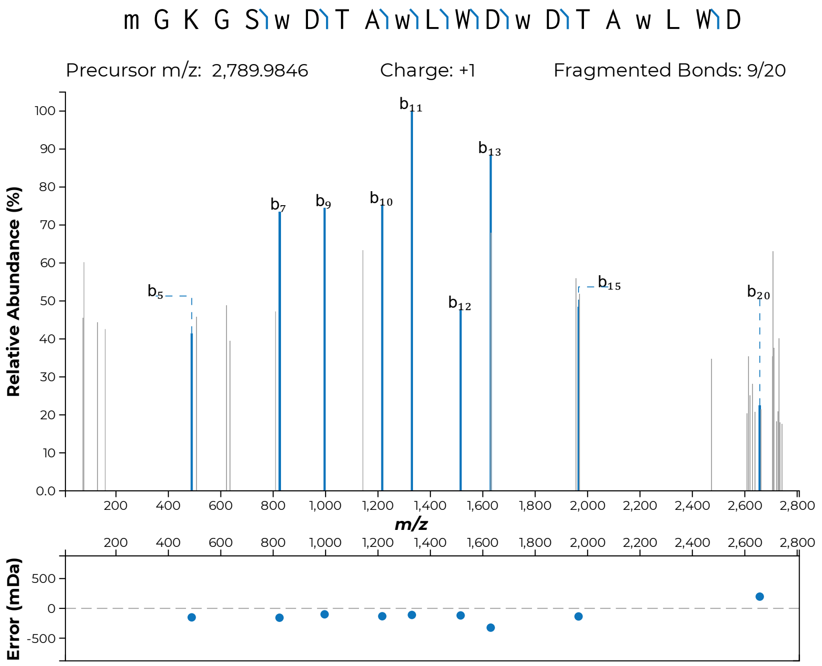


**Figure S36: MALDI-LIFT-MS for chlorinated ChlA$_{8\text{-}15}$ duplication substrate.** Following post-translational halogenation of the ChlA$_{8\text{-}15}$ duplication sequence, MS/MS indicated chlorinations was localized to Trp5, 9, 13, and 17 (although our assignment is not definitive). Lowercase letters indicate modified residues. For example, the N-terminal formylated Met is indicated as “m”. Chlorinated Trp is indicated as “w”.

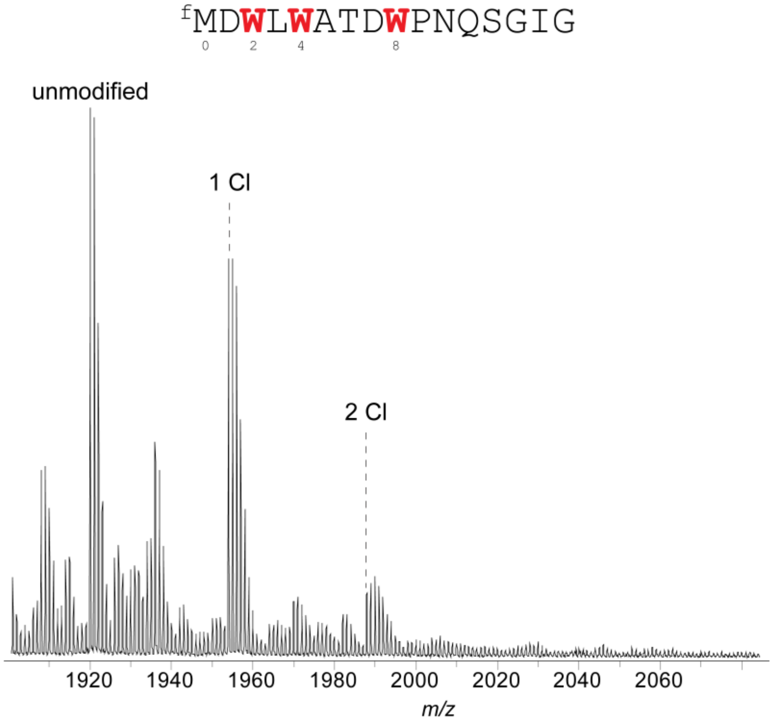


**Figure S37: MALDI-TOF-MS for ChlA$_{PE}$ inverted substrate chlorination.** Upon addition of ChlH/R to the translated peptide, mono- and dichlorination was observed within 1 h. Expected *m/z* values for unmodified, 1 Cl and 2 Cl are 1919.8, 1953.8 and 1987.8, respectively. $^{f}$M indicates formylation of the translation initiating Met residue, a standard occurrence for PURExpress translated polypeptides. All Trp residues are bolded, with presumed sites of chlorination additionally colored red.

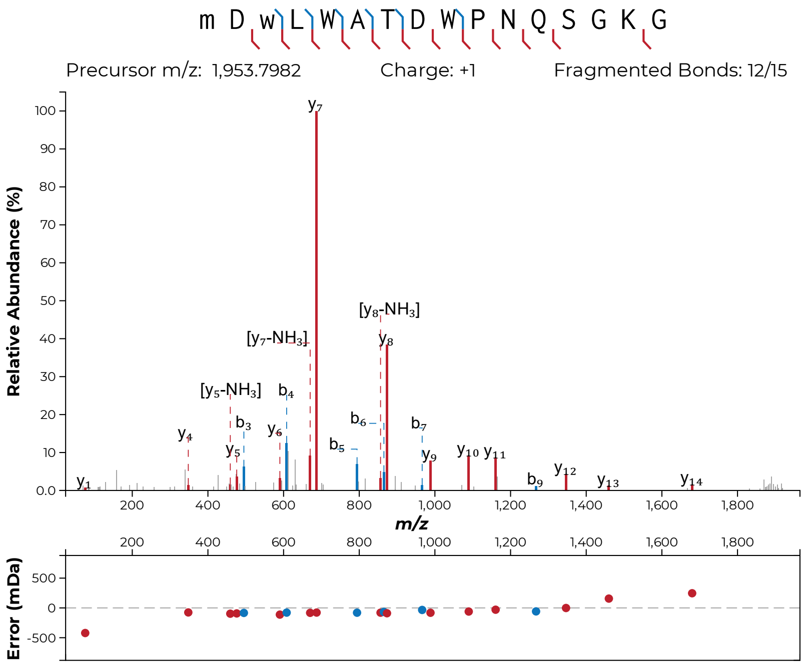


**Figure S38: MALDI-LIFT-MS for chlorinated ChlA$_{PE}$ inverted substrate.** Following post-translational halogenation of the ChlA$_{PE}$ reversed sequence, MS/MS indicated that the predominant monochlorinated species originated from Trp2 chlorination. Lowercase letters indicate modified residues. For example, the N-terminal formylated Met is indicated as “m”. Chlorinated Trp is indicated as “w”.

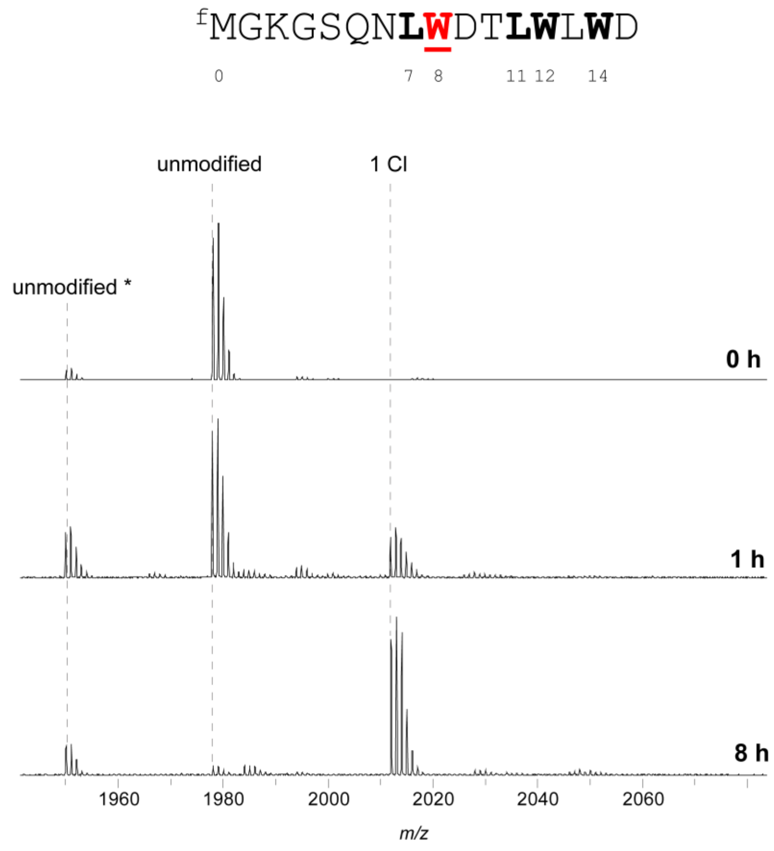


**Figure S39: MALDI-TOF-MS for ChlA$_{PE}$ P7L/A11L chlorination.** After an extended reaction time, ChlA$_{PE}$ P7L/A11L primarily yields a monochlorinated product (in comparison, ChlA$_{PE}$ was fully dichlorinated at just 1 h). Unmodified* refers to Met at the N-terminal position while unmodified refers to formylMet (from PURExpress in vitro transcription/translation). Expected *m/z* values for unmodified*, unmodified, and 1 Cl are 1949.9, 1977.9, 2011.9, respectively.

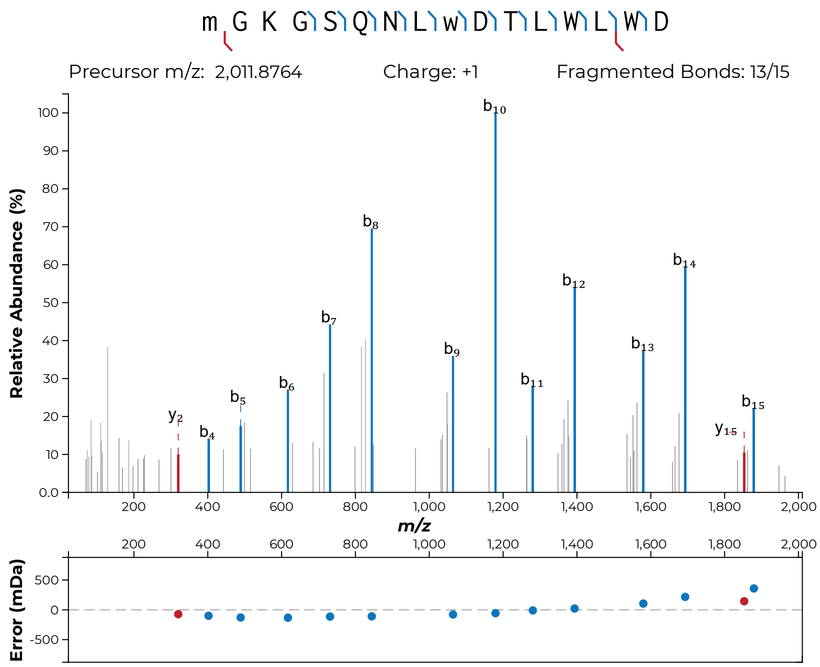


**Figure S40: MALDI-LIFT-MS for chlorinated ChlA$_{PE}$ P7L/A11L.** Following post-translational halogenation of the ChlA P7L/A11L peptide, MS/MS analysis indicated that the predominant monochlorinated species originated from Trp8 chlorination. Lowercase letters indicate modified residues. For example, the N-terminal formylated Met is indicated as “m”. Chlorinated Trp is indicated as “w”.

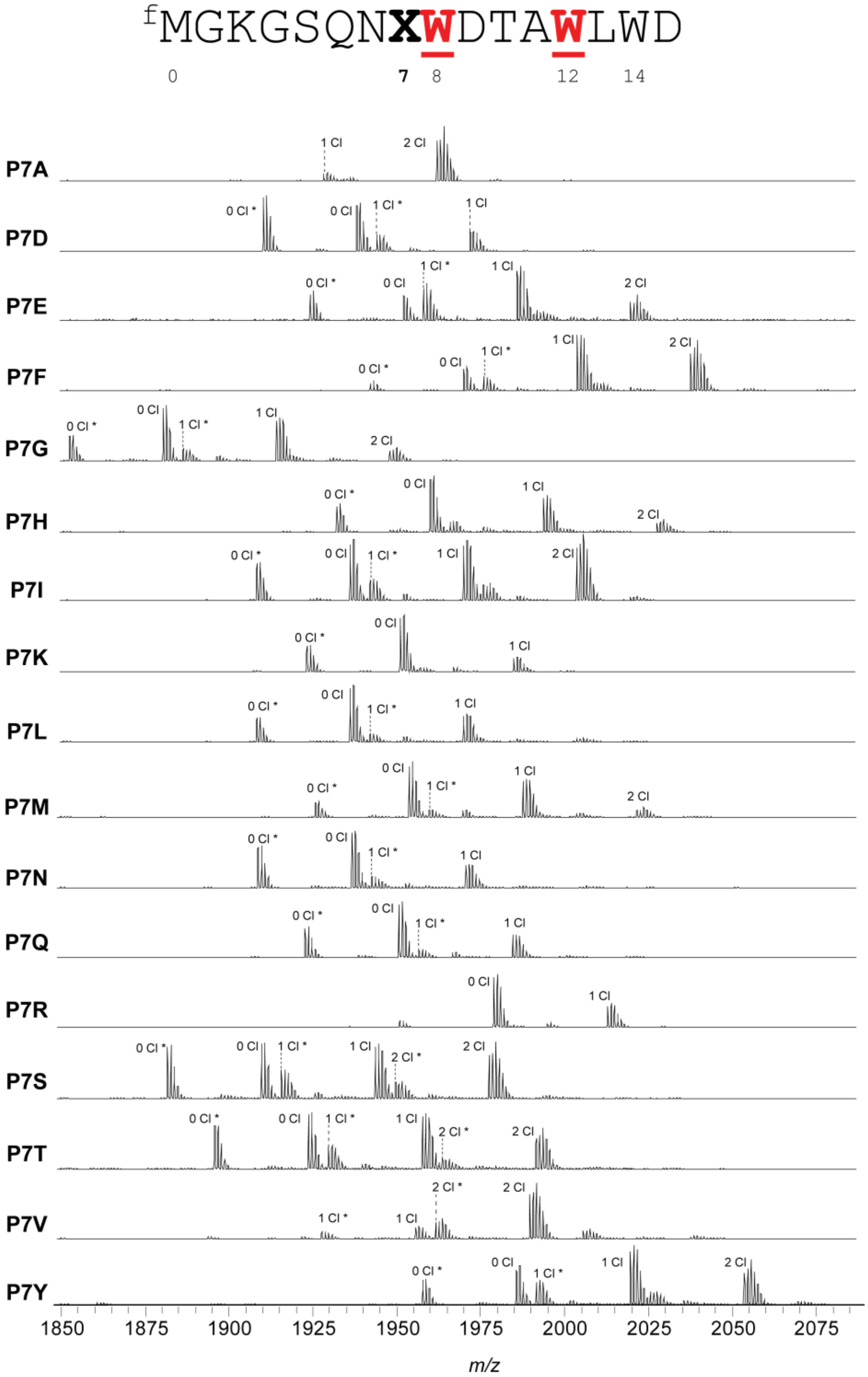


**Figure S41: MALDI-TOF-MS and -LIFT-MS for ChlA$_{PE}$ Pro7 variants.** (**a**) MALDI-TOF-MS spectra for all variants of position 7 of the ChlA$_{PE}$ substrate. Asterisks (*) indicate peptides lacking N-terminal formylation.

(**b-w, following pages**): MS/MS spectra acquired for this set of substrates. In all MS/MS spectra, lowercase letters indicate modified residues. For example, the N-terminal formylated Met is indicated as “m”. Chlorinated Trp is indicated as “w”.

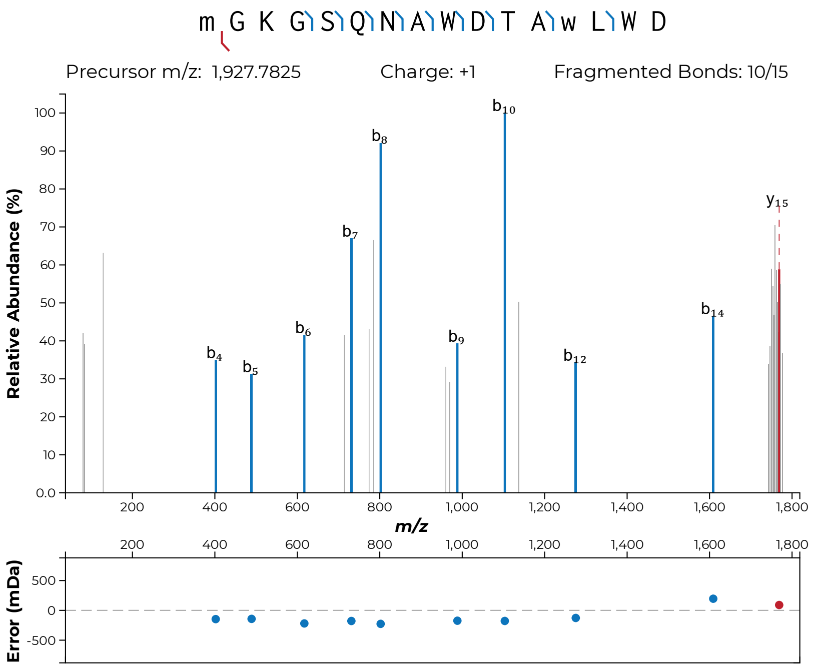

m G K G S Q N A W D T A w L W D
Precursor m/z: 1,927.7825
Charge: +1
Fragmented Bonds: 10/15
Relative Abundance (%)
b4
b5
b6
b7
b8
b9
b10
b12
b14
y15
m/z
Error (mDa)


**S41b**

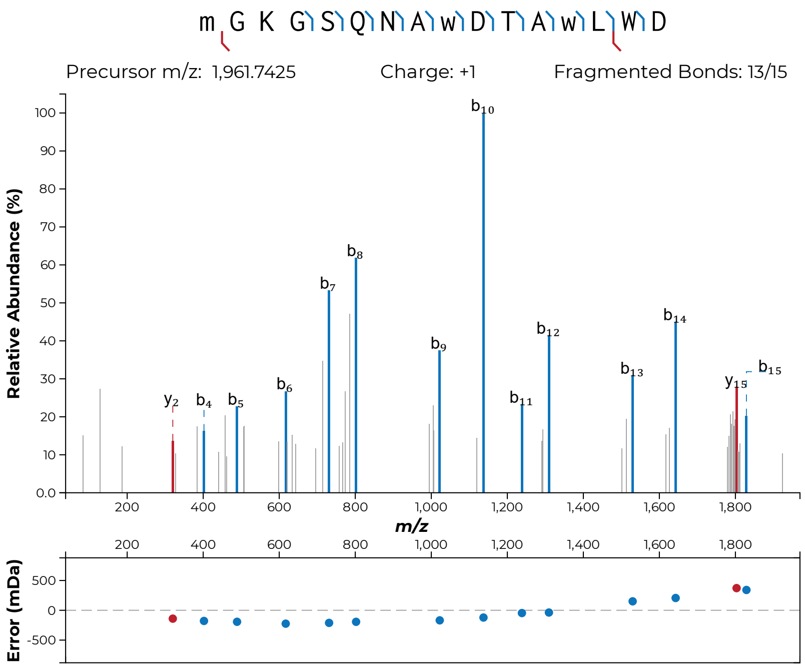
m G K G S Q N A w D T A w L W D
Precursor m/z: 1,961.7425
Charge: +1
Fragmented Bonds: 13/15
Relative Abundance (%)
m/z
Error (mDa)
y2
b4
b5
b6
b7
b8
b9
b10
b11
b12
b13
b14
y15
b15

**S41c**

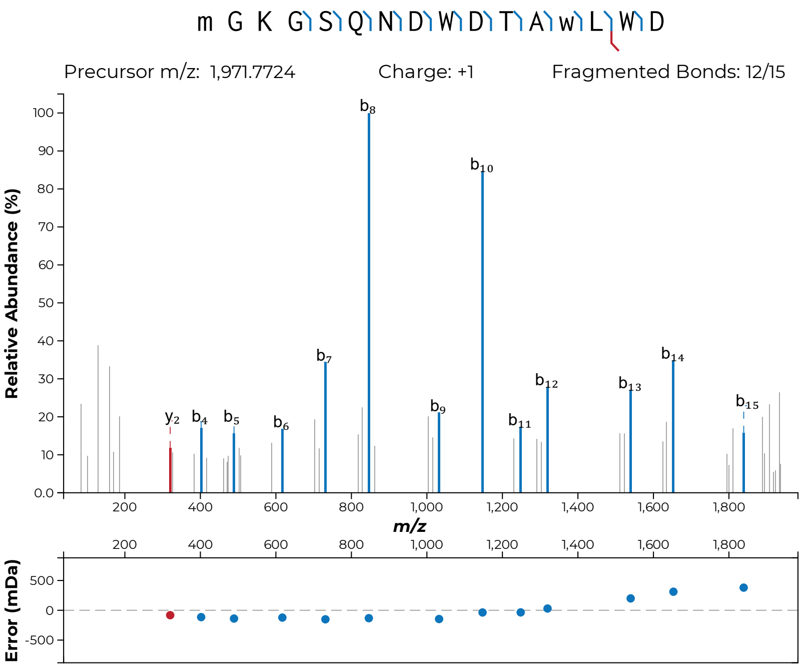
m G K G S Q N D W D T A w L W D
Precursor m/z: 1,971.7724
Charge: +1
Fragmented Bonds: 12/15
Relative Abundance (%)
m/z
Error (mDa)
y2
b4
b5
b6
b7
b8
b9
b10
b11
b12
b13
b14
b15

**S41d**

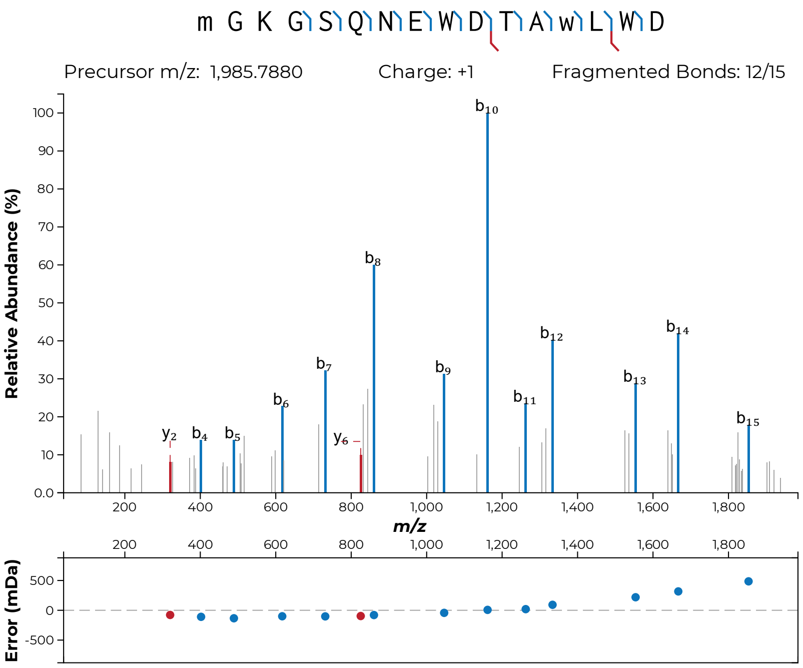

m G K G S Q N E W D T A w L W D
Precursor m/z: 1,985.7880
Charge: +1
Fragmented Bonds: 12/15
Relative Abundance (%)
m/z
Error (mDa)


**S41e**

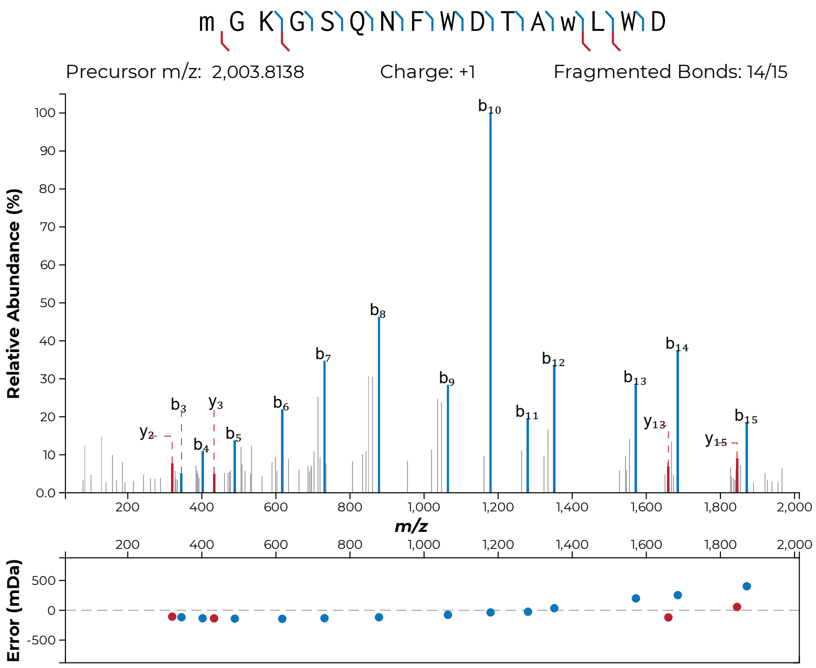
m G K G S Q N F W D T A w L W D
Precursor m/z: 2,003.8138
Charge: +1
Fragmented Bonds: 14/15
Relative Abundance (%)
m/z
Error (mDa)

**S41f**

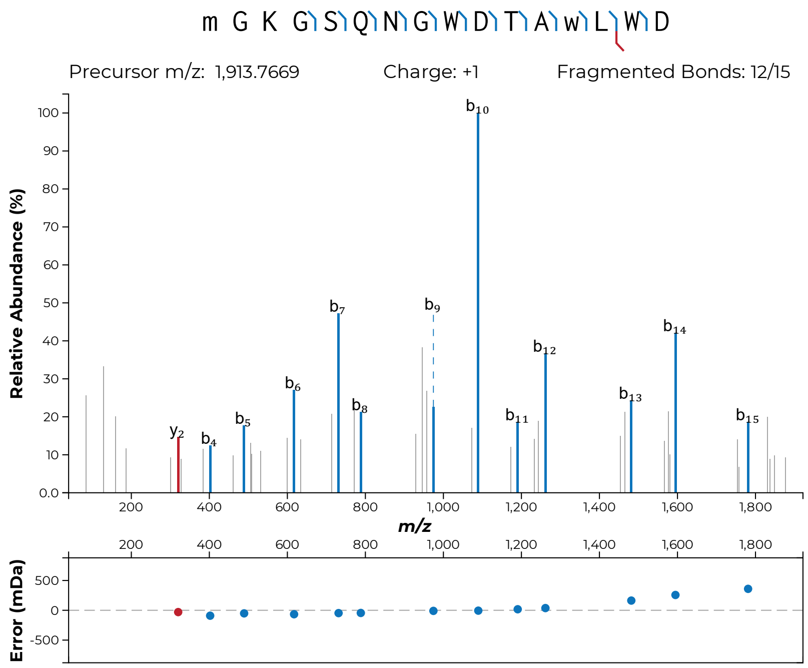

m G K G S Q N G W D T A w L W D
Precursor m/z: 1,913.7669
Charge: +1
Fragmented Bonds: 12/15
Relative Abundance (%)
m/z
Error (mDa)
y2
b4
b5
b6
b7
b8
b9
b10
b11
b12
b13
b14
b15


**S41g**

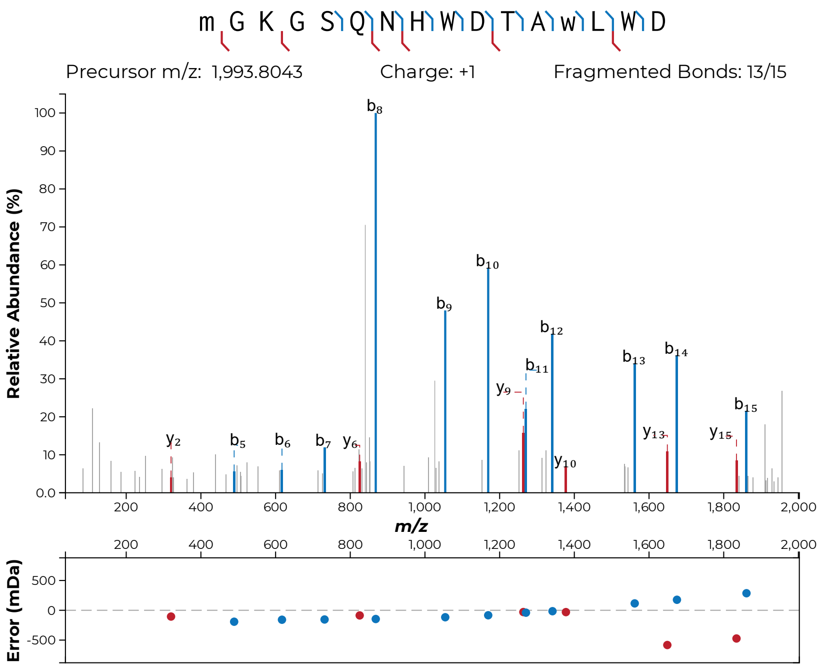
m G K G S Q N H W D T A w L W D
Precursor m/z: 1,993.8043
Charge: +1
Fragmented Bonds: 13/15
Relative Abundance (%)
m/z
Error (mDa)

**S41h**

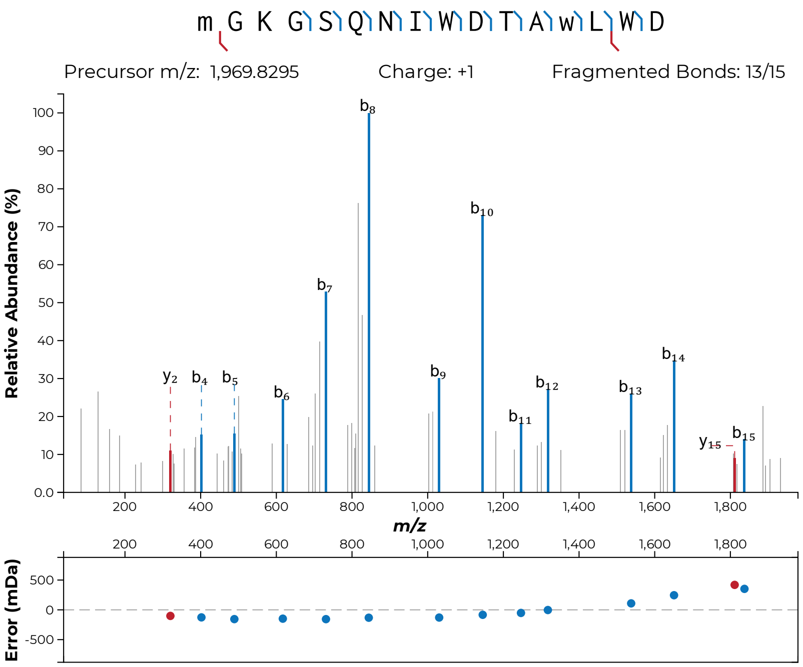
m G K G S Q N I W D T A w L W D
Precursor m/z: 1,969.8295
Charge: +1
Fragmented Bonds: 13/15
Relative Abundance (%)
m/z
Error (mDa)
y2
b4
b5
b6
b7
b8
b9
b10
b11
b12
b13
b14
y15
b15

**S41i**

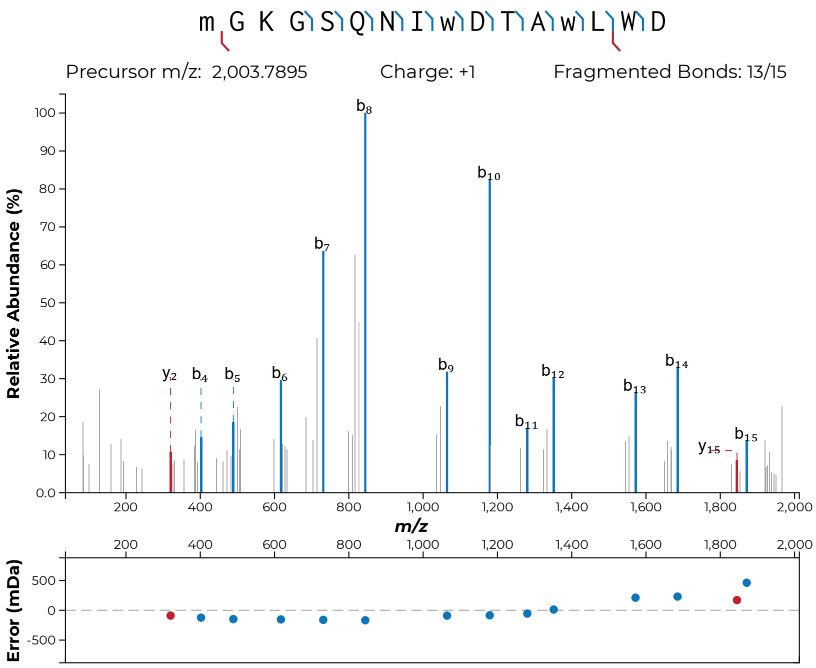
m G K G S Q N I w D T A w L W D
Precursor m/z: 2,003.7895
Charge: +1
Fragmented Bonds: 13/15
Relative Abundance (%)
m/z
Error (mDa)

**S41j**

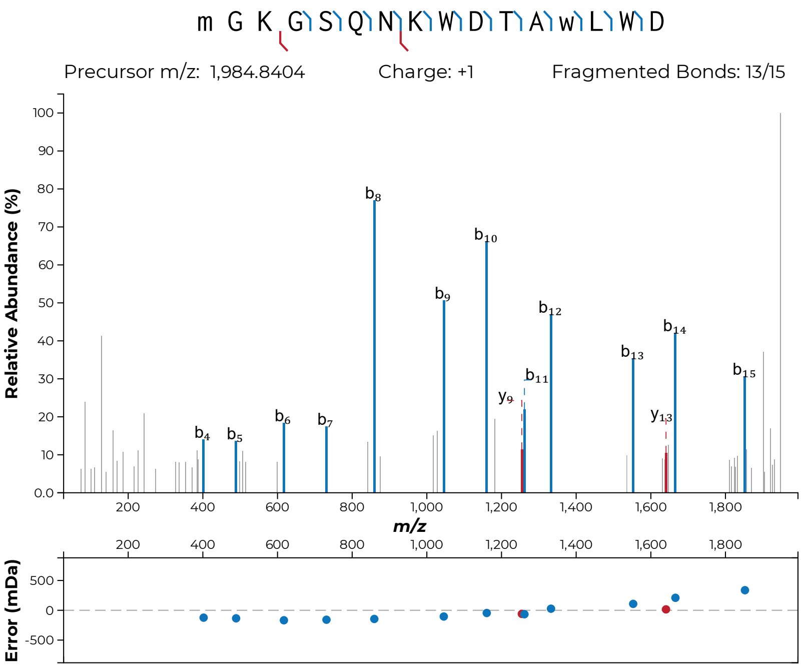
m G K G S Q N K W D T A w L W D
Precursor m/z: 1,984.8404
Charge: +1
Fragmented Bonds: 13/15
Relative Abundance (%)
m/z
Error (mDa)
b4
b5
b6
b7
b8
b9
b10
b11
b12
b13
b14
b15
y9
y13

**S41k**

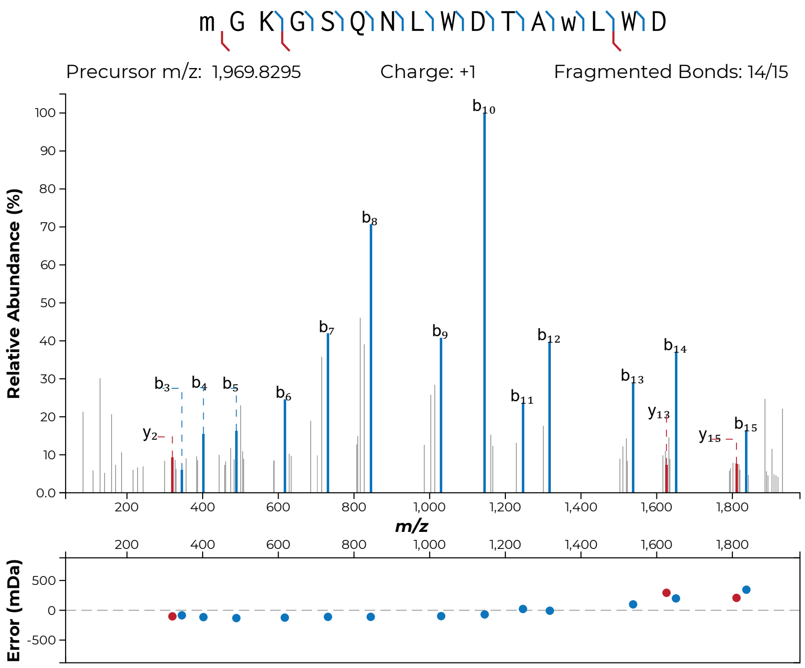

m G K G S Q N L W D T A w L W D
Precursor m/z: 1,969.8295
Charge: +1
Fragmented Bonds: 14/15
Relative Abundance (%)
m/z
Error (mDa)


**S41l**

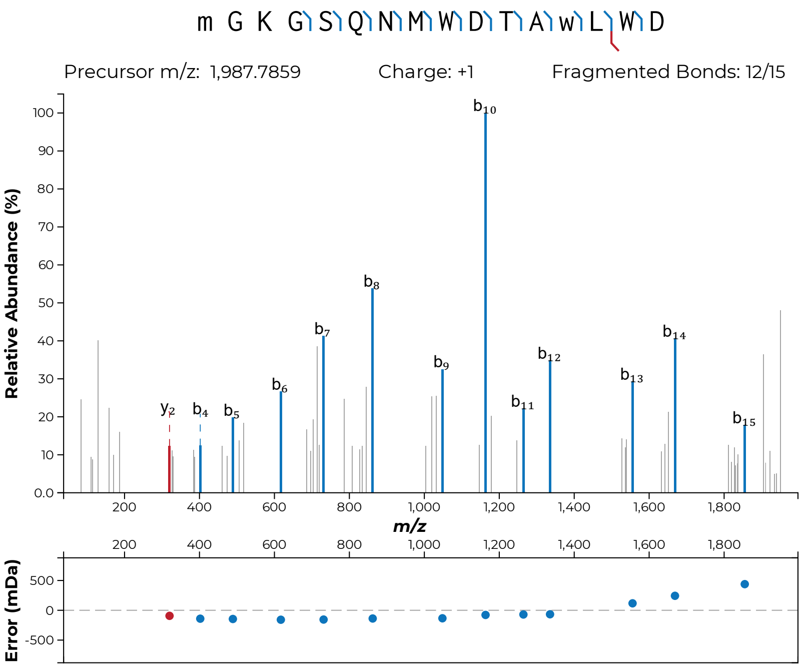
m G K G S Q N M W D T A w L W D
Precursor m/z: 1,987.7859
Charge: +1
Fragmented Bonds: 12/15
Relative Abundance (%)
m/z
Error (mDa)

**S41m**

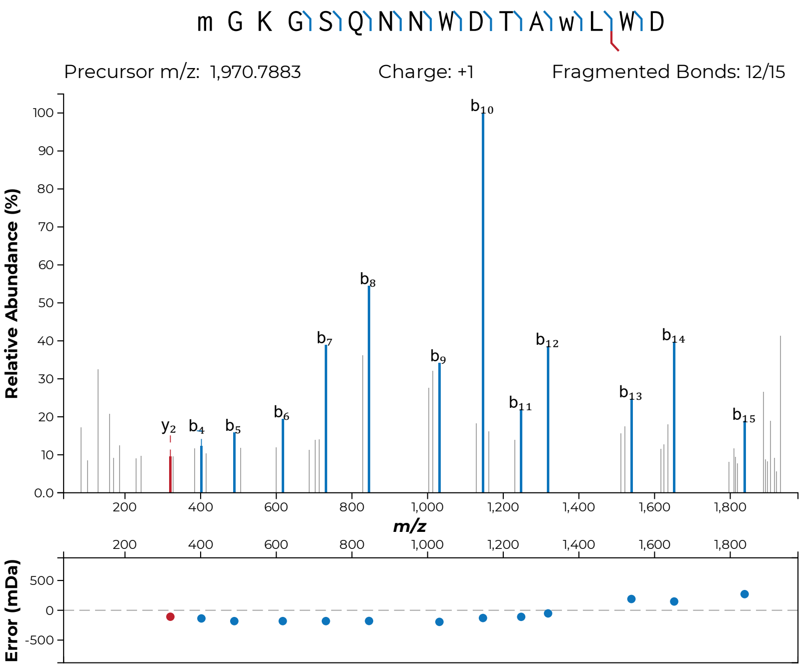
m G K G S Q N N W D T A w L W D
Precursor m/z: 1,970.7883
Charge: +1
Fragmented Bonds: 12/15
Relative Abundance (%)
m/z
Error (mDa)

**S41n**

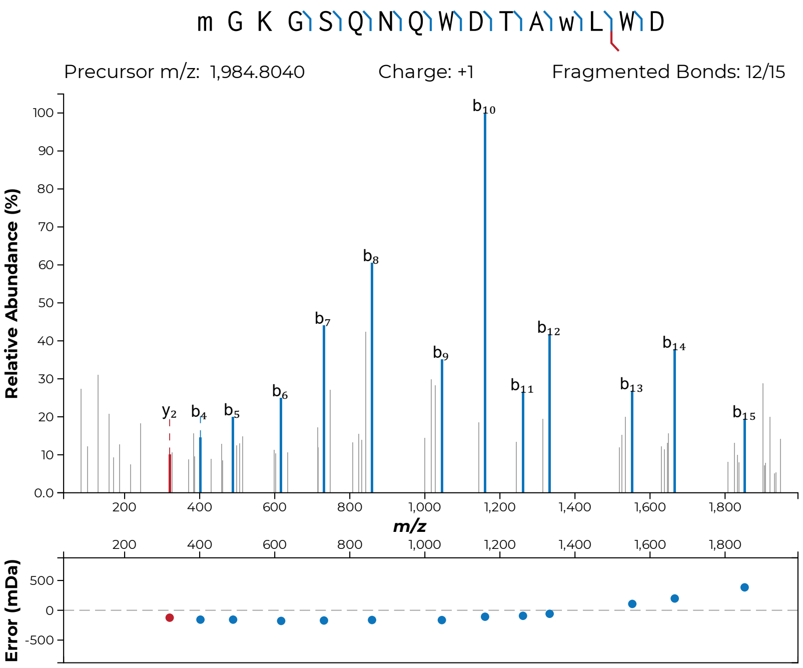

m G K G S Q N Q W D T A w L W D
Precursor m/z: 1,984.8040
Charge: +1
Fragmented Bonds: 12/15
Relative Abundance (%)
m/z
Error (mDa)
y2 b4 b5 b6 b7 b8 b9 b10 b11 b12 b13 b14 b15


**S41o**

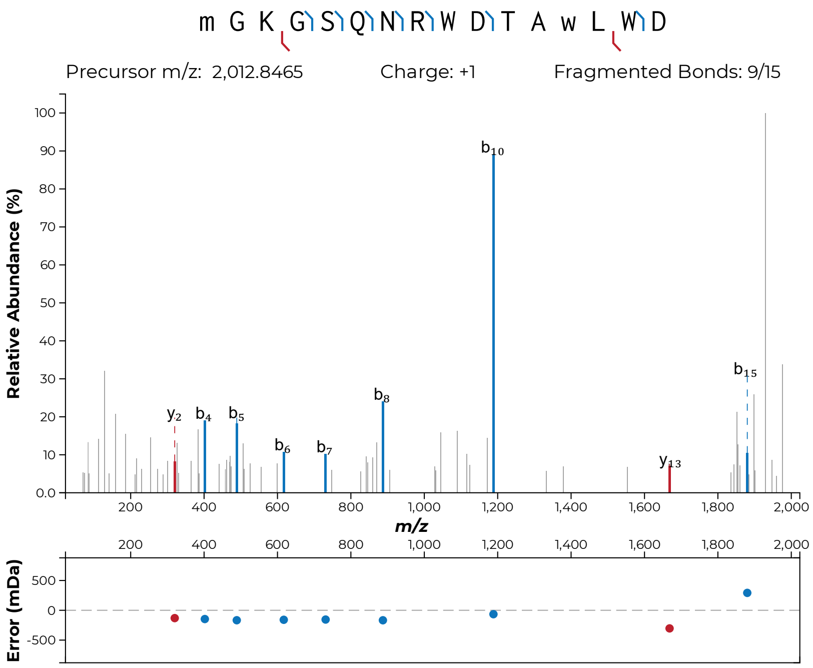

m G K G S Q N R W D T A w L W D
Precursor m/z: 2,012.8465
Charge: +1
Fragmented Bonds: 9/15
Relative Abundance (%)
m/z
Error (mDa)


**S41p**

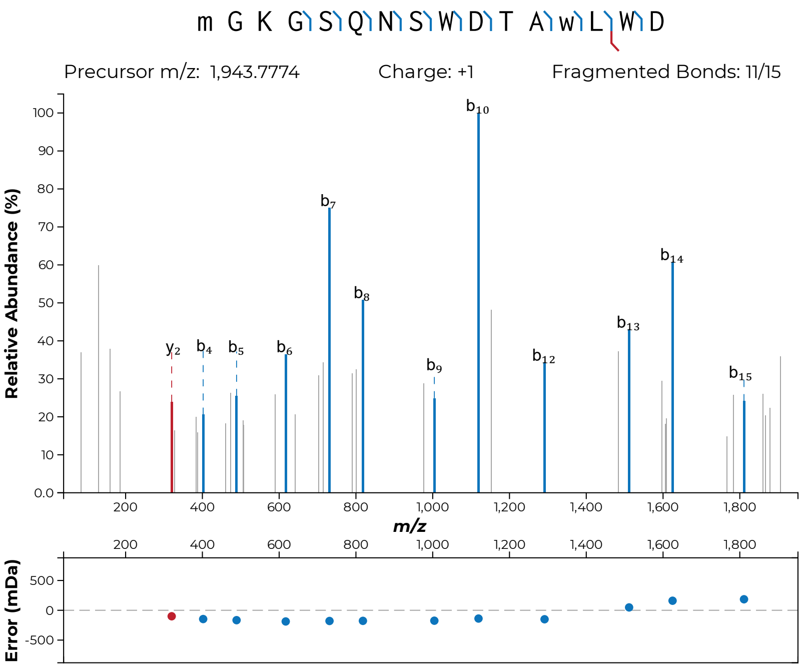
m G K G S Q N S W D T A w L W D
Precursor m/z: 1,943.7774
Charge: +1
Fragmented Bonds: 11/15
Relative Abundance (%)
m/z
Error (mDa)
y2
b4
b5
b6
b7
b8
b9
b10
b12
b13
b14
b15

**S41q**

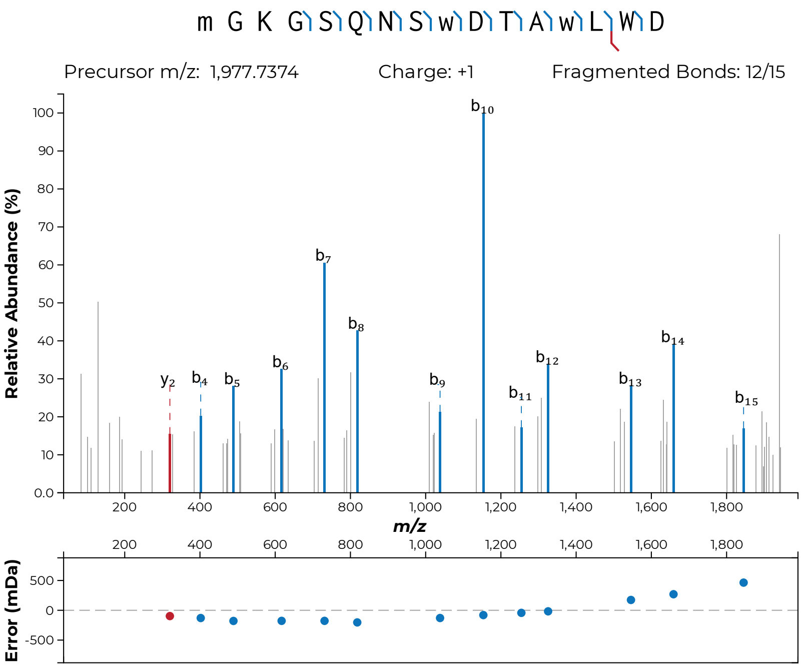
m G K G S Q N S w D T A w L W D
Precursor m/z: 1,977.7374
Charge: +1
Fragmented Bonds: 12/15
Relative Abundance (%)
m/z
Error (mDa)
y2
b4
b5
b6
b7
b8
b9
b10
b11
b12
b13
b14
b15

**S41r**

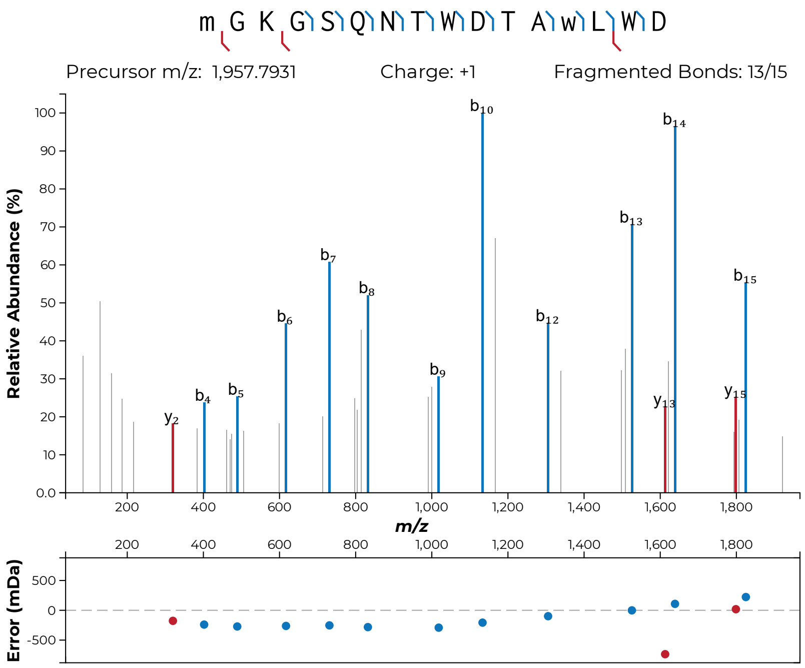
m G K G S Q N T W D T A w L W D
Precursor m/z: 1,957.7931
Charge: +1
Fragmented Bonds: 13/15
Relative Abundance (%)
m/z
Error (mDa)
b10
b14
b13
b7
b8
b6
b12
b15
b9
b4
b5
y2
y13
y15

**S41s**

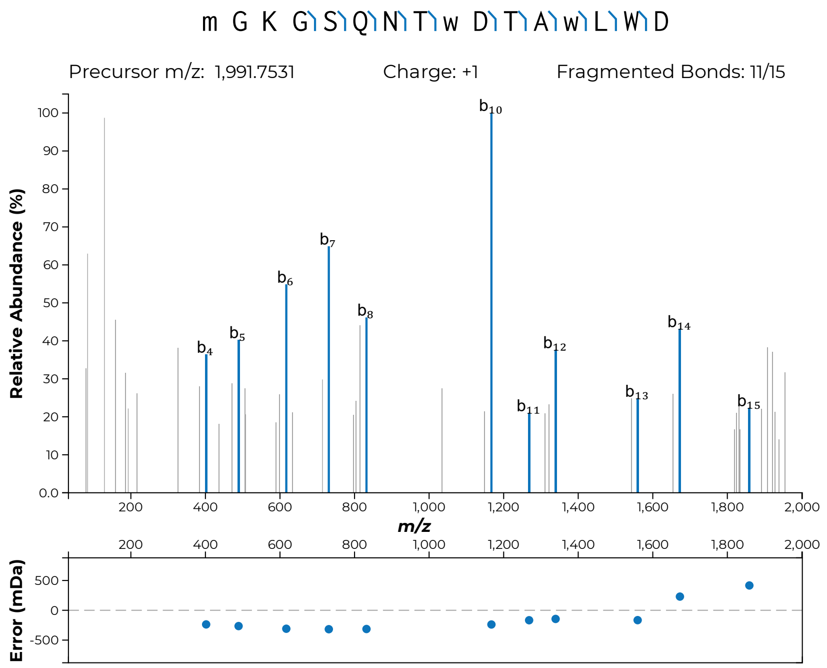
m G K G S Q N T w D T A w L W D
Precursor m/z: 1,991.7531
Charge: +1
Fragmented Bonds: 11/15
Relative Abundance (%)
b4
b5
b6
b7
b8
b10
b11
b12
b13
b14
b15
m/z
Error (mDa)

**S41t**

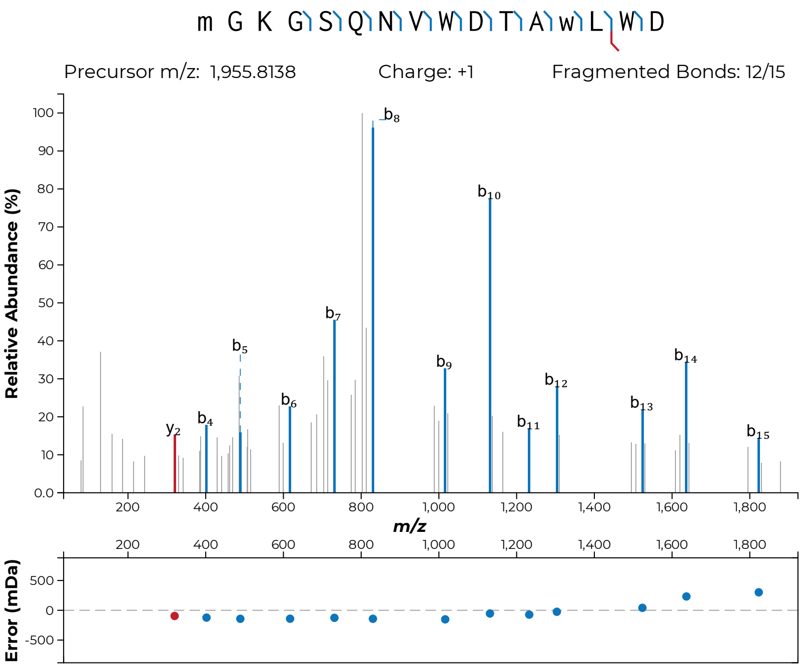
m G K G S Q N V W D T A w L W D
Precursor m/z: 1,955.8138
Charge: +1
Fragmented Bonds: 12/15
Relative Abundance (%)
m/z
Error (mDa)

**S41u**

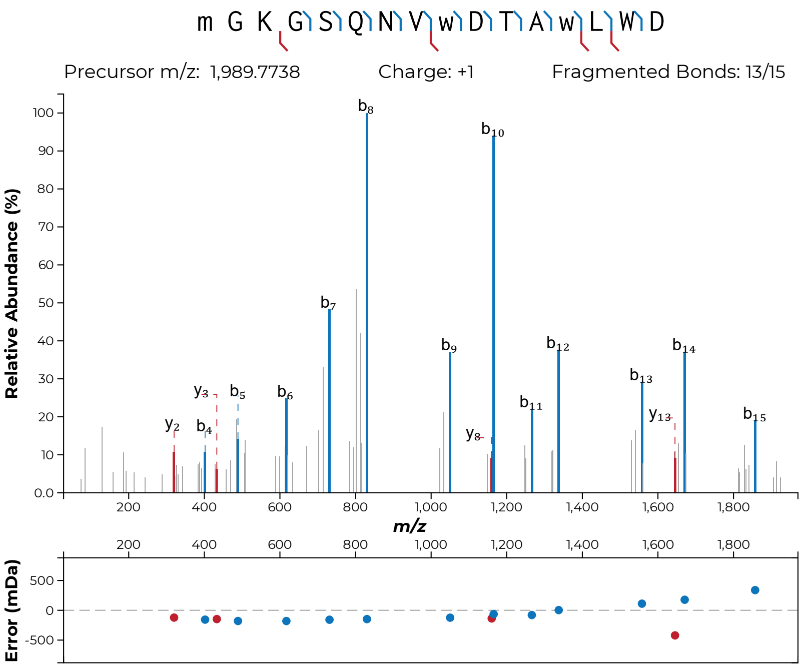

m G K G S Q N V w D T A w L W D
Precursor m/z: 1,989.7738
Charge: +1
Fragmented Bonds: 13/15
Relative Abundance (%)
m/z
Error (mDa)


S41v

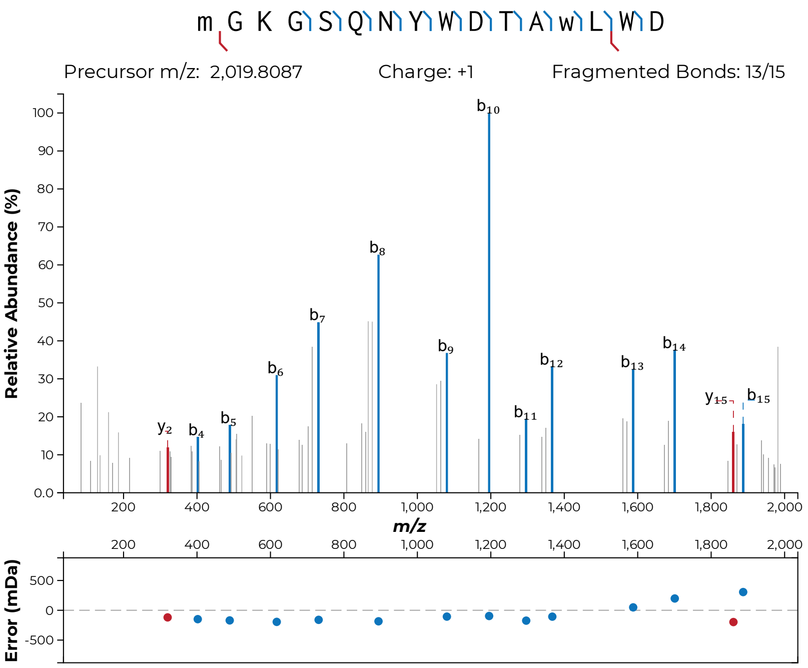

m G K G S Q N Y W D T A w L W D
Precursor m/z: 2,019.8087
Charge: +1
Fragmented Bonds: 13/15
Relative Abundance (%)
m/z
Error (mDa)
y2 b4 b5 b6 b7 b8 b9 b10 b11 b12 b13 b14 y15 b15


**S41w**

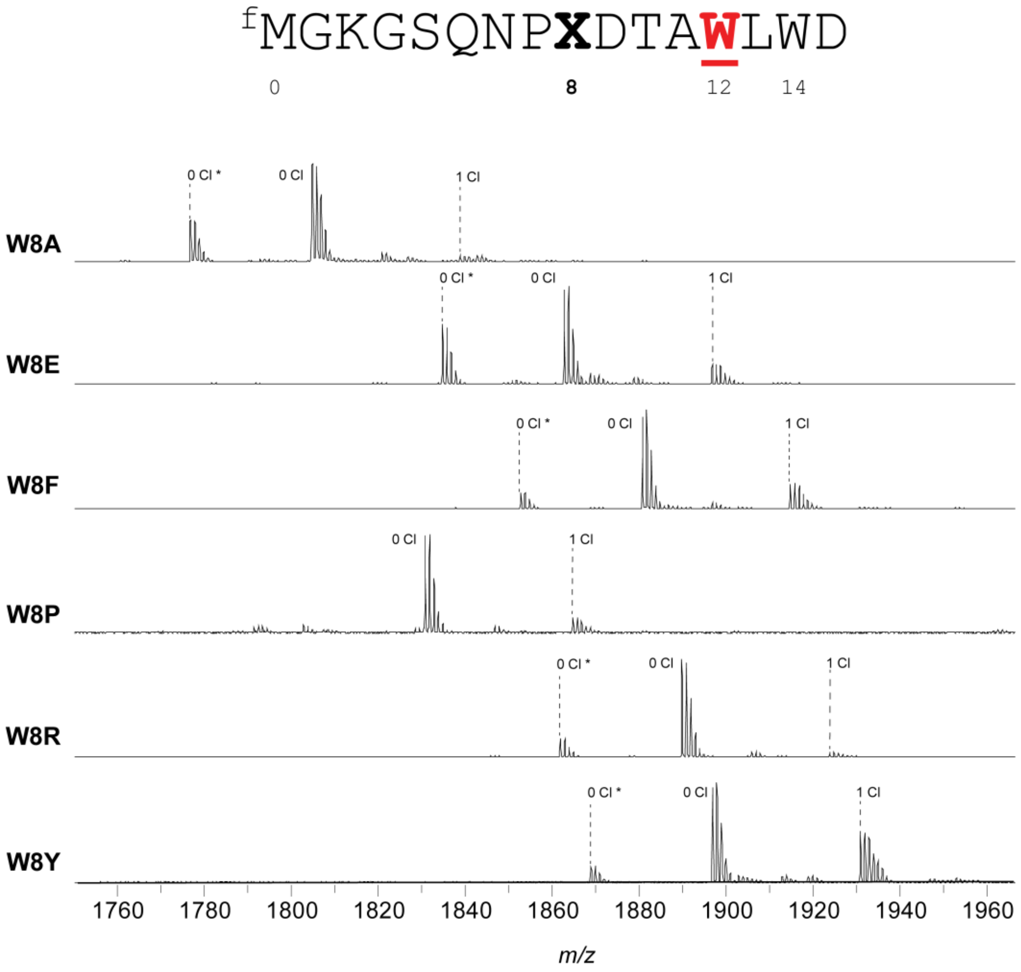


**Figure S42: MALDI-TOF-MS and -LIFT-MS for ChlA$_{PE}$ Trp8 variants.** (**a**) MALDI-TOF-MS spectra for all variants of position 8 of the ChlA$_{PE}$ substrate. Asterisks (*) indicate peptides lacking N-terminal formylation.

(**b-c, following pages**): MS/MS spectra acquired for this set of substrates. In all MS/MS spectra, lowercase letters indicate modified residues. For example, the N-terminal formylated Met is indicated as “m”. Chlorinated Trp is indicated as “w”.

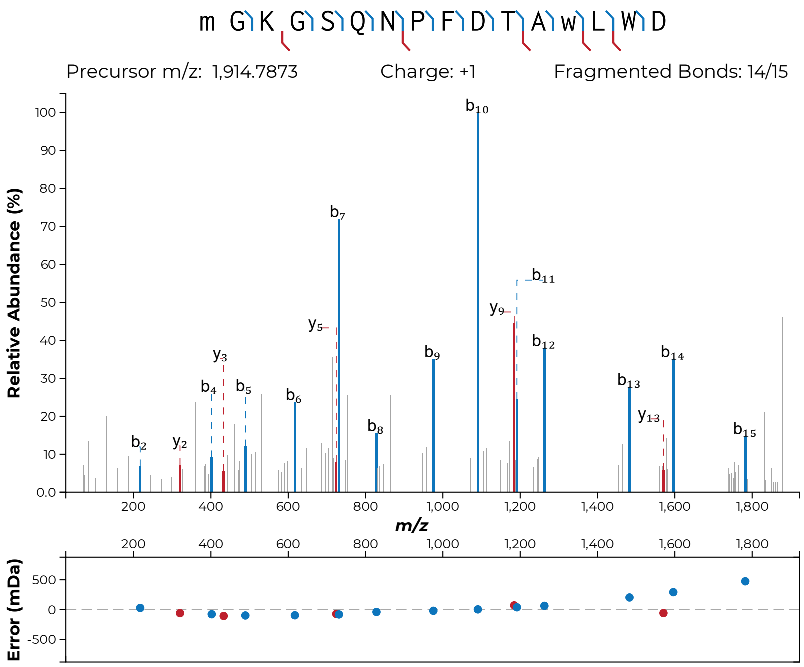
m G K G S Q N P F D T A w L W D
Precursor m/z: 1,914.7873
Charge: +1
Fragmented Bonds: 14/15
Relative Abundance (%)
m/z
Error (mDa)

**S42b**

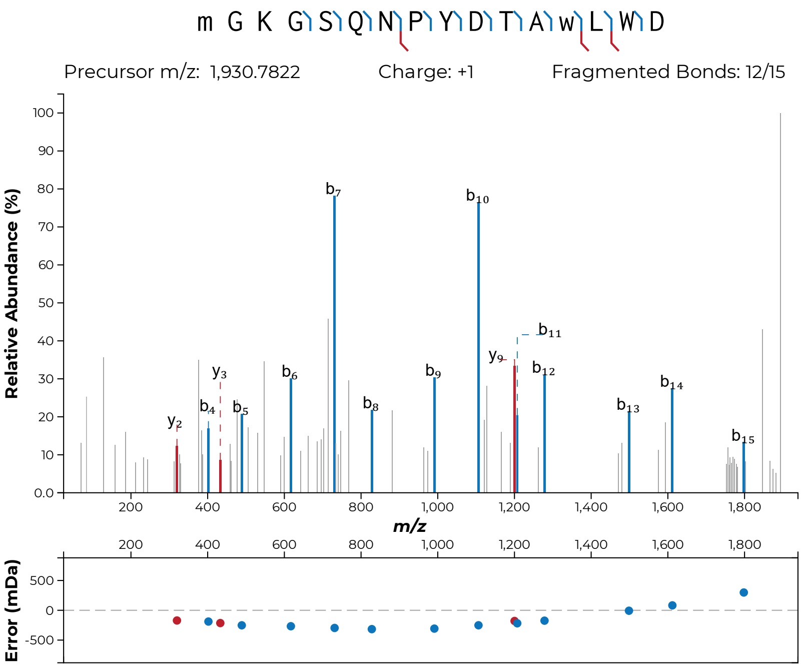

m G K G S Q N P Y D T A w L W D
Precursor m/z: 1,930.7822
Charge: +1
Fragmented Bonds: 12/15
Relative Abundance (%)
m/z
Error (mDa)


**S42c**

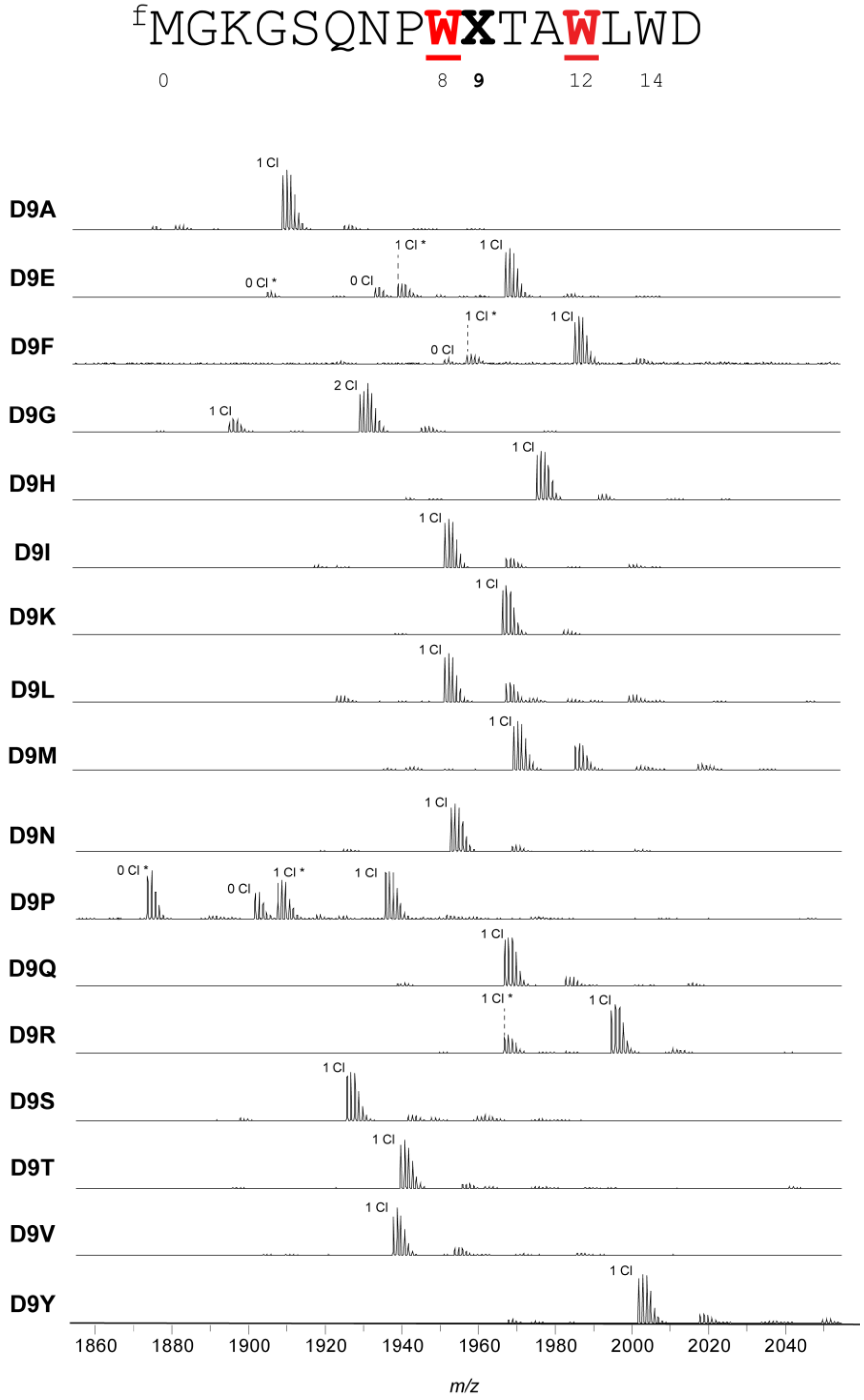


**Figure S43: MALDI-TOF-MS and -LIFT-MS for ChlA$_{PE}$ Asp9 variants.** (**a**) MALDI-TOF-MS spectra for all variants of position 9 of the ChlA$_{PE}$ substrate. Asterisks (*) indicate peptides lacking N-terminal formylation.

(**b-s, following pages**): MS/MS spectra acquired for this set of substrates. In all MS/MS spectra, lowercase letters indicate modified residues. For example, the N-terminal formylated Met is indicated as "m". Chlorinated Trp is indicated as "w".

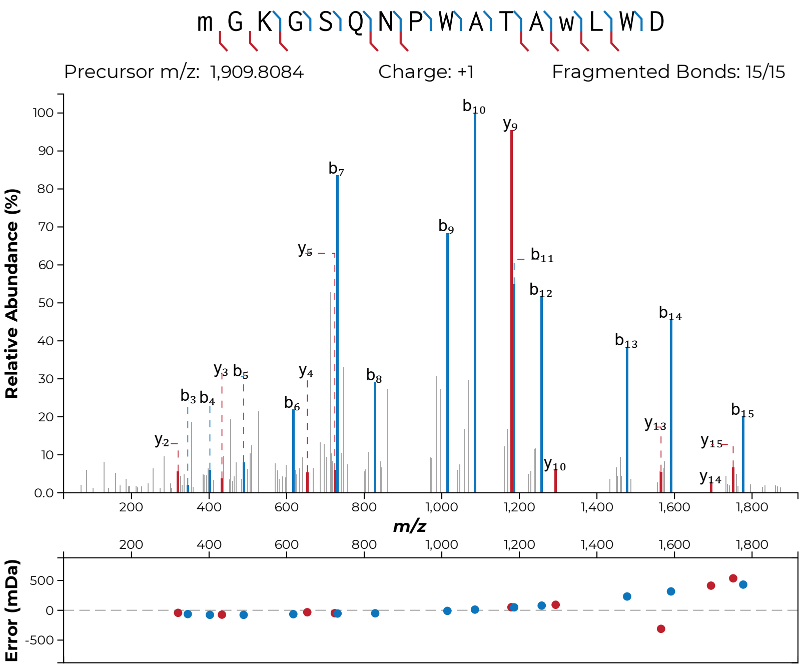
m G K G S Q N P W A T A w L W D
Precursor m/z: 1,909.8084
Charge: +1
Fragmented Bonds: 15/15
Relative Abundance (%)
m/z
Error (mDa)

**S43b**

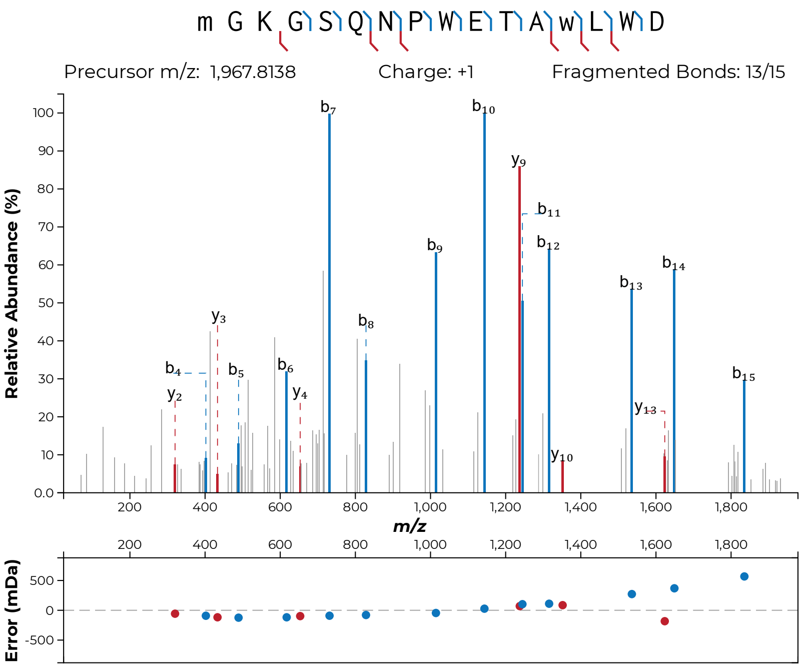

m G K G S Q N P W E T A w L W D
Precursor m/z: 1,967.8138
Charge: +1
Fragmented Bonds: 13/15
Relative Abundance (%)
m/z
Error (mDa)


**S43c**

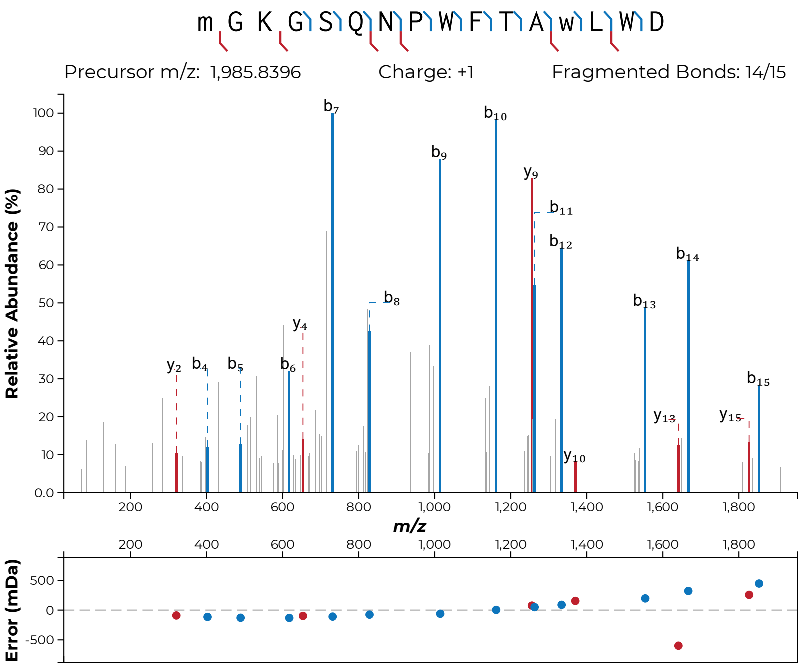
m G K G S Q N P W F T A w L W D
Precursor m/z: 1,985.8396
Charge: +1
Fragmented Bonds: 14/15
Relative Abundance (%)
m/z
Error (mDa)
b7
b10
b9
y9
b11
b12
b14
b13
b8
b15
y2
b4
b5
b6
y4
y10
y13
y15

**S43d**

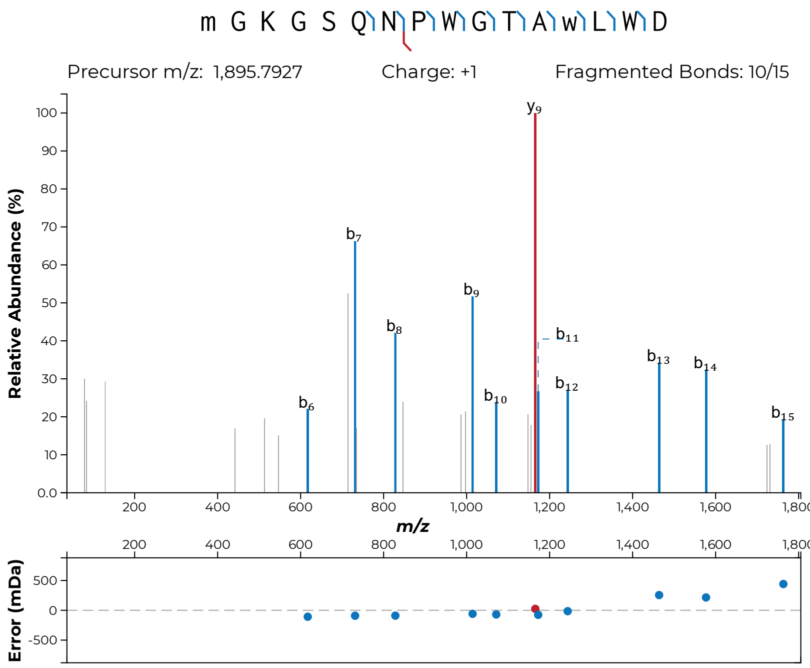

m G K G S Q N P W G T A w L W D
Precursor m/z: 1,895.7927
Charge: +1
Fragmented Bonds: 10/15
Relative Abundance (%)
y9
b7
b9
b8
b11
b13
b14
b12
b6
b10
b15
m/z
Error (mDa)


**S43e**

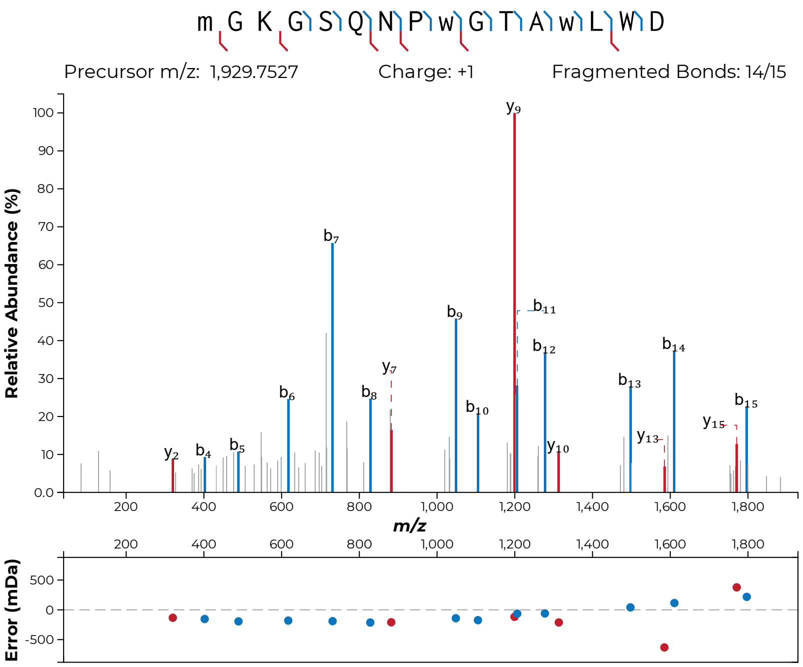
m G K G S Q N P w G T A w L W D
Precursor m/z: 1,929.7527
Charge: +1
Fragmented Bonds: 14/15
Relative Abundance (%)
m/z
Error (mDa)
y2
b4
b5
b6
b7
b8
y7
b9
b10
y9
b11
b12
y10
b13
y13
b14
y15
b15

**S43f**

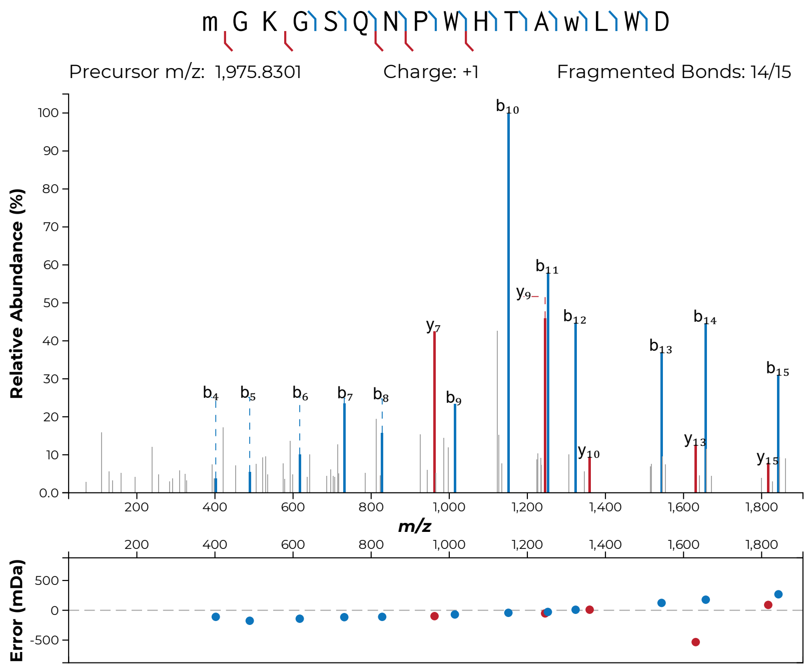
m G K G S Q N P W H T A w L W D
Precursor m/z: 1,975.8301
Charge: +1
Fragmented Bonds: 14/15
Relative Abundance (%)
m/z
Error (mDa)

**S43g**

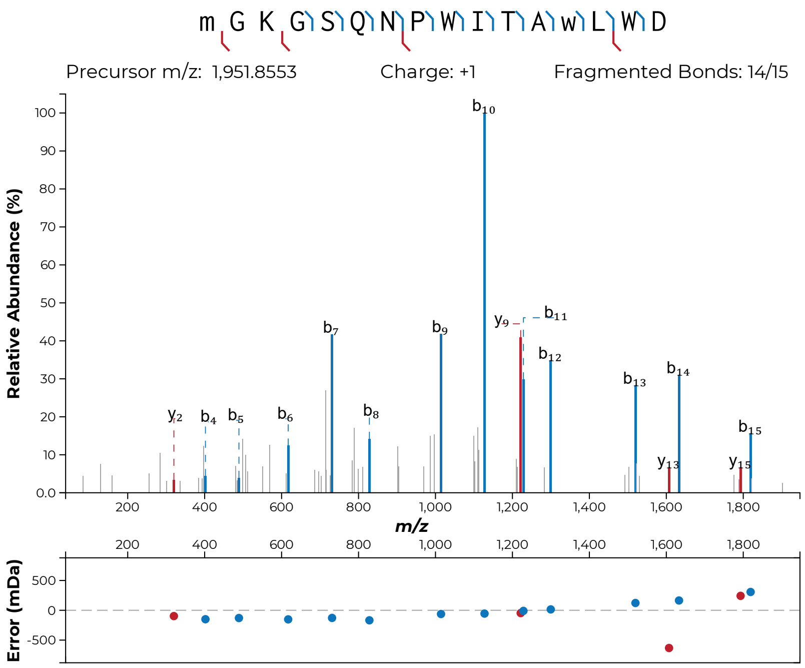
m G K G S Q N P W I T A w L W D
Precursor m/z: 1,951.8553
Charge: +1
Fragmented Bonds: 14/15
Relative Abundance (%)
m/z
Error (mDa)

**S43h**

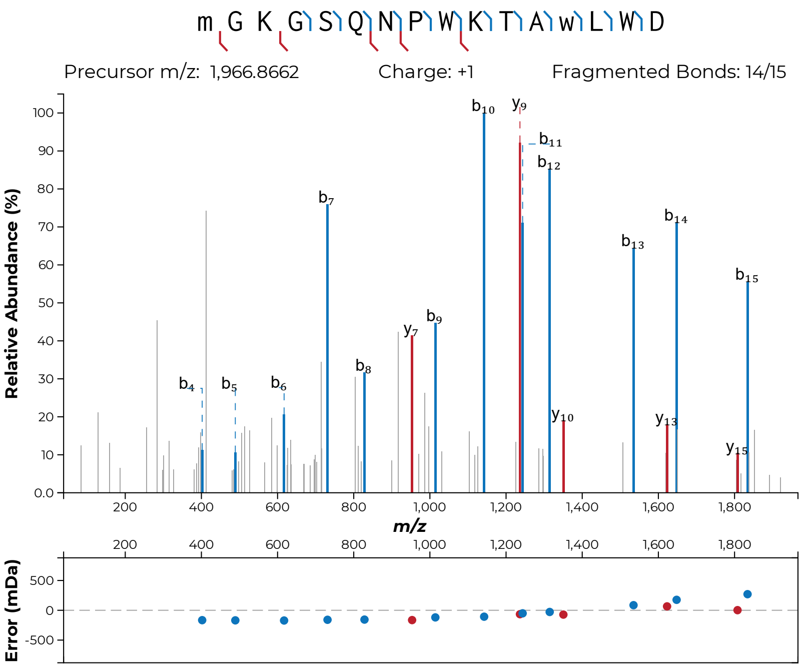

m G K G S Q N P W K T A w L W D
Precursor m/z: 1,966.8662
Charge: +1
Fragmented Bonds: 14/15
Relative Abundance (%)
m/z
Error (mDa)
b4
b5
b6
b7
b8
b9
b10
b11
b12
b13
b14
b15
y7
y9
y10
y13
y15


**S43i**

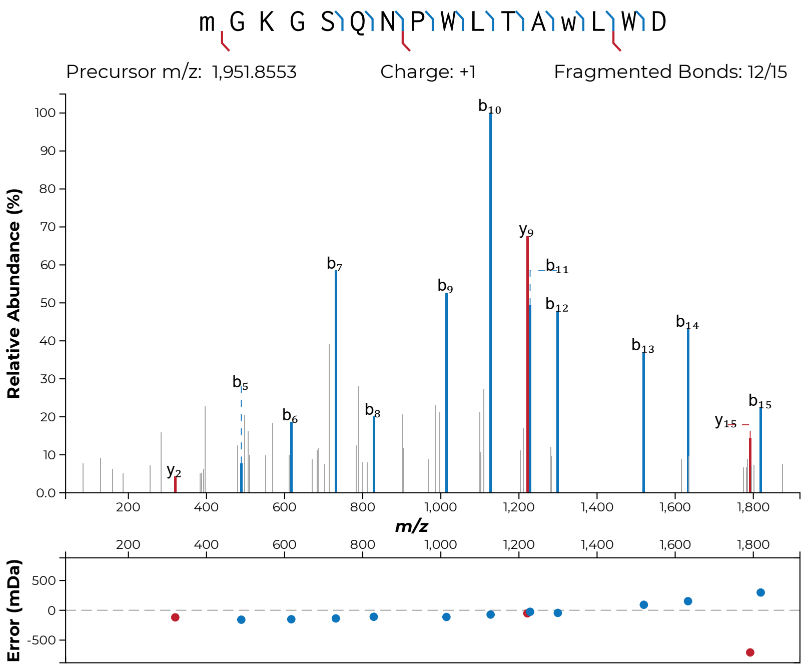
m G K G S Q N P W L T A w L W D
Precursor m/z: 1,951.8553
Charge: +1
Fragmented Bonds: 12/15
Relative Abundance (%)
m/z
Error (mDa)

**S43j**

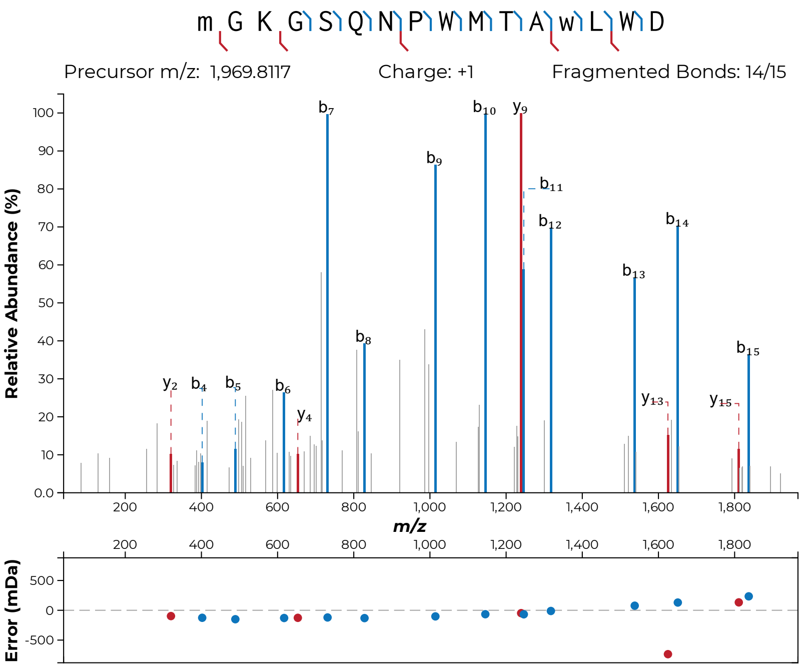

m G K G S Q N P W M T A w L W D
Precursor m/z: 1,969.8117
Charge: +1
Fragmented Bonds: 14/15
Relative Abundance (%)
m/z
Error (mDa)


**S43k**

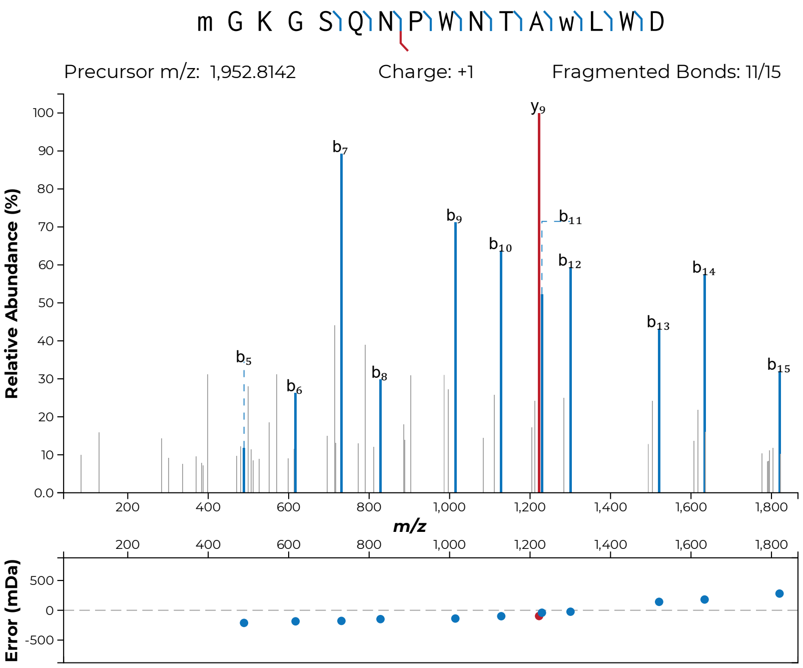

m G K G S Q N P W N T A w L W D
Precursor m/z: 1,952.8142
Charge: +1
Fragmented Bonds: 11/15
Relative Abundance (%)
m/z
Error (mDa)


**S43l**

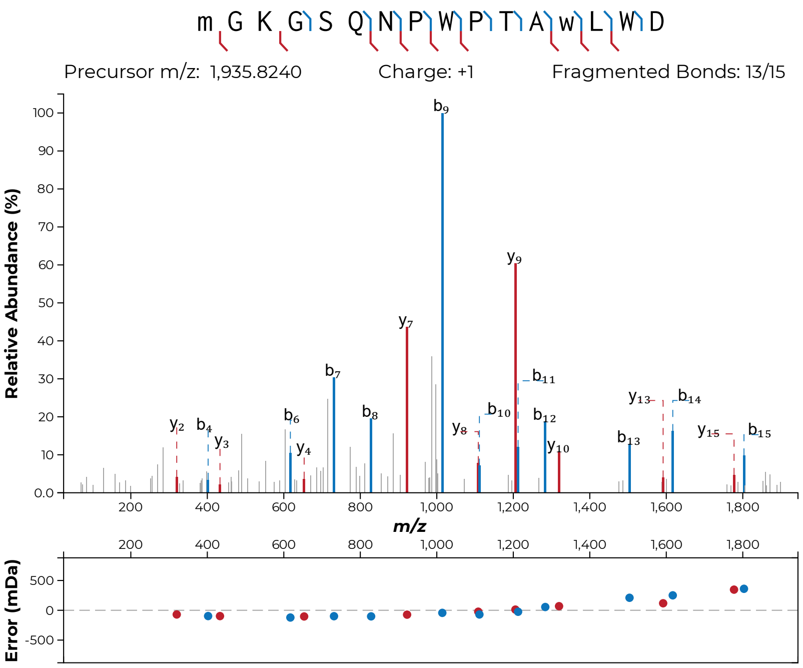
m G K G S Q N P W P T A w L W D
Precursor m/z: 1,935.8240
Charge: +1
Fragmented Bonds: 13/15
Relative Abundance (%)
m/z
Error (mDa)

**S43m**

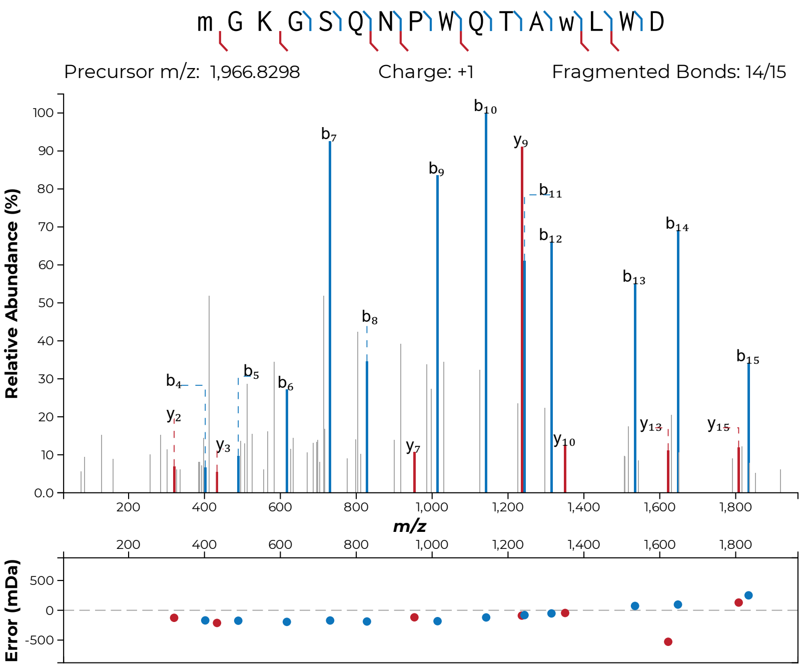
m G K G S Q N P W Q T A w L W D
Precursor m/z: 1,966.8298
Charge: +1
Fragmented Bonds: 14/15
Relative Abundance (%)
m/z
Error (mDa)

**S43n**

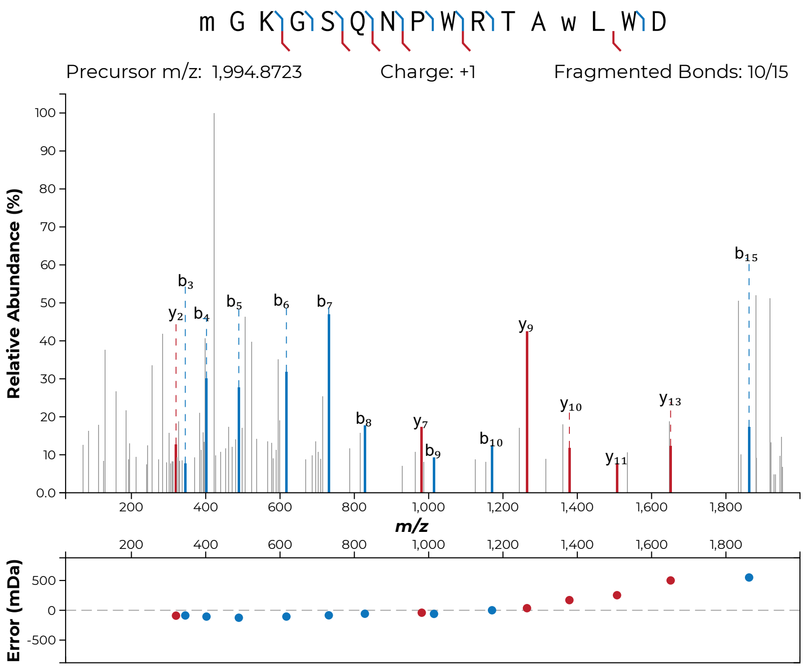

m G K G S Q N P W R T A w L W D
Precursor m/z: 1,994.8723
Charge: +1
Fragmented Bonds: 10/15
Relative Abundance (%)
m/z
Error (mDa)


**S43o**

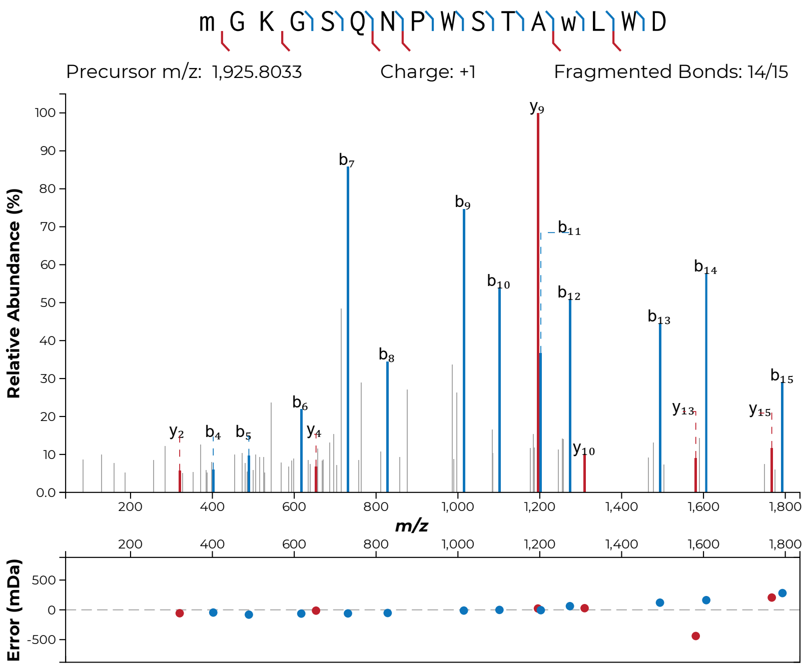

m G K G S Q N P W S T A w L W D
Precursor m/z: 1,925.8033
Charge: +1
Fragmented Bonds: 14/15
Relative Abundance (%)
m/z
Error (mDa)


**S43p**

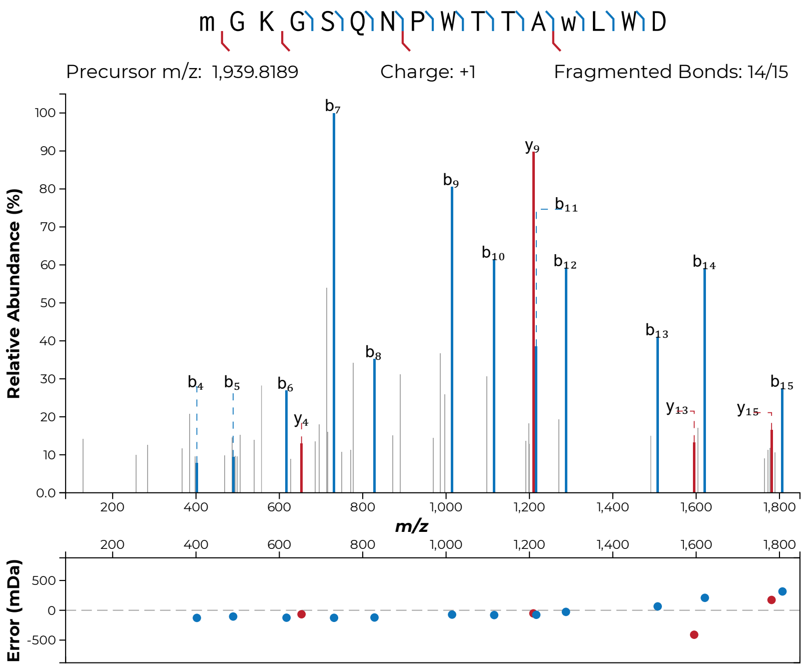
m G K G S Q N P W T T A w L W D
Precursor m/z: 1,939.8189
Charge: +1
Fragmented Bonds: 14/15
Relative Abundance (%)
m/z
Error (mDa)

**S43q**

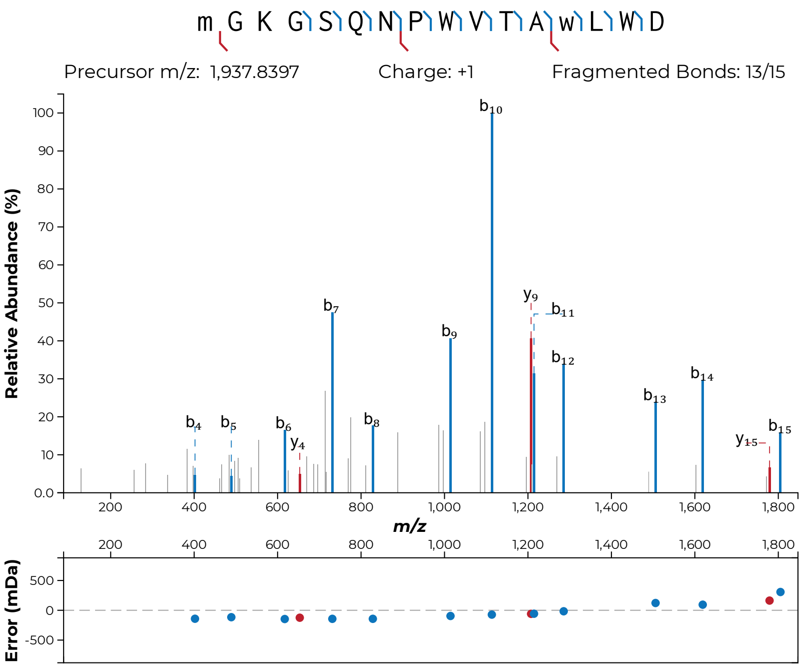
m G K G S Q N P W V T A w L W D
Precursor m/z: 1,937.8397
Charge: +1
Fragmented Bonds: 13/15
Relative Abundance (%)
m/z
Error (mDa)

**S43r**

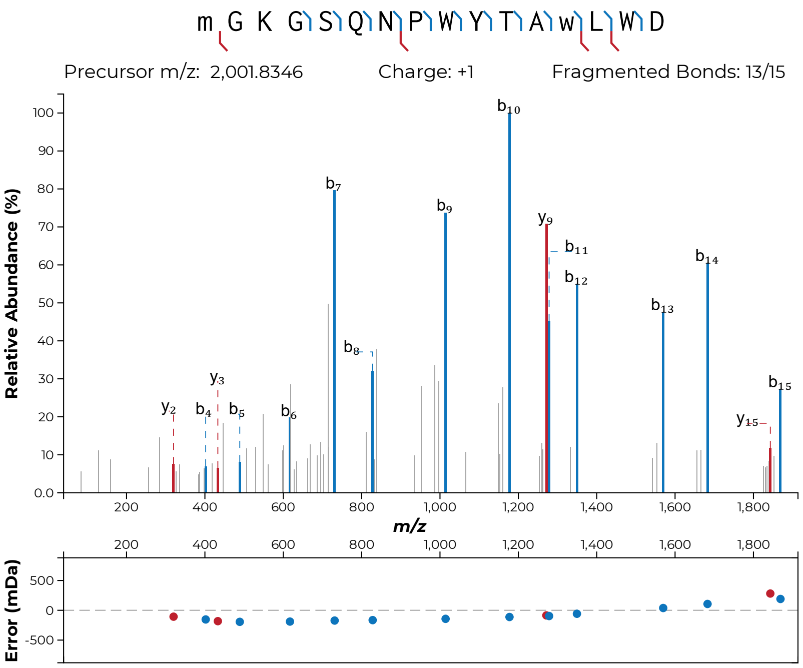

m G K G S Q N P W Y T A w L W D
Precursor m/z: 2,001.8346
Charge: +1
Fragmented Bonds: 13/15
Relative Abundance (%)
m/z
Error (mDa)


**S43s**

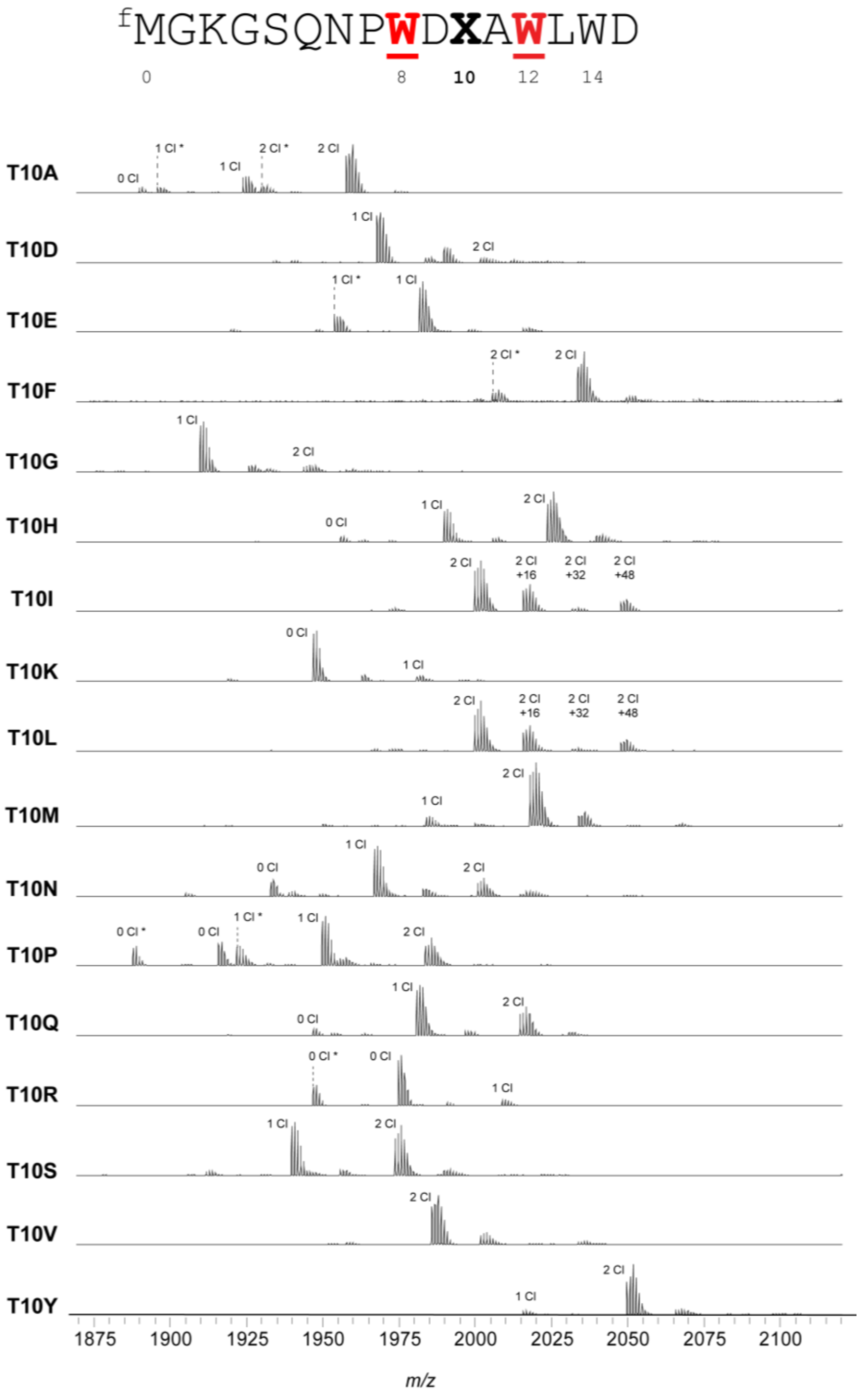


**Figure S44: MALDI-TOF-MS and -LIFT-MS for ChlA$_{PE}$ Thr10 variants.** (**a**) MALDI-TOF-MS spectra for all variants made of position 10 of the ChlA$_{PE}$ substrate. Asterisks (*) indicate peptides lacking N-terminal formylation.

(**b-x, following pages**): MS/MS spectra acquired for this set of substrates. In all MS/MS spectra, lowercase letters indicate modified residues. For example, the N-terminal formylated Met is indicated as “m”. Chlorinated Trp is indicated as “w”.

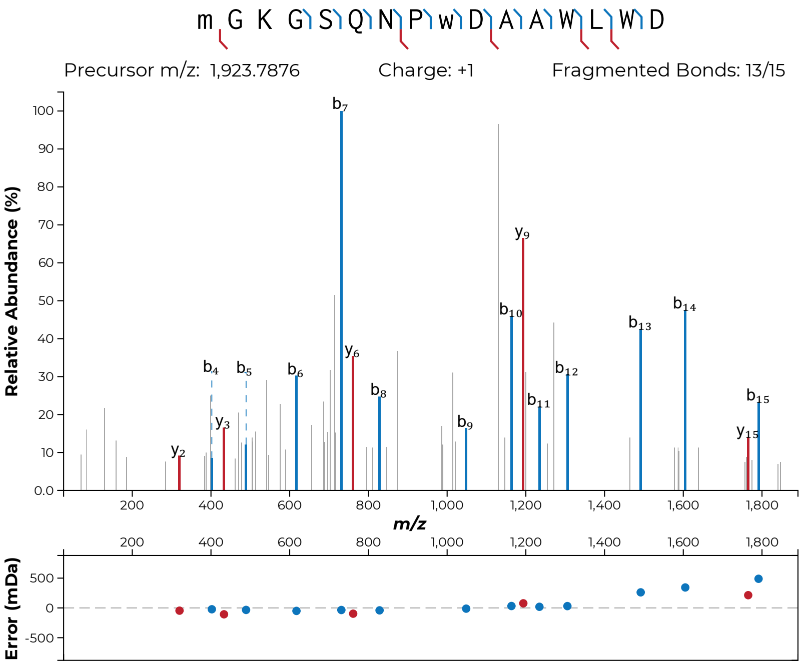
m G K G S Q N P w D A A W L W D
Precursor m/z: 1,923.7876
Charge: +1
Fragmented Bonds: 13/15
Relative Abundance (%)
m/z
Error (mDa)

**S44b**

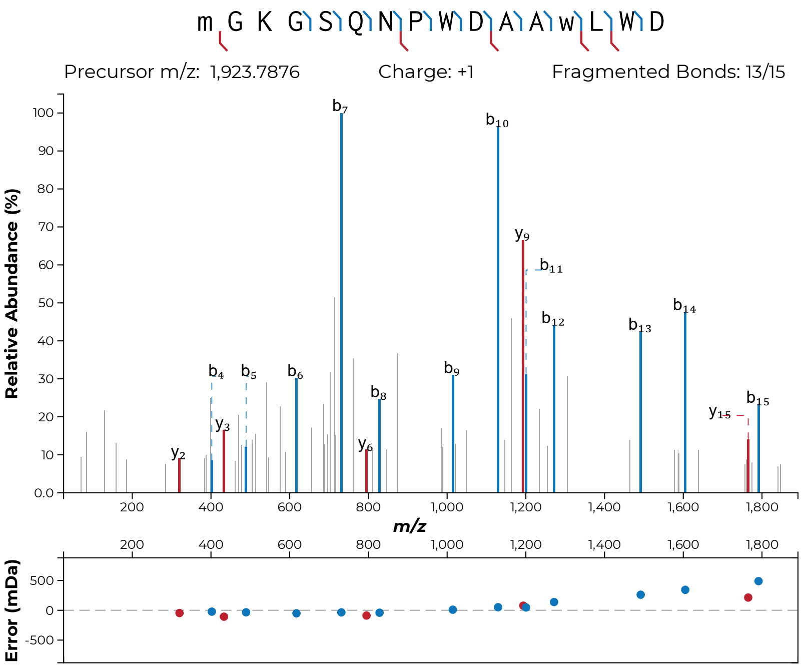
m G K G S Q N P W D A A w L W D
Precursor m/z: 1,923.7876
Charge: +1
Fragmented Bonds: 13/15
Relative Abundance (%)
m/z
Error (mDa)
b7
b10
y9
b11
b12
b13
b14
b4
b5
b6
b9
b8
y3
y2
y6
y15
b15

**S44c**

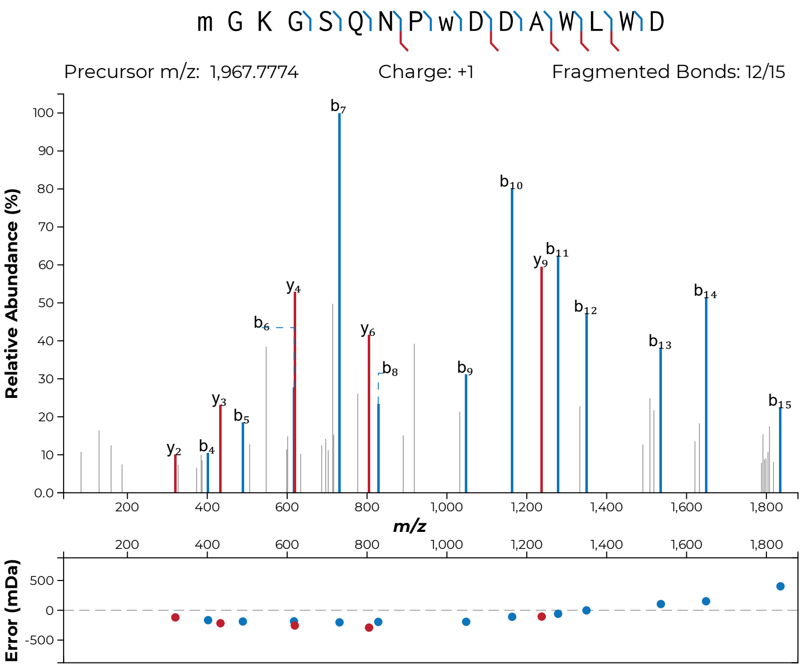
m G K G S Q N P w D D A W L W D
Precursor m/z: 1,967.7774
Charge: +1
Fragmented Bonds: 12/15
Relative Abundance (%)
m/z
Error (mDa)
b7
b10
b11
y9
b12
b14
y4
b6
y6
b8
b9
b13
b15
y3
b5
y2
b4

**S44d**

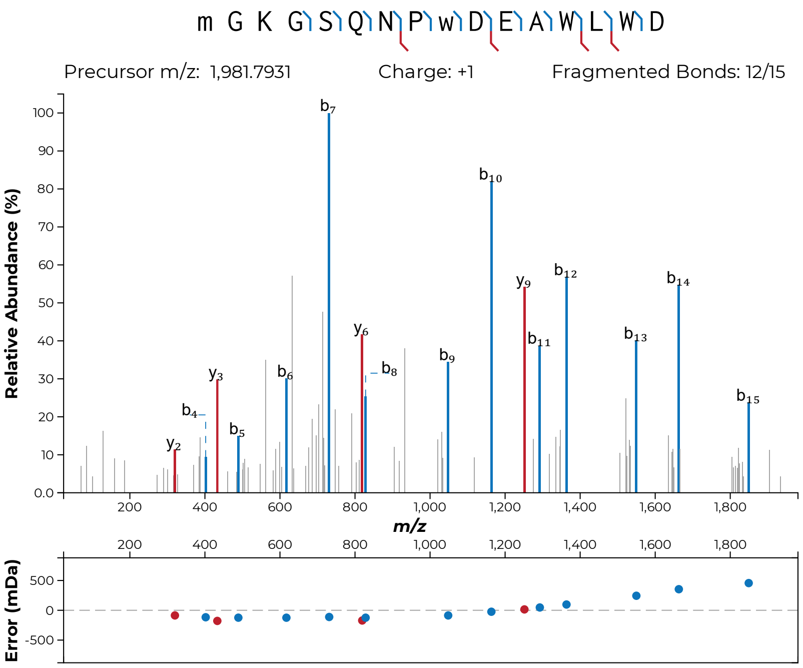
m G K G S Q N P w D E A W L W D
Precursor m/z: 1,981.7931
Charge: +1
Fragmented Bonds: 12/15
Relative Abundance (%)
m/z
Error (mDa)

**S44e**

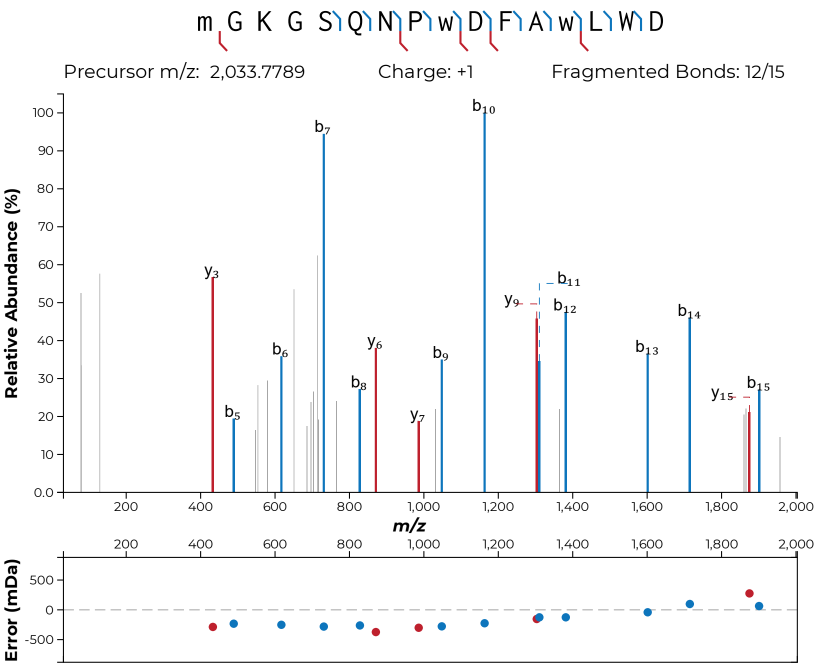
m G K G S Q N P w D F A w L W D
Precursor m/z: 2,033.7789
Charge: +1
Fragmented Bonds: 12/15
Relative Abundance (%)
m/z
Error (mDa)

**S44f**

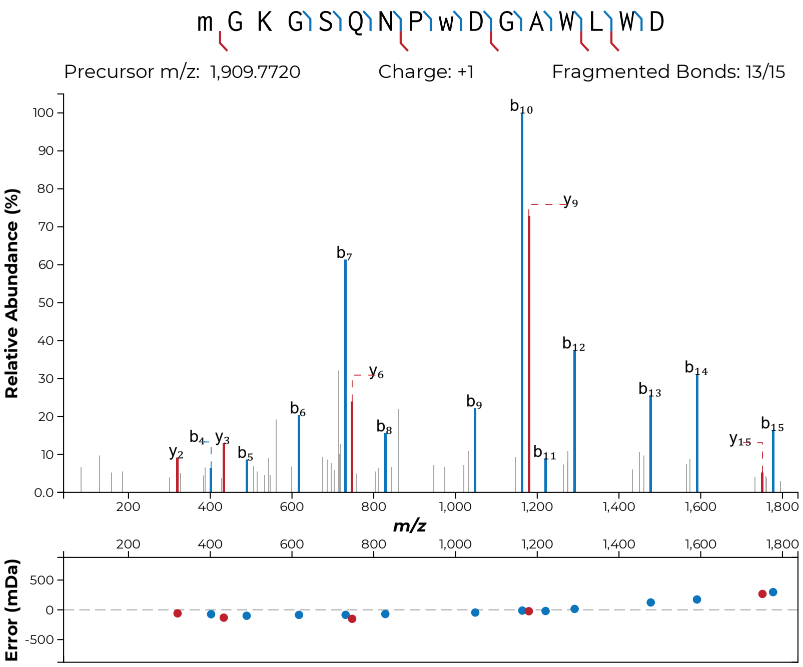

m G K G S Q N P w D G A W L W D
Precursor m/z: 1,909.7720
Charge: +1
Fragmented Bonds: 13/15
Relative Abundance (%)
m/z
Error (mDa)


**S44g**

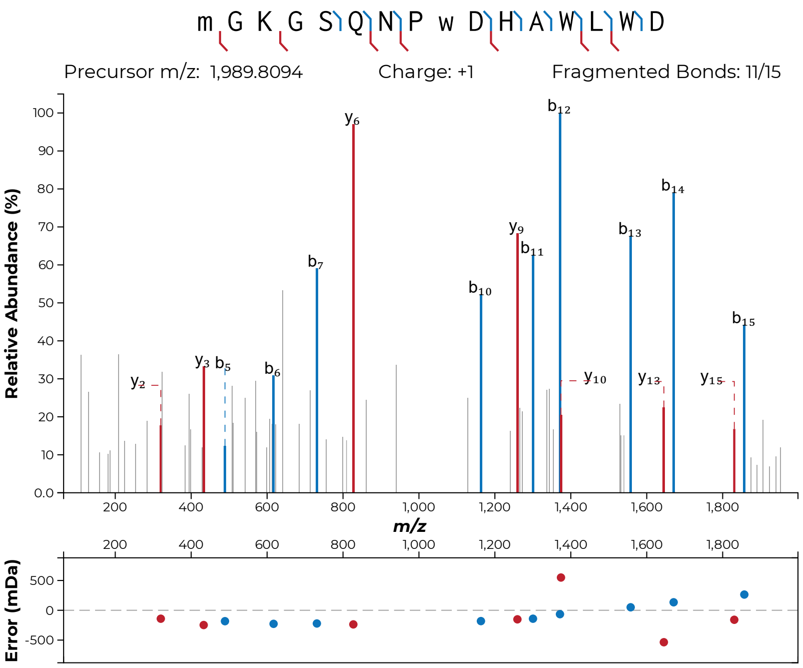
m G K G S Q N P w D H A W L W D
Precursor m/z: 1,989.8094
Charge: +1
Fragmented Bonds: 11/15
Relative Abundance (%)
m/z
Error (mDa)
y2
y3
b5
b6
b7
y6
b10
y9
b11
b12
y10
b13
y13
b14
y15
b15

**S44h**

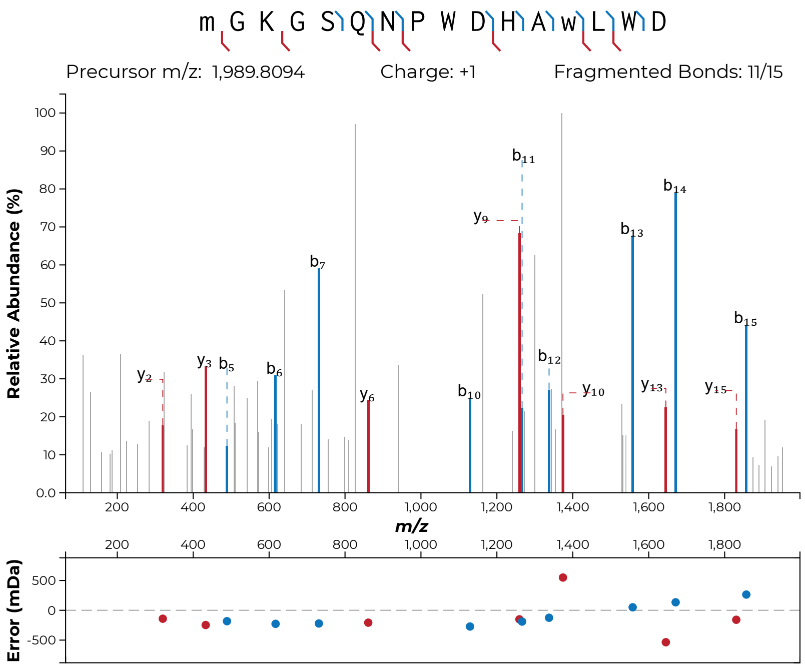

m G K G S Q N P W D H A w L W D
Precursor m/z: 1,989.8094
Charge: +1
Fragmented Bonds: 11/15
Relative Abundance (%)
m/z
Error (mDa)


**S44i**

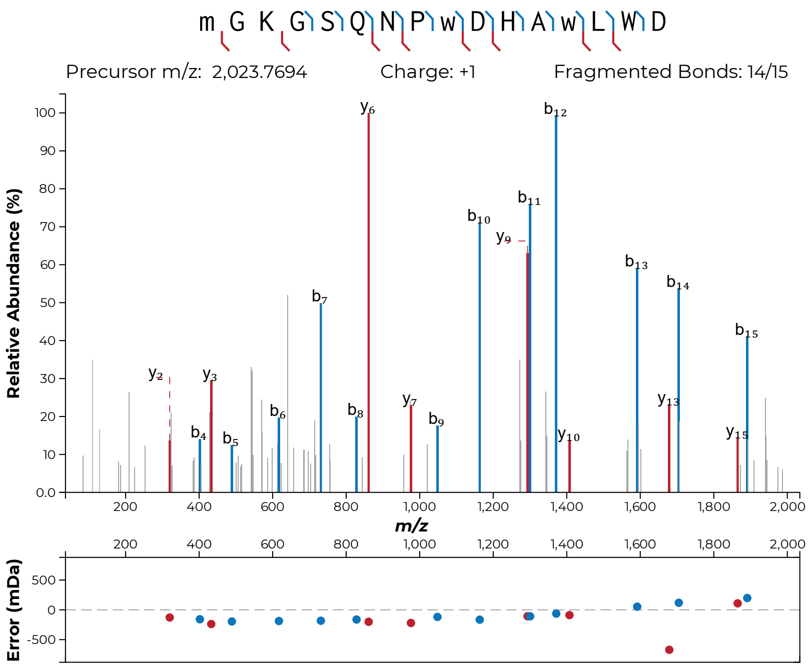
m G K G S Q N P w D H A w L W D
Precursor m/z: 2,023.7694
Charge: +1
Fragmented Bonds: 14/15
Relative Abundance (%)
m/z
Error (mDa)

**S44j**

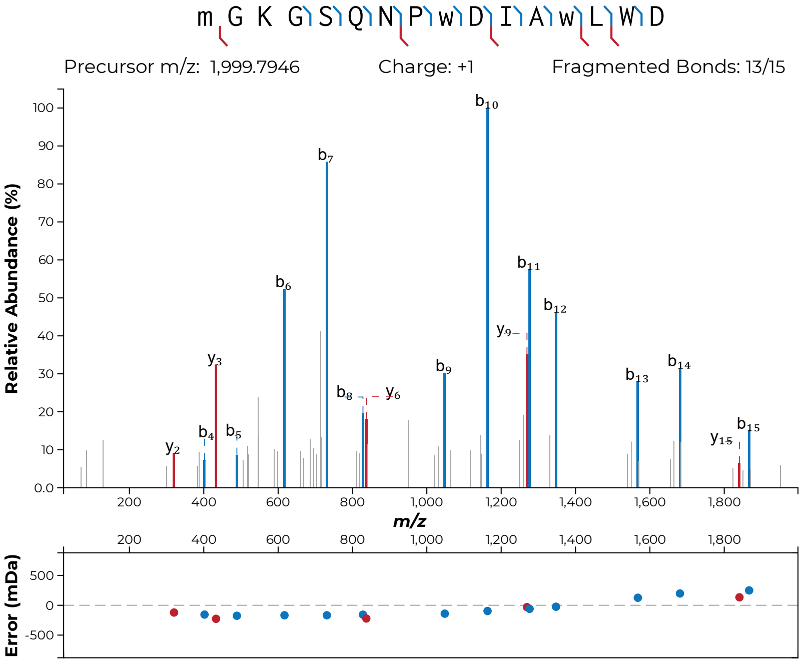
m G K G S Q N P w D I A w L W D
Precursor m/z: 1,999.7946
Charge: +1
Fragmented Bonds: 13/15
Relative Abundance (%)
m/z
Error (mDa)

**S44k**

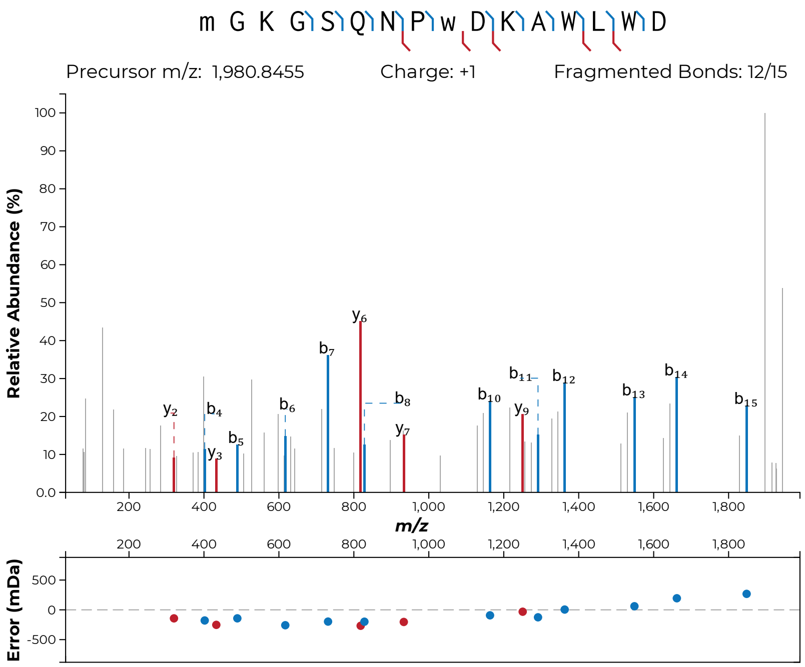
m G K G S Q N P w D K A W L W D
Precursor m/z: 1,980.8455
Charge: +1
Fragmented Bonds: 12/15
Relative Abundance (%)
m/z
Error (mDa)

**S44l**

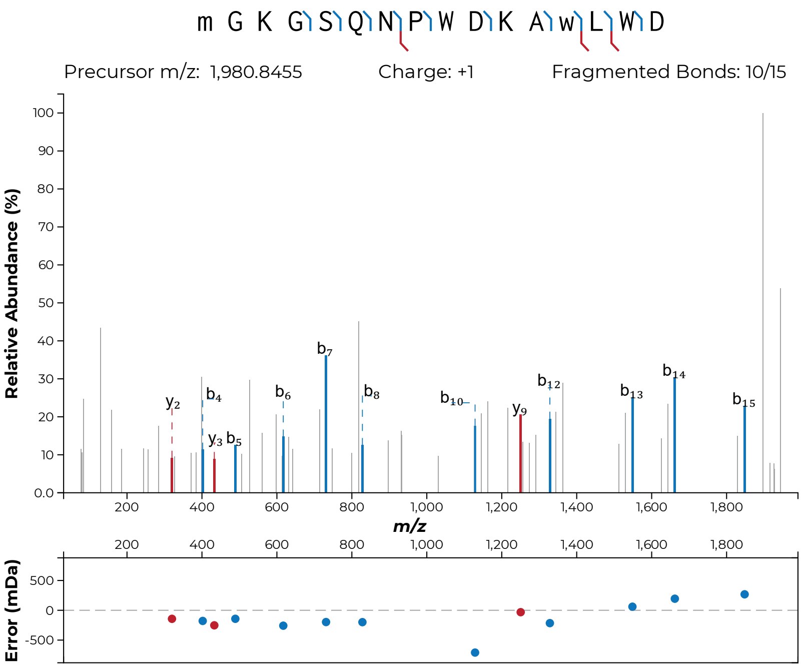
m G K G S Q N P W D K A w L W D
Precursor m/z: 1,980.8455
Charge: +1
Fragmented Bonds: 10/15
Relative Abundance (%)
m/z
Error (mDa)

**S44m**

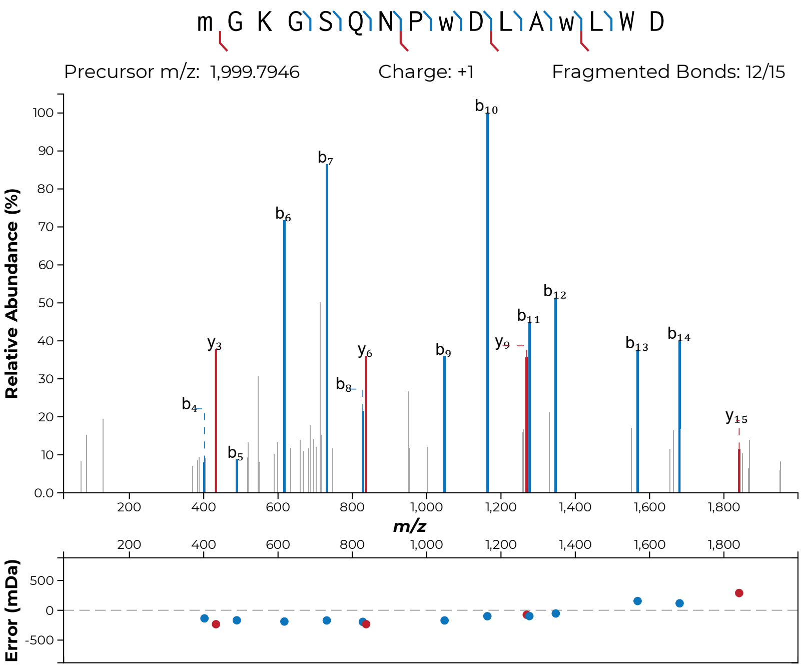

m G K G S Q N P w D L A w L W D
Precursor m/z: 1,999.7946
Charge: +1
Fragmented Bonds: 12/15
Relative Abundance (%)
m/z
Error (mDa)
b4
b5
b6
b7
b8
b9
b10
b11
b12
b13
b14
y3
y6
y9
y15


**S44n**

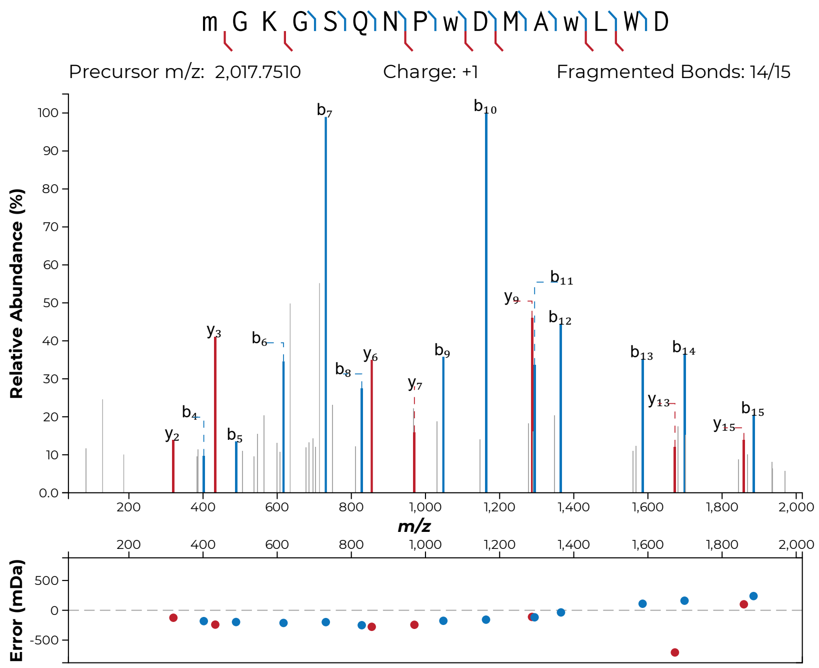
m G K G S Q N P w D M A w L W D
Precursor m/z: 2,017.7510
Charge: +1
Fragmented Bonds: 14/15
Relative Abundance (%)
m/z
Error (mDa)

**S44o**

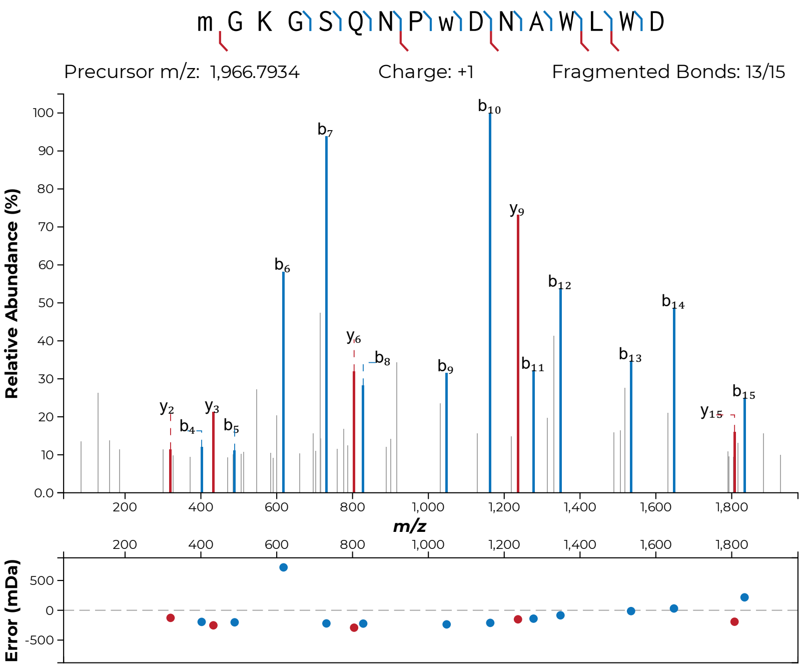
m G K G S Q N P w D N A W L W D
Precursor m/z: 1,966.7934
Charge: +1
Fragmented Bonds: 13/15
Relative Abundance (%)
m/z
Error (mDa)

**S44p**

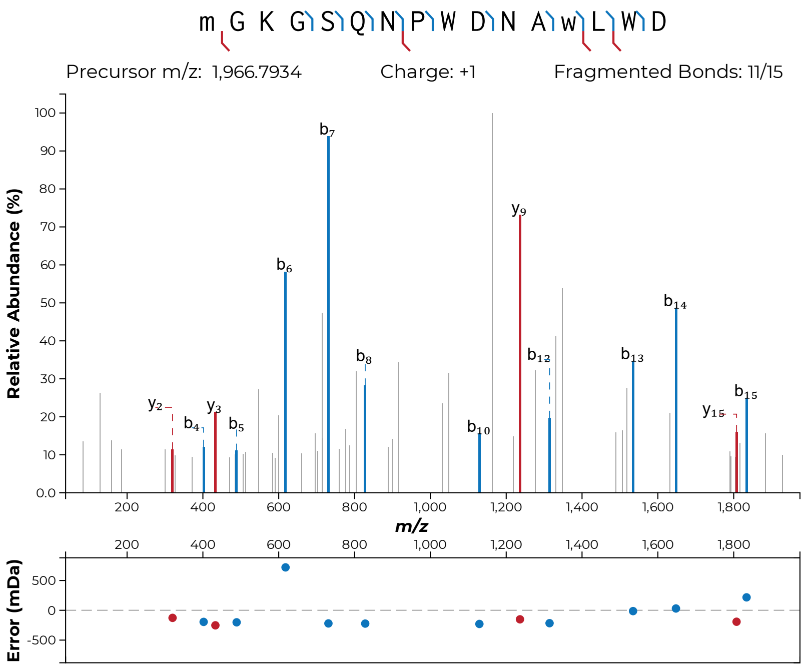
m G K G S Q N P W D N A w L W D
Precursor m/z: 1,966.7934
Charge: +1
Fragmented Bonds: 11/15
Relative Abundance (%)
m/z
Error (mDa)

**S44q**

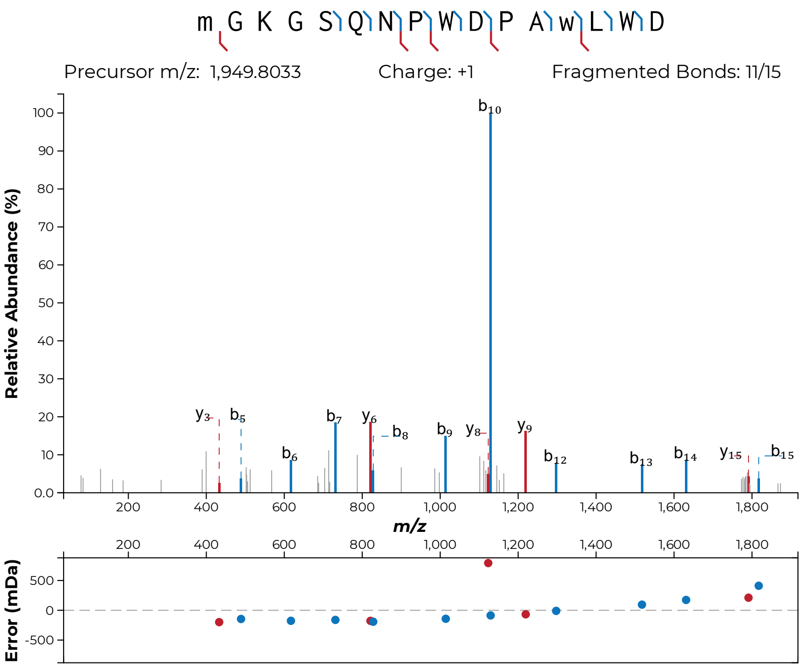
m G K G S Q N P W D P A w L W D
Precursor m/z: 1,949.8033
Charge: +1
Fragmented Bonds: 11/15
Relative Abundance (%)
m/z
Error (mDa)

**S44r**

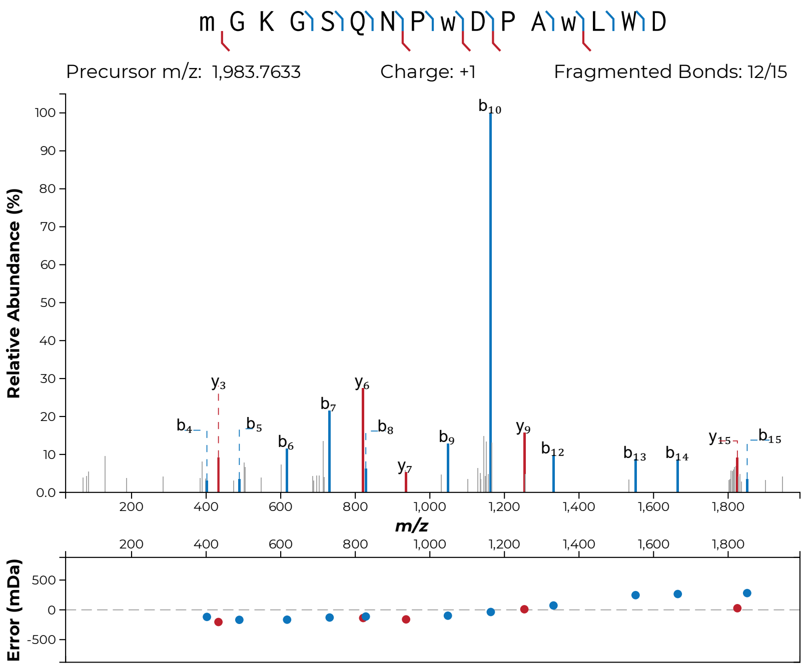

m G K G S Q N P w D P A w L W D
Precursor m/z: 1,983.7633
Charge: +1
Fragmented Bonds: 12/15
Relative Abundance (%)
m/z
Error (mDa)


**S44s**

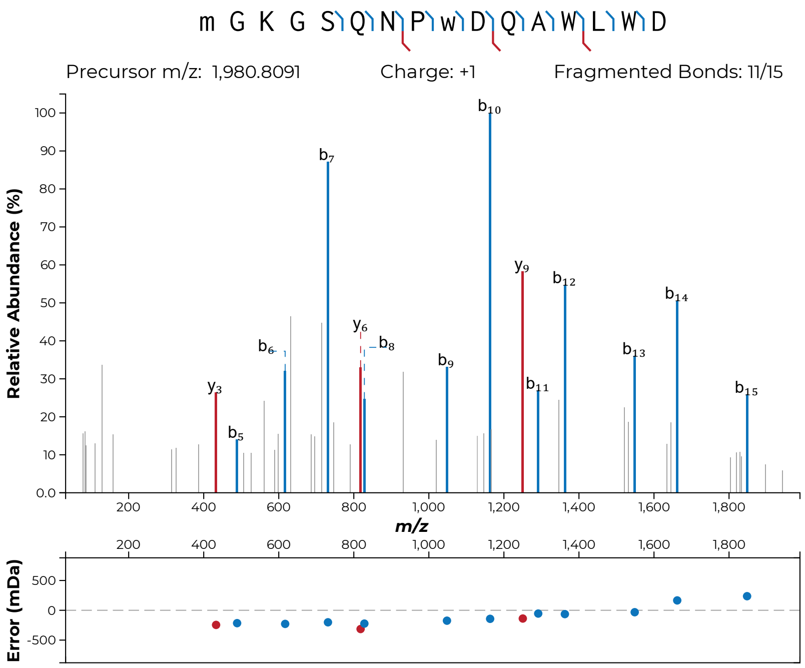
m G K G S Q N P w D Q A W L W D
Precursor m/z: 1,980.8091
Charge: +1
Fragmented Bonds: 11/15
Relative Abundance (%)
m/z
Error (mDa)

**S44t**

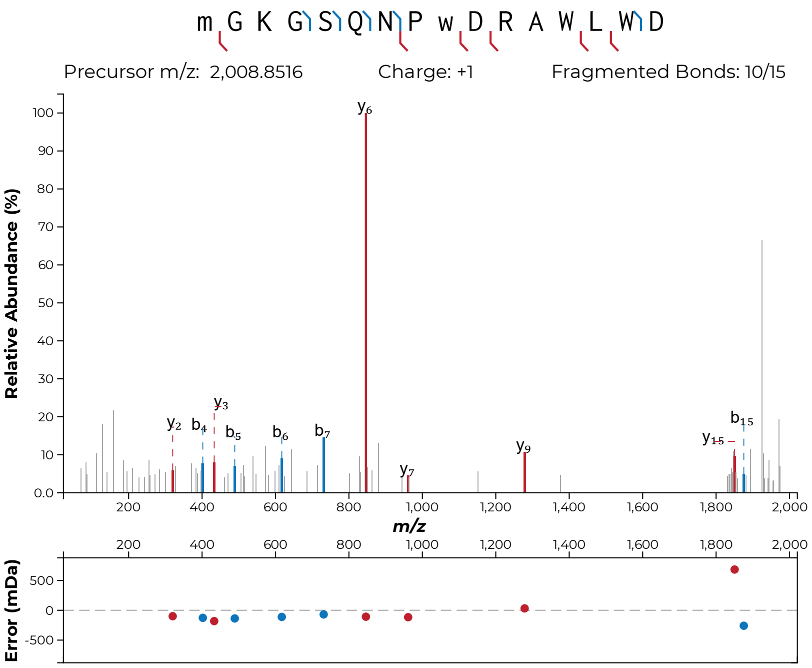

m G K G S Q N P w D R A W L W D
Precursor m/z: 2,008.8516
Charge: +1
Fragmented Bonds: 10/15
Relative Abundance (%)
m/z
Error (mDa)


**S44u**

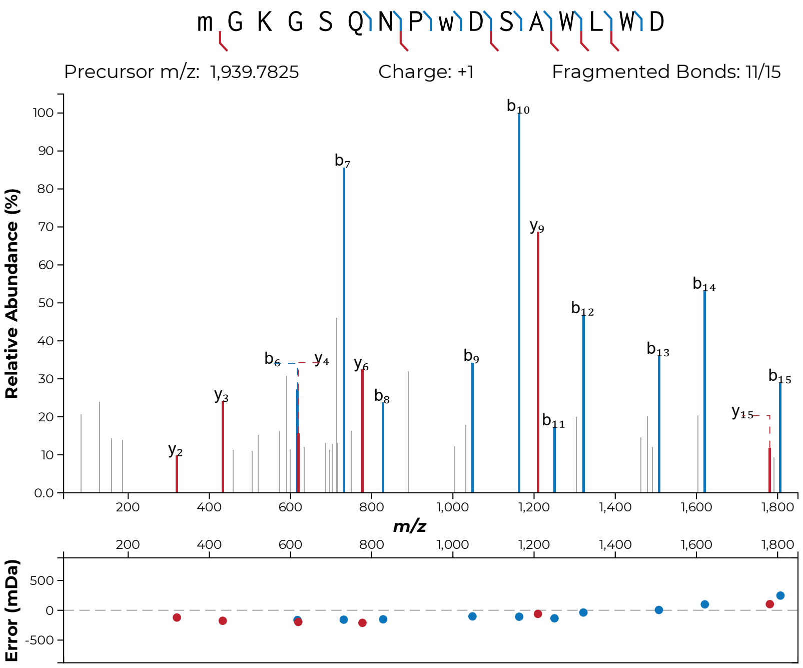
m G K G S Q N P w D S A W L W D
Precursor m/z: 1,939.7825
Charge: +1
Fragmented Bonds: 11/15
Relative Abundance (%)
m/z
Error (mDa)

**S44v**

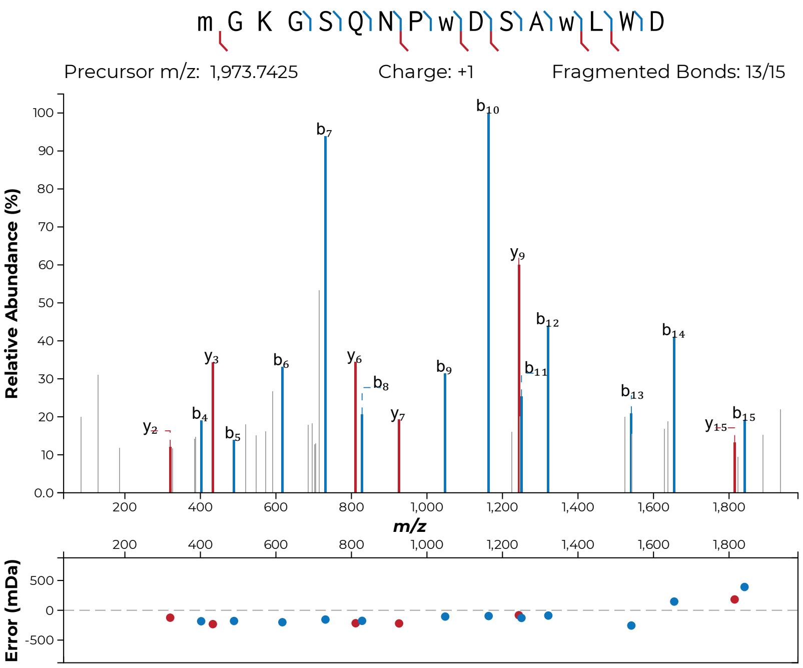
m G K G S Q N P w D S A w L W D
Precursor m/z: 1,973.7425
Charge: +1
Fragmented Bonds: 13/15
Relative Abundance (%)
m/z
Error (mDa)
b7
b10
y9
b12
b14
b11
y3
b6
y6
b8
b9
b13
b4
y7
b15
y2
b5
y15

**S44w**

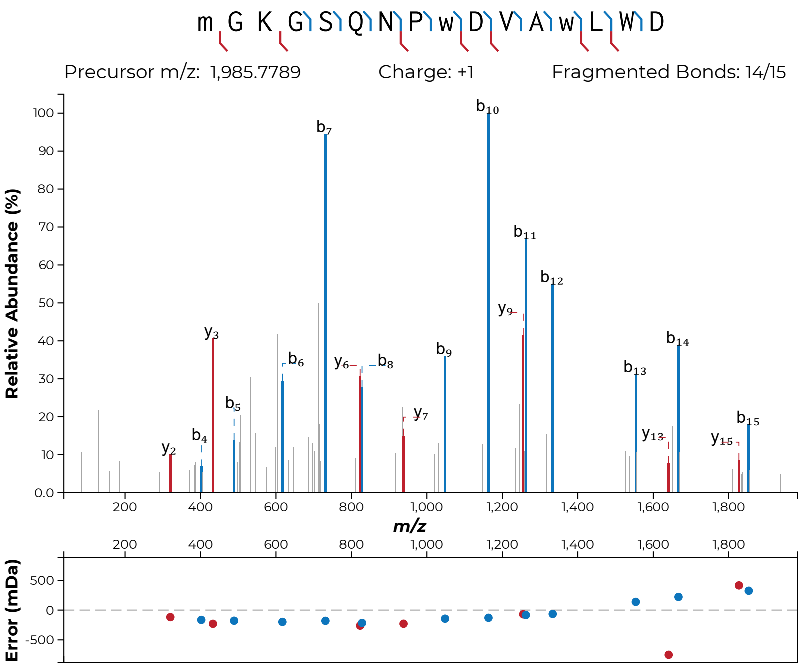

m G K G S Q N P w D V A w L W D
Precursor m/z: 1,985.7789
Charge: +1
Fragmented Bonds: 14/15
Relative Abundance (%)
m/z
Error (mDa)


**S44x**

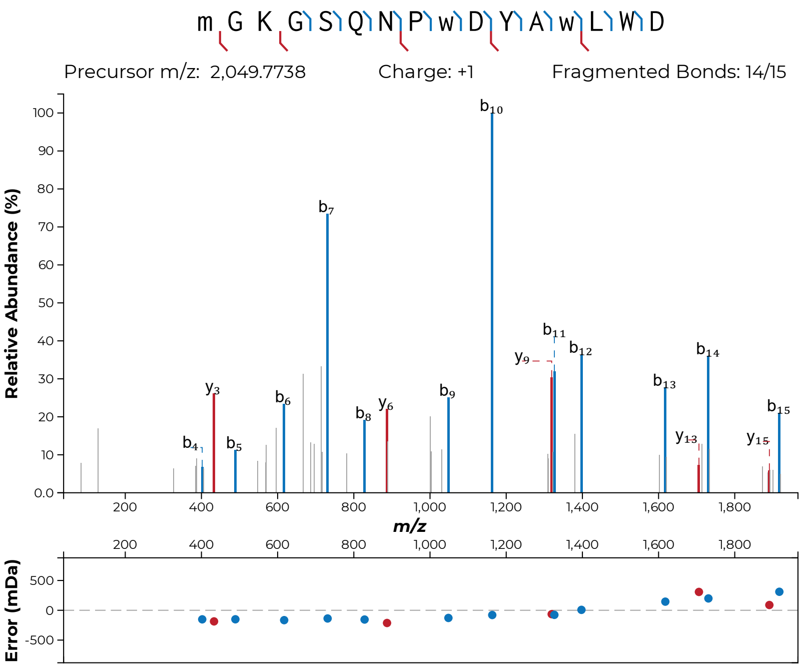

m G K G S Q N P w D Y A w L W D
Precursor m/z: 2,049.7738
Charge: +1
Fragmented Bonds: 14/15
Relative Abundance (%)
m/z
Error (mDa)


**S44y**

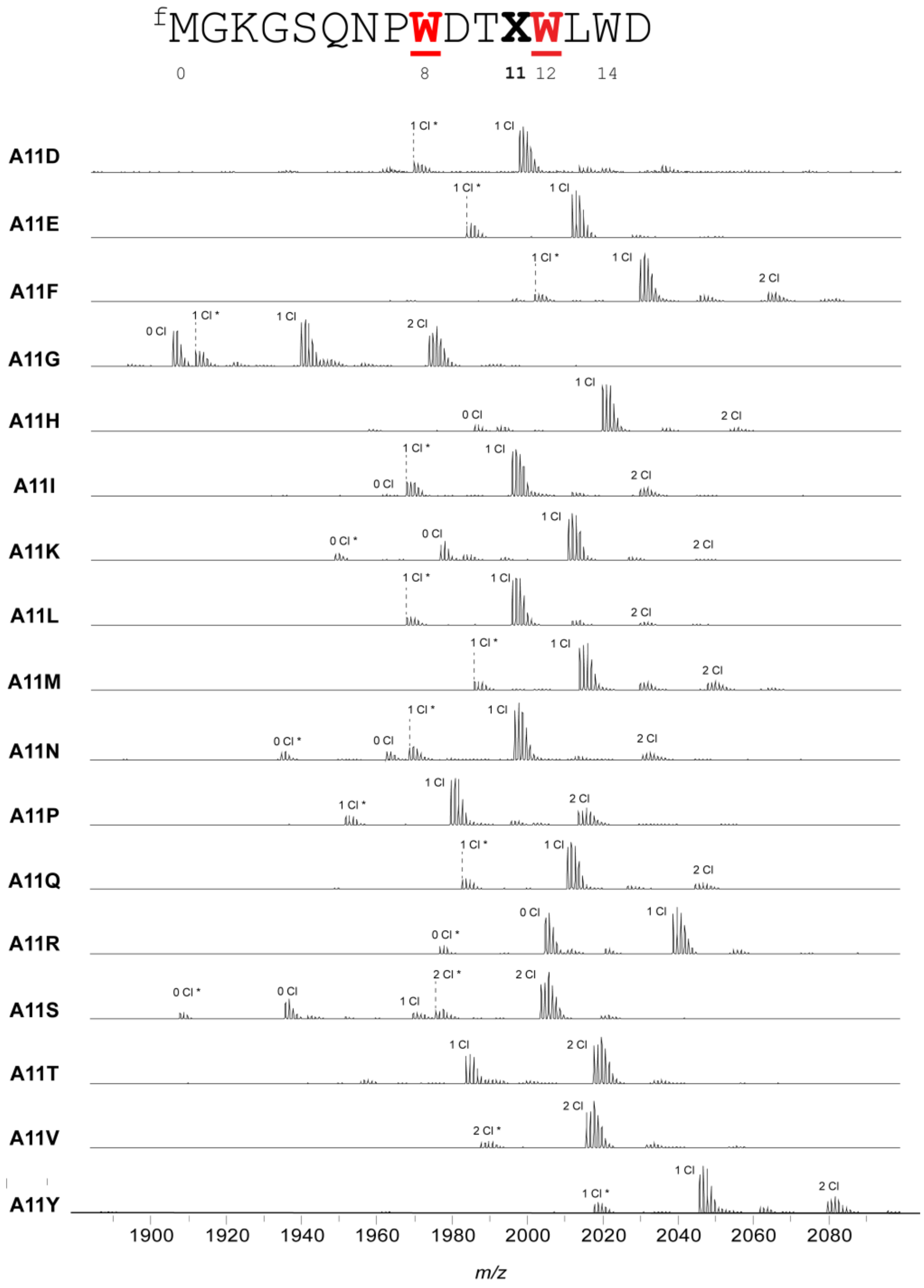


**Figure S45: MALDI-TOF-MS and -LIFT-MS for ChlA$_{PE}$ Ala11 variants.** (**a**) MALDI-TOF-MS spectra for all variants made of position 11 of the ChlA$_{PE}$ substrate. Asterisks (*) indicate peptides lacking N-terminal formylation.

(**b-s, following pages**): MS/MS spectra acquired for this set of substrates. In all MS/MS spectra, lowercase letters indicate modified residues. For example, the N-terminal formylated Met is indicated as “m”. Chlorinated Trp is indicated as “w”.

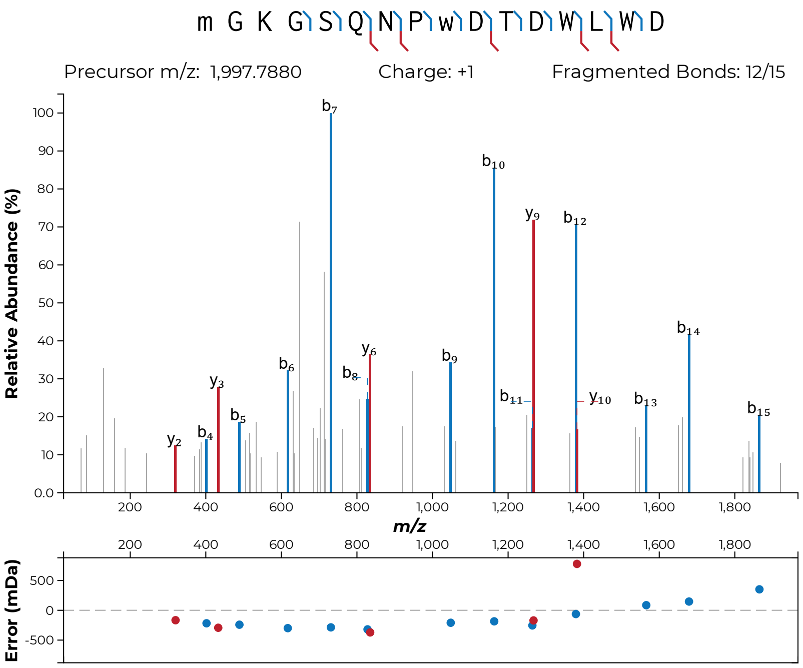

m G K G S Q N P w D T D W L W D
Precursor m/z: 1,997.7880
Charge: +1
Fragmented Bonds: 12/15
Relative Abundance (%)
m/z
Error (mDa)


**S45b**

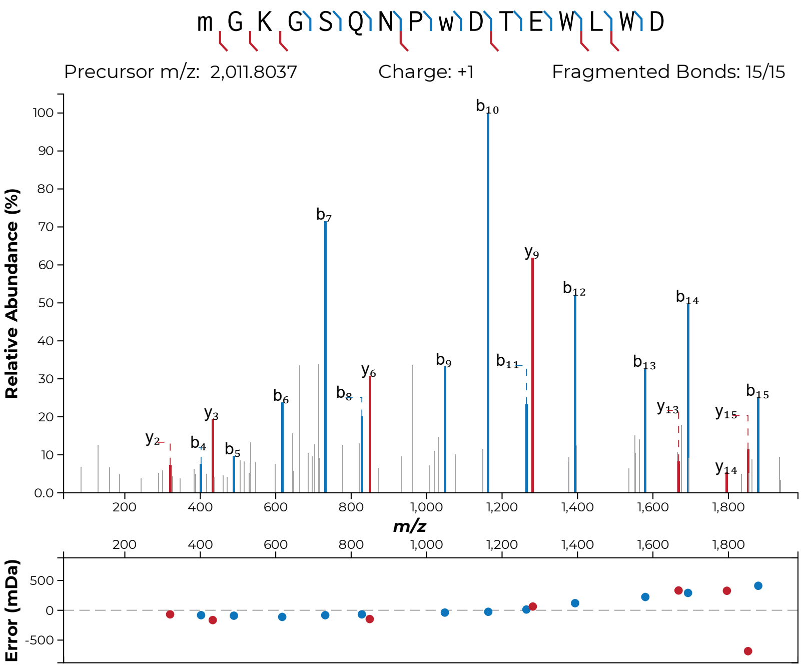

m G K G S Q N P w D T E W L W D
Precursor m/z: 2,011.8037
Charge: +1
Fragmented Bonds: 15/15
Relative Abundance (%)
m/z
Error (mDa)


**S45c**

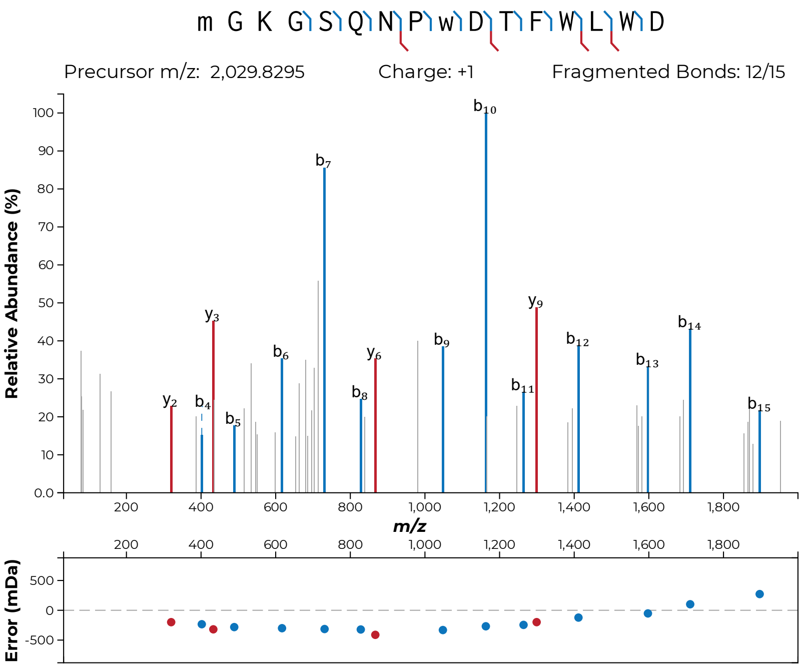
m G K G S Q N P w D T F W L W D
Precursor m/z: 2,029.8295
Charge: +1
Fragmented Bonds: 12/15
Relative Abundance (%)
m/z
Error (mDa)

**S45d**

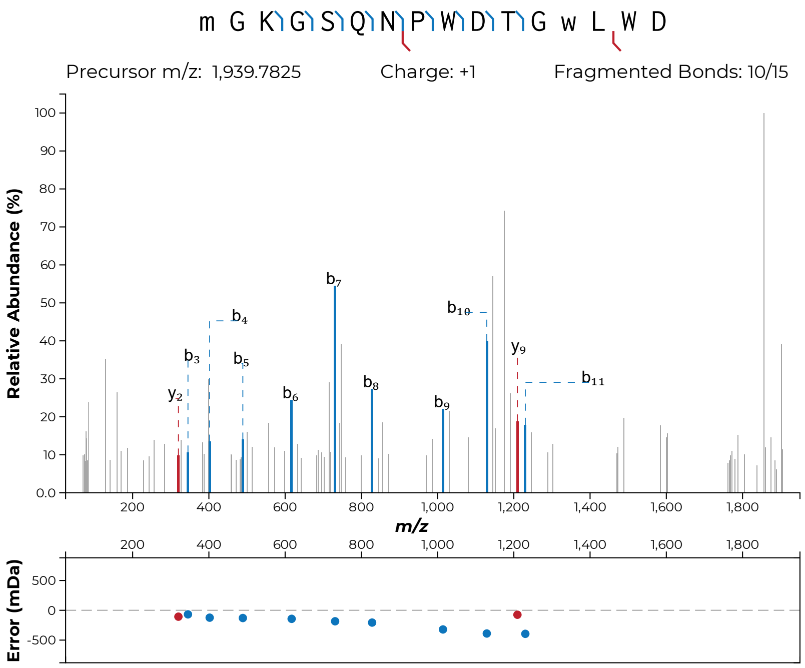
m G K G S Q N P W D T G w L W D
Precursor m/z: 1,939.7825
Charge: +1
Fragmented Bonds: 10/15
Relative Abundance (%)
m/z
Error (mDa)
y2
b3
b4
b5
b6
b7
b8
b9
b10
y9
b11

**S45e**

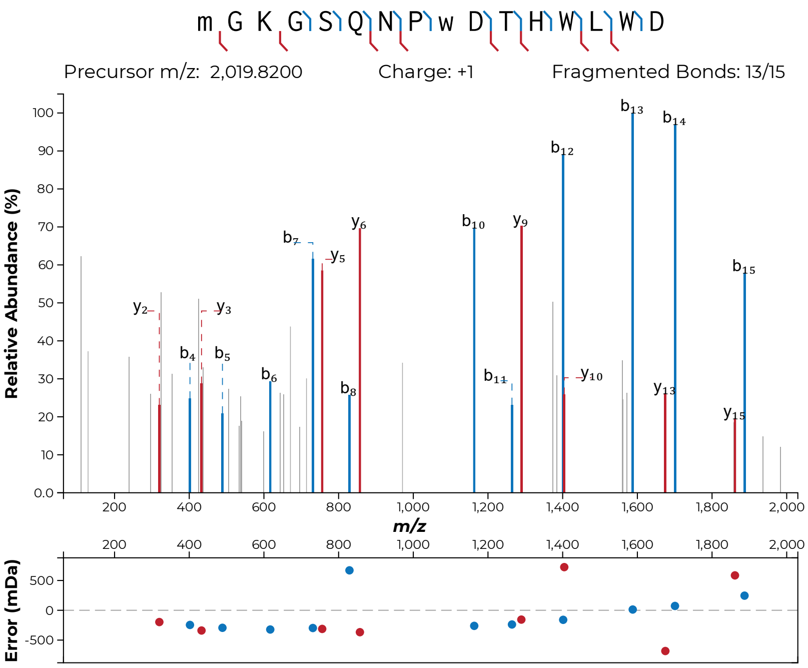

m G K G S Q N P w D T H W L W D
Precursor m/z: 2,019.8200
Charge: +1
Fragmented Bonds: 13/15
Relative Abundance (%)
m/z
Error (mDa)


**S45f**

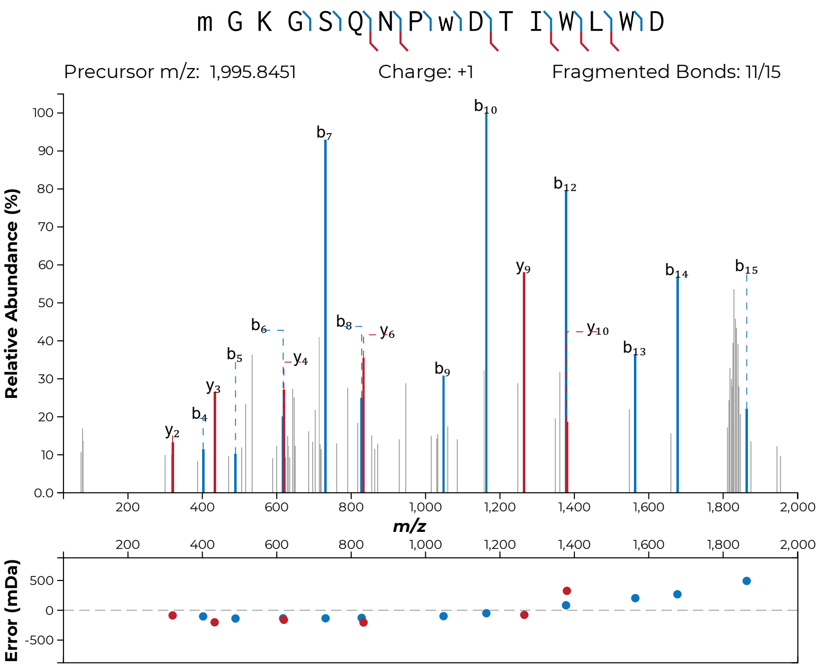
m G K G S Q N P w D T I W L W D
Precursor m/z: 1,995.8451
Charge: +1
Fragmented Bonds: 11/15
Relative Abundance (%)
m/z
Error (mDa)
b2 y2 y3 b4 b5 b6 y4 b7 b8 y6 b9 b10 y9 b12 y10 b13 b14 b15

**S45g**

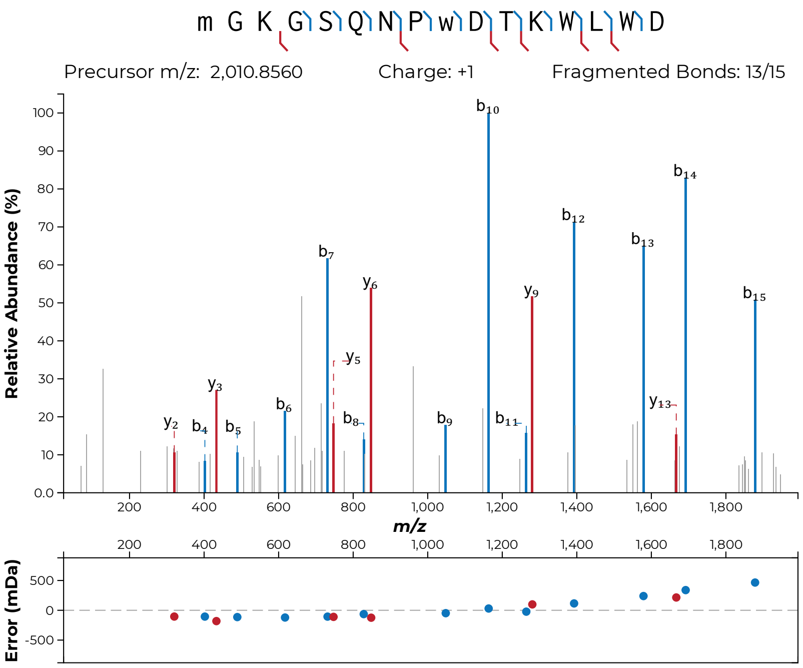

m G K G S Q N P w D T K W L W D
Precursor m/z: 2,010.8560
Charge: +1
Fragmented Bonds: 13/15
Relative Abundance (%)
m/z
Error (mDa)
b10
b14
b12
b13
b7
y6
y9
b15
y5
y3
b6
y13
y2
b4
b5
b8
b9
b11


**S45h**

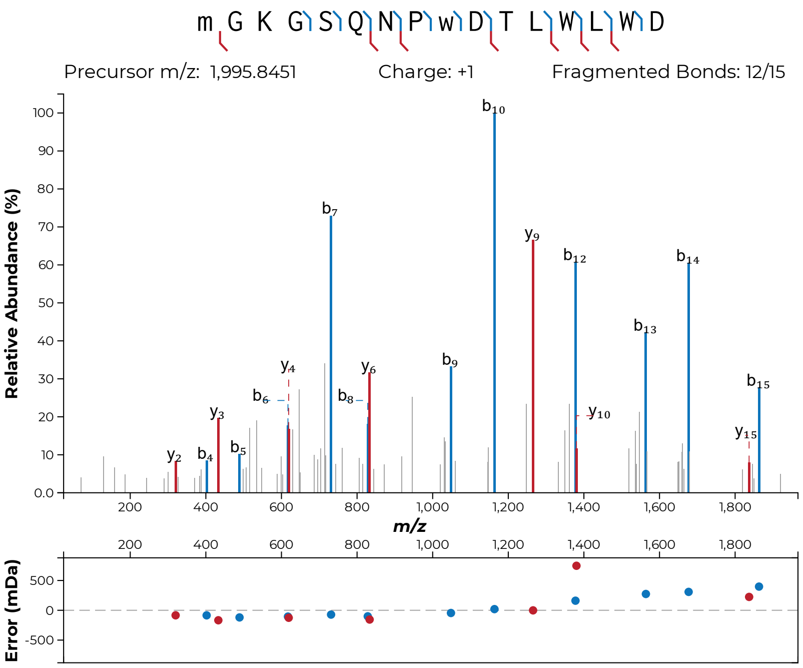
m G K G S Q N P w D T L W L W D
Precursor m/z: 1,995.8451
Charge: +1
Fragmented Bonds: 12/15
Relative Abundance (%)
m/z
Error (mDa)

**S45i**

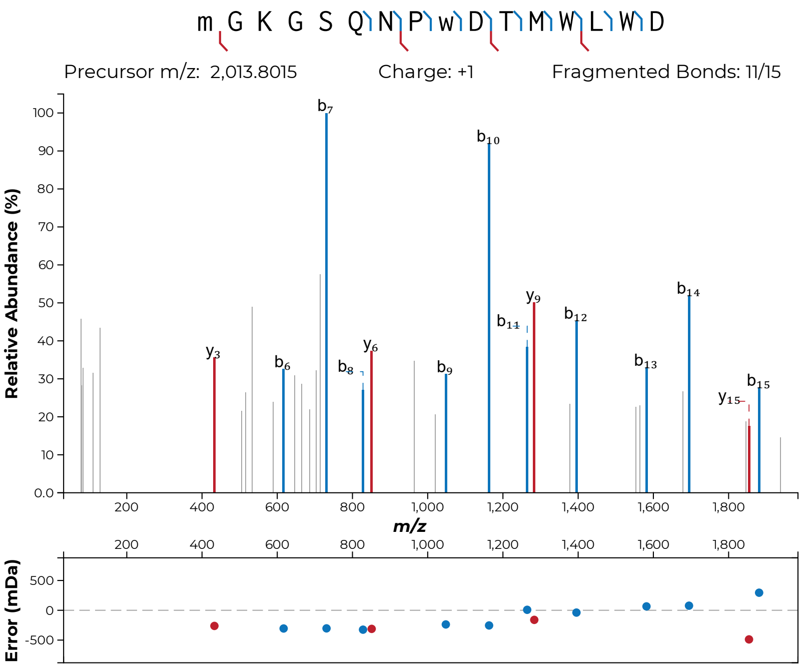
m G K G S Q N P w D T M W L W D
Precursor m/z: 2,013.8015
Charge: +1
Fragmented Bonds: 11/15
Relative Abundance (%)
m/z
Error (mDa)

**S45j**

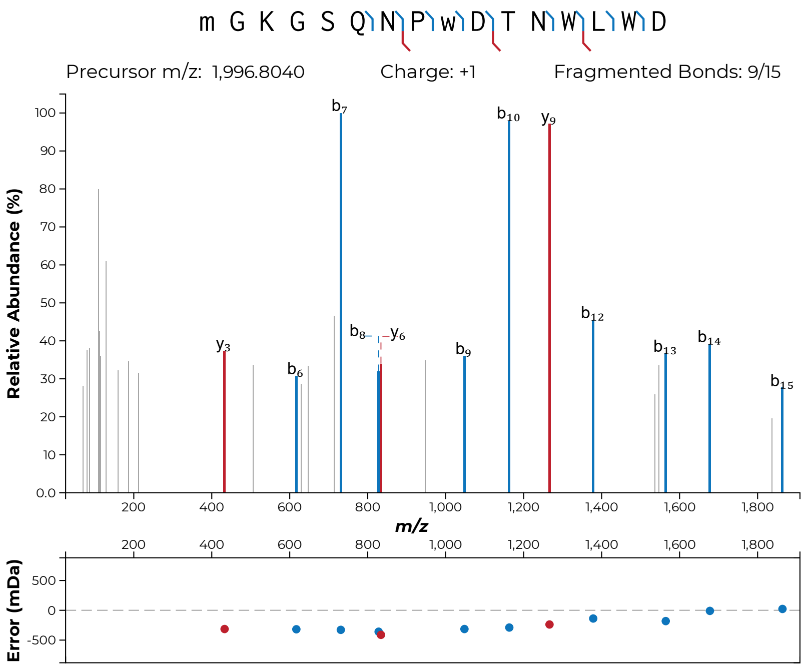
m G K G S Q N P w D T N W L W D
Precursor m/z: 1,996.8040
Charge: +1
Fragmented Bonds: 9/15
Relative Abundance (%)
m/z
Error (mDa)

**S45k**

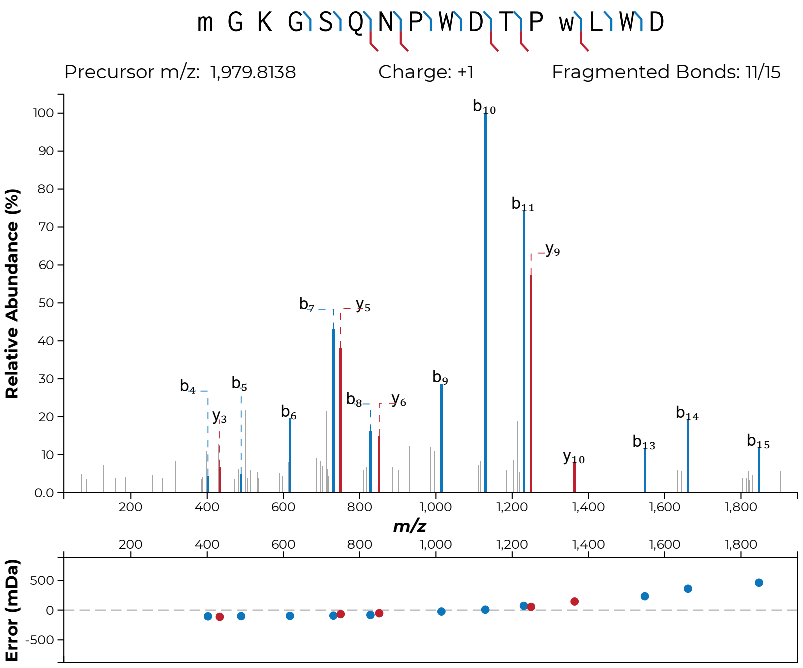
m G K G S Q N P W D T P w L W D
Precursor m/z: 1,979.8138
Charge: +1
Fragmented Bonds: 11/15
Relative Abundance (%)
m/z
Error (mDa)

**S45l**

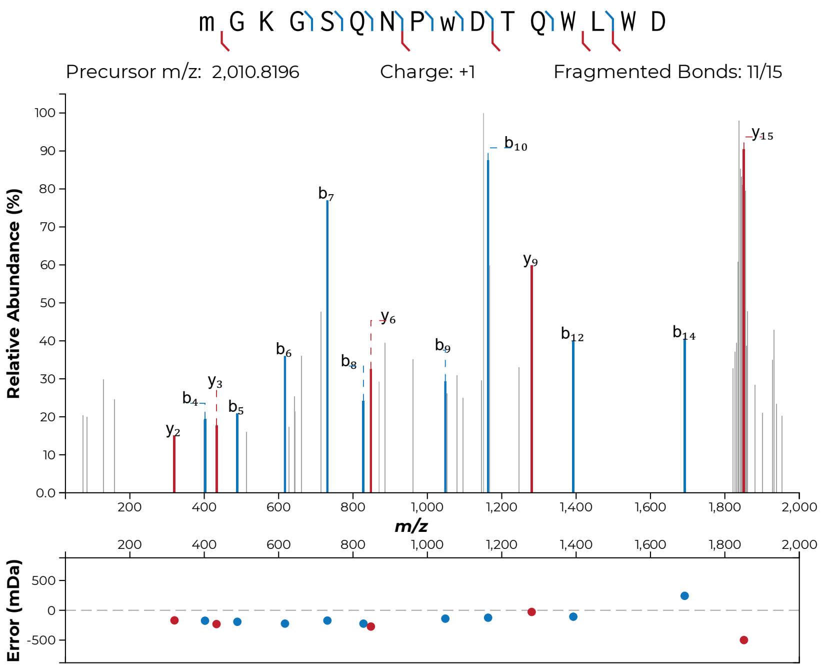
m G K G S Q N P w D T Q W L W D
Precursor m/z: 2,010.8196
Charge: +1
Fragmented Bonds: 11/15
Relative Abundance (%)
m/z
Error (mDa)
y2
b4
y3
b5
b6
b7
b8
y6
b9
b10
y9
b12
b14
y15

**S45m**

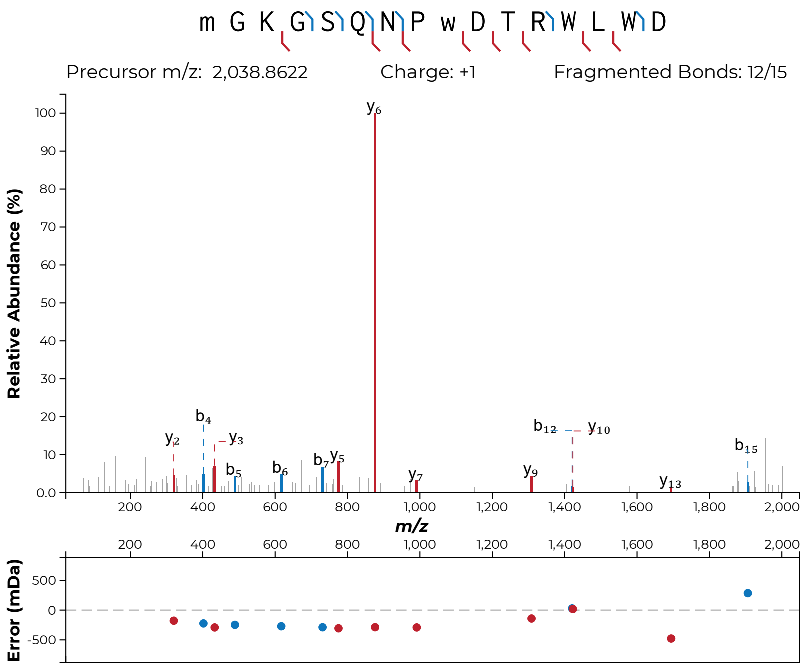
m G K G S Q N P w D T R W L W D
Precursor m/z: 2,038.8622
Charge: +1
Fragmented Bonds: 12/15
Relative Abundance (%)
m/z
Error (mDa)

**S45n**

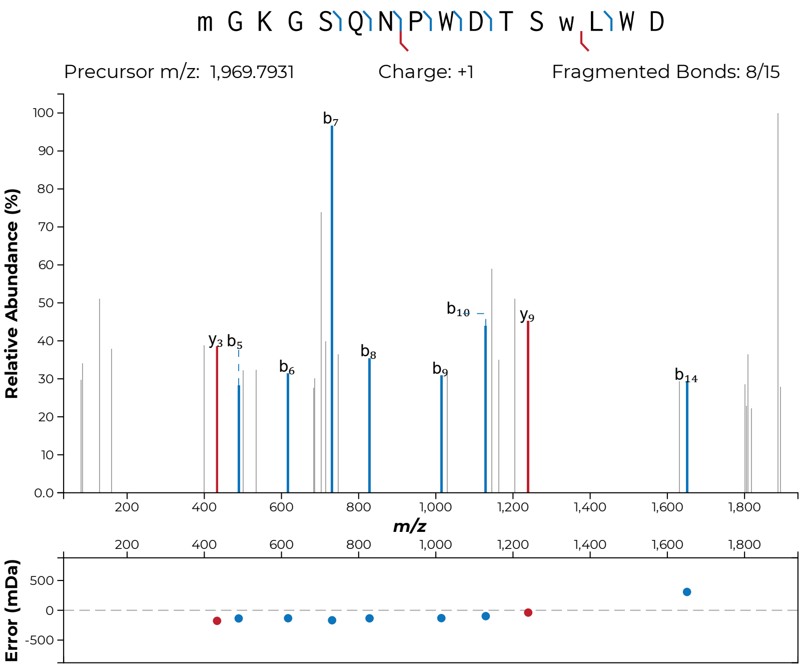

m G K G S Q N P W D T S w L W D
Precursor m/z: 1,969.7931
Charge: +1
Fragmented Bonds: 8/15
Relative Abundance (%)
m/z
Error (mDa)


**S45o**

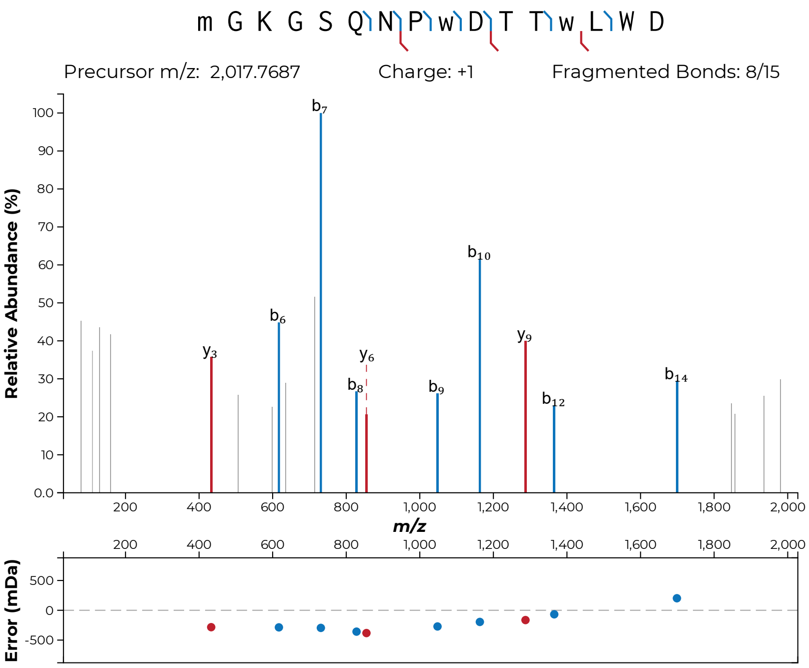
m G K G S Q N P w D T T w L W D
Precursor m/z: 2,017.7687
Charge: +1
Fragmented Bonds: 8/15
Relative Abundance (%)
m/z
Error (mDa)
b7
b10
b6
y9
y3
y6
b8
b9
b12
b14

**S45p**

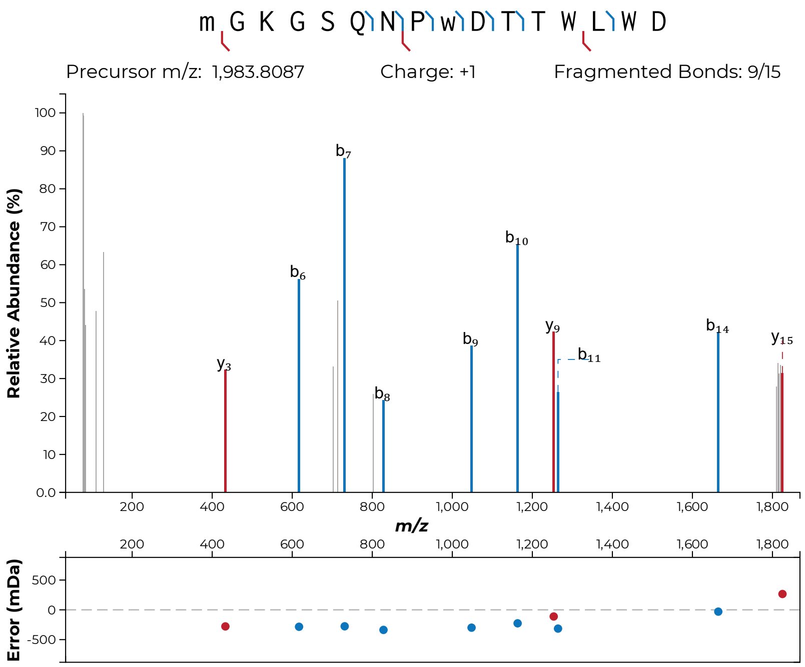

m G K G S Q N P w D T T W L W D
Precursor m/z: 1,983.8087
Charge: +1
Fragmented Bonds: 9/15
Relative Abundance (%)
m/z
Error (mDa)


**S45q**

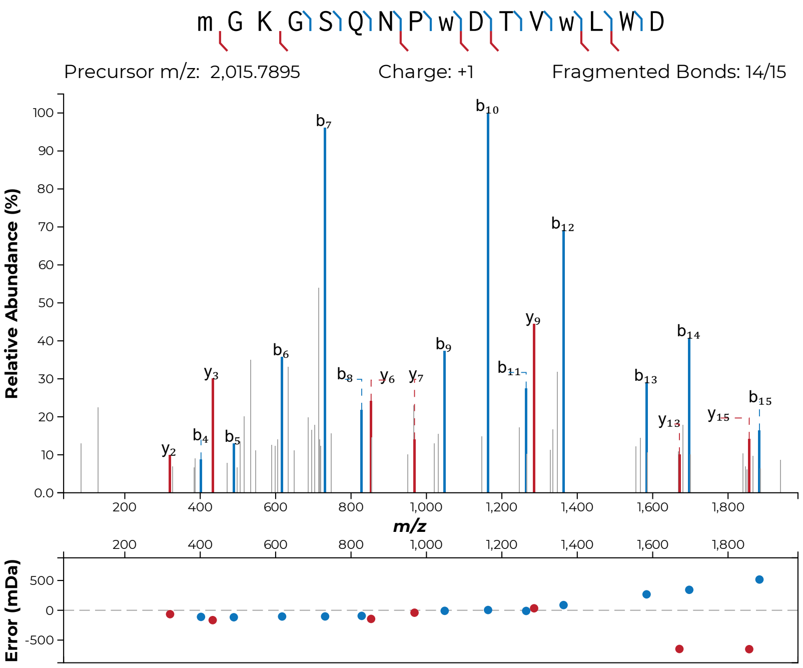
m G K G S Q N P w D T V w L W D
Precursor m/z: 2,015.7895
Charge: +1
Fragmented Bonds: 14/15
Relative Abundance (%)
m/z
Error (mDa)

**S45r**

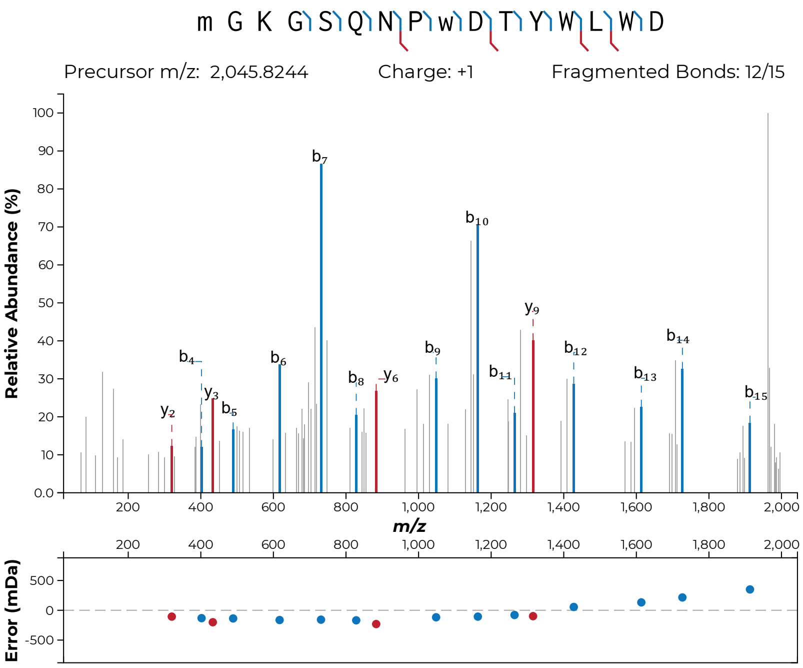
m G K G S Q N P w D T Y W L W D
Precursor m/z: 2,045.8244
Charge: +1
Fragmented Bonds: 12/15
Relative Abundance (%)
m/z
Error (mDa)

**S45s**

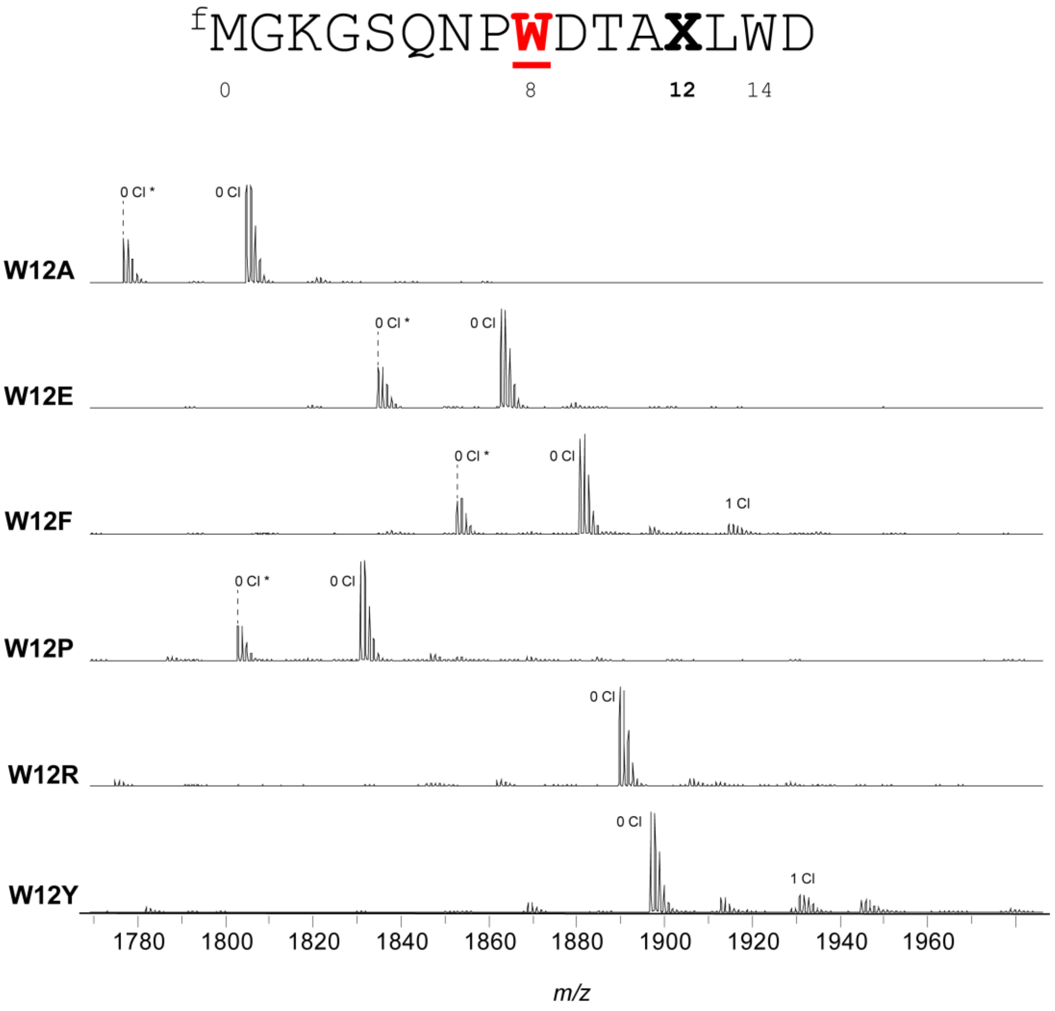


**Figure S46: MALDI-TOF-MS and -LIFT-MS for ChlA$_{PE}$ Trp12 variants.** (**a**) MALDI-TOF-MS spectra for all variants made of position 12 of the ChlA$_{PE}$ substrate. Asterisks (*) indicate peptides lacking N-terminal formylation.

(**b-c, following pages**): MS/MS spectra acquired for this set of substrates. In all MS/MS spectra, lowercase letters indicate modified residues. For example, the N-terminal formylated Met is indicated as "m". Chlorinated Trp is indicated as "w".

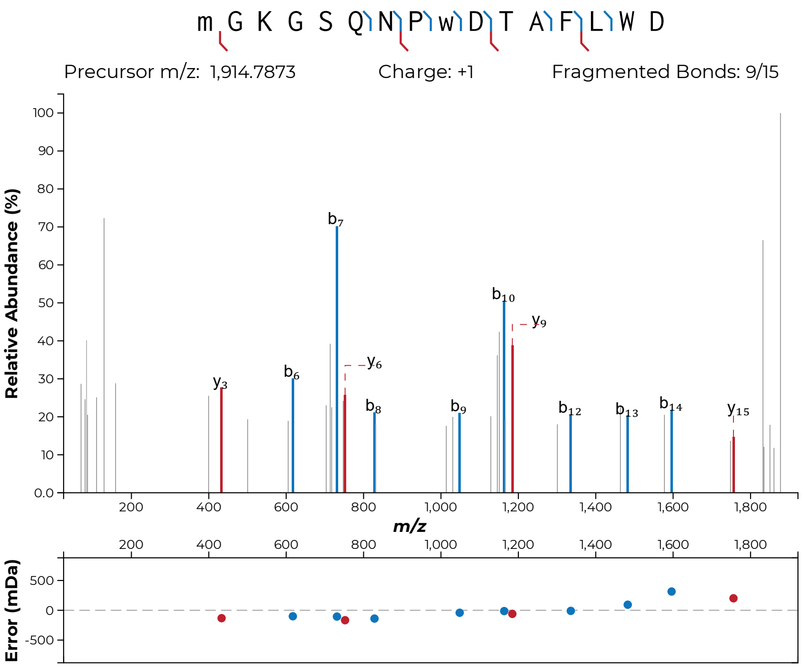
m G K G S Q N P w D T A F L W D
Precursor m/z: 1,914.7873
Charge: +1
Fragmented Bonds: 9/15
Relative Abundance (%)
m/z
Error (mDa)

**S46b**

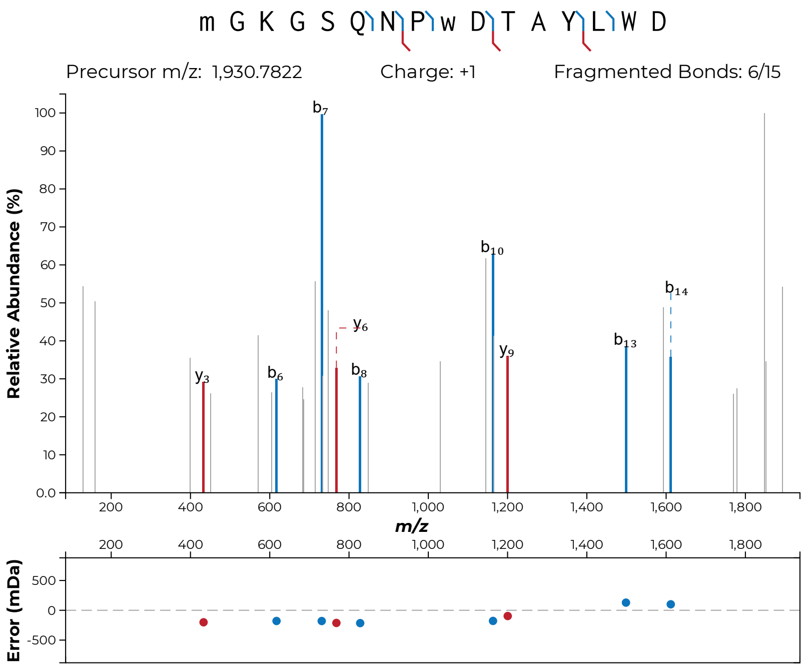
m G K G S Q N P w D T A Y L W D
Precursor m/z: 1,930.7822
Charge: +1
Fragmented Bonds: 6/15
Relative Abundance (%)
m/z
Error (mDa)
b7
b10
b14
b13
y6
y9
y3
b6
b8

**S46c**

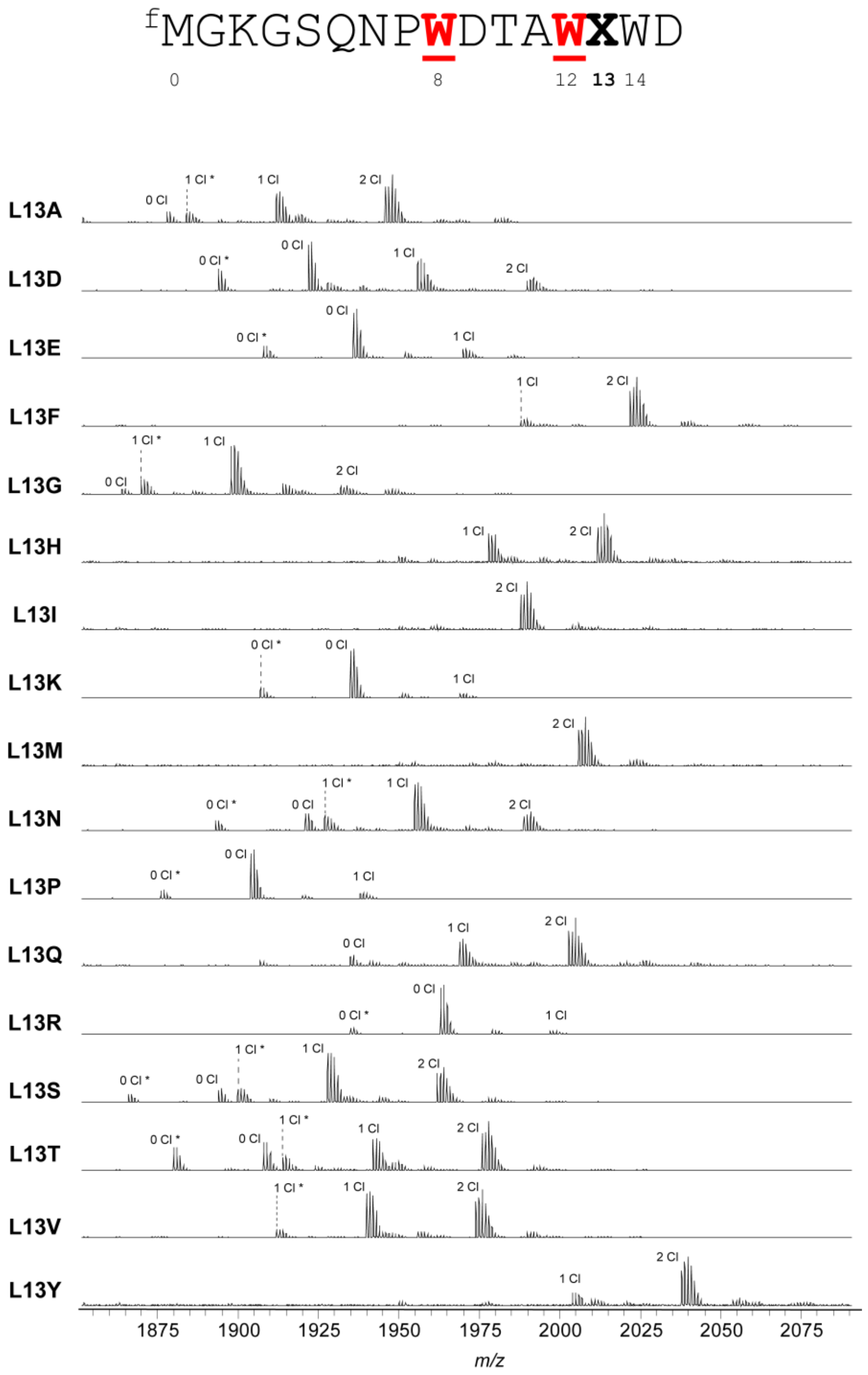


**Figure S47: MALDI-TOF-MS and -LIFT-MS for ChlA$_{PE}$ Leu13 variants.** (**a**) MALDI-TOF-MS spectra for all variants made of position 13 of the ChlA$_{PE}$ substrate. Asterisks (*) indicate peptides lacking N-terminal formylation.

(**b-aa, following pages**): MS/MS spectra acquired for this set of substrates. In all MS/MS spectra, lowercase letters indicate modified residues. For example, the N-terminal formylated Met is indicated as "m". Chlorinated Trp is indicated as "w".

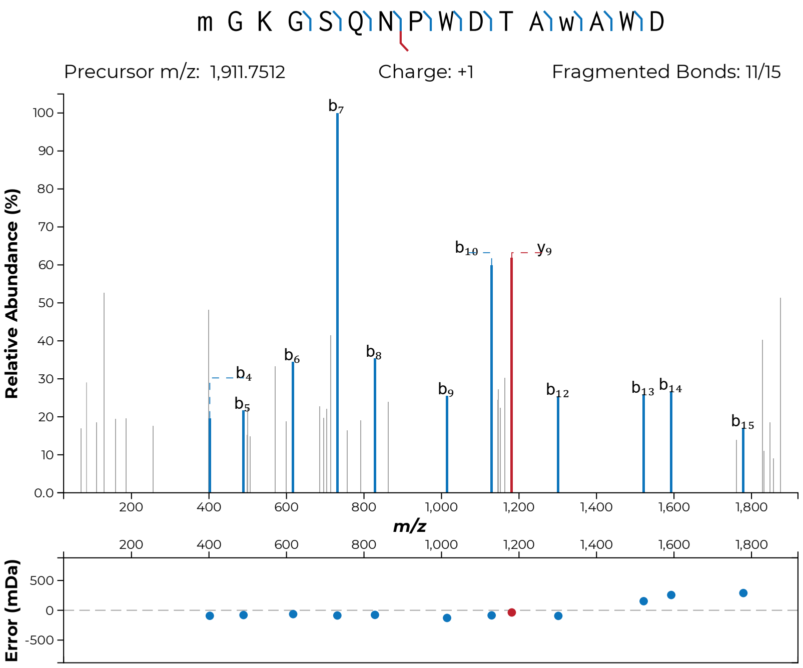
m G K G S Q N P W D T A w A W D
Precursor m/z: 1,911.7512
Charge: +1
Fragmented Bonds: 11/15
Relative Abundance (%)
m/z
Error (mDa)
b7
b10
y9
b4
b5
b6
b8
b9
b12
b13
b14
b15

**S47b**

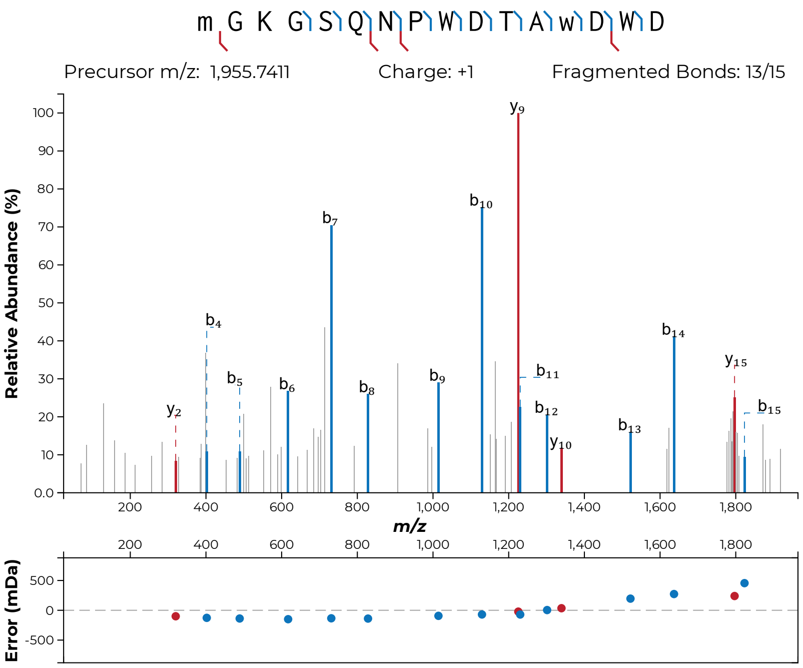
m G K G S Q N P W D T A w D W D
Precursor m/z: 1,955.7411
Charge: +1
Fragmented Bonds: 13/15
Relative Abundance (%)
m/z
Error (mDa)
y2
b4
b5
b6
b7
b8
b9
b10
y9
b11
b12
y10
b13
b14
y15
b15

**S47c**

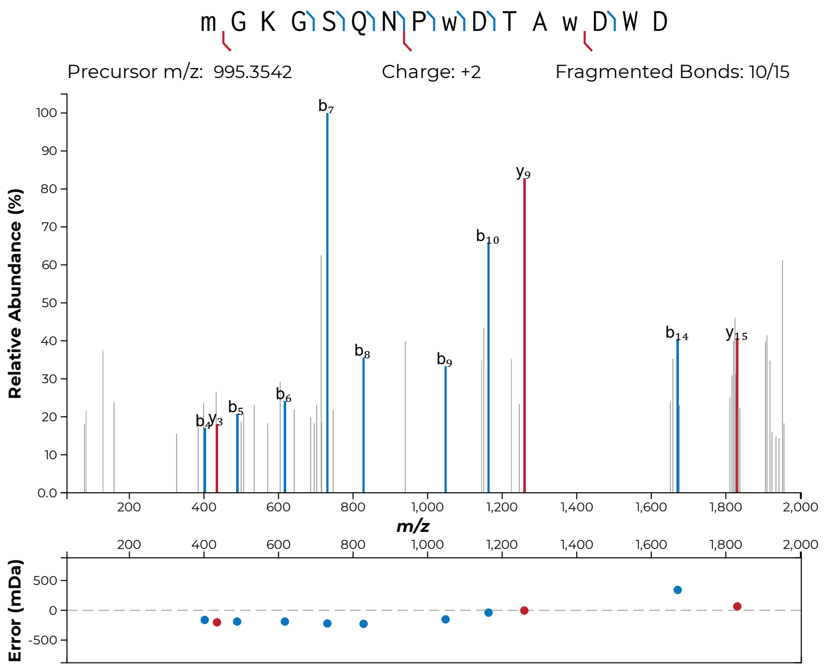
m G K G S Q N P w D T A w D W D
Precursor m/z: 995.3542
Charge: +2
Fragmented Bonds: 10/15
Relative Abundance (%)
m/z
Error (mDa)

**S47d**

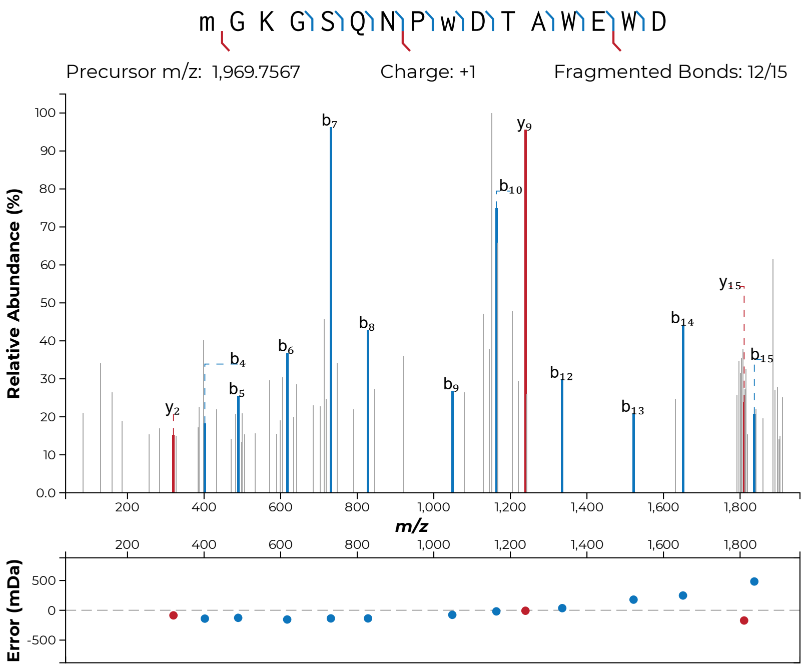

m G K G S Q N P w D T A W E W D
Precursor m/z: 1,969.7567
Charge: +1
Fragmented Bonds: 12/15
Relative Abundance (%)
m/z
Error (mDa)


**S47e**

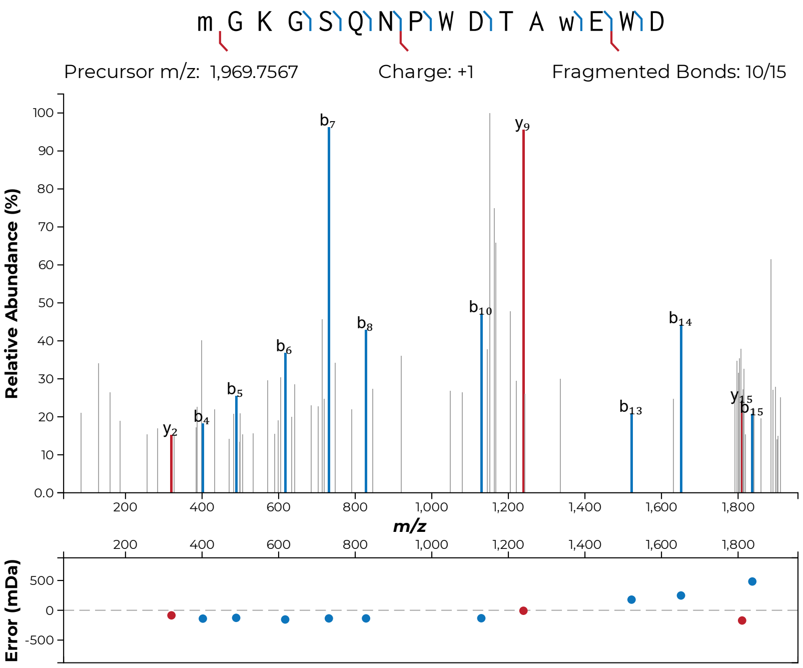

m G K G S Q N P W D T A w E W D
Precursor m/z: 1,969.7567
Charge: +1
Fragmented Bonds: 10/15
Relative Abundance (%)
m/z
Error (mDa)


**S47f**

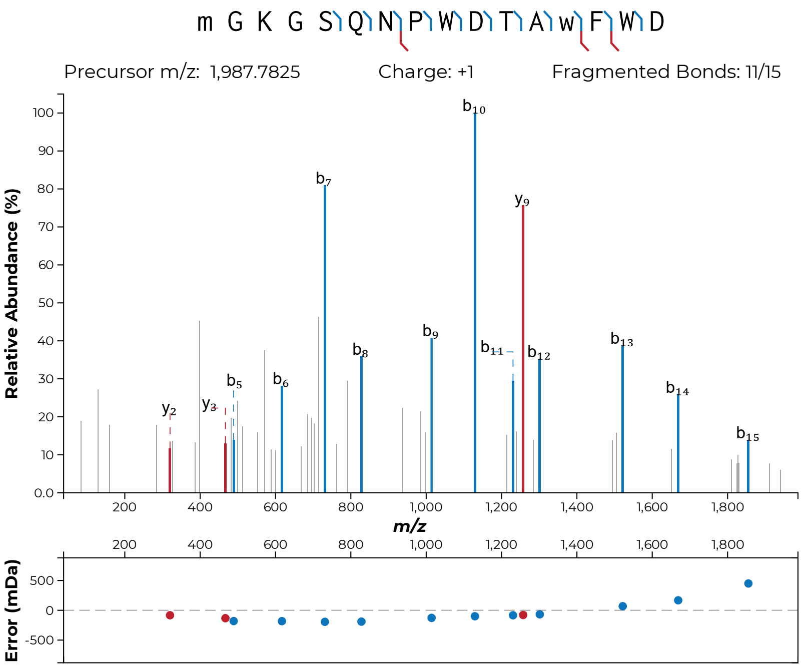
m G K G S Q N P W D T A w F W D
Precursor m/z: 1,987.7825
Charge: +1
Fragmented Bonds: 11/15
Relative Abundance (%)
m/z
Error (mDa)
b10
b7
y9
b9
b8
b11
b12
b13
b14
b15
b5
b6
y2
y3

**S47g**

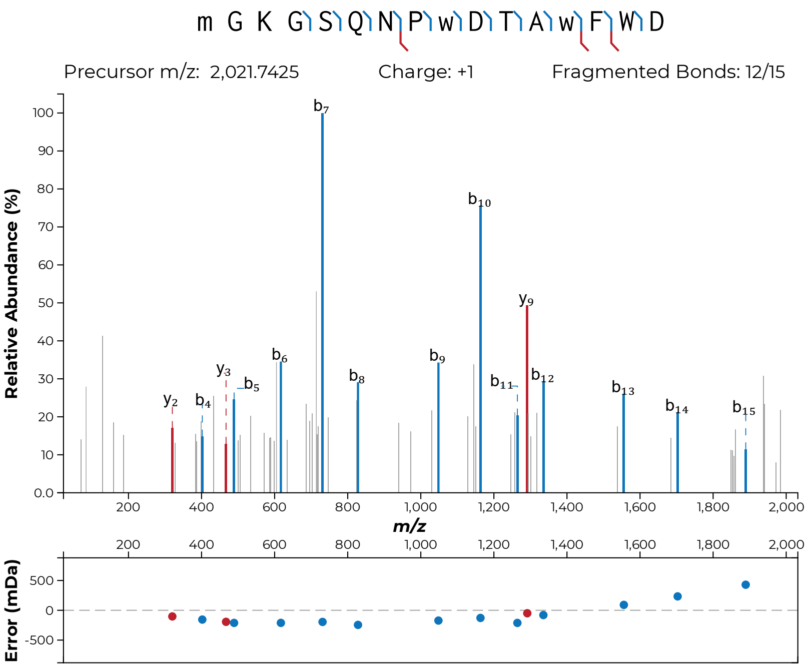
m G K G S Q N P w D T A w F W D
Precursor m/z: 2,021.7425
Charge: +1
Fragmented Bonds: 12/15
Relative Abundance (%)
m/z
Error (mDa)

**S47h**

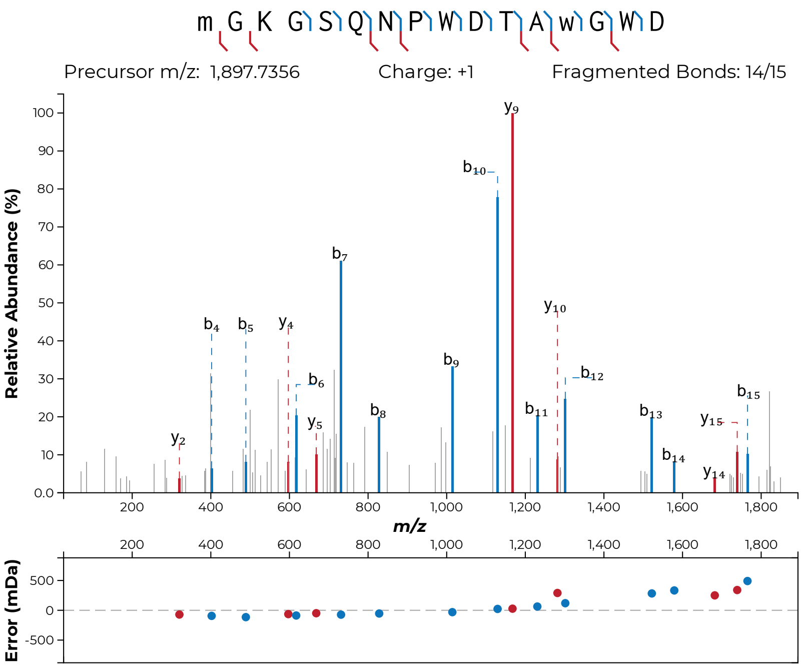

m G K G S Q N P W D T A w G W D
Precursor m/z: 1,897.7356
Charge: +1
Fragmented Bonds: 14/15
Relative Abundance (%)
m/z
Error (mDa)


**S47i**

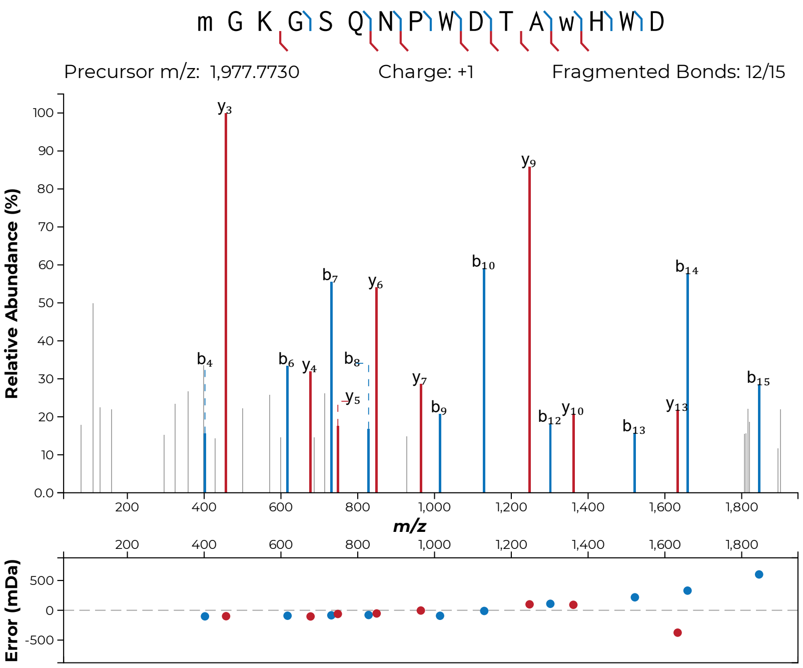

m G K G S Q N P W D T A w H W D
Precursor m/z: 1,977.7730
Charge: +1
Fragmented Bonds: 12/15
Relative Abundance (%)
m/z
Error (mDa)


**S47j**

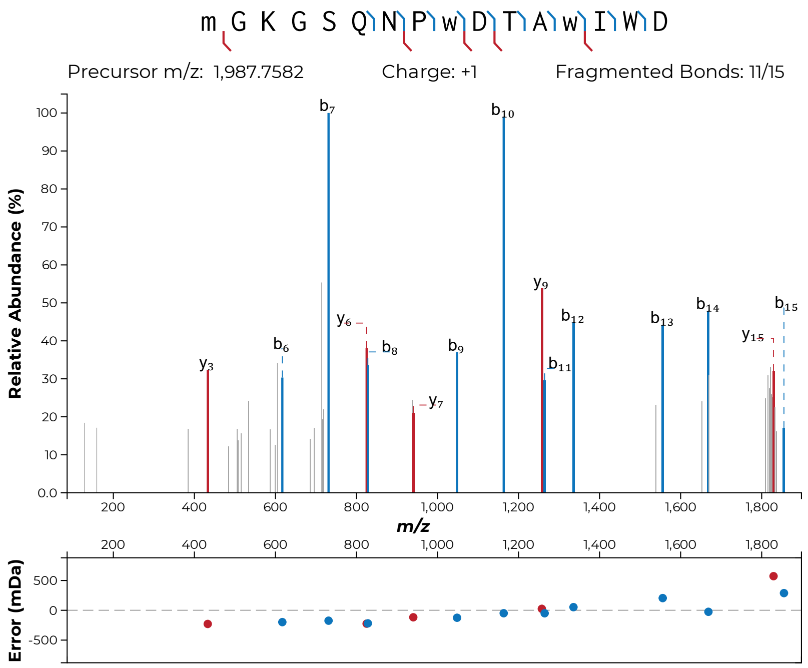
m G K G S Q N P w D T A w I W D
Precursor m/z: 1,987.7582
Charge: +1
Fragmented Bonds: 11/15
Relative Abundance (%)
m/z
Error (mDa)
b7
b10
y9
b12
b13
b14
b15
y15
y6
b8
b6
y3
b9
b11
y7

**S47k**

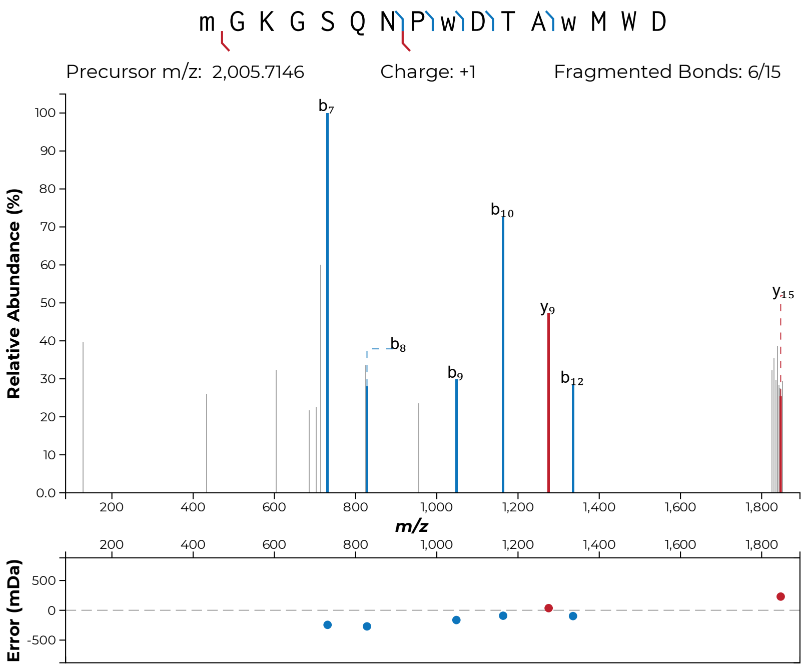

m G K G S Q N P w D T A w M W D
Precursor m/z: 2,005.7146
Charge: +1
Fragmented Bonds: 6/15
Relative Abundance (%)
m/z
Error (mDa)


**S47l**

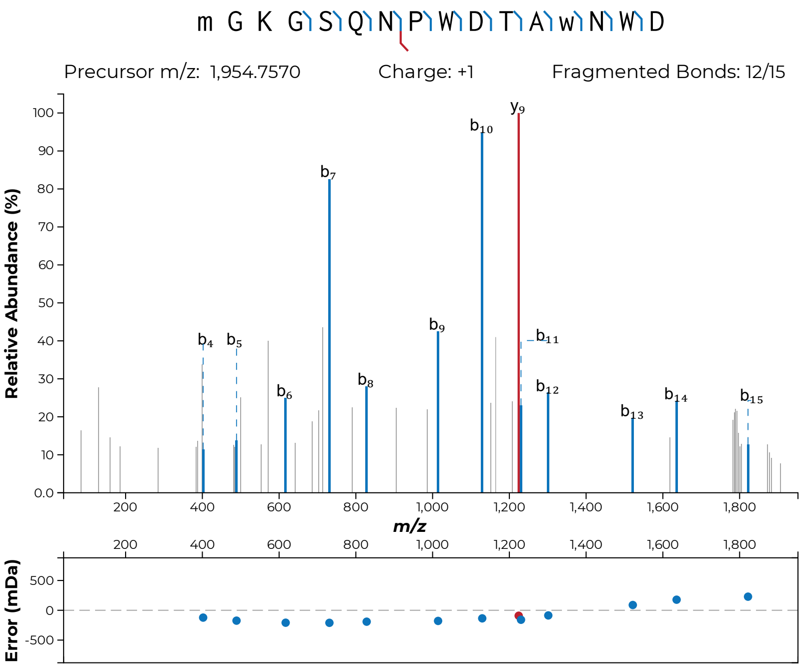

m G K G S Q N P W D T A w N W D
Precursor m/z: 1,954.7570
Charge: +1
Fragmented Bonds: 12/15
Relative Abundance (%)
m/z
Error (mDa)
b4
b5
b6
b7
b8
b9
b10
y9
b11
b12
b13
b14
b15


**S47m**

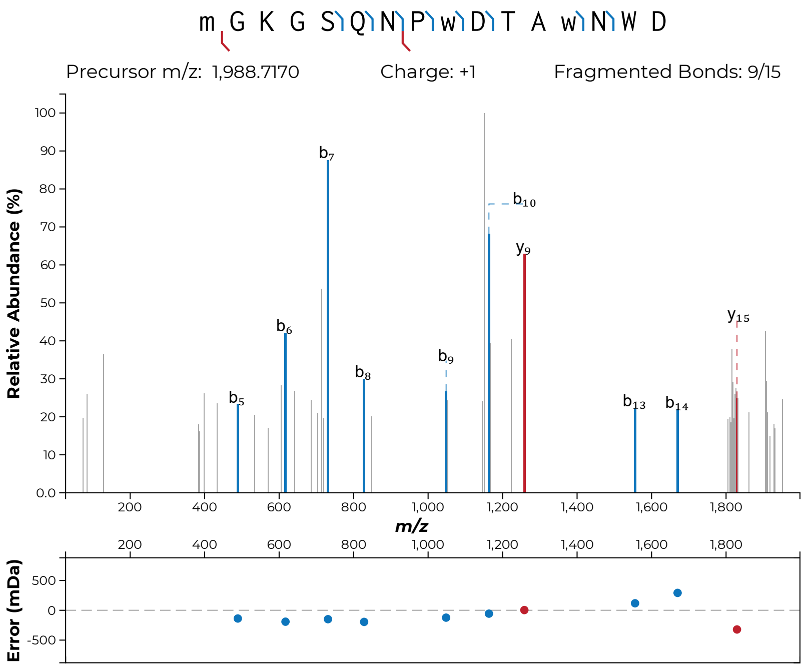

m G K G S Q N P w D T A w N W D
Precursor m/z: 1,988.7170
Charge: +1
Fragmented Bonds: 9/15
Relative Abundance (%)
m/z
Error (mDa)


**S47n**

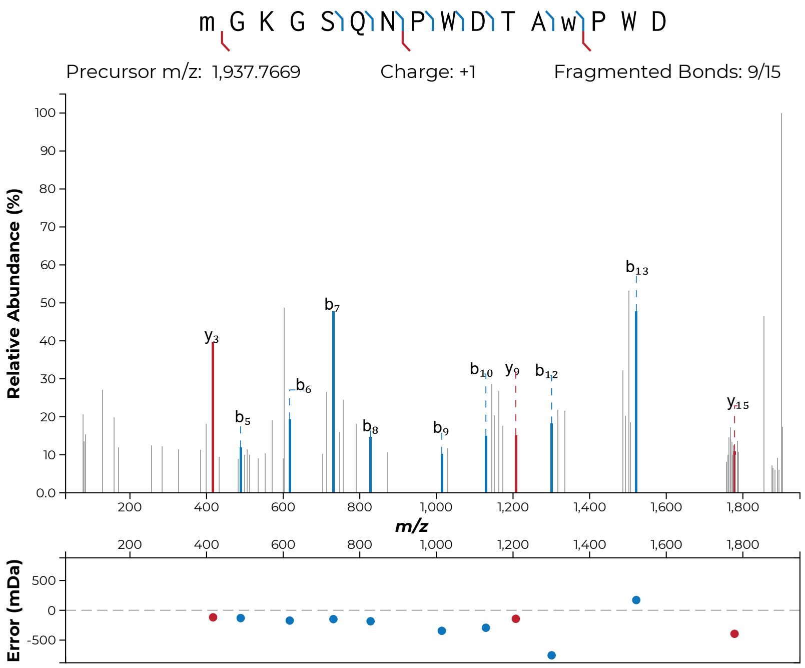

m G K G S Q N P W D T A w P W D
Precursor m/z: 1,937.7669
Charge: +1
Fragmented Bonds: 9/15
Relative Abundance (%)
m/z
Error (mDa)


**S47o**

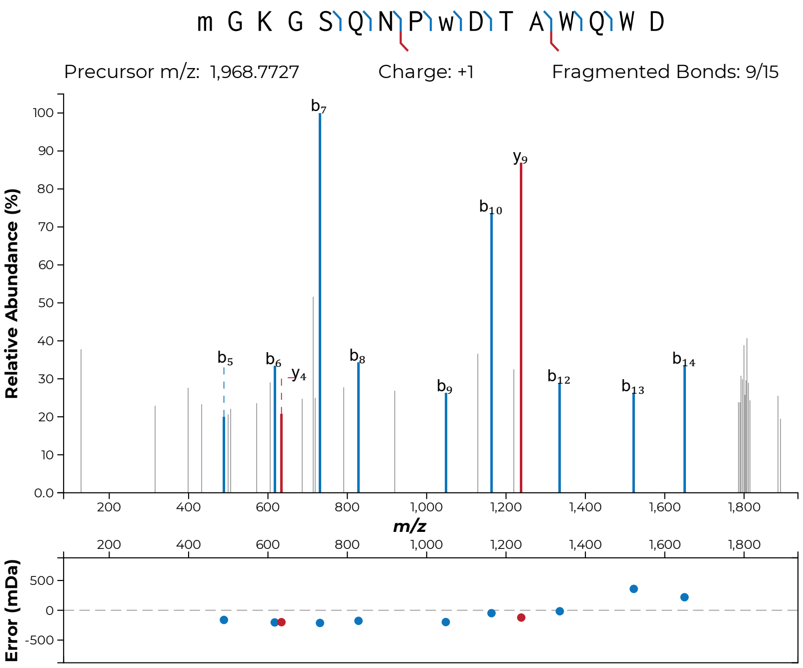

m G K G S Q N P w D T A W Q W D
Precursor m/z: 1,968.7727
Charge: +1
Fragmented Bonds: 9/15
Relative Abundance (%)
m/z
Error (mDa)


**S47p**

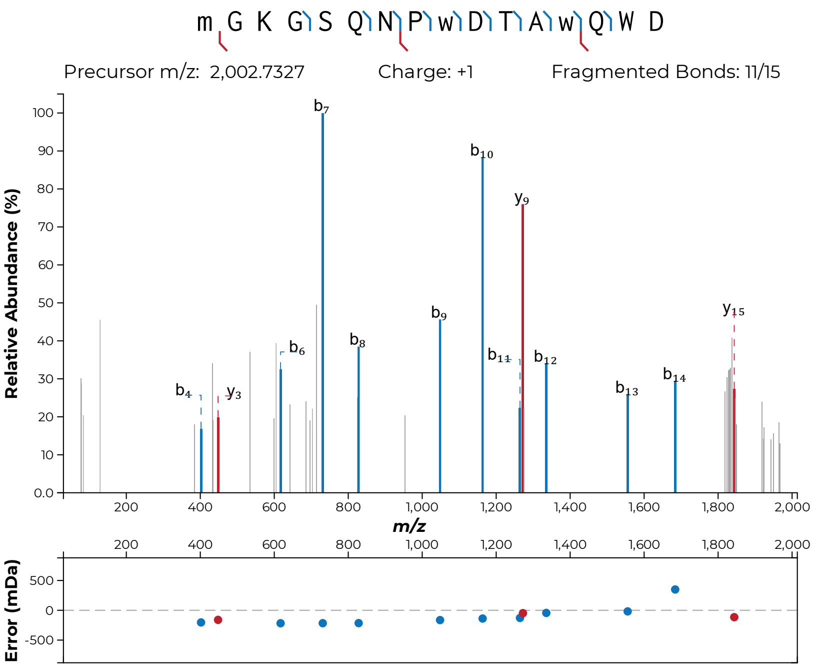

m G K G S Q N P w D T A w Q W D
Precursor m/z: 2,002.7327
Charge: +1
Fragmented Bonds: 11/15
Relative Abundance (%)
m/z
Error (mDa)
b4
y3
b6
b7
b8
b9
b10
y9
b11
b12
b13
b14
y15


**S47q**

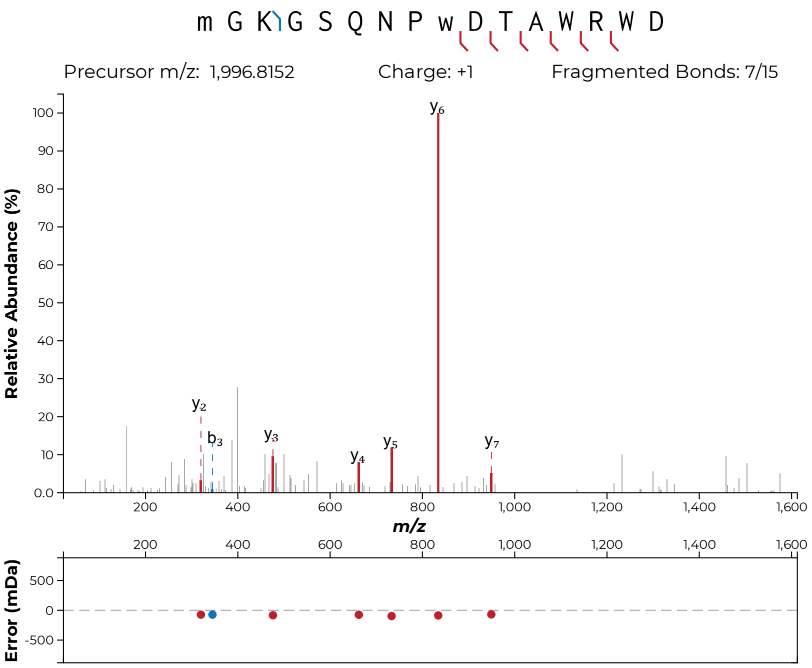
m G K G S Q N P w D T A W R W D
Precursor m/z: 1,996.8152
Charge: +1
Fragmented Bonds: 7/15
Relative Abundance (%)
y6
y2
b3
y3
y4
y5
y7
m/z
Error (mDa)

**S47r**

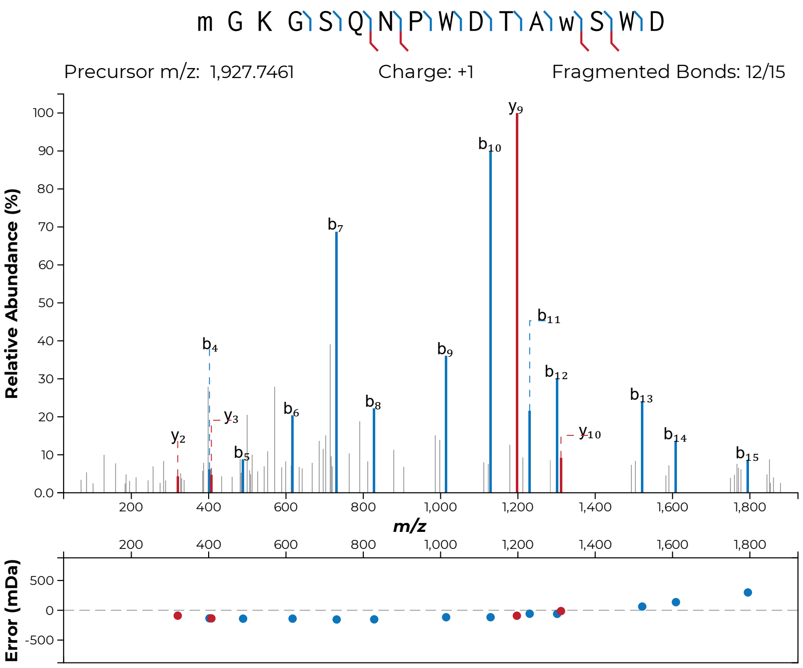
m G K G S Q N P W D T A w S W D
Precursor m/z: 1,927.7461
Charge: +1
Fragmented Bonds: 12/15
Relative Abundance (%)
m/z
Error (mDa)

**S47s**

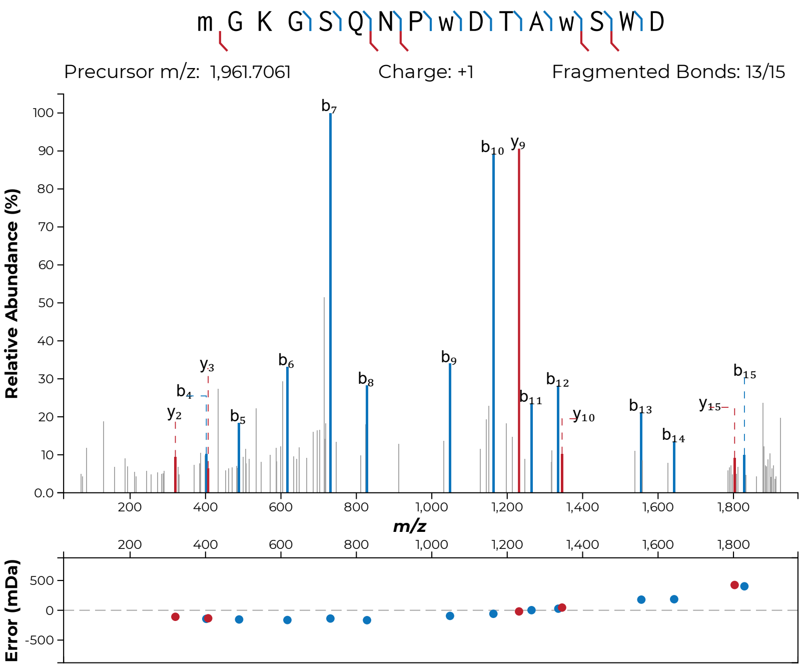

m G K G S Q N P w D T A w S W D
Precursor m/z: 1,961.7061
Charge: +1
Fragmented Bonds: 13/15
Relative Abundance (%)
m/z
Error (mDa)


**S47t**

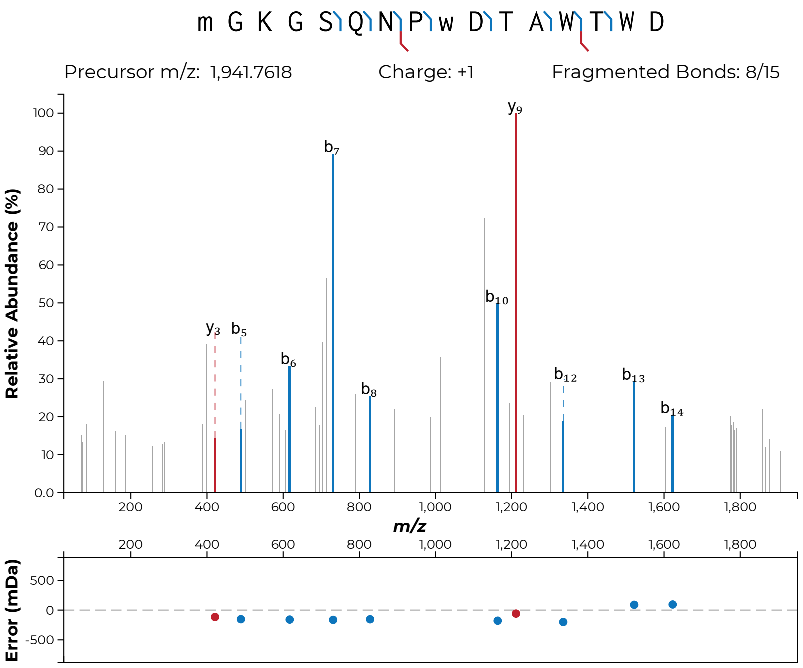
m G K G S Q N P w D T A W T W D
Precursor m/z: 1,941.7618
Charge: +1
Fragmented Bonds: 8/15
Relative Abundance (%)
m/z
Error (mDa)
y3
b5
b6
b7
b8
b10
y9
b12
b13
b14

**S47u**

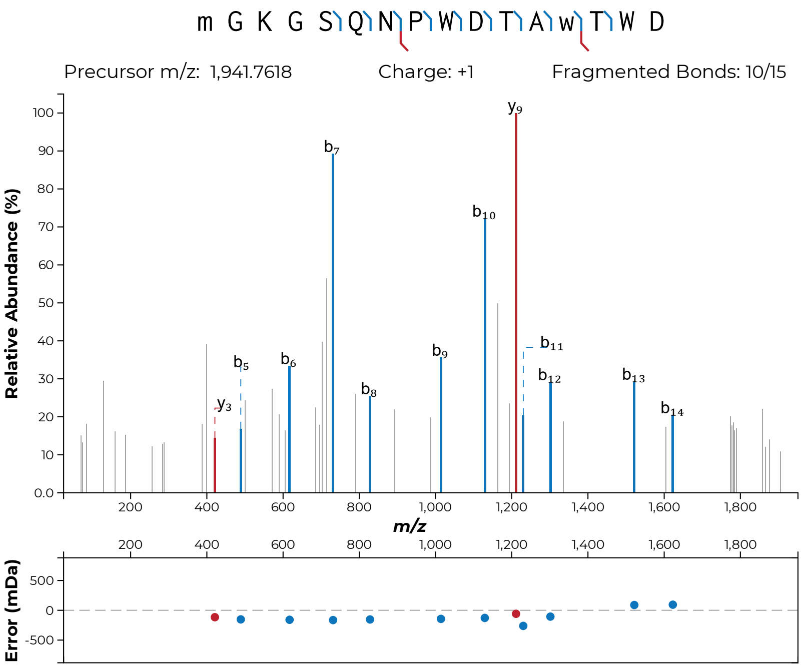

m G K G S Q N P W D T A w T W D
Precursor m/z: 1,941.7618
Charge: +1
Fragmented Bonds: 10/15
Relative Abundance (%)
m/z
Error (mDa)


**S47v**

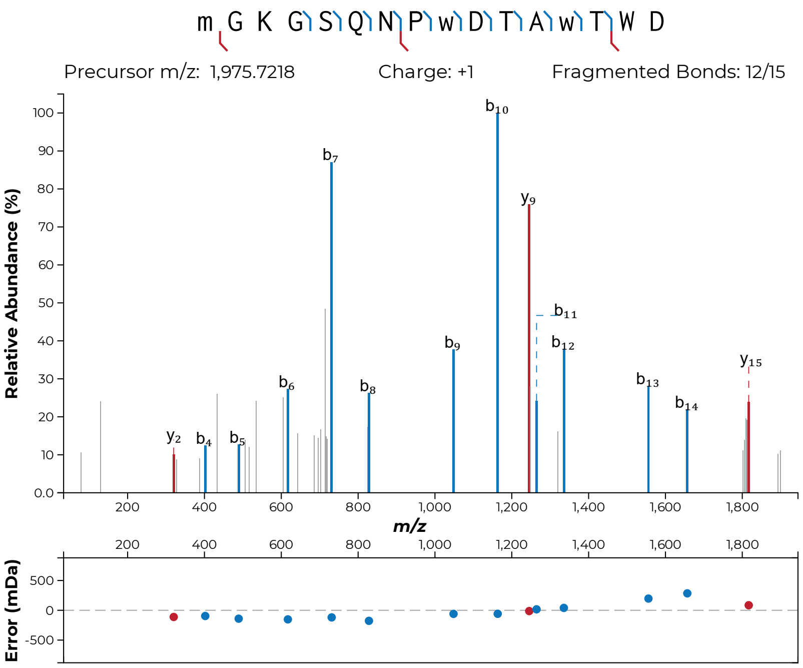
m G K G S Q N P w D T A w T W D
Precursor m/z: 1,975.7218
Charge: +1
Fragmented Bonds: 12/15
Relative Abundance (%)
m/z
Error (mDa)

**S47w**

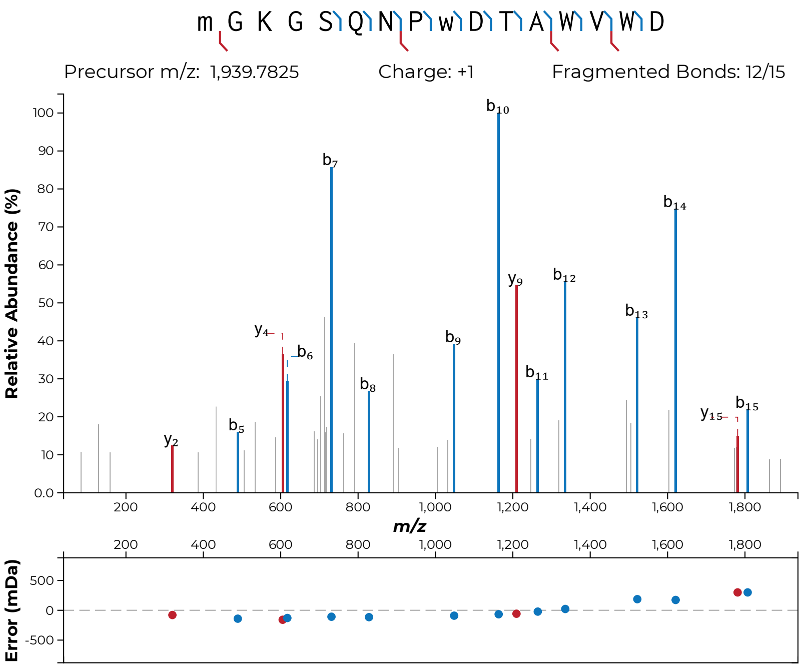

m G K G S Q N P w D T A W V W D
Precursor m/z: 1,939.7825
Charge: +1
Fragmented Bonds: 12/15
Relative Abundance (%)
m/z
Error (mDa)


**S47x**

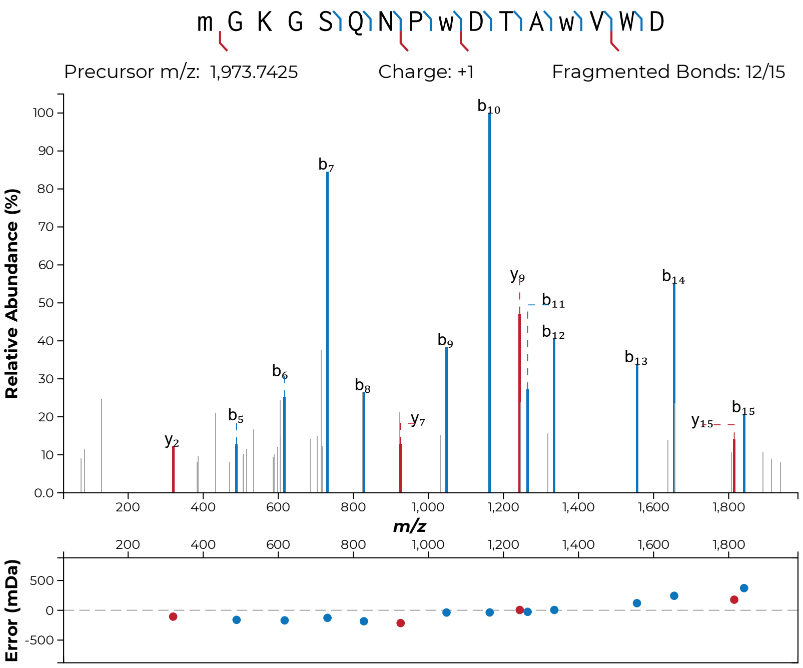
m G K G S Q N P w D T A w V W D
Precursor m/z: 1,973.7425
Charge: +1
Fragmented Bonds: 12/15
Relative Abundance (%)
m/z
Error (mDa)

**S47y**

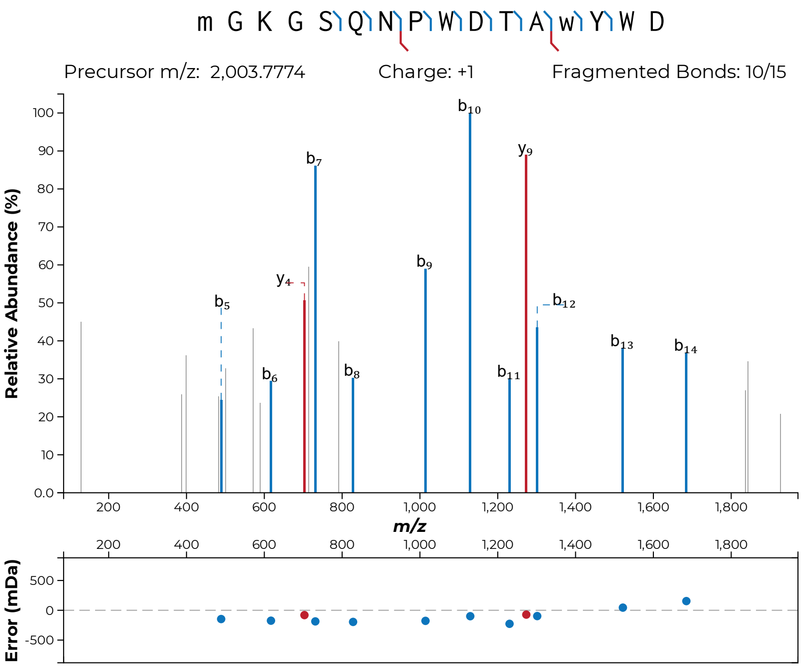
m G K G S Q N P W D T A w Y W D
Precursor m/z: 2,003.7774
Charge: +1
Fragmented Bonds: 10/15
Relative Abundance (%)
m/z
Error (mDa)
b5
b6
y4
b7
b8
b9
b10
b11
y9
b12
b13
b14

**S47z**

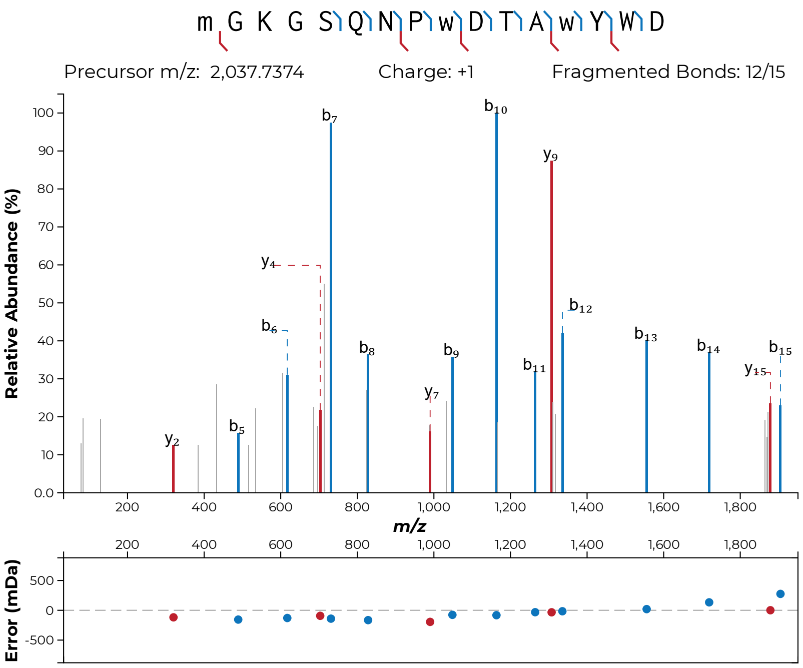

m G K G S Q N P w D T A w Y W D
Precursor m/z: 2,037.7374
Charge: +1
Fragmented Bonds: 12/15
Relative Abundance (%)
m/z
Error (mDa)


**S47aa**

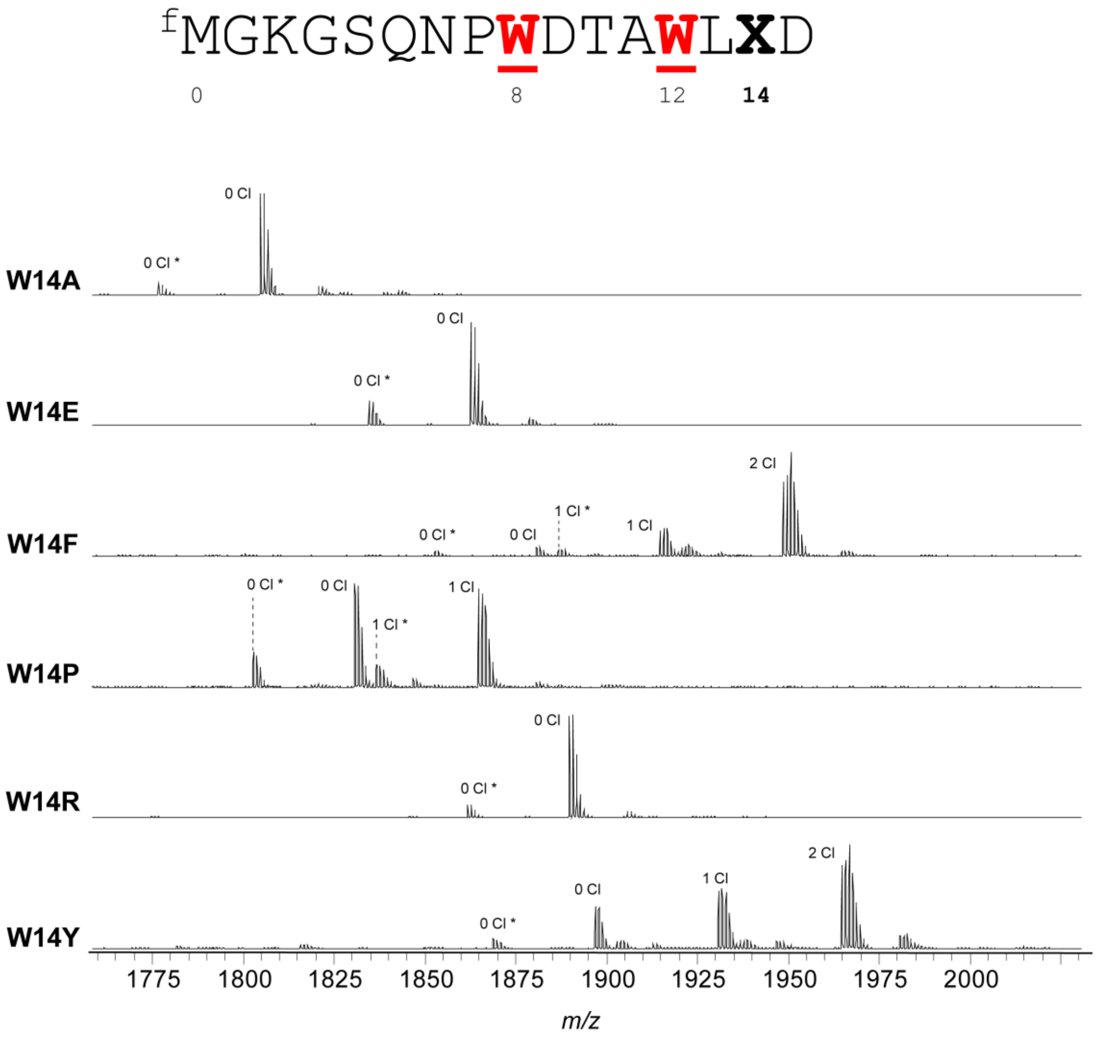


**Figure S48: MALDI-TOF-MS and -LIFT-MS for ChlA$_{PE}$ Trp14 variants.** (**a**) MALDI-TOF-MS spectra for all variants made of position 14 of the ChlA$_{PE}$ substrate. Asterisks (*) indicate peptides lacking N-terminal formylation.

(**b-g, following pages**): MS/MS spectra acquired for this set of substrates. In all MS/MS spectra, lowercase letters indicate modified residues. For example, the N-terminal formylated Met is indicated as “m”. Chlorinated Trp is indicated as “w”.

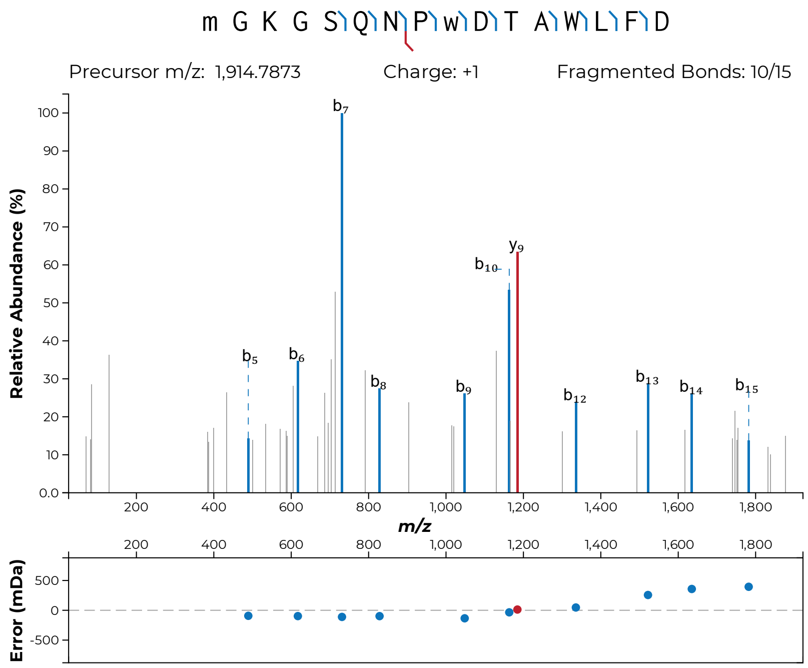

m G K G S Q N P w D T A W L F D
Precursor m/z: 1,914.7873
Charge: +1
Fragmented Bonds: 10/15
Relative Abundance (%)
m/z
Error (mDa)
b5
b6
b7
b8
b9
b10
y9
b12
b13
b14
b15


**S48b**

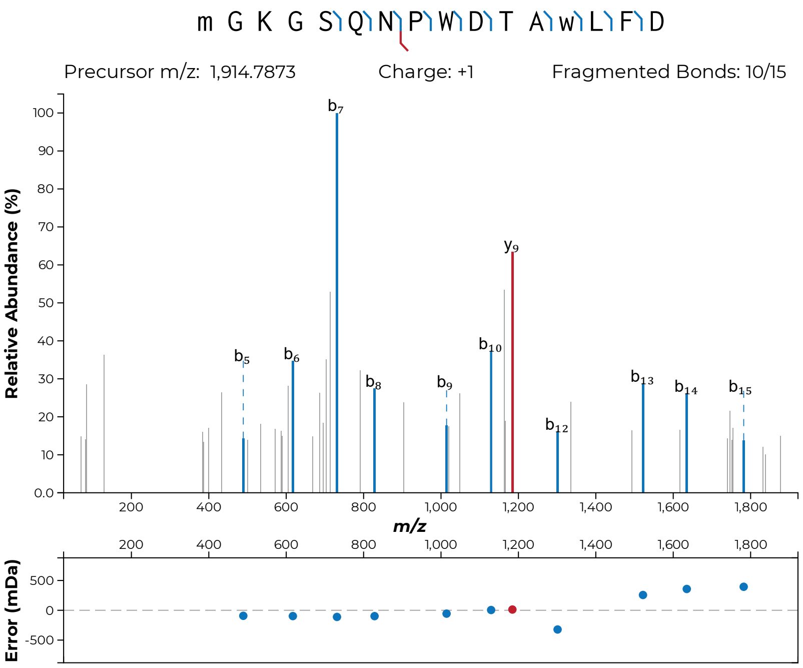
m G K G S Q N P W D T A w L F D
Precursor m/z: 1,914.7873
Charge: +1
Fragmented Bonds: 10/15
Relative Abundance (%)
m/z
Error (mDa)

**S48c**

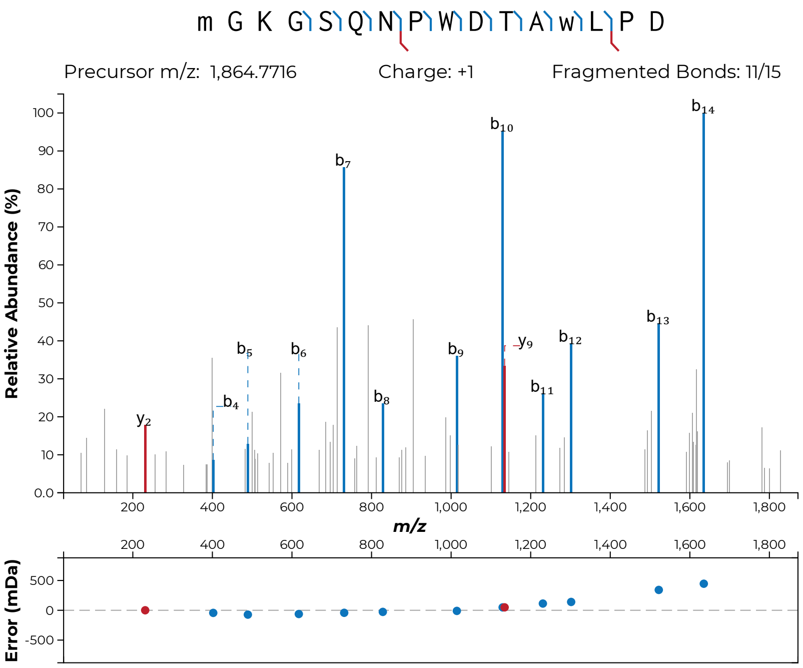
m G K G S Q N P W D T A w L P D
Precursor m/z: 1,864.7716
Charge: +1
Fragmented Bonds: 11/15
Relative Abundance (%)
m/z
Error (mDa)

**S48d**

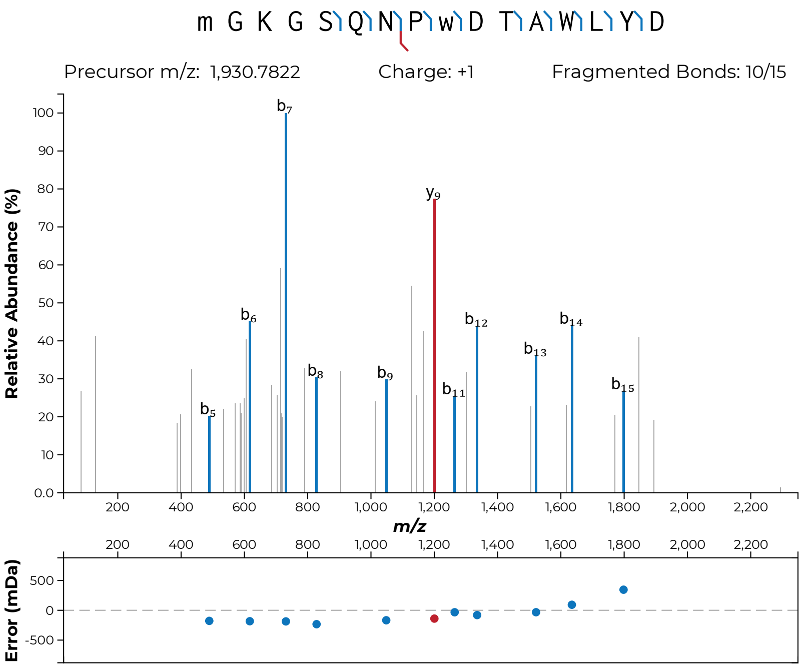
m G K G S Q N P w D T A W L Y D
Precursor m/z: 1,930.7822
Charge: +1
Fragmented Bonds: 10/15
Relative Abundance (%)
b7
y9
b6
b12
b14
b13
b8
b9
b11
b15
b5
m/z
Error (mDa)

**S48e**

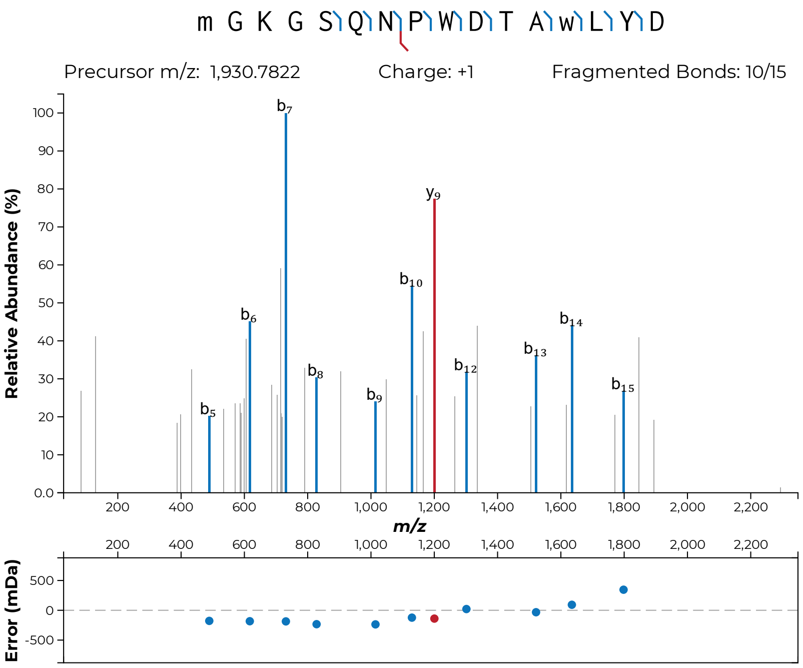
m G K G S Q N P W D T A w L Y D
Precursor m/z: 1,930.7822
Charge: +1
Fragmented Bonds: 10/15
Relative Abundance (%)
b7
y9
b10
b6
b14
b13
b8
b12
b15
b9
b5
m/z
Error (mDa)

**S48f**

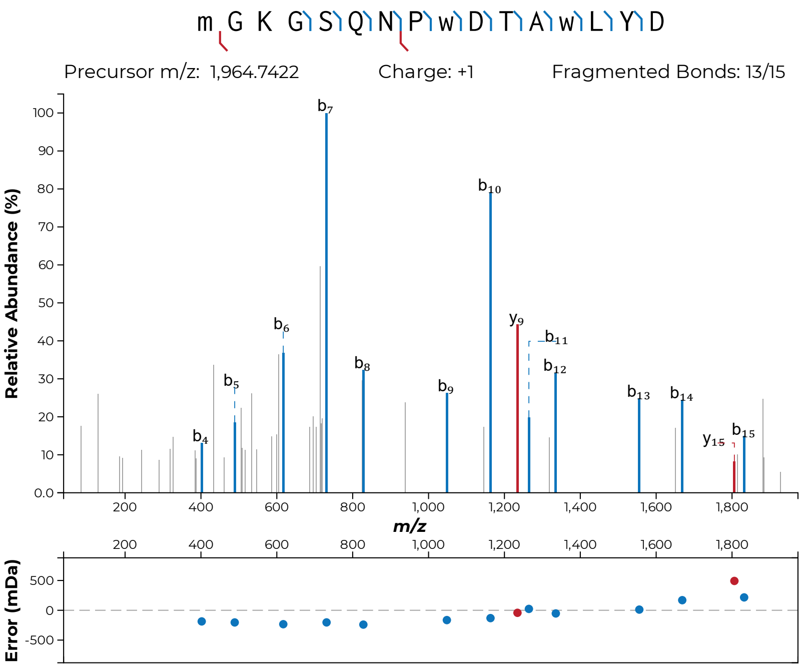

m G K G S Q N P w D T A w L Y D
Precursor m/z: 1,964.7422
Charge: +1
Fragmented Bonds: 13/15
Relative Abundance (%)
m/z
Error (mDa)


**S48g**

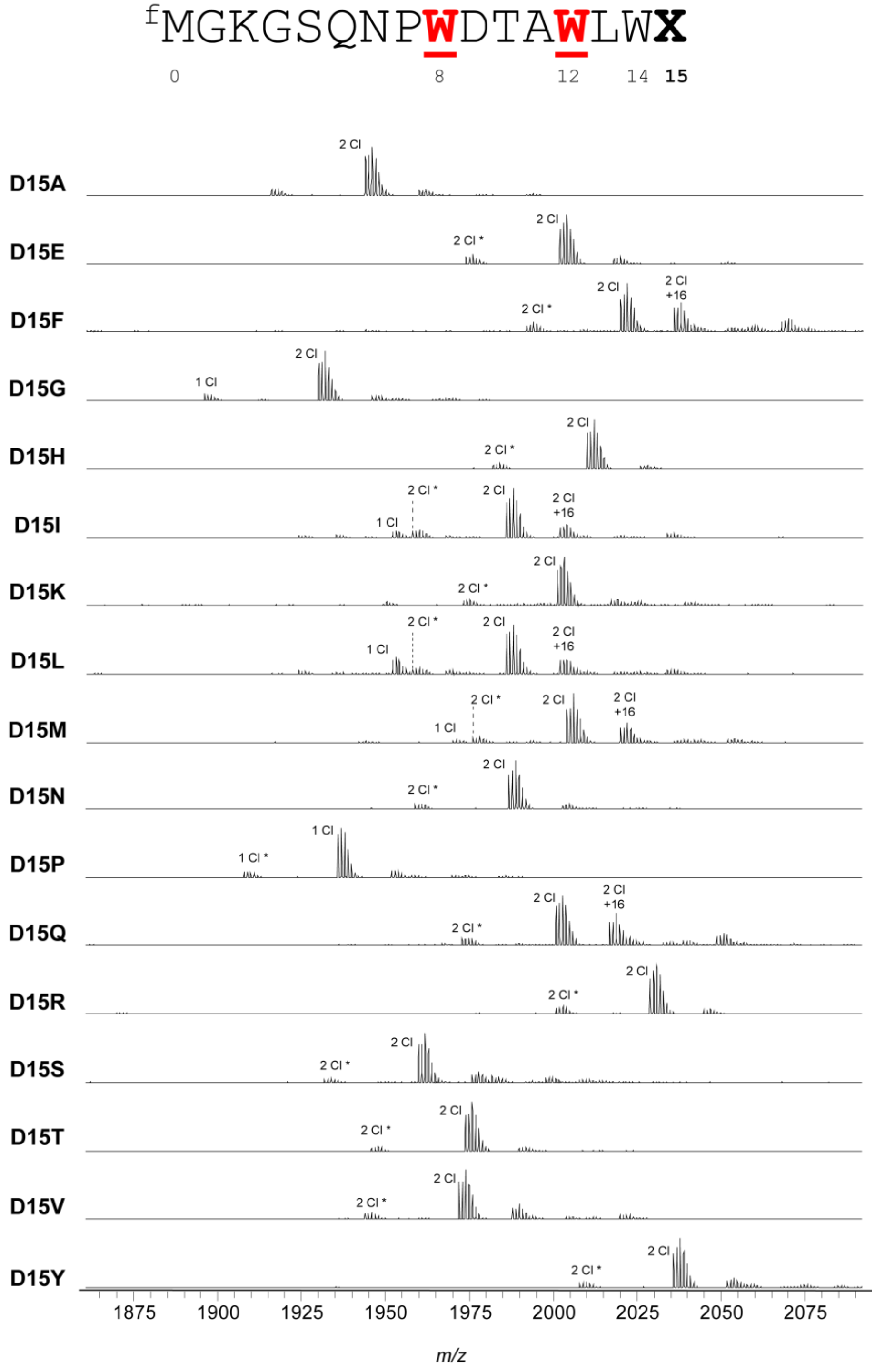


**Figure S49: MALDI-TOF-MS and -LIFT-MS for ChlA$_{PE}$ Asp15 variants.** (**a**) MALDI-TOF-MS spectra for all variants made of position 15 of the ChlA$_{PE}$ substrate. Asterisks (*) indicate peptides lacking N-terminal formylation.

(**b-s, following pages**): MS/MS spectra acquired for this set of substrates. In all MS/MS spectra, lowercase letters indicate modified residues. For example, the N-terminal formylated Met is indicated as “m”. Chlorinated Trp is indicated as “w”.

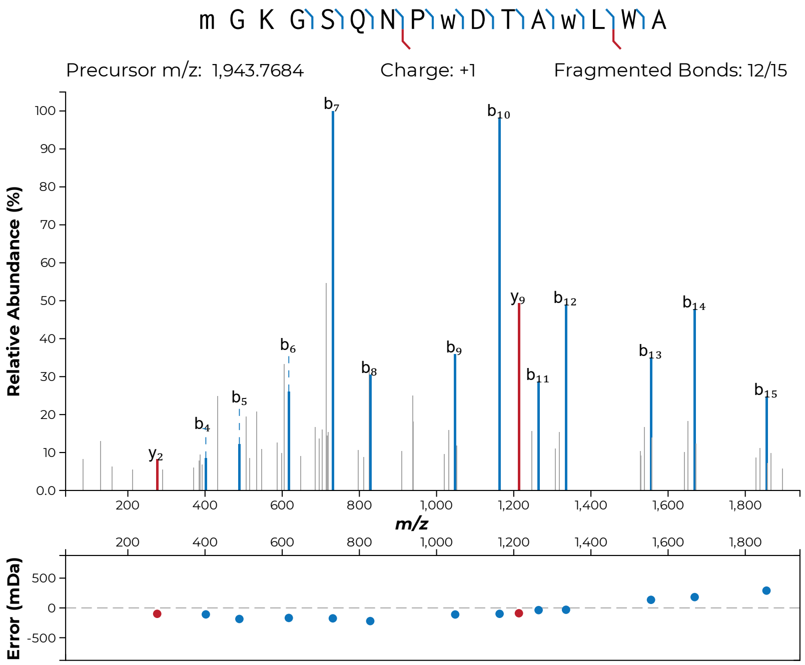
m G K G S Q N P w D T A w L W A
Precursor m/z: 1,943.7684
Charge: +1
Fragmented Bonds: 12/15
Relative Abundance (%)
m/z
Error (mDa)

**S49b**

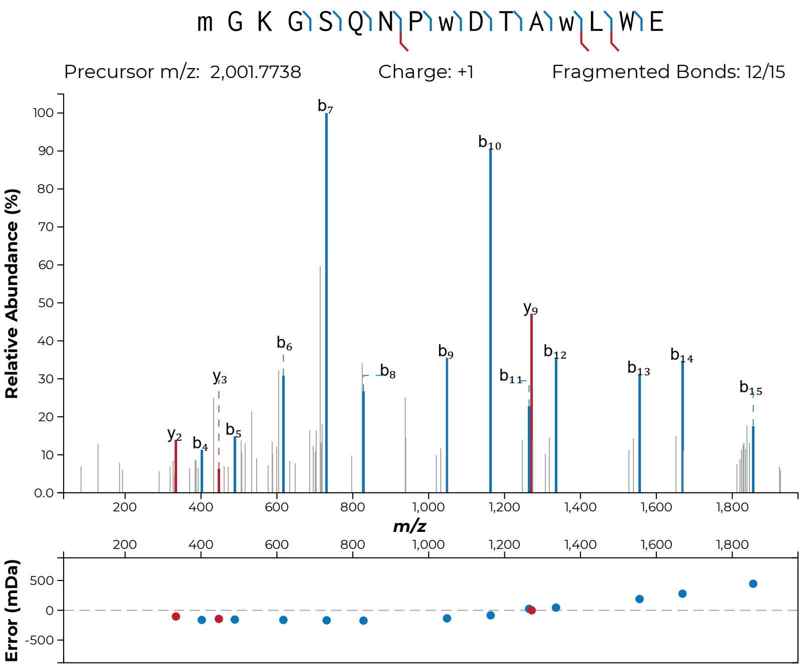
m G K G S Q N P w D T A w L W E
Precursor m/z: 2,001.7738
Charge: +1
Fragmented Bonds: 12/15
Relative Abundance (%)
m/z
Error (mDa)

**S49c**

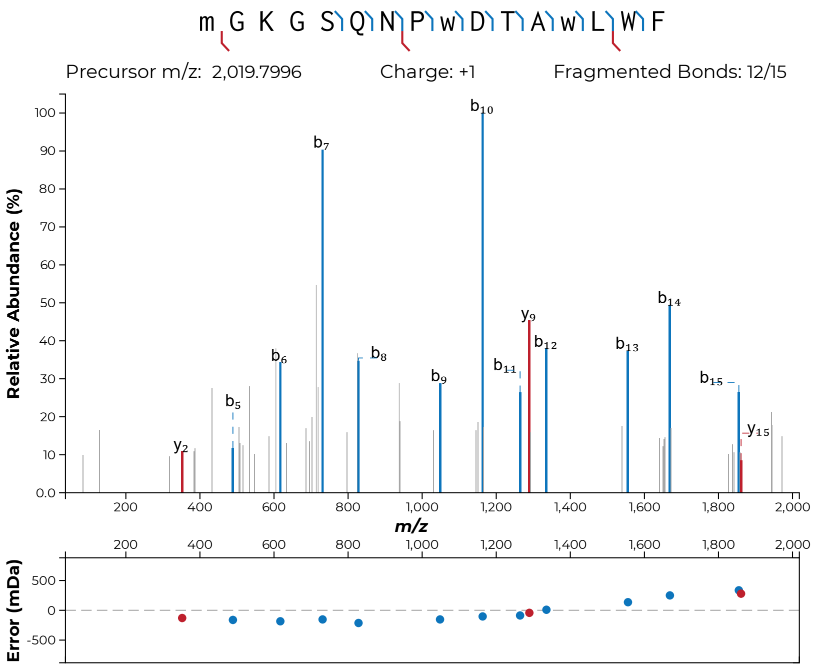
m G K G S Q N P w D T A w L W F
Precursor m/z: 2,019.7996
Charge: +1
Fragmented Bonds: 12/15
Relative Abundance (%)
m/z
Error (mDa)
y2
b5
b6
b7
b8
b9
b10
b11
y9
b12
b13
b14
b15
y15

**S49d**

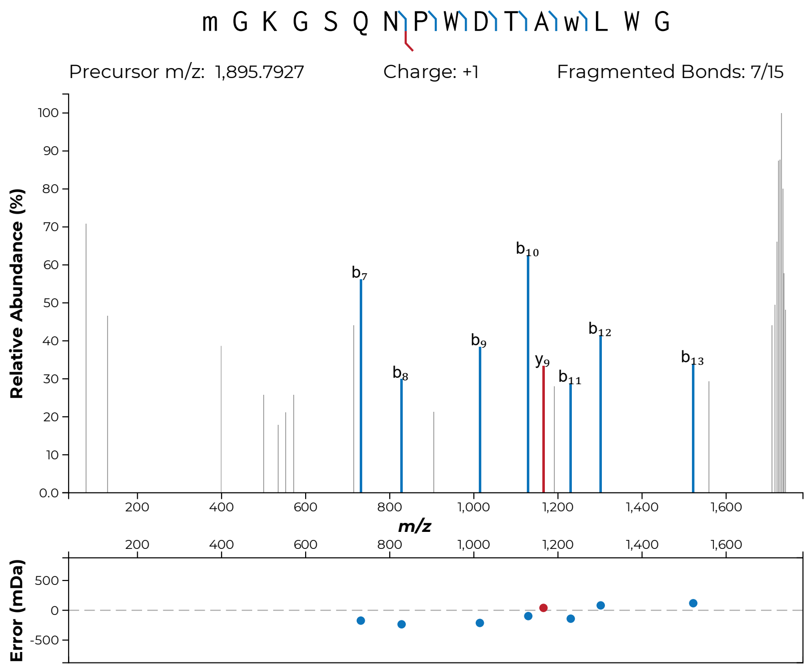

m G K G S Q N P W D T A w L W G
Precursor m/z: 1,895.7927
Charge: +1
Fragmented Bonds: 7/15
Relative Abundance (%)
m/z
Error (mDa)
b7
b8
b9
b10
y9
b11
b12
b13


**S49e**

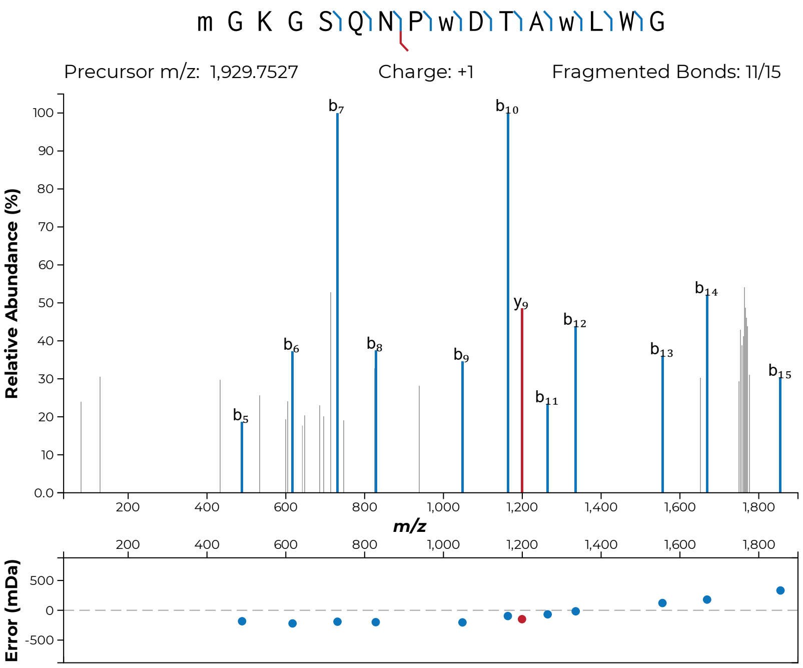
m G K G S Q N P w D T A w L W G
Precursor m/z: 1,929.7527
Charge: +1
Fragmented Bonds: 11/15
Relative Abundance (%)
m/z
Error (mDa)

**S49f**

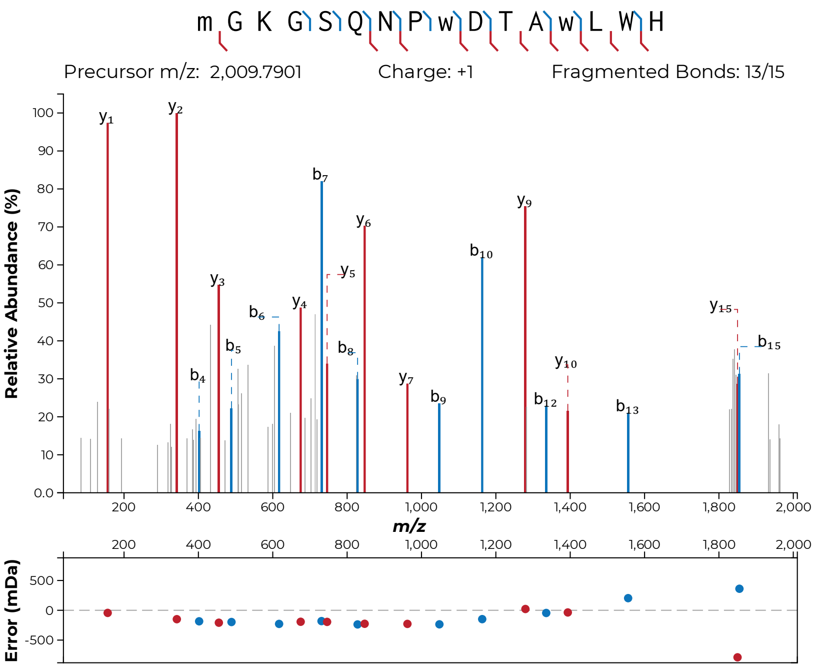
m G K G S Q N P w D T A w L W H
Precursor m/z: 2,009.7901
Charge: +1
Fragmented Bonds: 13/15
Relative Abundance (%)
m/z
Error (mDa)

**S49g**

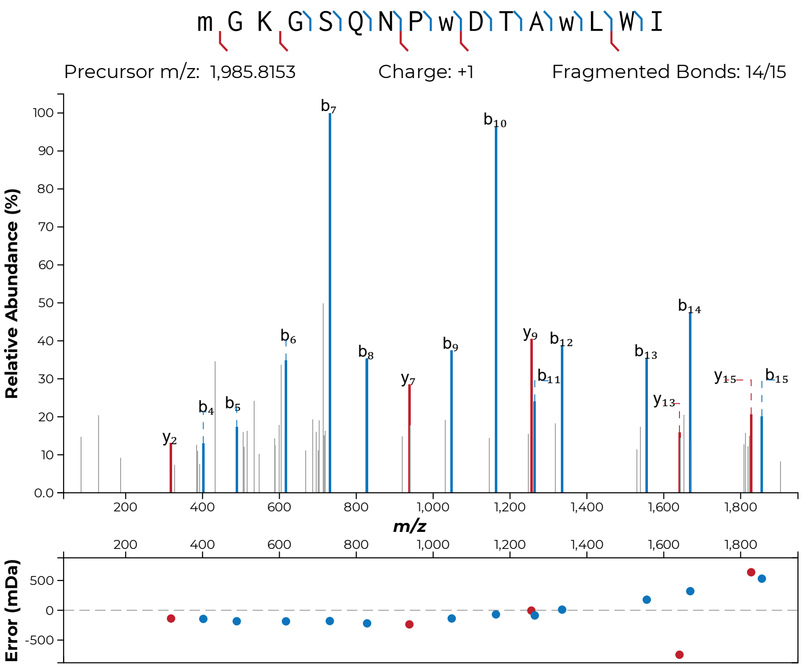
m G K G S Q N P w D T A w L W I
Precursor m/z: 1,985.8153
Charge: +1
Fragmented Bonds: 14/15
Relative Abundance (%)
m/z
Error (mDa)

**S49h**

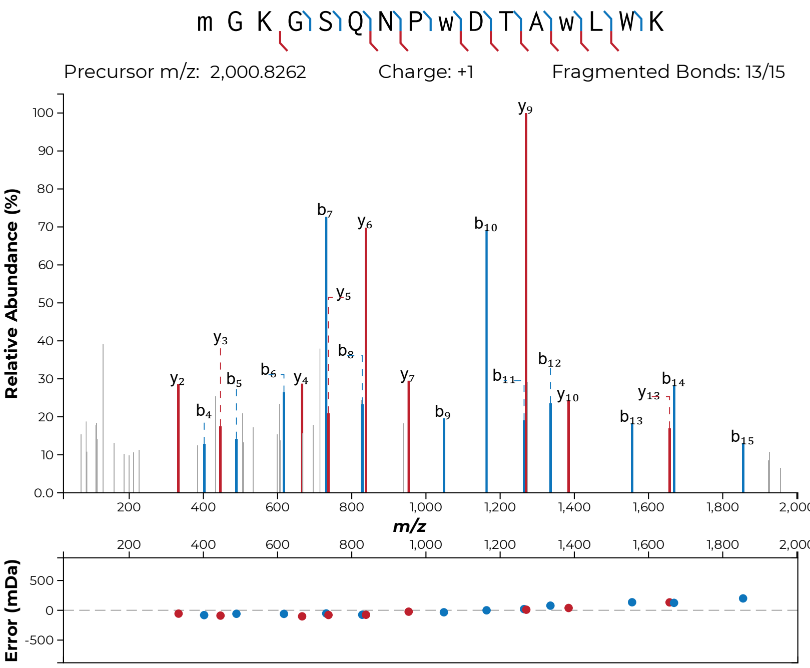

m G K G S Q N P w D T A w L W K
Precursor m/z: 2,000.8262
Charge: +1
Fragmented Bonds: 13/15
Relative Abundance (%)
m/z
Error (mDa)


**S49i**

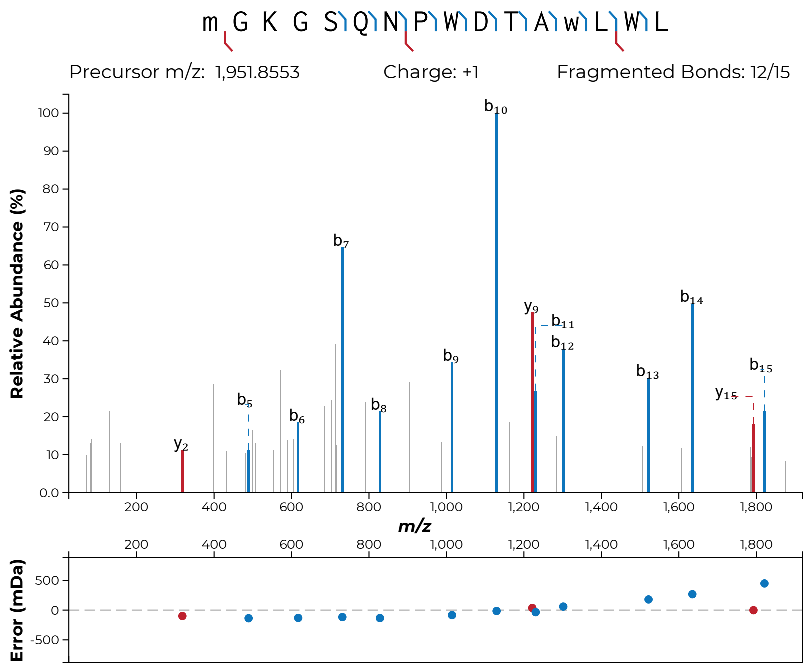

m G K G S Q N P W D T A w L W L
Precursor m/z: 1,951.8553
Charge: +1
Fragmented Bonds: 12/15
Relative Abundance (%)
m/z
Error (mDa)
b10
b7
y9
b11
b12
b14
b9
b13
b15
y15
b5
b6
b8
y2


**S49j**

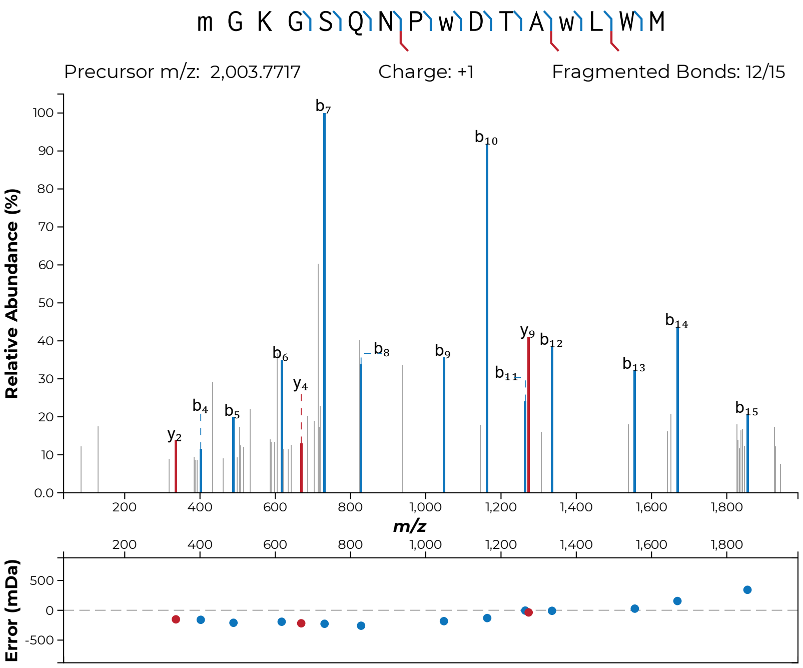
m G K G S Q N P w D T A w L W M
Precursor m/z: 2,003.7717
Charge: +1
Fragmented Bonds: 12/15
Relative Abundance (%)
m/z
Error (mDa)

**S49k**

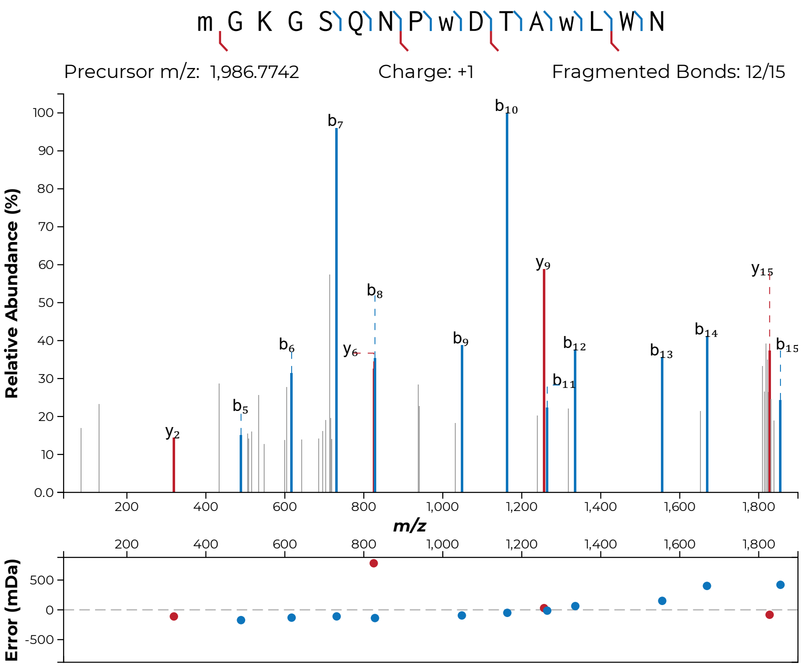
m G K G S Q N P w D T A w L W N
Precursor m/z: 1,986.7742
Charge: +1
Fragmented Bonds: 12/15
Relative Abundance (%)
m/z
Error (mDa)
b7
b10
y9
y15
b8
b6
y6
b9
b12
b14
b13
b15
b11
b5
y2

**S49l**

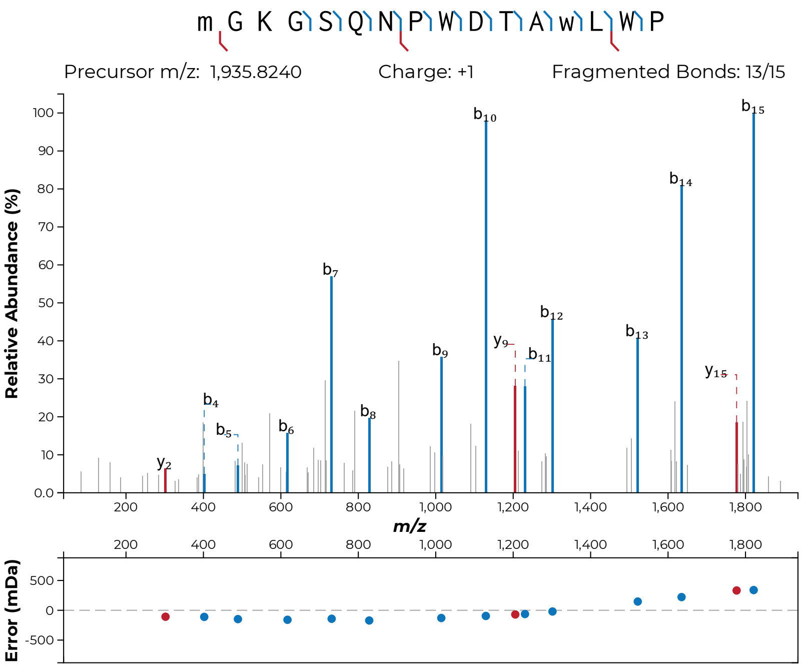

m G K G S Q N P W D T A w L W P
Precursor m/z: 1,935.8240
Charge: +1
Fragmented Bonds: 13/15
Relative Abundance (%)
m/z
Error (mDa)


**S49m**

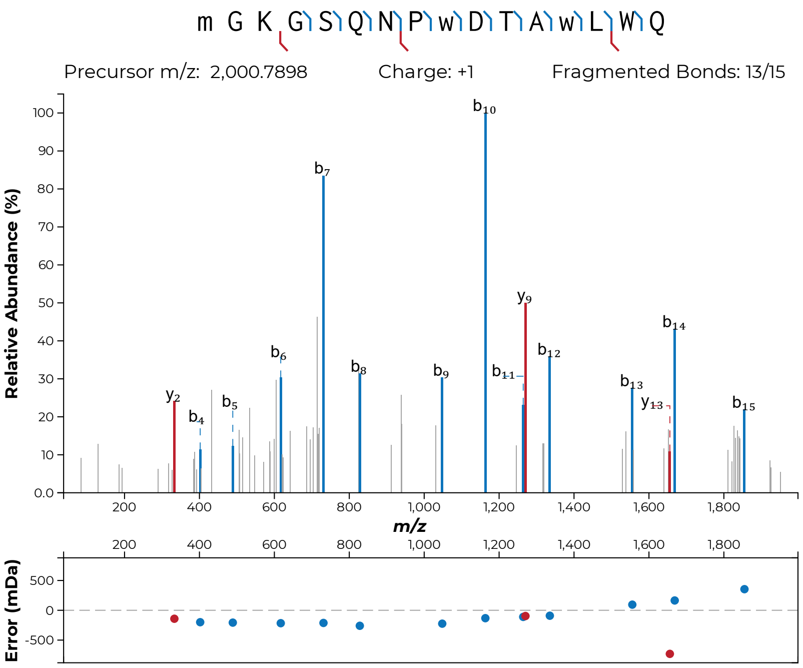
m G K G S Q N P w D T A w L W Q
Precursor m/z: 2,000.7898
Charge: +1
Fragmented Bonds: 13/15
Relative Abundance (%)
m/z
Error (mDa)
y2
b4
b5
b6
b7
b8
b9
b10
b11
y9
b12
b13
y13
b14
b15

**S49n**

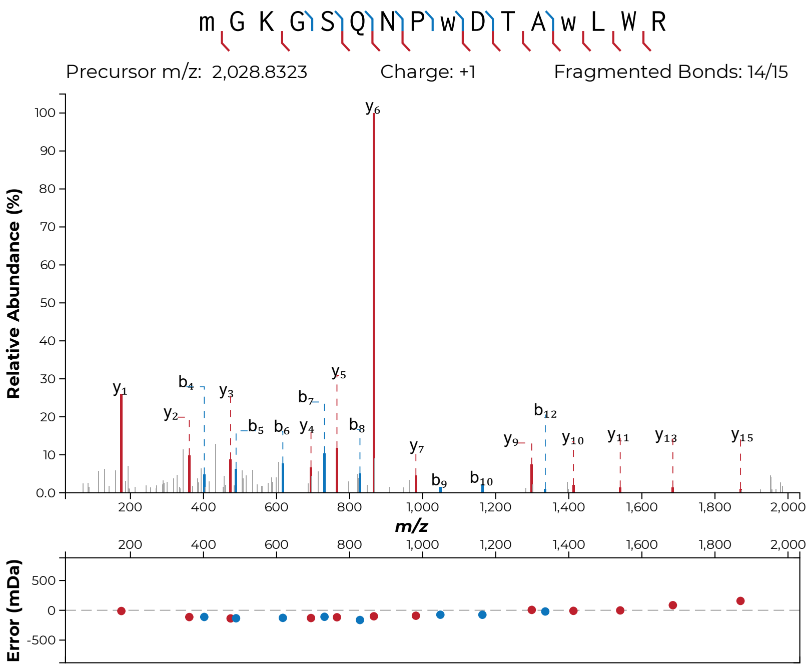
m G K G S Q N P w D T A w L W R
Precursor m/z: 2,028.8323
Charge: +1
Fragmented Bonds: 14/15
Relative Abundance (%)
m/z
Error (mDa)

**S49o**

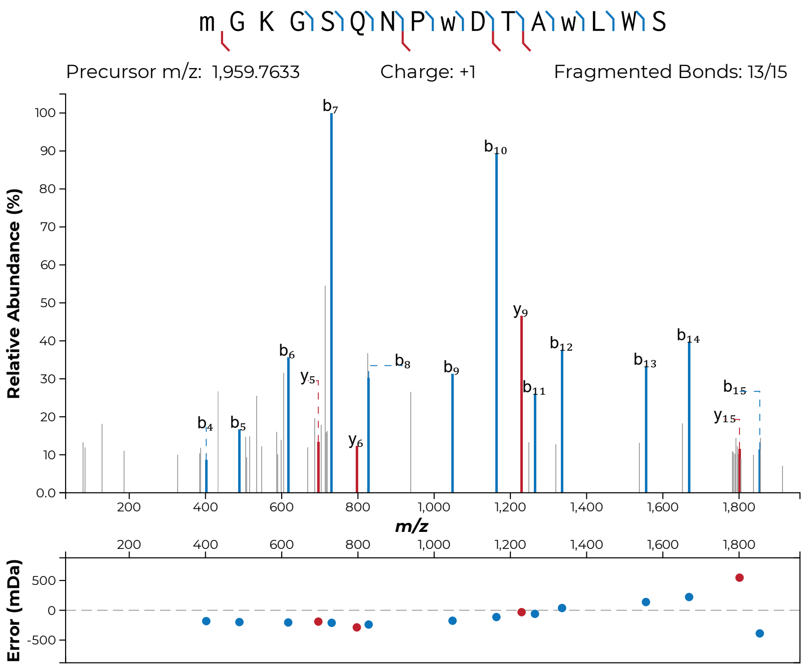
m G K G S Q N P w D T A w L W S
Precursor m/z: 1,959.7633
Charge: +1
Fragmented Bonds: 13/15
Relative Abundance (%)
m/z
Error (mDa)

**S49p**

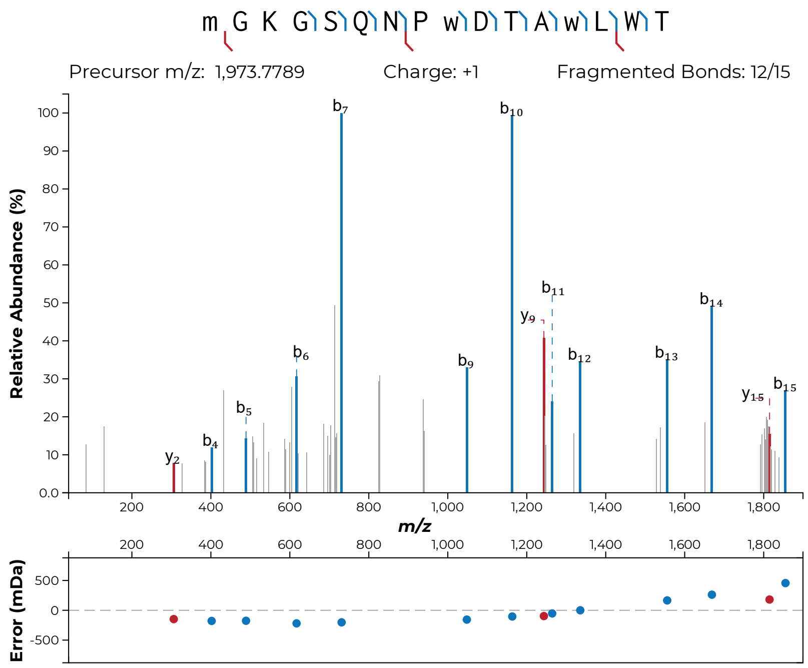
m G K G S Q N P w D T A w L W T
Precursor m/z: 1,973.7789
Charge: +1
Fragmented Bonds: 12/15
Relative Abundance (%)
m/z
Error (mDa)
b7
b10
b11
y9
b6
b9
b12
b13
b14
y15
b15
b5
b4
y2

**S49q**

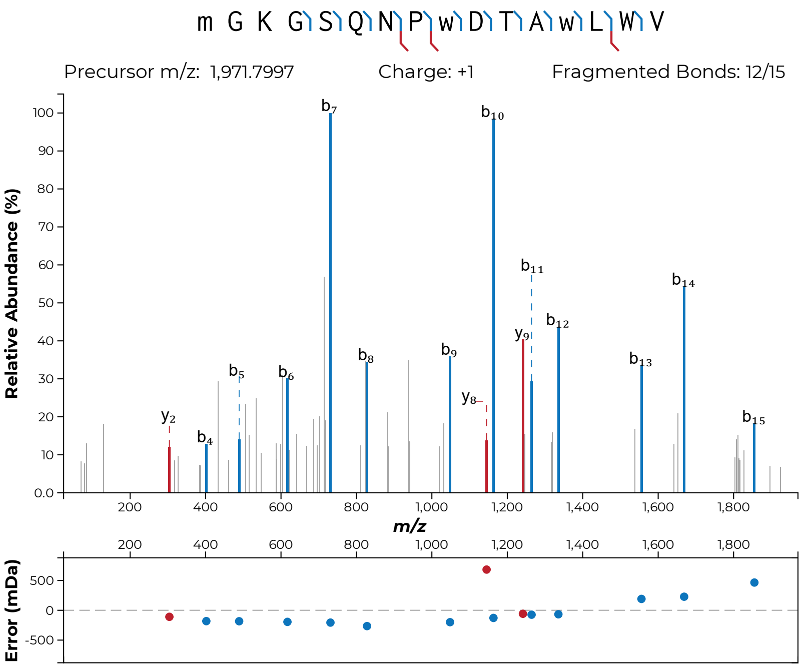
m G K G S Q N P w D T A w L W V
Precursor m/z: 1,971.7997
Charge: +1
Fragmented Bonds: 12/15
Relative Abundance (%)
m/z
Error (mDa)

**S49r**

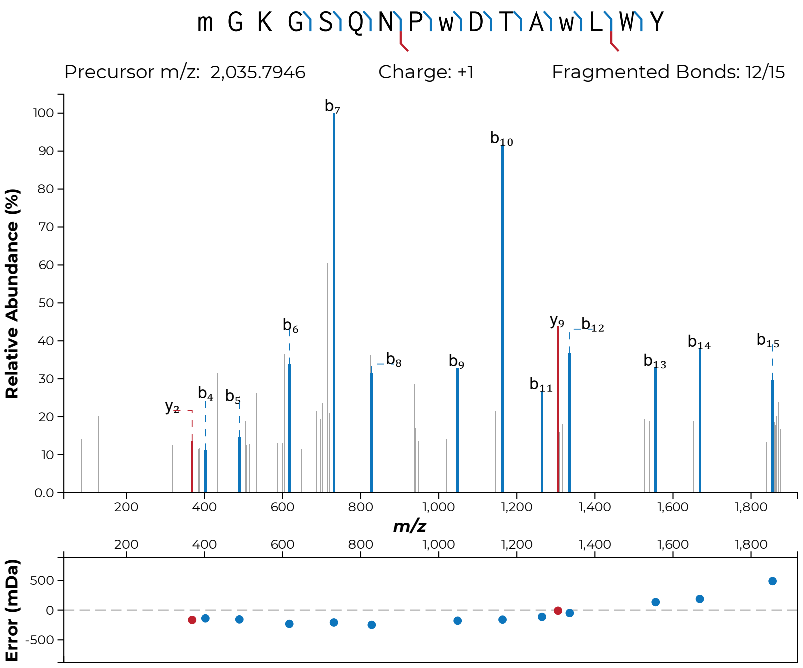
m G K G S Q N P w D T A w L W Y
Precursor m/z: 2,035.7946
Charge: +1
Fragmented Bonds: 12/15
Relative Abundance (%)
m/z
Error (mDa)
y2
b4
b5
b6
b7
b8
b9
b10
b11
y9
b12
b13
b14
b15

**S49s**

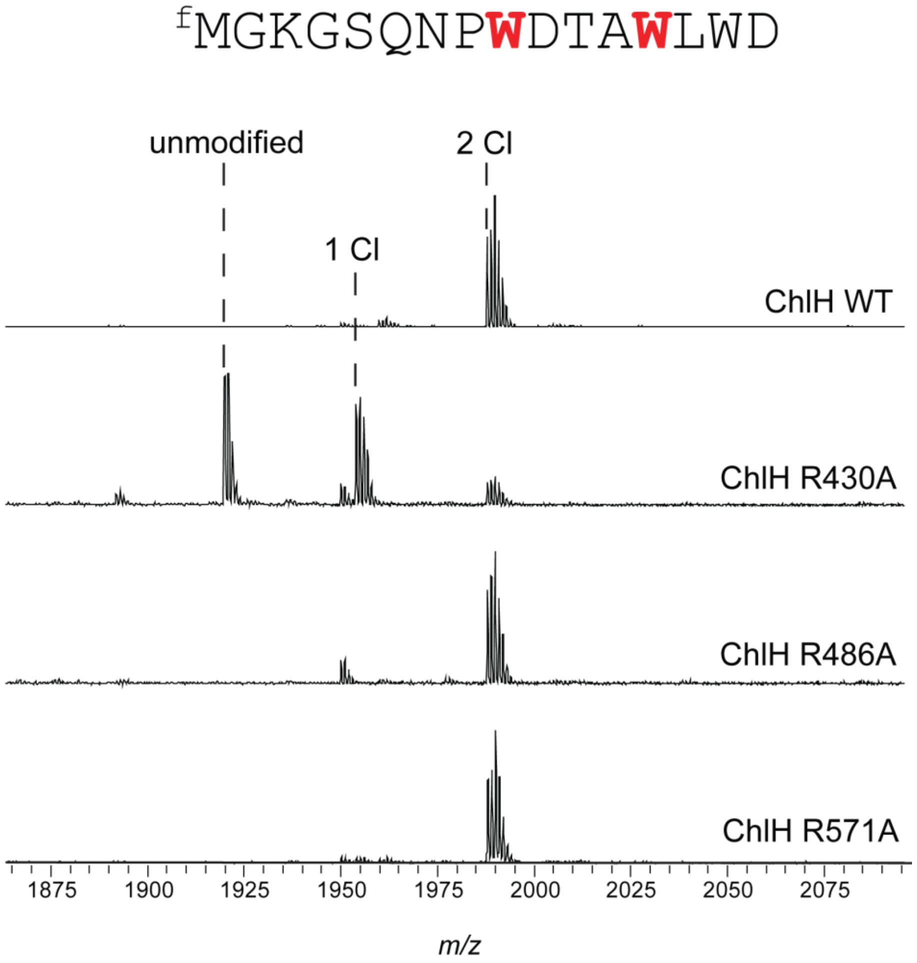


**Figure S50: ChlH Arg variant activity on ChlA$_{PE}$.** Wild-type ChlH and 3 site variants were reacted with ChlA$_{PE}$. Expected m/z: unmodified, 1920; 1 Cl, 1954; 2 Cl, 1988. ChlH R430A is significantly less active, while the R486A and 571A variants retain robust activity.

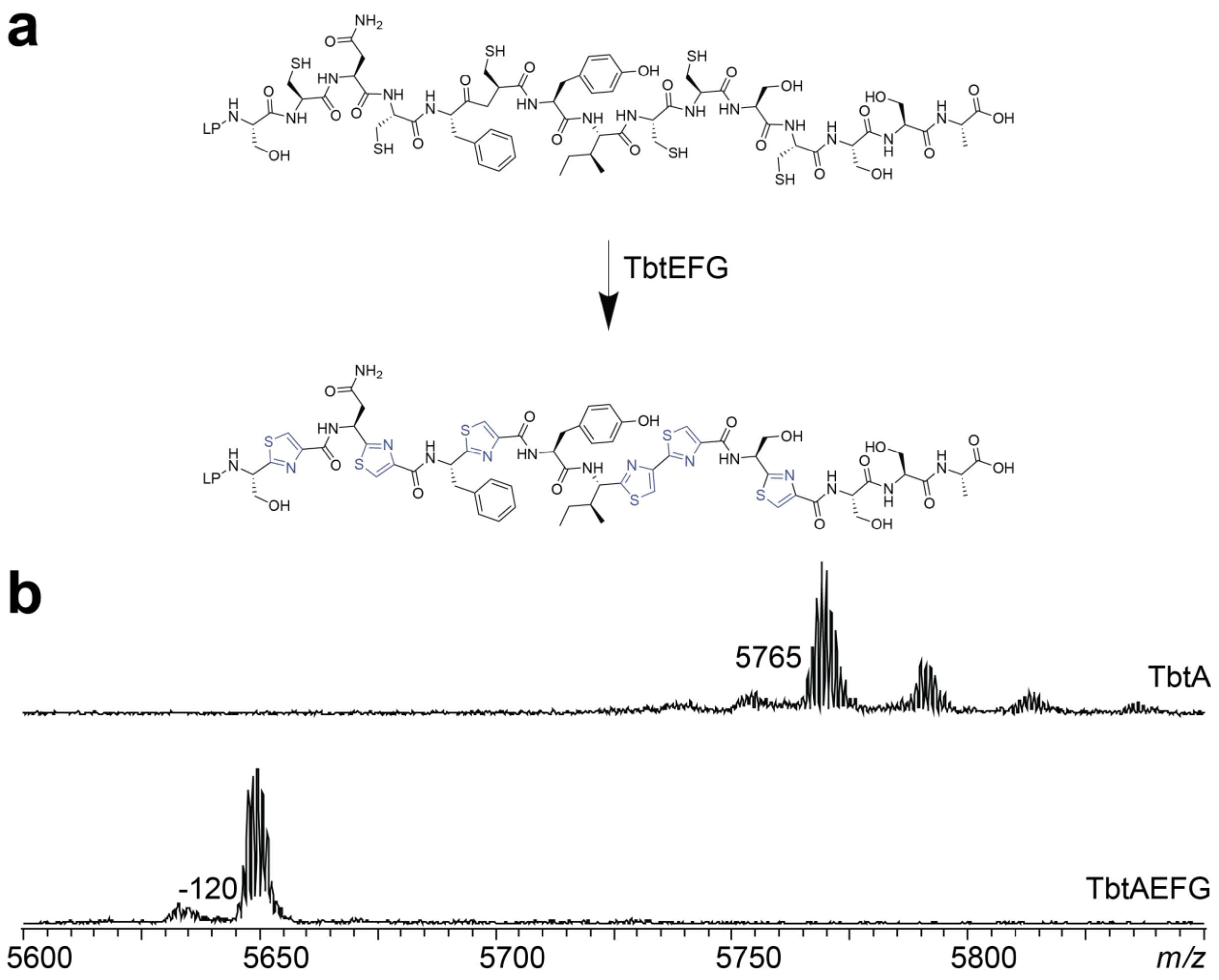


**Figure S51: Assessment of catalytic activity from MBP-TbtG.** The YcaO family member TbtG is a Cys-specific cyclodehydratase that forms a heterotrimeric synthetase complex responsible for peptidic thiazole formation.[4,16] TbtF is RRE-containing, directly engaging the leader peptide (LP) of the TbtA, and delivering this substrate peptide to TbtG, which forms six thiazoline heterocycles from the six Cys residues of TbtA in an ATP-dependent fashion. TbtE is a flavin-dependent thiazoline dehydrogenase that yields a six thiazole-modified product. (a) Reaction of TbtA with TbtE/F/G would generate six thiazoles if each protein were catalytically active. (bt) MALDI-TOF-MS spectrum depicting the formation of six thiazoles on TbtA (-20 Da per thiazole for -120 Da mass loss). This reaction was run using the conditions previously described.[4]

```
ChlA: ----GIGSQNPWDTAWLWD
TbtG: -------LGIPWLPVR---
ChlA: GIGSQNPWDTAWLWD----
```

**Figure S52: Alignment highlighting similarities between TbtG-Trp74 tand ChlA.** ChlA and the tryptic fragment of TbtG were manually aligned. Trp74, the modified Trp residue in TbtG, is flanked on the N-terminal side by Pro, as is Trp8 of ChlA, and on the C-terminal side by Leu, as is Trp12 of ChlA. Modified Trp residues are bolded and depicted in red, and the aligned Pro and Leu residues are shown in blue.

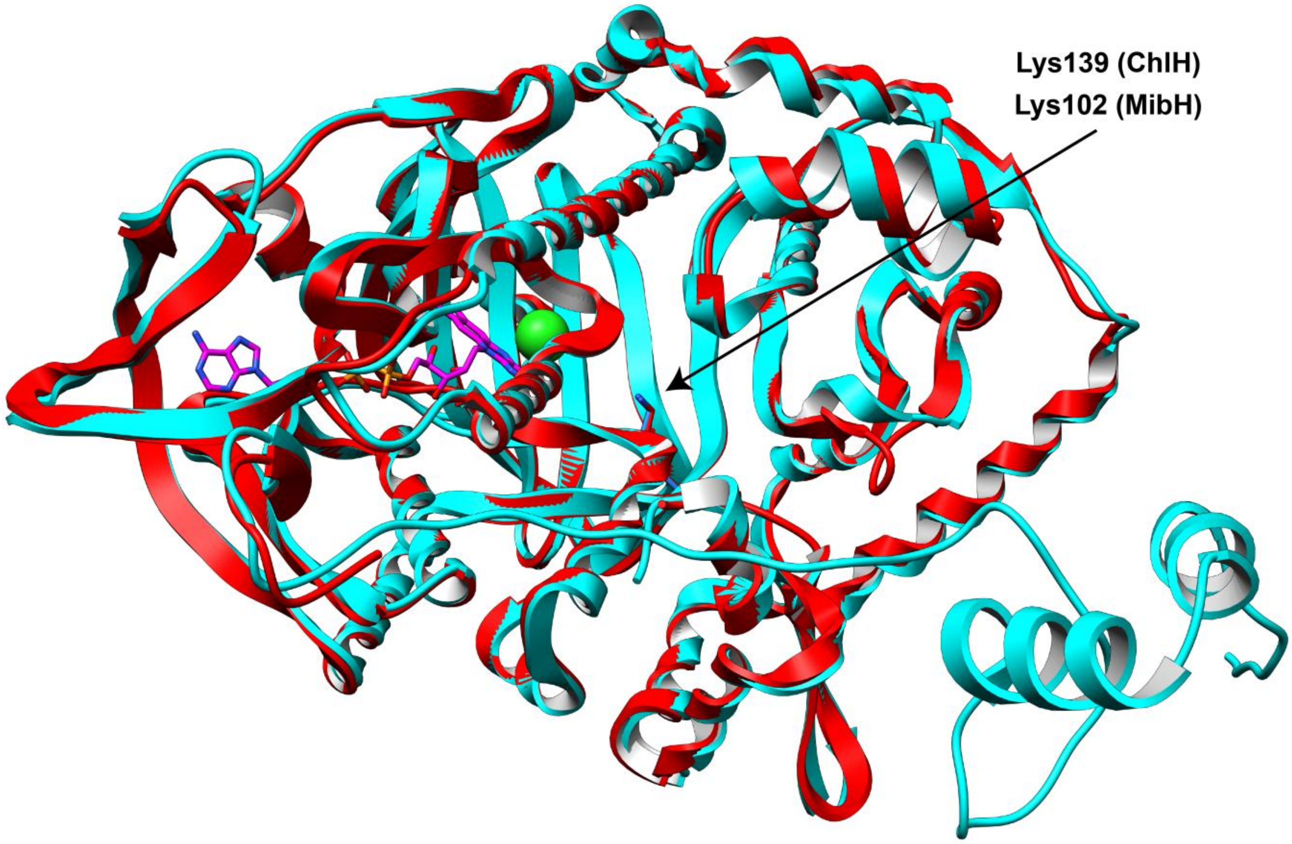


**Figure S53: Alignment of MibH crystal structure and ChlH AlphaFold 3 structure.** An AlphaFold 3[15] structure for ChlH, depicted in cyan, was aligned with the MibH crystal structure (PDB:5UAO),[17] depicted in red. $Cl^-$ and FAD are shown in green and magenta, respectively. The proteins aligned with a global RMSD of 0.507 Å over 2885 atoms. This figure was generated using Chimera.[18]

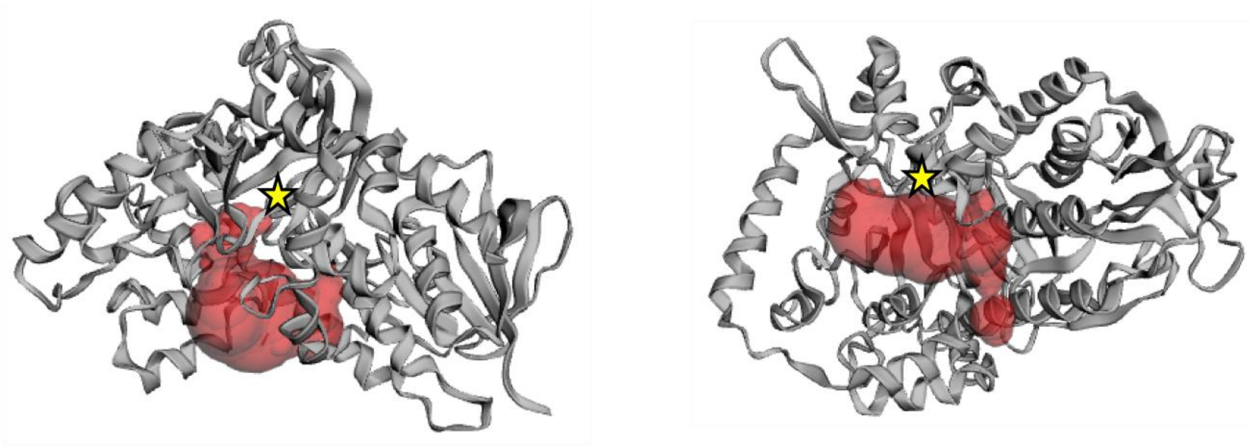

MibH
Surface Area: 553.9 Å$^2$
Volume: 652.7 Å$^3$

ChlH$_{Δ1\text{-}55}$ (1)
Surface Area: 517.9 Å$^2$
Volume: 513.1 Å$^3$

**Figure S54: Active site cavity measurement for ChlH AlphaFold 3 structure.** CASTpFold was used to measure the FDH's active site surface areas and volumes. The crystal structure of MibH (PDB: 5UAO)[17] was used, while an AlphaFold3 predicted structure of ChlH was used. For ChlH$_{Δ1\text{-}55}$, residues 1-55 were excised prior to AlphaFold3 structure prediction. The yellow star denotes the location of the active site Lys in each structure. These data were generated using the CASTpFold web server.[19]

| | ChlH | MibH | SrpI | DarH | ThaI | RebH | PrnA |
|---|---|---|---|---|---|---|---|
| ChlH | 100 | 58 | 21 | 15 | 27 | 27 | 29 |
| MibH | 71 | 100 | 22 | 15 | 27 | 26 | 28 |
| SrpI | 32 | 32 | 100 | 15 | 21 | 23 | 23 |
| DarH | 27 | 27 | 30 | 100 | 16 | 17 | 16 |
| ThaI | 41 | 40 | 33 | 30 | 100 | 63 | 55 |
| RebH | 40 | 40 | 35 | 30 | 77 | 100 | 53 |
| PrnA | 42 | 41 | 36 | 32 | 71 | 70 | 100 |

Amino acid sequence of SrpI:

```
MIQPGSESLRKIAVIGRGTAGSLAAASVTRLHPDADHELHHIYDSRIPVIGVGEGSWPSLVQEVQQLTGLPHETVQQRLKGTRKYG
VAFEGWGRRGRDFTHYFTPQQVSYAYHLSADLLADMLHESSRARHIDAKVLDIARVDGGARVEFEGRAPERYDLVFDARGFPRELD
TDEHIDISFIPTNTAVIRRCPAIVEEAAGPVLQHTYTRAVARPHGWIFVIPLAVHTSYGYIFNRDVTGLDEVESDFDAFLETDGVP
EFEQRAVLRFPNFVHRRIYDGAVARIGNAAAFMEPLEATAIVSAQIQIGMVLKTRLGRSVEHLDRDAPAVNRFLVKNVLRYGLFVG
WHYSCGSRYDSPFWRFARDRTWPRYRSAADPAAVDCNALGEFDEMIRLLHQPVIDQGDWHRMCAVPLTSYAQMSQGLGC
```

**Figure S55: Sequence identity/similarity matrix for flavin-dependent Trp halogenases.** The percent sequence identity (above diagonal) and similarity (below diagonal) between two Trp-modifying FDHs are given. Accession identifiers for each protein are: ChlH (WP_090100303.1), MibH (WP_036325874.1), DarH (WP_063364335.1), ThaI (ABK79936.1), RebH (CAC93722.1), and PrnA (AAB97504.1). As SrpI does not have an NCBI or UniProt accession code, the amino acid sequence has been provided above.

### Supplemental References


(1) Brademan, D. R.; Riley, N. M.; Kwiecien, N. W.; Coon, J. J. Interactive Peptide Spectral Annotator: A Versatile Web-Based Tool for Proteomic Applications*. *Mol. Cell. Proteomics* **2019**, *18* (8, Supplement 1), S193–S201. https://doi.org/10.1074/mcp.TIR118.001209.

(2) Nishihara, K.; Kanemori, M.; Kitagawa, M.; Yanagi, H.; Yura, T. Chaperone Coexpression Plasmids: Differential and Synergistic Roles of DnaK-DnaJ-GrpE and GroEL-GroES in Assisting Folding of an Allergen of Japanese Cedar Pollen, Cryj2, inEscherichia Coli. *Appl. Environ. Microbiol.* **1998**, *64* (5), 1694–1699. https://doi.org/10.1128/AEM.64.5.1694-1699.1998.

(3) Nishihara, K.; Kanemori, M.; Yanagi, H.; Yura, T. Overexpression of Trigger Factor Prevents Aggregation of Recombinant Proteins in Escherichia Coli. *Appl. Environ. Microbiol.* **2000**, *66* (3), 884–889. https://doi.org/10.1128/AEM.66.3.884-889.2000.

(4) Hudson, G. A.; Zhang, Z.; Tietz, J. I.; Mitchell, D. A.; van der Donk, W. A. In Vitro Biosynthesis of the Core Scaffold of the Thiopeptide Thiomuracin. *J. Am. Chem. Soc.* **2015**, *137* (51), 16012–16015. https://doi.org/10.1021/jacs.5b10194.

(5) Kretsch, A. M.; Gadgil, M. G.; DiCaprio, A. J.; Barrett, S. E.; Kille, B. L.; Si, Y.; Zhu, L.; Mitchell, D. A. Peptidase Activation by a Leader Peptide-Bound RiPP Recognition Element. *Biochemistry* **2023**. https://doi.org/10.1021/acs.biochem.2c00700.

(6) Rice, A. J.; Pelton, J. M.; Kramer, N. J.; Catlin, D. S.; Nair, S. K.; Pogorelov, T. V.; Mitchell, D. A.; Bowers, A. A. Enzymatic Pyridine Aromatization during Thiopeptide Biosynthesis. *J. Am. Chem. Soc.* **2022**, *144* (46), 21116–21124. https://doi.org/10.1021/jacs.2c07377.

(7) DiCaprio, A. J.; Firouzbakht, A.; Hudson, G. A.; Mitchell, D. A. Enzymatic Reconstitution and Biosynthetic Investigation of the Lasso Peptide Fusilassin. *J. Am. Chem. Soc.* **2019**, *141* (1), 290–297. https://doi.org/10.1021/jacs.8b09928.

(8) Gaussian 16, Revision A.03, Frisch, M. J.; Trucks, G. W.; Schlegel, H. B.; Scuseria, G. E.; Robb, M. A.; Cheeseman, J. R.; Scalmani, G.; Barone, V.; Petersson, G. A.; Nakatsuji, H.; Li, X.; Caricato, M.; Marenich, A. V.; Bloino, J.; Janesko, B. G.; Gomperts, R.; Mennucci, B.; Hratchian, H. P.; Ortiz, J. V.; Izmaylov, A. F.; Sonnenberg, J. L.; Williams-Young, D.; Ding, F.; Lipparini, F.; Egidi, F.; Goings, J.; Peng, B.; Petrone, A.; Henderson, T.; Ranasinghe, D.; Zakrzewski, V. G.; Gao, J.; Rega, N.; Zheng, G.; Liang, W.; Hada, M.; Ehara, M.; Toyota, K.; Fukuda, R.; Hasegawa, J.; Ishida, M.; Nakajima, T.; Honda, Y.; Kitao, O.; Nakai, H.; Vreven, T.; Throssell, K.; Montgomery, J. A., Jr.; Peralta, J. E.; Ogliaro, F.; Bearpark, M. J.; Heyd, J. J.; Brothers, E. N.; Kudin, K. N.; Staroverov, V. N.; Keith, T. A.; Kobayashi, R.; Normand, J.; Raghavachari, K.; Rendell, A. P.; Burant, J. C.; Iyengar, S. S.; Tomasi, J.; Cossi, M.; Millam, J. M.; Klene, M.; Adamo, C.; Cammi, R.; Ochterski, J. W.; Martin, R. L.; Morokuma, K.; Farkas, O.; Foresman, J. B.; Fox, D. J. Gaussian, Inc., Wallingford CT, 2016.

(9) Wang, J.; Wang, W.; Kollman, P. A.; Case, D. A. Automatic Atom Type and Bond Type Perception in Molecular Mechanical Calculations. *J. Mol. Graph. Model.* **2006**, *25* (2), 247–260. https://doi.org/10.1016/j.jmgm.2005.12.005.

(10) Wang, J.; Wolf, R. M.; Caldwell, J. W.; Kollman, P. A.; Case, D. A. Development and Testing of a General Amber Force Field. *J. Comput. Chem.* **2004**, *25* (9), 1157–1174. https://doi.org/10.1002/jcc.20035.

(11) Russo, J. D.; Zhang, S.; Leung, J. M. G.; Bogetti, A. T.; Thompson, J. P.; DeGrave, A. J.; Torrillo, P. A.; Pratt, A. J.; Wong, K. F.; Xia, J.; Copperman, J.; Adelman, J. L.; Zwier, M. C.; LeBard, D. N.; Zuckerman, D. M.; Chong, L. T. WESTPA 2.0: High-Performance Upgrades for Weighted Ensemble Simulations and Analysis of Longer-Timescale Applications. *J. Chem. Theory Comput.* **2022**, *18* (2), 638–649. https://doi.org/10.1021/acs.jctc.1c01154.

(12) Harris, L. A.; Saad, H.; Shelton, K. E.; Zhu, L.; Guo, X.; Mitchell, D. A. Tryptophan-Centric Bioinformatics Identifies New Lasso Peptide Modifications. *Biochemistry* **2024**. https://doi.org/10.1021/acs.biochem.4c00035.

(13) Madeira, F.; Madhusoodanan, N.; Lee, J.; Eusebi, A.; Niewielska, A.; Tivey, A. R. N.; Lopez, R.; Butcher, S. The EMBL-EBI Job Dispatcher Sequence Analysis Tools Framework in 2024. *Nucleic Acids Res.* **2024**, *52* (W1), W521–W525. https://doi.org/10.1093/nar/gkae241.

(14) Yeh, E.; Blasiak, L. C.; Koglin, A.; Drennan, C. L.; Walsh, C. T. Chlorination by a Long-Lived Intermediate in the Mechanism of Flavin-Dependent Halogenases,. *Biochemistry* **2007**, *46* (5), 1284–1292. https://doi.org/10.1021/bi0621213.

(15) Abramson, J.; Adler, J.; Dunger, J.; Evans, R.; Green, T.; Pritzel, A.; Ronneberger, O.; Willmore, L.; Ballard, A. J.; Bambrick, J.; Bodenstein, S. W.; Evans, D. A.; Hung, C.-C.; O'Neill, M.; Reiman, D.; Tunyasuvunakool,

K.; Wu, Z.; Žemgulytė, A.; Arvaniti, E.; Beattie, C.; Bertolli, O.; Bridgland, A.; Cherepanov, A.; Congreve, M.; Cowen-Rivers, A. I.; Cowie, A.; Figurnov, M.; Fuchs, F. B.; Gladman, H.; Jain, R.; Khan, Y. A.; Low, C. M. R.; Perlin, K.; Potapenko, A.; Savy, P.; Singh, S.; Stecula, A.; Thillaisundaram, A.; Tong, C.; Yakneen, S.; Zhong, E. D.; Zielinski, M.; Žídek, A.; Bapst, V.; Kohli, P.; Jaderberg, M.; Hassabis, D.; Jumper, J. M. Accurate Structure Prediction of Biomolecular Interactions with AlphaFold 3. *Nature* **2024**, *630* (8016), 493–500. https://doi.org/10.1038/s41586-024-07487-w.

(16) Zhang, Z.; Hudson, G. A.; Mahanta, N.; Tietz, J. I.; van der Donk, W. A.; Mitchell, D. A. Biosynthetic Timing and Substrate Specificity for the Thiopeptide Thiomuracin. *J. Am. Chem. Soc.* **2016**, *138* (48), 15511–15514. https://doi.org/10.1021/jacs.6b08987.

(17) Ortega, M. A.; Cogan, D. P.; Mukherjee, S.; Garg, N.; Li, B.; Thibodeaux, G. N.; Maffioli, S. I.; Donadio, S.; Sosio, M.; Escano, J.; Smith, L.; Nair, S. K.; van der Donk, W. A. Two Flavoenzymes Catalyze the Post-Translational Generation of 5-Chlorotryptophan and 2-Aminovinyl-Cysteine during NAI-107 Biosynthesis. *ACS Chem. Biol.* **2017**, *12* (2), 548–557. https://doi.org/10.1021/acschembio.6b01031.

(18) Pettersen, E. F.; Goddard, T. D.; Huang, C. C.; Couch, G. S.; Greenblatt, D. M.; Meng, E. C.; Ferrin, T. E. UCSF Chimera—A Visualization System for Exploratory Research and Analysis. *J. Comput. Chem.* **2004**, *25* (13), 1605–1612. https://doi.org/10.1002/jcc.20084.

(19) Ye, B.; Tian, W.; Wang, B.; Liang, J. CASTpFold: Computed Atlas of Surface Topography of the Universe of Protein Folds. *Nucleic Acids Res.* **2024**, *52* (W1), W194–W199. https://doi.org/10.1093/nar/gkae415.
